# Identification of miRNAs and their corresponding mRNA targets from chickpea nodules and functional characterization of candidate miRNAs by overexpression in chickpea roots

**DOI:** 10.1101/2020.01.12.903260

**Authors:** Manish Tiwari, Baljinder Singh, Manisha Yadav, Vimal Pandey, Sabhyata Bhatia

## Abstract

- Legumes developed symbiotic associations with rhizobia to meet its nitrogen requirement. The nitrogen fixation takes place in root nodules which involves bacterial colonization, organogenesis and nitrogen fixation.
- One microRNA and four parallel analysis of RNA ends (PARE) libraries were sequenced to unravel the miRNA mediated regulation of symbiosis in chickpea.
- Sequencing of microRNA library identified a set of 91 miRNAs comprising of 84 conserved and 7 novel miRNAs. Additionally, PARE library analysis revealed 564 genes being targeted by 85 miRNAs.
- Phylogenetic analysis of the precursor sequences of the 91 miRNAs clearly indicated a clustering of two distinct miRNAs in the same clade representing a close ancestral precursor.
- Further, biogenesis of miRNAs was predicted using the miRNAs identified from different legume genomes.
- The miRNA reads from the nodule library were also mapped onto bacterial genomes from which bacterial small RNA were predicted.
- The antagonistic expression of some of the miRNA-target pairs was investigated and the negative co-related expression profiling proved the validity of the libraries and the miRNA-target pairs. Four miRNAs were selected based on the antagonistic expression profiling and were ectopically expressed in chickpea roots by hairy root transformation.
- The overexpression lines showed significant change in nodule numbers. The target of miR171f (*NRK*), miR394 (*HP*) and miR1509 (*AK*) are novel ones being reported for the first time. This analysis opens a wide arena for investigation of the novel miRNAs and target pairs, polycistronic miRNAs and the bacterial derived smRNAs predicted in this study.

## Introduction

Most legumes inherently meet their nitrogen requirement through symbiotic association with rhizobia. The symbiosis results in fixation of atmospheric nitrogen to ammonia. Initiation of symbiosis involves secreted flavonoids mediated attraction of rhizobia towards legumes. This attraction leads to activation of nodulation genes in rhizobia and secretion of nod factors which establishes communication between the host and symbiont by nod factor receptors (Tiwari and Bhatia, 2019). The rhizobia enters the root hair cells after this preliminary signal exchange. This interaction activates a signaling cascade in epidermal cells which activates various genes like, CCaMK, NSP2, NIN and ENOD etc. The signaling in epidermal cells further alters the phytohormone balance of auxin and cytokinin and results in corticular cell division involving dedifferentiation and redifferentiation of cells to form a niche where rhizobia resides and fix nitrogen for the plant (Bazin *et al*., 2012). One of the indispensable components of the PTGS include microRNAs (miRNAs) that are a class of small endogenous RNAs, approximately 20-24 nt in length. They negatively regulate the expression of a gene either by mRNA degradation, translational inhibition or DNA methylation (Bartel, 2004; Voinnet, 2009). Some miRNAs also act by producing secondary small interfering RNAs (siRNA and tasiRNA) (Chen *et al*., 2010). miRNAs are generated by the action of RNA polymerase II that transcribes microRNA encoding genes, which undergo further processing by 5’ capping, splicing, and polyadenylation at 3’end to form pri-miRNA. These are further processed/cleaved to produce a precursor RNA (pre-miRNA) with a stem-loop structure, which is the cleaved by DCL1 to give rise to the mature miRNA-miRNA* duplex having a two nucleotide overhang at 3’ end (Finnegan and Matzke, 2003). One of the ways in which the mature miRNAs bring about PTGS is through the small RNA guided cleavage that is mediated using the Argonaute protein that has an endonuclease domain and a RNA binding domain (Song *et al*., 2004). A RISC complex is formed which comprises of mature miRNA and AGO protein. This RISC complex along with the region of complementarity between the miRNA and target mRNA determines whether the cleavage pathway or translational repression of target mRNA has to occur (van den Berg *et al*., 2008).

Nodule development is a tightly regulated molecular dialogue between the host plant and rhizobia. Though several mechanisms exist to govern and modulate the gene cascades during the symbiosis process, regulation mediated through miRNAs has been the focus of attention recently (Hoang *et al*., 2020; Zanetti *et al*., 2020). Studies in model crop *Medicago* revealed that overexpression of miRNA166 and miRNA169 led to reduction of nodule formation by cleavage of their respective targets i.e. class III HD-zip transcription factor and *MtHAP2*.*1* (Boualem *et al*., 2008; Combier *et al*., 2006). Similarly, it has been shown that the GRAS family transcription factor *NSP2* (Nodulation Signaling Pathway 2) which is involved in nod factors signaling is targeted by miRNA171 and ectopic expression of miR171 resulted in a significant decrease in nodule numbers (Hofferek *et al*., 2014).

Recently, due to the emergence of next-generation sequencing technologies and the development of *in-silico* approaches, it has become possible to sequence miRNA libraries and predict conserved and novel miRNAs. Further, deep sequencing has also facilitated the development of an approach for the high-throughput identification of miRNA targets. This approach known as Parallel Analysis of RNA Ends (PARE) analyses the RNA degradome generated through miRNA derived cleavage products (German *et al*., 2009). Several studies reporting the sequencing of root nodule specific microRNA libraries are available in legumes such as *Medicago*, common bean and soybean (Devers *et al*., 2011; Formey *et al*., 2014, 2015; Lelandais-Brière *et al*., 2009; Subramanian *et al*., 2008; Wang *et al*., 2009), however very few studies of the miRNA target identification by PARE library sequencing are available from nodules. Therefore, in this study, deep sequencing of a miRNA library from chickpea nodules was carried out and the conserved and novel miRNAs were predicted. The miRNA targets were also identified and annotated by high-throughput sequencing of the RNA degradome libraries from different stages of nodule development. Based on the combined analysis of this data, miRNAs with nodule specific targets were functionally characterized through ectopic expression using hairy root transformation in chickpea.

## Results

### Sequencing and identification of miRNAs from chickpea root nodule tissue

To gain a comprehensive insight into the landscape of miRNAs involved in root nodulation in chickpea, a miRNA library was generated from pooled nodule tissue at different stages. Sequencing resulted in 21,760,971 small RNA reads. After Q20 filtering, adapter removal, sorting reads with length between 20-24 nt and mapping to rRNA, tRNA, snRNA and snoRNA, a total of 4,445,569 reads resulting in 1,315,289 unique tags were obtained (**Table S1)**. The 24 nt reads were most abundant amongst the raw reads as well as the unique tags followed by 21 nt reads (∼30% and 23% unique tags respectively) (**Figure 1A**). The reads were mapped to the chickpea kabuli genome to extract the 250 nt flanking regions which were folded to predict the secondary structures. Structures fulfilling all the latest criteria for miRNA annotation were chosen for miR prediction using ShortStack and mirDeep-P (Axtell, 2013; Yang and Li, 2018).

**Figure 1.**
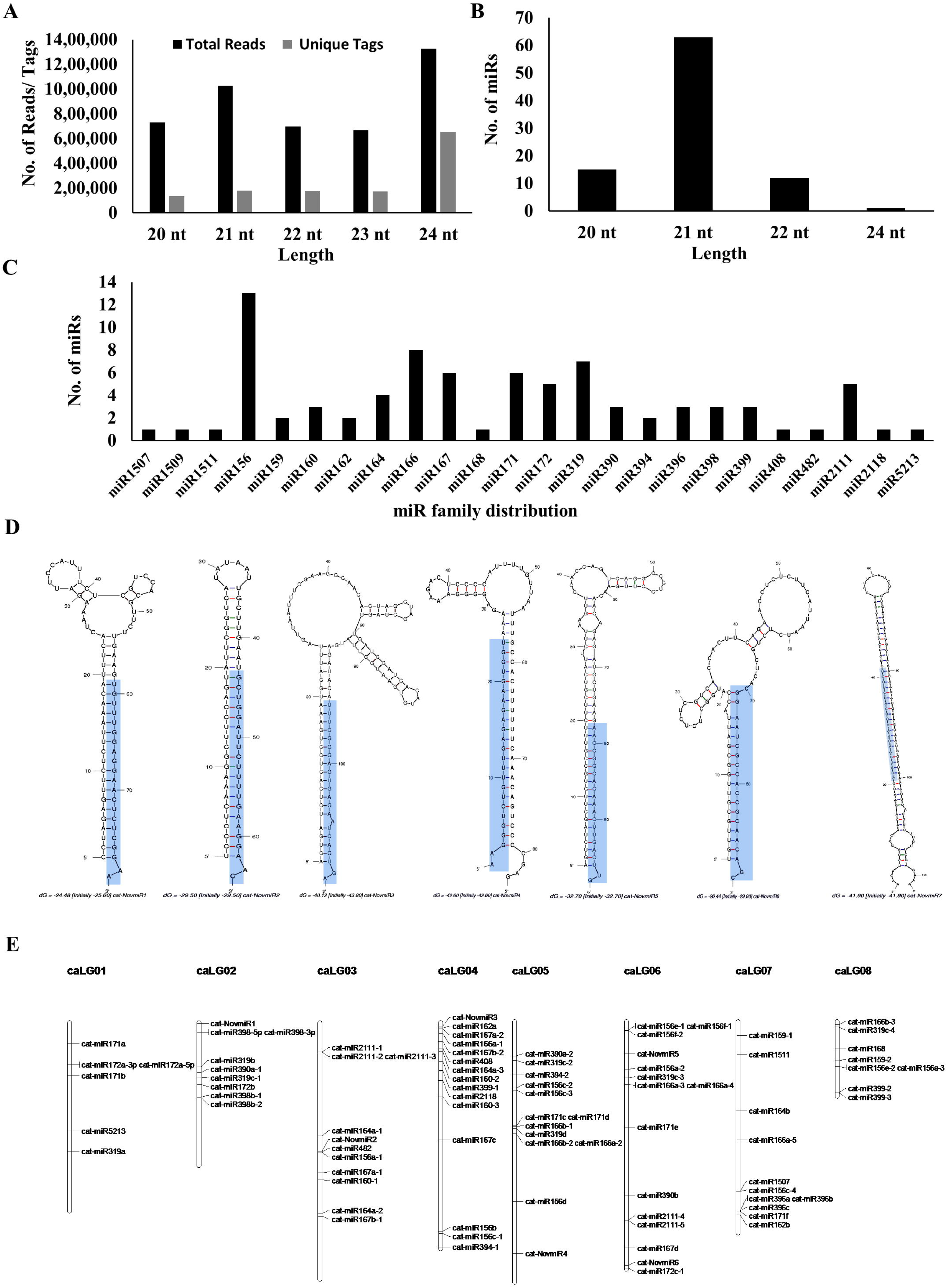
Nodule microRNA library stats, miR length and family distribution and chromosomal mapping. **(A)** Length distribution of reads between 20-24 nucleotides used in analysis. **(B)** Length distribution of miRNA reads into unique tags. **(C)** Length distribution of the identified miRNAs. **(D)** Secondary structures of putative novel miRNAs, the highlighted region represents mature miRNA in hairpin structure (cat-NovmiR1-7). **(E)** The chromosomal mapping of the identified miRNAs on 8 chickpea chromosomes.

A total of 91 miRNAs were identified through a cumulative curation from both mirDeep-P and ShortStack. For prediction of Novel miRNAs, only those passing the stringent criteria for miRNA annotation through ShortStack were retained (**Table S2**). The miRNAs identified were most prevalent in 21 nt category (∼69%), followed by 20 nt (16%) (**Figure 1B**). The first nucleotide at 5’ end of miRNA is crucial for biogenesis and processing of miRNAs. Characteristic association specificity of miRNA’s with AGO1 and AGO4 is provided by the presence of either U or A as the first nucleotide (O’Brien *et al*., 2018). It was found that Uridine was the most abundant nucleotide at the first position of 5’end (∼78%) (**Table S3**). Further, miRNA’s with nucleotide length 20, 21, and 22 all had a greater fraction of U as the first nucleotide (93.3%, 73% and 91.6% respectively) and these particularly associate with AGO1 protein (**Table S3**). In contrast, miRNA with nucleotide length 24 has A as first nucleotide and was reported to associate with AGO4 (**Table S3**). The 24 nt miRNAs are reported to mediate DNA methylation through RNA-directed DNA methylation (RdDM) pathway (Yu, 2015).

Of the 91 miRNAs identified, 84 were conserved and 7 were novel. The 84 conserved miRNAs belonged to 24 miRNA families with maximum representation of miR156 family (∼14%) followed by miR166 (8.8%) (**Figure 1C**). The secondary stem-loop structures of all the miRNAs were analysed carefully following the latest criteria of miRNA annotation (Axtell and Meyers, 2018) (**Figure 1D**). Comparison with the miRBase revealed 18 miRNAs to be legume specific of which 13 were found to be chickpea specific (**Table S2**). Though miR159 and the miR166 were the most abundant miRNAs among the conserved miRNAs, miR319 and miR1511 were most prevalent among the legume specific conserved miRNAs (**Table S2**). Amongst the lowest present miRNAs, miR172 and miR164 were the least in the conserved miRNA pool and miR171 among legume specific conserved ones.

To map the mature miRNAs on chickpea chromosomes the genomic locations of the precursor miRNA were used to position the miRNAs on the eight chickpea chromosomes. Distribution of miRNAs across the chickpea genome revealed that they were located on all 8 chromosomes with maximum miRs being positioned on chromosomes 4 and 6 whereas minimum was located on chromosome 1 (**Figure 1E**).

Nine pairs of polycistronic miRNAs were identified within 10 kb genomic loci region (**Table S2**). These included miR156, miR166, miR171, miR172, miR2111, miR396 and miR398 which represented ∼21% of the total miRNAs which is in accordance with earlier reported loci in *Populus trichocarpa, O. sativa* and Arabidopsis (20%, 19% and 22% respectively) (Merchan *et al*., 2009).

### Phylogenetic analysis of precursor miRNAs in chickpea

To gain an insight into the phylogenetic relationship among the 91 miRNAs identified, a neighbour joining phylogenetic tree was constructed using the precursor sequences of all the miRNAs using MEGA (Kumar *et al*., 2018) (**Figure 2**). The tree revealed distribution of precursors sequences into 22 different clades and interestingly, cat-miR482 did not cluster with any of the miRNA precursor sequences. In most cases, miRNAs of the same family clustered in the same clade. However, there were exceptions and several interesting pairs were identified such as cat-MIR408/156c-2, cat-MIR399/1509, cat-NovMIR4/164b, cat-Nov-MIR6/172c/319/394, cat-MIR162a/166b, cat-MIR162b/171f and cat-MIR1507/2118 etc., which were phylogenetically closer to other miRNA instead of their own family member (**Figure 2**) thereby indicating that they may have evolved from common ancestors.

**Figure 2.**
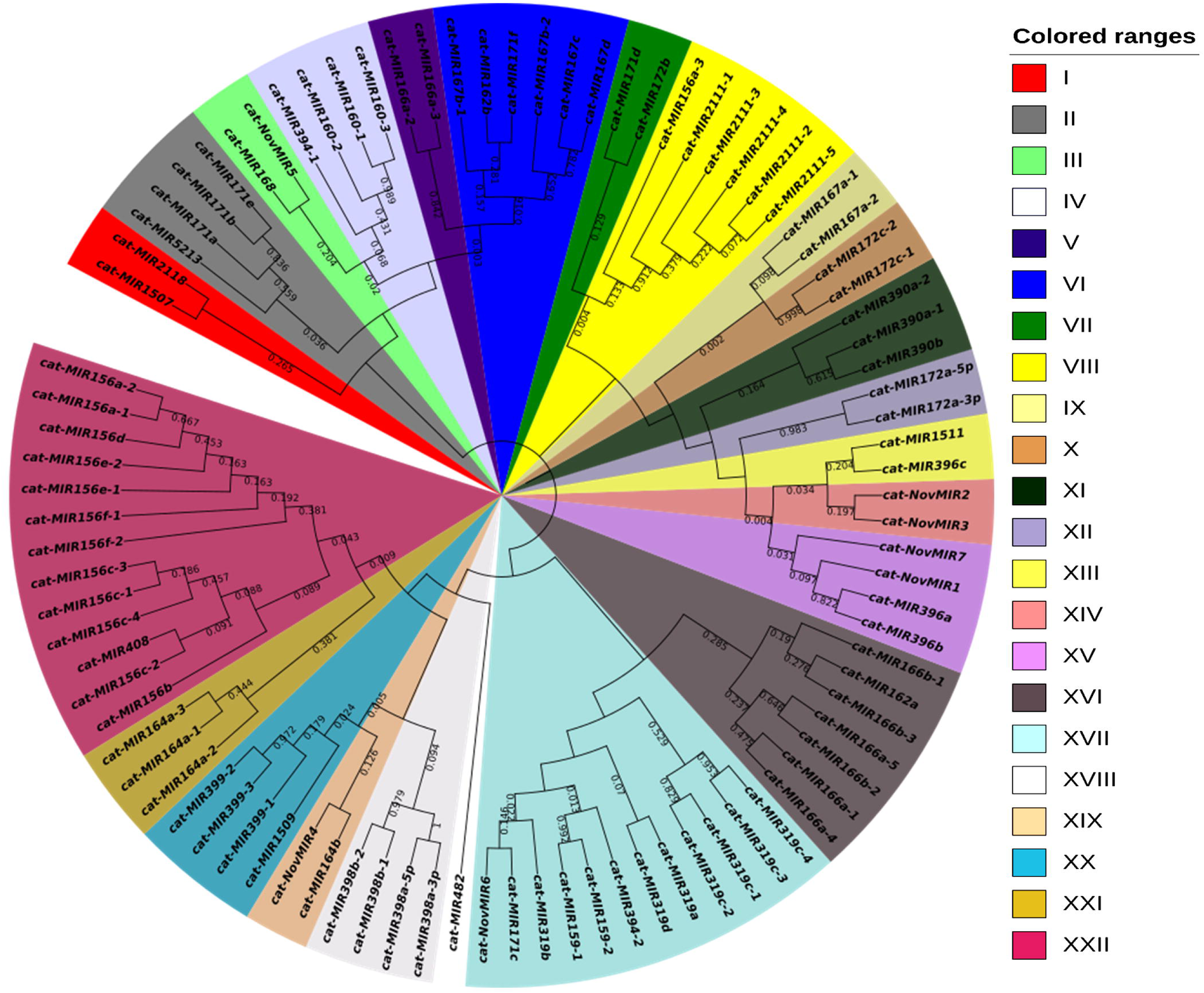
Phylogenetic analysis of the predicted pre-miRNA sequences from chickpea. The phylogenetic tree revealing clustering of 91 miRNA precursor sequences from chickpea in 22 different subclades using MEGA. The tree was constructed using neighbour joining method. The bootstrap values are marked as numbers (Clades I to XXII represented with coloured blocks).

### Prediction of miRNA processing during biogenesis

It is known that during the biogenesis of miRNAs, processing of the precursors occurs in different ways (Bologna *et al*., 2013; Chorostecki *et al*., 2017). Briefly, three types of processing are known to occur during miRNA biogenesis. Firstly, the base to loop miRNA processing involves recognition of ∼15-17 nt conserved stem below the miRNA/miRNA* and first cut by DCL1 below the miRNA/miRNA* region and second cut ∼21 nt away from the first cleavage site (Bologna *et al*., 2013; Chorostecki *et al*., 2017). The second type was loop to base processing that involves ∼15-17 nt conserved region above the miRNA/miRNA* inflicting first cut below the terminal loop region and next cleavage at the base (Bologna *et al*., 2013; Chorostecki *et al*., 2017). The third type of processing involves, a sub-type of loop to base or base to loop processing with multiple cuts instead of two cuts mentioned above. This results in a sequential loop to base or sequential base to loop processing (Bologna *et al*., 2013; Chorostecki *et al*., 2017). The fourth type of miRNA processing during biogenesis includes mixed processing pattern in which there is a short ∼6 nt conserved region above and ∼4 nt conserved region below the miRNA/miRNA* duplex (Chorostecki *et al*., 2017). Therefore, in order to understand the mode of miRNA biogenesis in chickpea and other legumes, the chickpea miRNA reads were mapped to the different plant genomes of legumes, Arabidopsis and rice (see materials and method). Reads mapped with different percentages on to different plant genomes with maximum reads (84%) mapping to chickpea followed by the closest relative *Medicago* (53%) and minimum on the genomes of Arabidopsis (32%) and rice (29%), the distant species (**Figure S1**). miRNAs were predicted, from these plant species (**Table S4**) and the precursor sequences of the similar miRNAs from legumes were aligned using MUSCLE in order to predict the type of processing for miRNA biogenesis. The alignment of precursors from legumes revealed that cat-miR160, cat-miR162b and cat-miR171f undergoes short loop to base processing (**Figure 3A, Figure S2**) and cat-miR167a-d, cat-miR168, cat-miR390a and cat-miR396a-b undergoes short base to loop processing (**Figure 3B, Figure S3**). Interestingly, cat-miR166a displays the conserved ∼4 nt sequences below the miRNA/miRNA* duplex and other conservations were missing. Therefore, it undergoes mixed processing pattern (**Figure 3C**). Additionally, cat-miR319c undergoes sequential loop to base processing involving multiple sequential cuts (**Figure 3D**).

**Figure 3.**
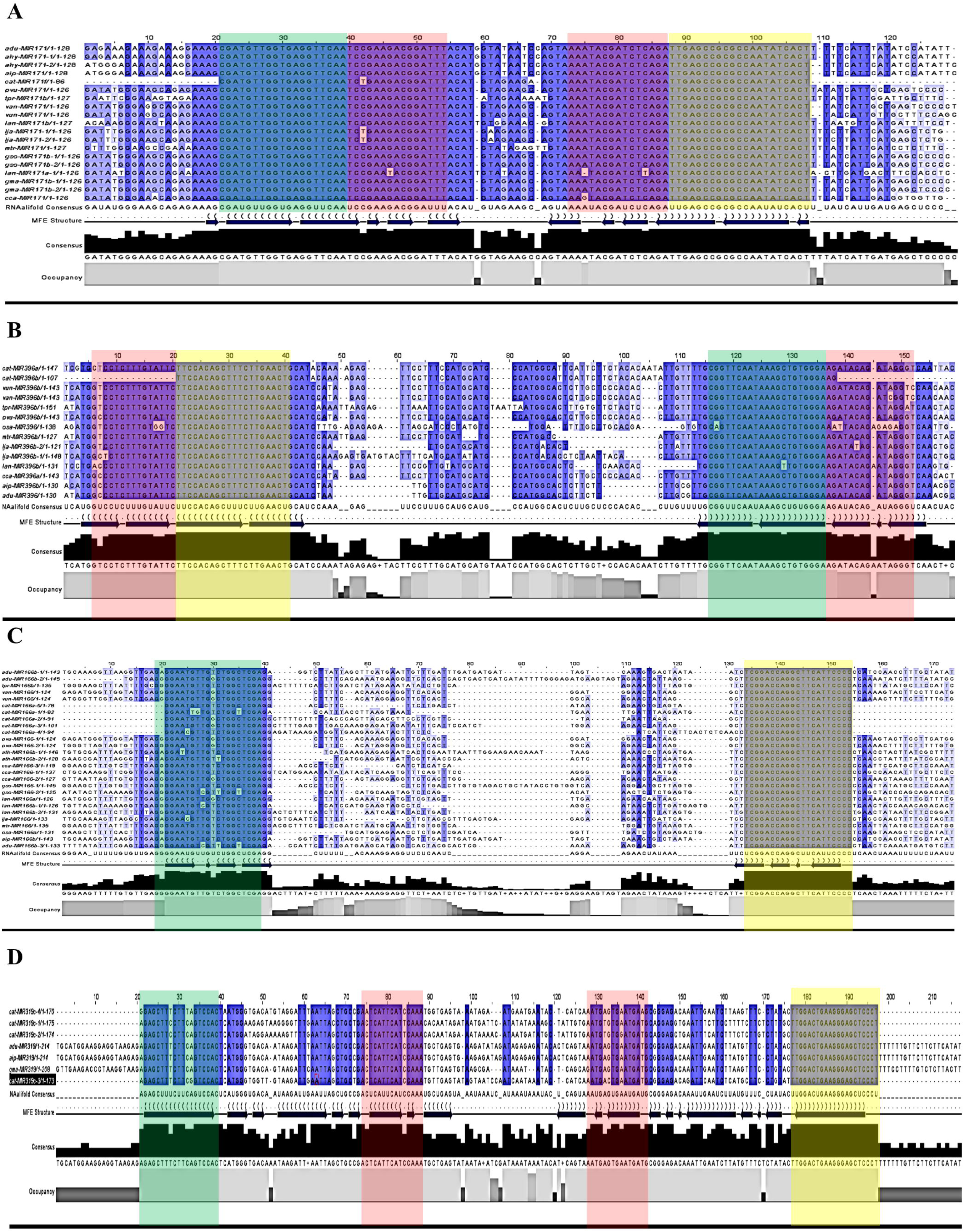
Representation of miRNA precursors processing. **(A)** Alignment of precursor sequences of miR171 representing a short loop-to-base processing. **(B)** Alignment of precursor sequences of miR396 representing a short base-to-loop processing. **(C)** Alignment of precursor sequences of miR166 representing a mixed processing and **(D)** Alignment of precursor sequences of miR319 representing a sequential loop -to-base processing. Yellow marking indicate mature miRNA sequences, green marking indicate miRNA* sequence. Red marking indicate conserved sequences apart from miRNA/miRNA* (*adu- Arachis duranensis, ahy- Arachis hypogea, aip- Arachis ipaensis, ath- Arabidopsis thaliana, cat- Cicer arietinum, cca- Cajanus cajan, gma- Glycine max, gso- glycine soja, lan- Lupinus angustifolius, lja- Lotus japonicas, mtr- Medicago truncatula, osa- Oryza sativa, pvu- Phaseouls vulgaris, tpr- Trifolium pretense, van- Vigna angularis, vun- Vigna unguiculata*)

### Bacterial small RNA prediction

There is growing evidence, such as the recent study in soybean, to show that the rhizobial tRNA derived small RNAs regulate plant genes during nodulation process using the host Argonaute1 machinery in soybean (Ren *et al*., 2019). Following the same line of evidence, the reads obtained from chickpea nodule library were also mapped on bacterial genomes which resulted in maximum reads being mapped to *M. ciceri (CC1192)* genome (13%) and least to *E. coli (strain K12 substr. MG1655)* genome (3%). Similarly, reads from a related study comprising of nodule miRNA from common bean mapped maximally on its host bacterium *Rhizobium tropici (CIAT 899)* (5%) and minimum again on *E. coli* genome (0.4%) (**Figure S2B**) thereby validating the chickpea result. The chickpea miRNA reads, not mapping to the chickpea genome, but mapping onto the *M. ciceri* bacterial genome were used to predict small RNAs from *M. ciceri*. Fifty-three bacterial small RNAs were predicted using mirDeep-P (**Table S5**). Of these small RNAs, 11 precursor sequences of the bacterial small RNAs were PCR amplified from bacterial RNA (**Figure S4A**). Further, qRT-PCR expression analysis of the smRNAs predicted from bacteria was done using RNA obtained from bacterial cells and nodule tissue. The expression analysis revealed very high expression of the smRNA in nodule tissue compared to the bacterial tissue indicating their presence in root nodule (**Figure S4B**).

### *In silico* expression analysis and identification of chickpea nodule specific miRNAs

In order to estimate the expression of miRNAs and to identify nodule specific miRNAs, an earlier study reporting the identification of miRNAs from different chickpea tissues except nodule was used (Jain *et al*., 2014). Reads mapping and *in silico* analysis revealed that of the 91 miRNAs, 74 miRNAs were present in all the tissues and 7 were found to be nodule specific (6 conserved miRNAs and 1 novel) (**Figure 4, Table S6**). 79, 79, 83, 79, 81, 82 and 84 miRNAs were found to be expressed in tissues such as flower-bud, flower, leaf, young-pod, root, stem and shoot respectively (**Figure 4, Table S6**). However, in the nodule tissue, 16 (14 conserved and 2 novel) miRNAs were found to be highly expressing and especially members of miR166 family were most highly expressed (**Figure 4, Table S7**). Interestingly, comparison of root and nodule miRNAs revealed that of the 91 miRNAs, 67 miRNAs were downregulated whereas 24 were upregulated in nodule tissue (**Figure 4, Table S8**). Amongst these, 28 miRNAs were downregulated and only 11 were upregulated in nodule tissue in comparison to root (**Figure 4, Table S8**).

**Figure 4.**
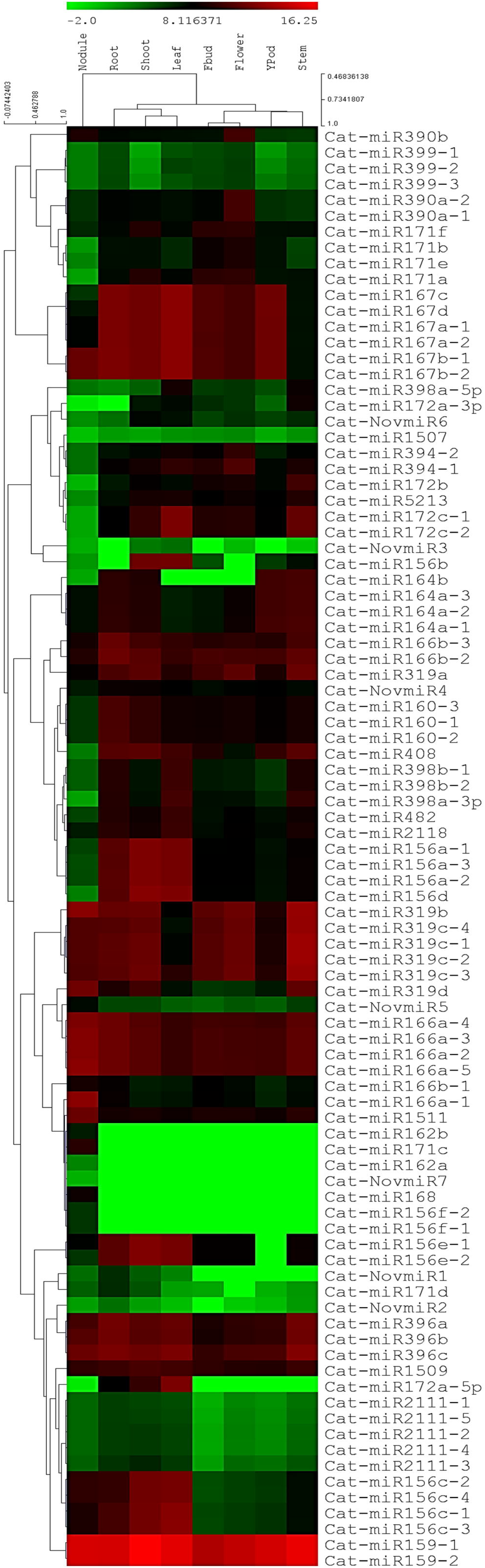
In-silico expression profile of miRNAs in different chickpea tissues. A, heat-map and 2-way hierarchical clustering based on 91 miRNAs that were differentially expressed between nodule and other chickpea tissues (Tissues from left to right: Nodule, Root, Shoot, Leaf, Flower bud (fbud), Young pod (ypod) and Stem).

### Degradome sequencing for target identification

The miRNA prediction data becomes biologically more relevant and significant after identification of the corresponding mRNA target transcripts. Even though, targets can be predicted from available databases such as the psRNA, this does not provide an experimental validation. Therefore, in this study, to simultaneously identify and validate targets at global level, high throughput degradome sequencing of four different libraries from uninfected root and *M. ciceri* infected root nodules at various stages (see materials and method) was performed. After quality filtering and adapters trimming, a total of 5,0,517,249, 4,3,578,672, 4,3,870,856 and 5,2,803,247 reads were obtained from the 4 libraries respectively. The CleaveLand pipeline was used to predict targets of the 91 miRs identified from nodules in this study. The abundance of cleavage tags at the target site determined the category of the cleavage targets and based on this the targets were divided into five categories from 0 to 4 with ‘0’ being the best category (see material and method). More than one raw read was present in categories 0, 1, 2 and 3 whereas category 4 had presence of only single read at the cleavage position **(Figure S5A)**. The targets obtained from each library were sorted based on p-value < 0.05, Category < 3 and Allen score < 5. Target plots (T-plots) representative of cleavage events was also generated (**Figure S5C-F)**. However, the stage-wise distribution showed that 351, 387, 421 and 339 cleavage sites present in 331, 362, 395 and 319 genes respectively were identified from early nodule, late nodule, pooled nodule and root samples that were targeted by 68, 72, 70 and 67 miRNAs (**Figure S5B, Table S9A-D**). These targeted transcripts were part of the non-redundant set of 564 genes in total, which were cleaved by 85 miRNAs out of the 91 identified. The total sites predicted to be cleaved by miRNAs were found in genes from early, mid-late, nodule and root respectively (**Figure S5B, Table S9A-D**). 165 transcripts and 54 miRNAs were present in all the 4 PARE libraries suggesting the strong likelihood of their being significant nodulation related miRNAs-target pairs (**Figure 5A-B**). Further, comparison showed that 245 transcripts were found to be specifically cleaved in nodule libraries in comparison to root and represented nodule specific targets. Similarly, 18 miRNAs were found to be specifically targeting transcripts in nodule tissue only (**Figure 5A-B**). The conserved miR156 family targeted maximum number of transcripts in each library (∼60) and miR1511 had only one target in each library (**Table S9A-D**). Some of the miRNA-target pairs predicted were miR319-*TCP*, miR156-*SPL*, miR164-*NAC*, miR171-*GRAS* and miR172-*AP2* etc., (**Table S9A-D**) which have been previously very well characterized and hence validated the results obtained from the PARE library sequencing in this study.

**Figure 5.**
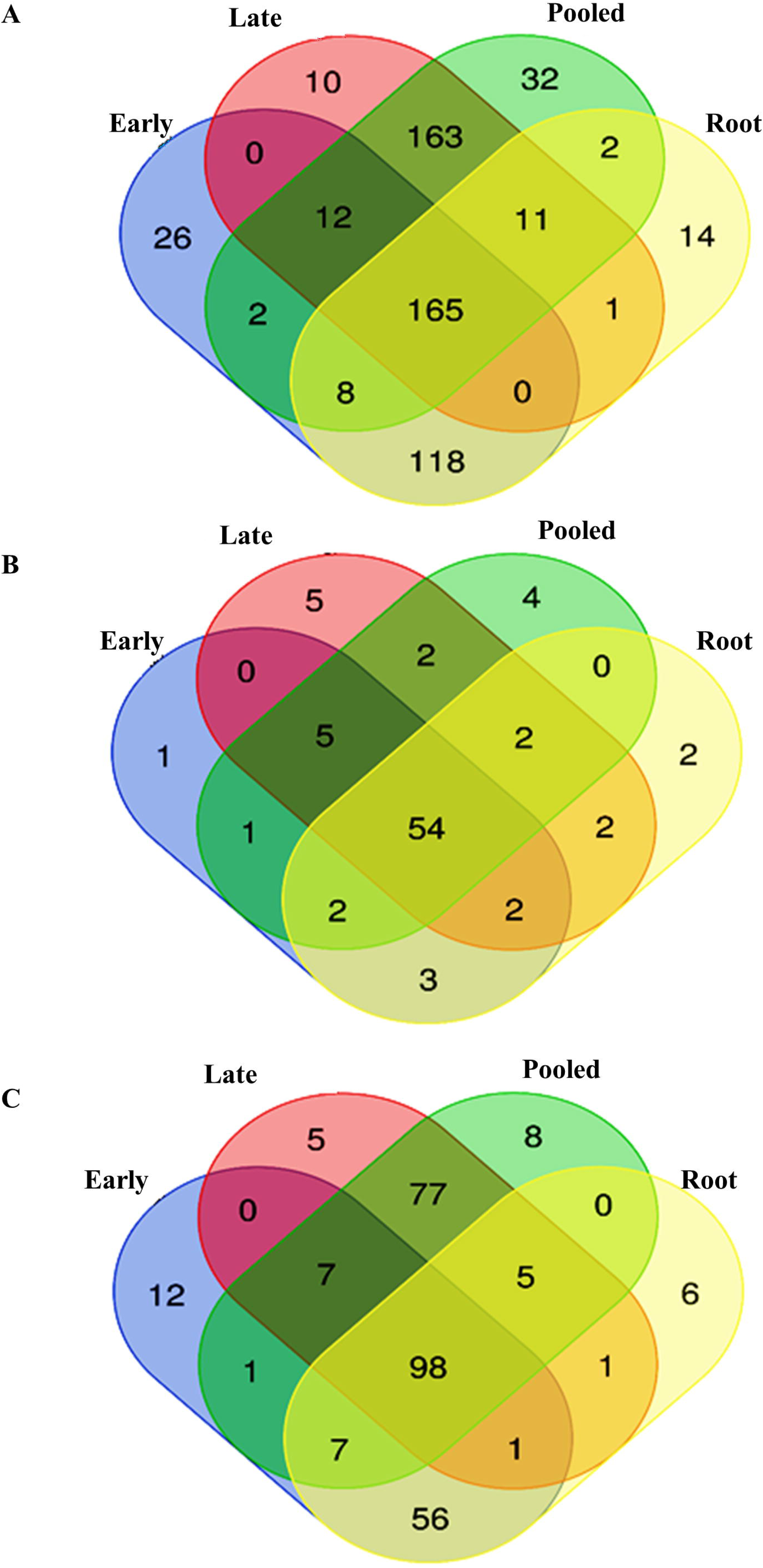
Venn diagram representation of the cleaved targets, the miRNAs found to cleave transcripts in each PARE library and the associated KEGG pathways. **(A)** The targets cleaved by miRNAs in the 4 PARE libraries. (**B**) The miRNAs responsible for cleavage of transcripts identified through PARE libraries **(C)** The differentiation of the KEGG pathways associated with the target transcripts (Early- Early stage of nodule: 0 hpi to 3 dpi, Late-Late stage of nodule: 3 dpi to 28 dpi, Pooled-Pooled nodule: 0 hpi to 28 dpi and Control Root: 0 hpi to 28 dpi)

### Annotation of target transcripts

The target genes were subjected to Blast2GO annotation which classified them into the following important categories: transferases (Aspartate aminotransferase, Histone-lysine N-methyltransferase etc.), kinases (adenylate kinase, PTI1-like tyrosine-protein kinase 6, Nodulation receptor kinase (NRK), CBL-interacting protein kinase 2, etc.), transcription factors and their associated subunits (TCP, ARF, bHLH, NSP, NAC, etc.) resistance protein (TIR-NBS class, LRR and NB-ARC domains-containing disease resistance protein etc.) transporters (ABC transporter, High-affinity nitrate transporter 3.3, Ammonium transporter 1 member 1, Auxin transporter-like protein 6 etc.), cell division cycle (Cell division cycle protein 48 homolog), ribosomal protein (40s and 60s) and other structural and functional proteins (**Table S9A-D**).

The miRNA targets identified using the PARE libraries were subjected to functional annotation by Blast2GO gene ontologies (GO) were assigned. The target genes were mainly predicted to function in cellular components on the basis of their localization in nucleus, cytoplasm, mitocondria, and vesicle during nodule and root development (**Figure S6A**). The miRNA target genes were also predicted to be involved in various biological processes such as protein transport and modification, RNA metabolic process and amide transport (**Figure S6B**). Notably, target genes were also involved in molecular processes such as nucleic acid binding and enzyme activity in both nodule and root tissues. In addition to the binding activity, target genes were also predicted to be involved in hydrolase activity (**Figure S6C**).

The KEGG pathway analysis revealed an involvement of the cleaved targets in 284 pathways of which 98 had common transcripts from all the 4 libraries (**Figure 5C**). Further, genes present in 6 pathways were associated with root tissue only whereas 110 transcripts were involved in pathways associated only with nodulation process (**Figure 5C**). Some of the important pathways include calcium signaling pathway (**Figure S7A**), Nod-like receptor signaling pathway (**Figure S7B**), plant hormone signal transduction (**Figure S7C**) and plant pathogen interaction (**Figure S7D**).

### Expression profiling of conserved and novel miRs and their corresponding targets

In order to estimate the abundance of miRNA transcripts across chickpea tissues, (nodule, root, seed, leaf and flower) a stem-loop qRT-PCR was performed. Expression analysis was carried out for 8 conserved miRNAs (having indicative role in nodule development) and the 7 novel miRNAs. The expression profile revealed that cat-miR1509, cat-miR398a-5p, cat-NovmiR1, cat-NovmiR2, cat-NovmiR4 and cat-NovmiR5 showed fairly high expression in nodule tissue (**Figure 6A**).

**Figure 6.**
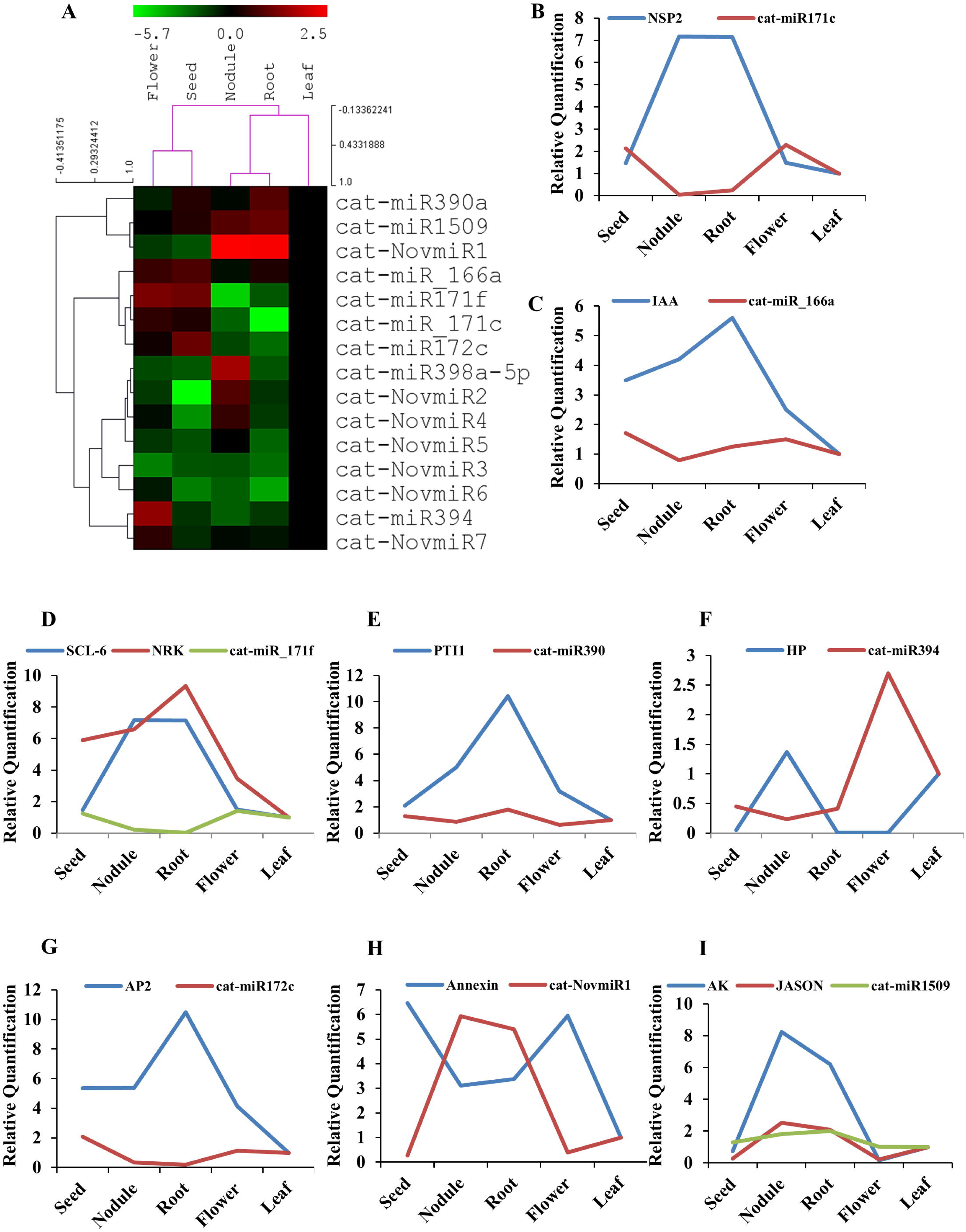
qRT-PCR based expression profiling. **(A)** Heat map illustration of the qRT-PCR of the 15 miRNAs representing the expression profiling of cat-miRs in 5 different tissues (Flower, Seed, Nodule, Root and Leaf). (**B-I**) qRT-PCR based expression analysis of miRNA-Target pair in 5 different tissues of chickpea. **(B)** cat-miR171c-Nodulation signalling pathway 2 (NSP2). **(C)** cat-miR166a- auxin-responsive protein IAA8-like (IAA). **(D)** cat-miR171f-scarecrow-like (SCL) transcription factors (SCL6) and Nodulation receptor kinase (NRK). **(E)** cat-miR390- Pti1-like kinase (PTI1). **(F)** cat-miR394- Histidine phosphotransferase (HP). **(G)** cat-miR172c- *APETALA2*-*like* (AP2). **(H)** cat-NovmiR1- Annexin. **(I)** cat-miR1509-Adenylate kinase (AK) and Jason.

The miRNAs are known to target genes for degradation and hence are expected to exhibit expression patterns that are antagonistic to the expression of their target genes. Therefore, to validate the targets predicted from PARE libraries and the negatively correlated expression patterns of the miRNAs, qRT-PCR was employed. Results revealed that the abundant expression of cat-miR166a, cat-miR171c, cat-miR172c, cat-miR390, cat-miR394 and cat-NovmiR1 was inversely correlated to the expression of targets *IAA, NSP2, AP2, PTI1, HP1* and *Annexin* respectively. Additionally, expression of miR171f and its targets *SCL-6* and *NRK*, were found to be antagonistic. The legume specific miR1509 target its transcripts of *AK* and *Jason* also exhibited expressions that were inversely correlated. (**Figure 6B-I**)

### Ectopic overexpression of cat-miR171, cat-miR172, cat-miR394 and cat-miR1509 by hairy root transformation

The NGS sequencing and analysis of miRNA and PARE libraries revealed several important miRNA-mRNA target pairs. However, to clearly establish the functional role of miRNAs in regulating nodulation, four miRNAs, cat-miRs171f, cat-miR172c, cat-miR394 and cat-miR1509 were selected based on the significance of their target genes (SCL-6/NRK, AP2, HP and adenylate kinase respectively). Of these SCL-6 and AP2 were known targets whereas NRK, HP and AK were novel targets predicted in this study. The 4 pre-miRNAs were cloned and transformed into chickpea roots through *A. rhizogenes* based on hairy root transformation. The ectopically overexpressing lines of 4 miRNAs were obtained as shown in (**Figure 7A**). The effect of overexpression of the miRNAs on the root and nodule phenotype as well as on the expression levels of the corresponding targets were analysed in the overexpression lines (**Figure 7A-F**). The overexpression lines of cat-miRs171f, displayed more than fivefold increase in cat-miRs171f levels with a concomitant decrease in targets *SCL-6* and *NRK* to 0.67 and 0.55 in comparison to vector control (**Figure 7C**). The effect of the cat-miRs171f overexpression and downregulation of corresponding target resulted in a significant decrease in nodule number to 0.71 times in comparison to vector control (**Figure 7B**).

**Figure 7.**
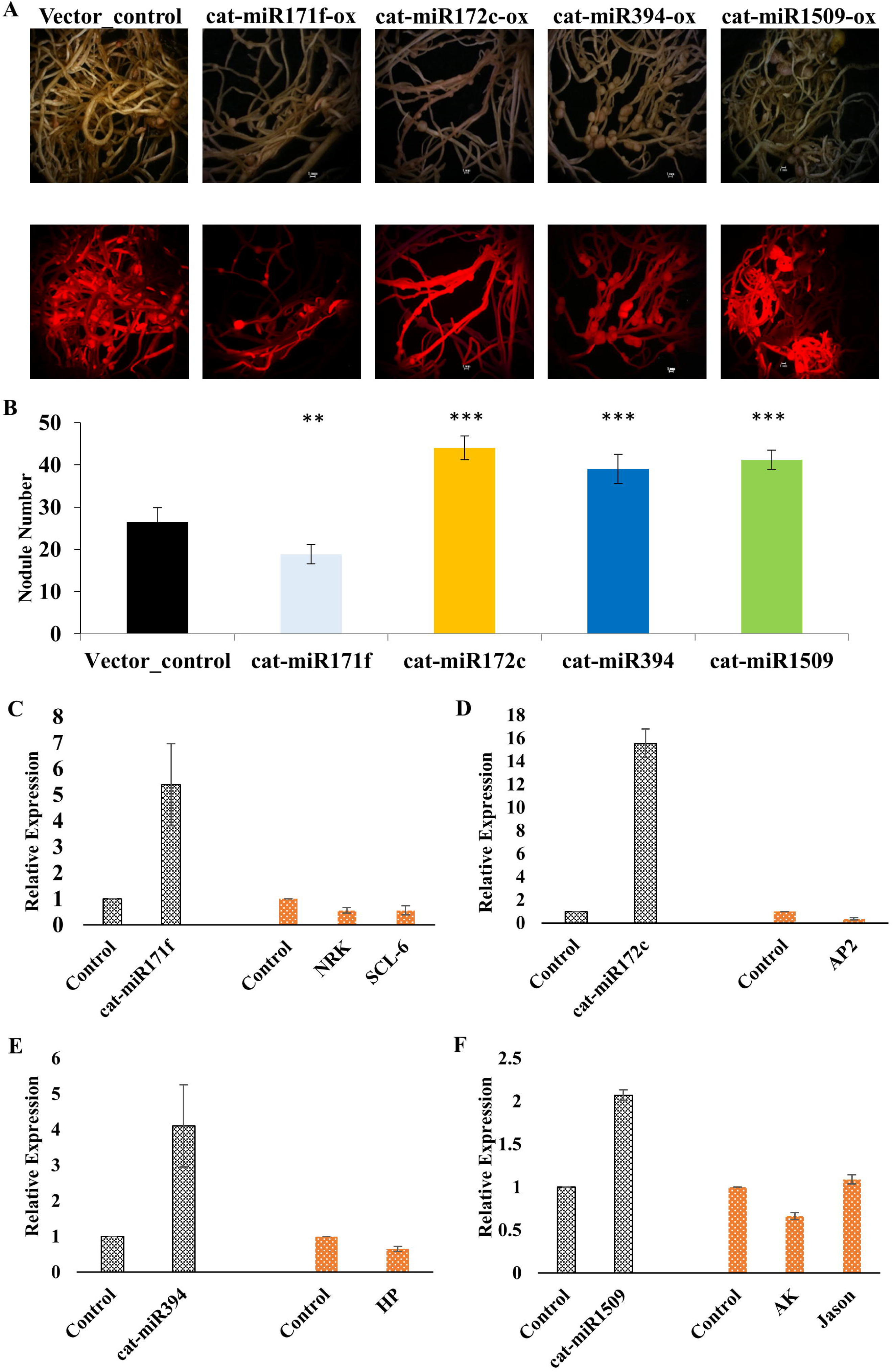
Ecotopic expression of cat-miRs in chickpea hairy roots and qRT-PCR based expression of miRNAs and the corresponding target genes in miRNA overexpression lines. (**A**) dsRED-expressing root nodules in transgenic roots transformed with vector control and respective miRNAs. (**B**) Relative nodule number formed on chickpea hairy roots expressing cat-miR171f, cat-miR172c, cat-miR394 and cat-miR1509 as compared to roots transformed with the empty vector control. (** indicate that the difference is statistically significant at P < 0.05 level and *** indicate that the difference is statistically significant at P < 0.01 level. Error bar represents SE.) **(C)** qRT-PCR analysis of cat-miR171f and its two targets (SCL-6 and NRK) in cat-miR171f overexpression lines. **(D)** qRT-PCR analysis of cat-miR172c and its target AP2 in cat-miR172c overexpression lines. **(E)** qRT-PCR analysis of cat-miR394 and its target HP1 in cat-miR394 overexpression lines. **(F)** qRT-PCR analysis of cat-miR1509 and its target AP2 in cat-miR1509 overexpression lines. Three biological replicates were used. Error bar represents SE. Black bar represent control and miRNA, orange bar represent control and the corresponding targets

Similarly, chickpea roots transformed with overexpression construct of miR172c displayed enhancement of the miRNA upto ∼15-fold in comparison to the vector control with subsequent reduction of its target AP2 transcript by 0.37 times (**Figure 7D**). This ultimately led to an increase in nodule number by 1.67-fold (**Figure 7B**).

The role of miR394 in nodule symbiosis remains largely unexplored till date. Therefore, to investigate its role in chickpea nodulation, ectopic expression of cat-miR394 was carried out. Phenotypic analysis of the overexpression lines showed 1.48 times increase in nodule number (**Figure 7B)**. This could be due to the observed 4-fold increase in expression of cat-miR394 coupled with a 0.65-fold decrease in expression of the target *HP* gene (**Figure 7E)**.

Likewise, the legume specific miRNA, cat-miR1509 was also analysed for its *in-vivo* effect on the nodulation process. The overexpression of cat-miR1509 resulted in a 2.07-fold increase in its expression and a concomitant decrease in the expression of target *AK* transcript by 0.67 times in comparison to the vector control (**Figure 7F**). The overexpression of cat-miR1509 also resulted in an increase in nodule number by 1.56 times of vector control (**Figure 7B)**.

## Discussion

The crop legume is not only valued for its protein content, but for the biological nitrogen fixation (BNF) that it can bring about with the help of rhizobia in its nodules. Reports of spatial and temporal expression of miRNAs are available in chickpea (Jain *et al*., 2014; Khandal *et al*., 2017; Kohli *et al*., 2014). However, miRNA identification and functional characterization has not been performed in crop chickpea nodules. Hence, for identification of miRNAs involved in chickpea nodulation, a high-throughput NGS based sequencing was carried out. A small RNA library from nodules was sequenced and analysed resulting in identification of 84 conserved and 7 novel miRNAs. Moreover, as expected from a chickpea nodule library, legume, chickpea and nodule specific miRNAs were also identified. The first nucleotide of the miRNAs determines the biogenesis and final processivity. Both strands of the miRNA duplex can be loaded into the Argonaute (AGO). The U at the first nucleotide position at the 5’ end of the miRNA duplex decreases the thermodynamic stability making it more favourable to be loaded into AGO as the guide strand (O’Brien *et al*., 2018). In our study 71 miRs had U as the first nucleotide at 5’ end and were hence amenable to DCL1 cleavage and AGO1 association. The 21-nt miRNAs were the most predominant miRNAs in this study however, interestingly 12 miRNAs were also identified with 22-nt length. Such miRNAs are reported to trigger the biogenesis of secondary siRNAs from target mRNAs (Chen *et al*., 2010). Our analysis identified novel polycistronic precursors with non-homologous miRNA (miR156, miR166, miR171, miR172, miR2111, miR396 and miR398). These polycistron harbouring miRNAs target functionally related genes and hence can be used as a tool to regulate gene expression (Merchan *et al*., 2009).

To evaluate the genetic relatedness of the miRNAs, a phylogenetic tree based on the precursor miRNA sequences was constructed. Each subclade contained miRNA members of the same family, however in some cases, very distinct unrelated miRNAs clustered closely together. This close relationship indicated a common ancestor gene that later underwent functional diversification to enable cleavage of different targets. The miRNA reads were mapped onto genomes of legumes and non-legumes. The percentage of reads mapped was in accordance with the evolutionary distance of the organisms from each other indicating that the same pattern of evolution was followed by the small RNAs as other genes. Similarly, the abundance of reads mapped onto bacterial genomes also followed similar pattern indicating a co-evolution of the rhizobia along with the host legumes. Further, the miRNA reads mapping to the *M. ciceri* genome were used to predict bacterial small RNA. Recently it has been shown that rhizobial tRNA derived small RNAs regulate the plant genes governing nodule numbers (Ren *et al*., 2019). Our data of bacterial smRNA prediction and their expression validation opens up several new areas for investigation including the cross-kingdom biogenesis and translocation of small RNAs.

An evolutionary footprint analysis of the precursor miRNA alignments showed that there were conserved regions other than miRNA/miRNA* present in the plant precursors. Their relative positions and length vary among different miRNAs, which determines the processing mechanism of miRNAs. An alignment analysis of precursor sequences from miRNAs identified in chickpea and other legumes revealed regions with conserved sequences beyond the miRNA/miRNA* and these evolutionary footprints were linked to mechanistic processing during miRNA biogenesis. The findings of this investigation revealed that the conserved processing mechanism of the precursor cannot be uncoupled from the miRNA sequence and the evolutionary footprints that determine the biogenesis (Bologna *et al*., 2013; Chorostecki *et al*., 2017).

The *in-silico* expression analysis of the miRNAs indicated a spatio-temporal expression pattern of the miRNAs with several miRNAs recruited to specifically regulate the plant developmental processes. Expression analysis of the cat-miRs revealed 7 to be nodule specific (cat-miR156f-1, cat-miR156f-2, cat-miR162a, cat-miR162b, cat-miR168, cat-miR171c and cat-NovmiR7). The nodule specific cat-miR171c targets *NSP2* gene, which is an important TF involved in nodulation. Overexpression of miR171 leads to a significant decrease in nodule number (Ariel *et al*., 2012; Hofferek *et al*., 2014; Hossain *et al*., 2019). Following this, the other six miRNAs could also regulate expression of key targets regulating nodulation and could be interesting candidates to be characterized for their role during nodulation process. Of the 16 highly expressing miRNAs in nodule tissue, the important member miR166 has been shown to have a polycistonic precursor. In the model legume *Medicago truncatula*, a single transcriptional unit, the MtMIR166a precursor consisted of tandem copies of miR166. The miR166 family regulates a nodule development related TF family, the class-III homeodomain-leucine zipper (HD-ZIP III) genes. *In-planta* overexpression of miR166 causes, miR166 levels to accumulate leading to downregulation of several class-III HD-ZIP genes and a reduction of the number of nodules and lateral roots in the transgenics (Boualem *et al*., 2008). This study established that even though the plant polycistronic miRNA precursors, are rare, they can be processed to regulate the genes and associated pathways. Out study also identified the polycistronic cat-miR166 and its family members that showed high expression in chickpea nodule tissue. Hence it could be suggested that a similar mechanism for processing of polycistronic miRNAs was in place in chickpea nodules also.

To lend biological significance to the study of miRNA identification, their target prediction becomes necessary. In this study analysis of the 4 PARE libraries identified several significant miRNA-target pairs, some of which were earlier well characterized for their role in nodulation. This gave us the confidence to pick up other novel miRNA-target pairs identified in this study as candidates for analysis. Our study showed that 165 targets of 54 miRNAs were found to occur in all the 4 PARE libraries and could serve as high confidence pairs for further characterization. From the venn diagram analysis (**Figure 5A-C**) of the cleaved targets and targeting miRNAs it could be deduced that the 12 cleaved targets were likely to be targeted by the 5 miRNAs regulating 7 functional pathways commonly found in all 4 libraries.

In the past decade, the miRNA mediated regulation has been well established in several plant species including characterization of miRNAs regulating symbiosis at different stages (Boualem et al., 2008; De Luis et al., 2012; Nizampatnam et al., 2015; Nova-Franco et al., 2015). However, functional characterization of a miRNA involved in nodulation in the important crop plant chickpea has never been done. Therefore, in order to functionally decode the regulatory circuitry of miRNA and their targets, 4 miRNA candidates were selected for functional characterization based on the importance of their targets in nodulation.

The miR171 family has been documented to negatively impact the nodulation process and mycorrhizal symbiosis by targeting *NSP2* gene and *SCL-6* (Hofferek *et al*., 2014; Hossain *et al*., 2019; Lauressergues *et al*., 2012; De Luis *et al*., 2012). The ectopic expression of gma-miR171o and gma-miR171q resulted in a significant inhibition of the symbiosis with very few nodules devoid of nitrogen fixation (Hossain *et al*., 2019). In *M. truncatula* it has been reported that cytokinin regulates *NSP2* post-transcriptionally during nodule organogenesis (Ariel *et al*., 2012) through miR171h. The PARE library analysis in our study revealed two targets of cat-miR171f, i.e. *SCL-6* and *NRK*. Through transient *in planta* expression in chickpea, the cleavage of the two targets by cat-miR171f was confirmed. The expression levels of both the targets displayed negative correlation with the cat-miR171f in the overexpression lines. Additionally, the overexpression lines resulted in a significant reduction in nodule numbers. *SCL-6* belongs to the GRAS family of transcription factor which are known to regulate the shoot and root radial patterning (Lim *et al*., 2000). It has been reported in soybean that the miR171 targeting *SCL-6* reduced the nodule numbers but role of *SCL-6* in nodulation is yet to be characterized. Similarly the other target *NRK*, which is a LRR-RLK has been reported to function in Nod-factor perception and downstream transduction system that initiates a signal cascade leading to nodulation (Endre *et al*., 2002). The ortholog of *NRK, DMI2* has been reported to act downstream of Nod-factor receptors and is activated by rhizobial inoculation (Pan *et al*., 2018). Overexpression of full length protein or the kinase domain can induce nodulation even in absence of bacteria (Antolín-Llovera *et al*., 2014). Therefore, reduction in nodule number in the overexpression lines of cat-miR171f were amply supported by previous literature.

The miR172 is well known for its role in flowering time and phase transition process, tuberization of potato and also recently its role in nodule symbiosis has been elucidated (Holt *et al*., 2015; Martin *et al*., 2009; Nova-Franco *et al*., 2015; Wang *et al*., 2014; Yan *et al*., 2013; Zhu and Helliwell, 2011). The miR172 is known to target AP2 family member named *nodule number control1* (NNC1). The miR172c-*NNC1* pair acts as a critical regulatory module of symbiosis which regulates *ENOD40* downstream which leads to nodule initiation and organogenesis (Wang *et al*., 2014). Recently it has also been established that miR172c/*NNC1* module directly controls the transcription of *GmRIC1* and *GmRIC2*, (the cle peptides) which mediate AON activation and attenuation. Based on the importance, cat-miR172-c was selected for overexpression analysis to validate its role in chickpea. Five members of the cat-miR172 miRNA were identified in our study of which miR172c specifically targets the AP2 TF superfamily. The target AP2 transcript was identified through PARE library analysis. Overexpression of cat-miR172c resulted in an increase in nodule numbers which could be explained by the cat-miR172c-NNCI mediated ENOD activation.

The analysis of miRNA-PARE libraries revealed cat-miR394 to target an important cytokinin signaling molecule *histidine phosphotransferase* (HP) with a significant p-value in all the PARE libraries. This target (*HP*) of cat-miR394 was a novel target reported in this study for the first time. HPs are important regulatory molecules which relay the transfer of the phosphate group from the histidine kinases (HKs) to response regulators (RRs) (Hwang *et al*., 2002). Although the mechanism by which HPs regulate the symbiosis process has not been elucidated. The *in-planta* expression studies overexpressing cat-miR394 revealed a significant phenotype (increase in nodule number). The mechanism underlying the cat-miR394-*HP* network mediated nodulation need to be further addressed.

The legume-specific cat-miR1509 targets *AK* and *Jason* as revealed by the degradome analysis. (convert ADP to ATP) and AMP (2ADP ↔ ATP + AMP). The enzyme is known to regulate the level of phosphorus (Carrari *et al*., 2005; Regierer *et al*., 2002). There are no reports available currently to decipher the role of *AK* during nodulation. In present study the cat-miR1509 which target *AK* was overexpressed and resulted in an increase in nodule numbers. This result was indicative of the role of cat-miR1509 mediated cleavage of *AK* and hence a phenotype change in form of increased nodulation. The other target *Jason* did not show an antagonistically correlated expression with the miRNA in the cat-miR1509 overexpression lines. The regulation of nodule phenotype by miR1509 and *AK* needs to be deal with a deeper insight in future.

Overall, the following study presents a holistic view of the miRNAs governing the post-transcriptional gene regulation in chickpea nodulation. A miRNA and 4 PARE libraries were sequenced from the nodule tissues leading to the identification of a repertoire of conserved and the novel miRNAs and their targets implicated to have a role in nodulation. The miRNAs identified were extensively studied for their phylogenetic relationships, prediction of biogenesis from precursors, and their *in-silico* expression patterns. Further, the reads were mapped onto rhizobia genome to predict the bacterial small RNAs with putative cross-kingdom roles that need to be investigated. Based on the significance of the identified miRNAs and targets involved in nodulation, 4 miRNAs were chosen for functional *in-planta* studies. This revealed a novel target, *NRK*, an ortholog of *DMI2* belonging to the LRR-RLK family. The *NRK* was targeted by miR171f and the overexpression of the miR suppressed the nodule number. The miR171f is a cytokinin regulated miRNA and hence this miR and its target pair (a LRR-RLK) explains the effect played by cytokinin on nodulation as proposed earlier. Additionally, a well characterized target of miR172c has also been functionally validated that showed an increase in nodule number as has been previously reported. Two conserved miRNAs (cat-miR394, cat-miR1509) which have not been previously assigned any role during nodulation, nor have their target genes been characterized, were also chosen for investigation of their *in-vivo* role in chickpea nodulation. Their overexpression resulted in an increase in nodule number. The present study provides several novel clues about the conserved and the novel miRNAs, bacterial small RNAs and important nodulation related unexplored target genes. Further, the miRNAs characterized in the present study could be investigated in detail to understand their involvement in regulatory pathways and the mechanism of their action. The genome-wide resources generated in the present study would serve as a foundation to be can be utilized by the research community to enhance the understanding related to modulation of PTGS by miRNAs during symbiosis.

## Experimental procedures

### Tissue generation

Root and nodule tissues were harvested from chickpea (*Cicer aritienum)* cv BGD256. Briefly, seeds were surface sterilized with 0.1% HgCl_2_ and germinated in dark at room temperature. Germinated seeds were transferred to 1% Agar plate. After 4 days of growth, they were inoculated with *Mesorhizobium ciceri* strain TAL620. *M. ciceri* was cultured in Yeast Mannitol Broth and grown at 30 °C for 3 d. Secondary culture with an OD_600_ (1.0) was used for infection. Infected tissues were collected at 1, 3, 6, 12, 24 hpi (hours post infection) and 3, 7, 14, 21 and 28 dpi (days post infection). Root tissue without infection was also collected at the same time points to serve as control tissue.

### RNA extraction for construction of miRNA library and sequencing

Total RNA was isolated from rhizobia infected roots using LiCl precipitation method. The concentration and integrity of RNA were determined. Library preparation was performed following NEBNext® Multiplex Small RNA Library Prep Set from Illumina. The sample was sequenced using Illumina Next Seq 500 Sequencer with 75 SE chemistry.

### microRNA prediction

The reads obtained after sequencing the miRNA library were quality filtered using FastP (Chen *et al*., 2018). Filtered reads were trimmed of the adapter sequences and reads with length between 20 to 24 nt were retained and used for mapping on the genomes (using Bowtie) of chickpea, *M. truncatula*, pigeonpea, soybean, Arabidopsis, rice, *L. japonicus, Arachis hypogea, Arachis duranensis, Arachis ipaensis, Glycine soja, Phaseolus vulgaris, Trifolium pratense, Lupinus angustifolius, Vigna angularis* and *Vigna unguiculata* downloaded from Legume Information System (https://legumeinfo.org/). Sequences that were found to match with those of tRNA, rRNA, snRNAs and snoRNAs were excluded from further analysis. microRNA abundance was normalized into transcripts per million (TPM) for expression analysis.

Moreover, the miRNA library reads were also mapped onto bacterial genomes of *Agrobacterium tumefaciens, Bradyrhizobium japonicum (USDA 6), E. coli (strain K12 substr. MG1655), Mesorhizobium ciceri (biovar biserrulae WSM1271), Mesorhizobium ciceri (CC1192), Sinorhizobium meliloti (1021)* and *Rhizobium tropici (CIAT 899)*. Further, the reads from a miRNA library of common bean nodules were also mapped on the same bacterial genomes (Formey *et al*., 2015).

In order to predict the miRNAs from chickpea nodules 2 pipelines i.e. ShortStack with default parameters and miRDeep-P with core algorithms and plant specific criteria were used (Axtell, 2013; Yang and Li, 2018). For using either of algorithms, reads were mapped to chickpea and other plant genomes and flanking 250 bp window was used for RNA secondary structure prediction using RNAfold. The folded RNA secondary structures which fulfill all the latest criteria for miRNA annotation as described in (Axtell and Meyers, 2018) were chosen for mature miRNA prediction.

### Phylogenetic analysis and prediction of miRNAs biogenesis

The precursor sequences of the chickpea miRNAs identified were aligned using MUSCLE program and a neighbor joining phylogenetic tree was constructed in MEGA using default parameters (Kumar *et al*., 2018).

The precursor sequences of the miRNAs from the genomes of other legumes were mined using the chickpea nodule library reads and were aligned using MUSCLE program and viewed in Jalview (https://www.jalview.org/). The modes of biogenesis of mature miRNA from precursor were predicted based on these alignments.

### Generation of degradome libraries from chickpea nodules

For degradome sequencing, total RNA was isolated at different time intervals ranging from (1 hr to 28 days) from uninfected root (control) and *M. ciceri* infected root (nodule tissue). RNA from root tissues were pooled into one whereas nodule tissues RNAs were pooled to constitute 3 samples i.e. early infection (0 hpi to 3 dpi), late infection (3 dpi to 28 dpi) and a cumulative pooled sample. PolyA RNA was purified using OligodT Dynabeads followed by 5’sRNA adapter ligation. The ligated product was fragmented and purified which was subsequently treated for phosphate removal from 3’ end. After phosphate removal, the 3’sRNA adapter was ligated. Reverse transcription of the RNA library accompanied with PCR and finally sequencing was performed using Illumina GA II sequencing system.

### miRNA target prediction

Sequence data from the PARE libraries of nodule tissues and roots described above was used for target prediction. Reads obtained after degradome sequencing were filtered for removal of low-quality reads and adapter trimming using FastP. Reads less than 10 bp were discarded from further analysis. Targets of microRNA’s were predicted from chickpea cDNA sequences using CleaveLand 4.0 pipeline and the miRNAs predicted in our analysis (Addo-Quaye *et al*., 2009; Parween *et al*., 2015). Reads were mapped to cDNA sequences of chickpea draft2 genome, obtained from a GFF file information. Targets obtained were grouped into 5 categories, namely, 0, 1, 2, 3 and 4. Category 4: Presence of a single raw read. Category 3, 2, 1 and 0 have presence of more than 1 read and in category 3 abundance of the reads is less than both maximum the transcripts and the median, category 2 have abundance of the reads less than the maximum number of reads mapped on transcript but more than the median, category 1 have abundance equal to the maximum which has more than 1 maximum and category 0 have abundance equal to maximum on the transcript having only one maximum (Addo-Quaye *et al*., 2009). Finally, all of the identified potential targets were annotated and characterized using Blast2GO.

### *In silico* differential expression analysis of miRNAs

The expression profile of miRNA across different tissues provide us with an information regarding spatio-temporal post transcriptional gene regulation in chickpea. The digital expression profile of miRNAs was obtained in 9 different tissues of chickpea and compared for differential regulation during plant developmental stages. The miRNA library sequenced in chickpea root, shoot, leaf, flower bud, flower, young pod and stem with SRA number SRP030647 (Jain *et al*., 2014) was used for expression analysis and compared with nodule library performed in present study. Heat-map representing the expression of miRNAs across different libraries was generated by MeV software (http://mev.tm4.org/#/welcome).

### Stemloop and qRT-PCR validation

To validate the miRNAs predicted from the chickpea nodule libraries, stem-loop PCR amplification was performed. Further, expression of the identified targets was detected using qRT-PCR. Briefly, 2µg total RNA was used for cDNA synthesis of microRNA and target using Takara SMARTScribe cDNA synthesis kit and BioRad iScript cDNA synthesis kit as per manufacturer’s protocol respectively. SYBR green master mix was used for Real Time validation. Each PCR was performed with three technical and biological replicates.

### Ectopic expression of cat-miRNAs in chickpea nodules

In order to investigate the functional role of miRNAs, four candidate miRNAs were selected and were overexpressed in chickpea roots by hairy root transformation (Mandal and Sinharoy, 2018). The 200 nt sequences upstream and downstream of the mature miRNAs were extracted from the chickpea kabuli genome (https://legumeinfo.org/). These regions were amplified from the genomic DNA and cloned into the pUb-cGFP-dsRED vector downstream of ubiquitin promoter. The construct containing the transgene was mobilized into *A. rhizogenes* which was then used for hairy root transformation of chickpea roots (Mandal and Sinharoy, 2018). The transformed roots were screened using stereo fluorescence microscope Nikon AZ100 equipped with Nikon digital camera (Nikon digital sight DS-Ri1) for red fluorescence. The chickpea plants with transformed hairy roots overexpressing the desired miRNA precursors were identified by their constitutive DsRed expression and the nodule number was recorded for each of the overexpressing lines in comparison with vector control.

## Data availability statement

All the sequence data sets generated during the current study are available in the NCBI Sequence Read Archive (SRA) under accession PRJNA606204.

## Author’s contributions

Conceived and designed the experiments: SB and MT. Performed the experiments: MT, BS and MY. Analyzed the data: MT, and BS. Wrote the paper: SB and MT. MT, VP and BS performed the computational analysis, MT, BS and MY prepared all the figures and tables. MT, MY, BS and VP performed the chickpea transformation. Critical evaluation, funding resources and all the facilities were arranged by SB.

## Acknowledgements

We kindly acknowledge National Institute of Plant Genome Research (NIPGR) and Department of Biotechnology, Govt. of India (http://www.dbtindia.nic.in). The study was funded by grant (BT/PR3305/AGR/2/816/2011) from Department of Biotechnology, India.

## Supporting information legends

**Supplementary Figures S1-S7**

**Figure S1. Mapping percentage of reads on various plant and bacterial genomes**.

**Figure S2. Representation of miRNA precursors processed in a loop-to-base direction**.

**Figure S3. Representation of miRNA precursors processed in a base-to-loop direction**.

**Figure S4. Validation of bacterial smRNAs**.

**Figure S5. Confirmed microRNA targets using degradome sequencing**.

**Figure S6. Blast2GO based functional annotation of target transcripts**.

**Figure S7A-D. KEGG pathway analysis of targets**.

**Supplementary Tables S1-S9**

**Table S1. The length distribution of sequenced reads of microRNA in library**.

**Table S2. The detailed information about the miRNAs, chromosomal location, start and end position, length, sequence information of mature and precursors miRNAs with their corresponding RPM values from nodule library**.

**Table S3. The detailed information of length distribution of mature miRNAs and their first nucleotide distribution**

**Table S4. The detailed information about the other plant species miRNAs, chromosomal location, start and end position, length, sequence information of mature and precursors miRNAs with their corresponding RPM values from nodule library**

**Table S5. The detailed information about the bacterial small RNAs, chromosomal location, start and end position, length, sequence information of mature and precursors miRNAs with their corresponding RPM values from nodule library**

**Table S6. The distribution of 91 microRNA in different tissues of chickpea**

**Table S7. In silico tissue-wise expression in different tissues of chickpea**

**Table S8. The list of expression of microRNA in nodule tissue in comparison to root**

**Table S9. The information of miRNA, target, annotation and category**

