## Supplementary material for "Identification of miRNAs and their corresponding mRNA targets from chickpea nodules and functional characterization of candidate miRNAs by overexpression in chickpea roots": Figure S1-S7

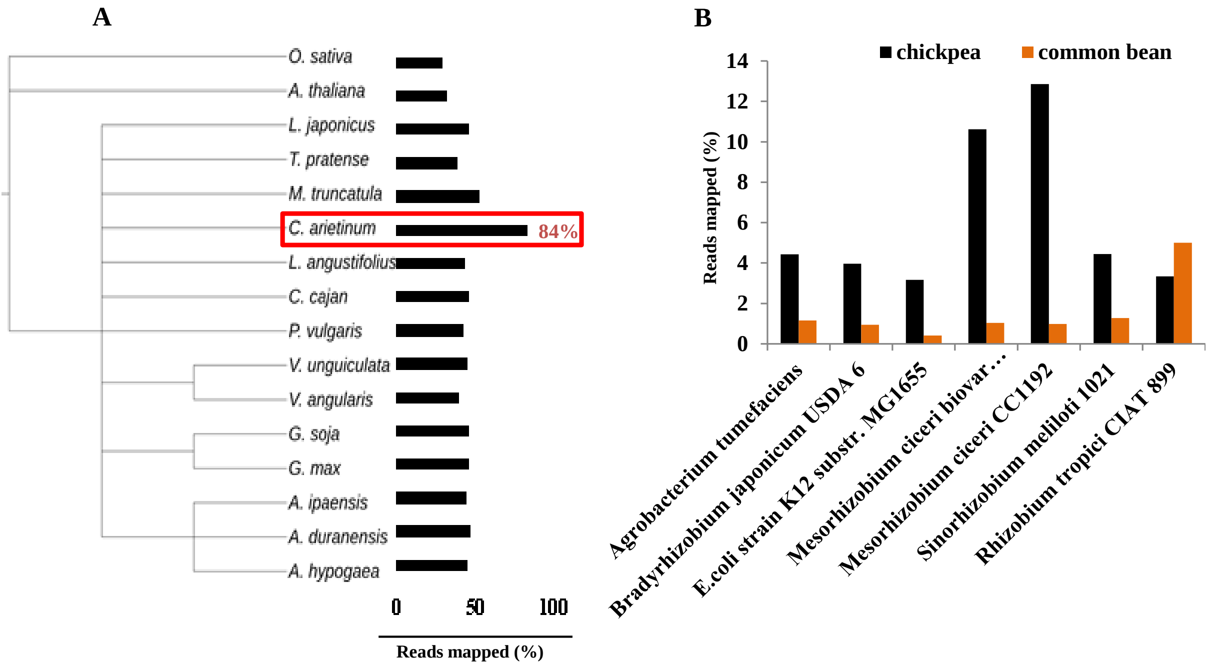


**Figure S1. Mapping percentage of reads on various plant and bacterial genomes.** (**A**) A phylogenetic tree based on the %age of reads mapped on genomes of different plant species (*Oryza sativa, Arabidopsis thaliana, Lotus japonicus, Trifolium pratense, Medicago truncatula, Cicer arietinum, Lupinus angustifolius, Cajanus cajan, Phaseolus vulgaris, Vigna unguiculata, Vigna angularis, Glycine soja, Glycine max, Arachis ipaensis, Arachis duranensis and Arachis hypogea*). (**B**) Bar diagram represent the %age of reads (nodule library chickpea and common bean) mapped on different bacterial genomes including the rhizobia *(Agrobacterium tumefaciens, Bradyrhizobium japonicum- USDA 6, E. coli- K12, Mesorhizobium ciceri- WSM1271 and CC1192, Sinorhizobium melioti- 1021 and Rhizobium tropici- CIAT899).*

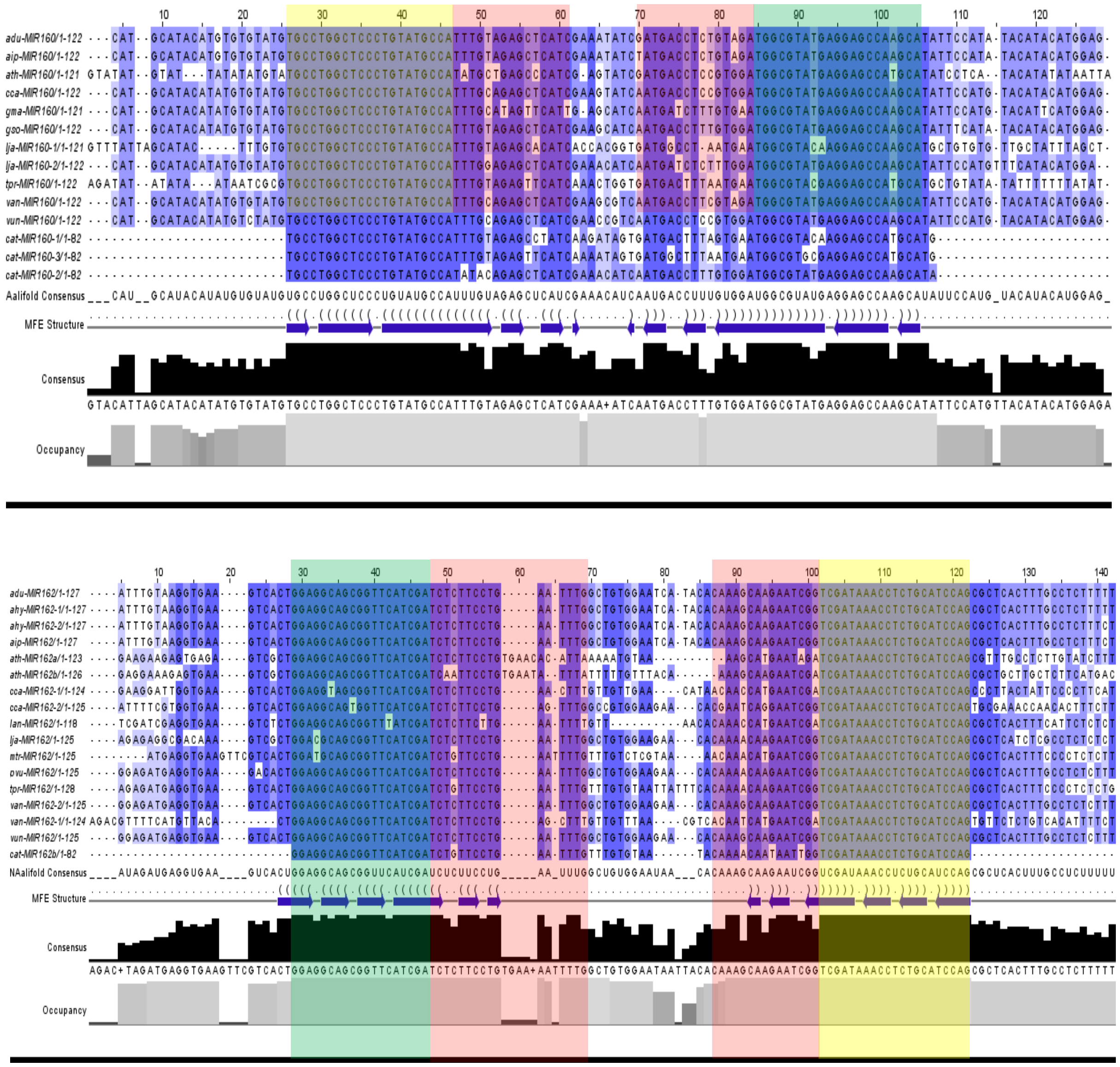

**Figure S2. Representation of miRNA precursors processed in a loop-to-base direction.** Alignment of precursor sequences of miR160, and miR162 representing a short loop-to-base processing. Yellow marking indicate mature miRNA sequences, green marking indicate miRNA* sequence. Red marking indicate conserved sequences above miRNA/miRNA* (*adu- Arachis duranensis, ahy- Arachis hypogea, aip- Arachis ipaensis, ath- Arabidopsis thaliana, cat- Cicer arietinum, cca- Cajanus cajan, gma- Glycine max, gso- glycine soja, lan- Lupinus angustifolius, lja- Lotus japonicas, mtr- Medicago truncatula, osa- Oryza sativa, pvu- Phaseouls vulgaris, tpr- Trifolium pretense, van- Vigna angularis, vun- Vigna unguiculata* )

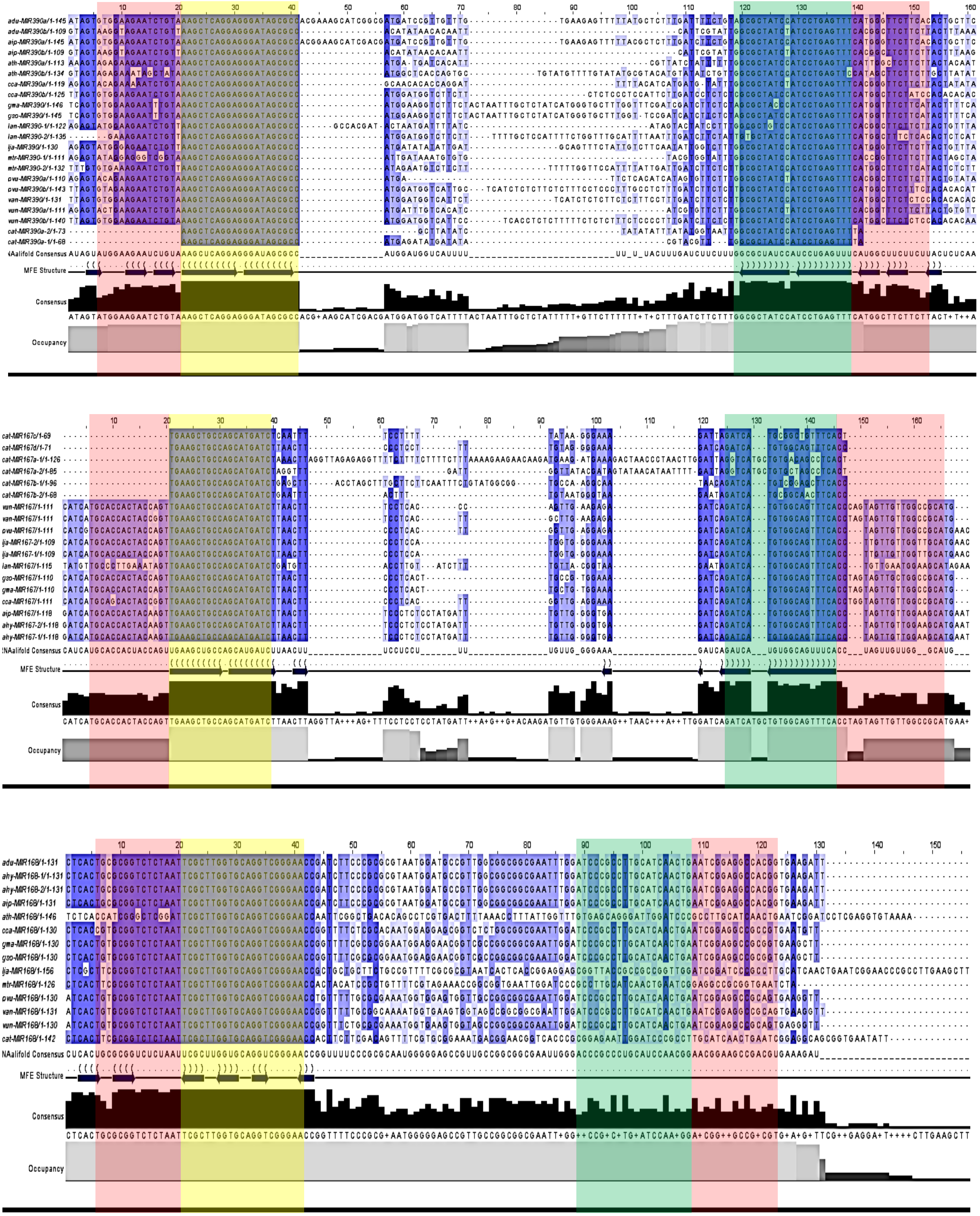


**Figure S3. Representation of miRNA precursors processed in a base-to-loop direction.** Alignment of precursor sequences of miR167, miR168, and miR390 representing a short base-to-loop processing. Yellow marking indicate mature miRNA sequences, green marking indicate miRNA* sequence. Red marking indicate conserved sequences above miRNA/miRNA* (*adu- Arachis duranensis, ahy- Arachis hypogea, aip- Arachis ipaensis, ath- Arabidopsis thaliana, cat- Cicer arietinum, cca- Cajanus cajan, gma- Glycine max, gso- glycine soja, lan- Lupinus angustifolius, lja- Lotus japonicas, mtr- Medicago truncatula, osa- Oryza sativa, pvu- Phaseouls vulgaris, tpr- Trifolium pretense, van- Vigna angularis, vun- Vigna unguiculata* )

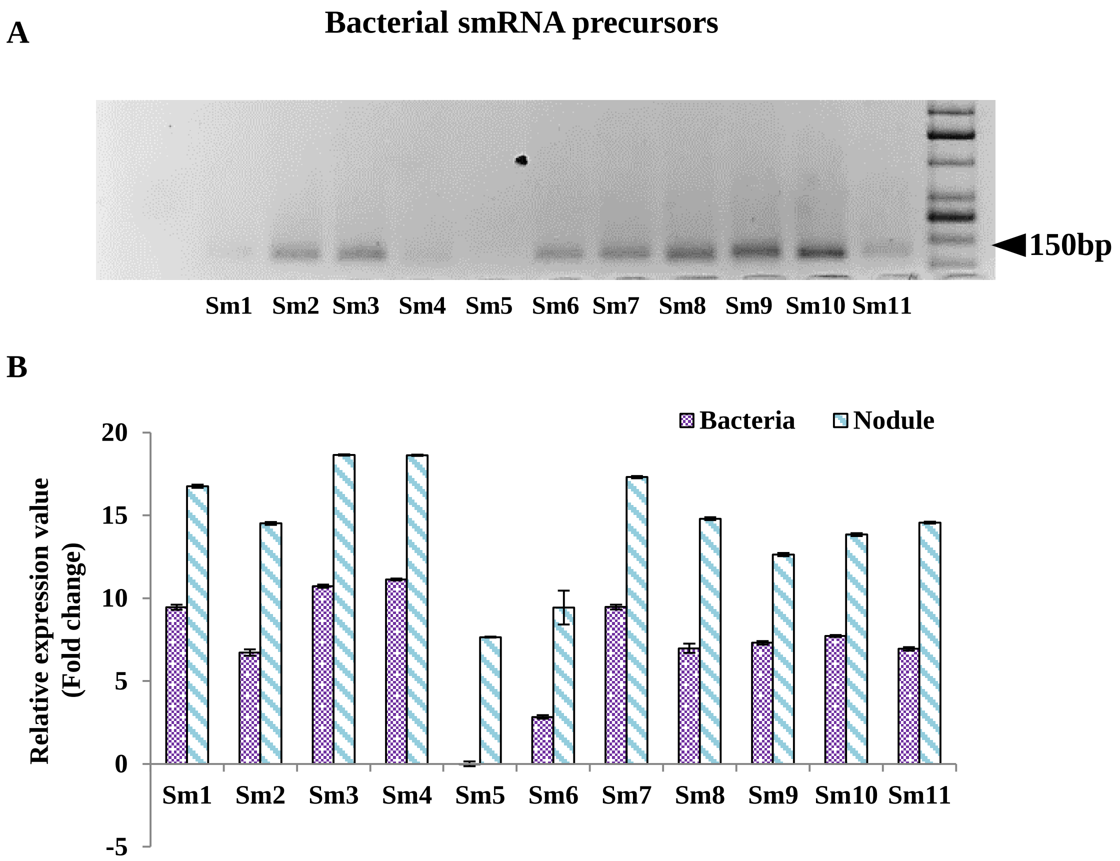


**Figure S4. Validation of bacterial smRNAs. (A)** Amplification of the precursor mRNAs of the bacterial smRNAs from the *M. ciceri* bacteria. (**B**) qRT-PCR based expression profile of bacterial smRNA in bacterial RNA and chickpea nodule tissue.

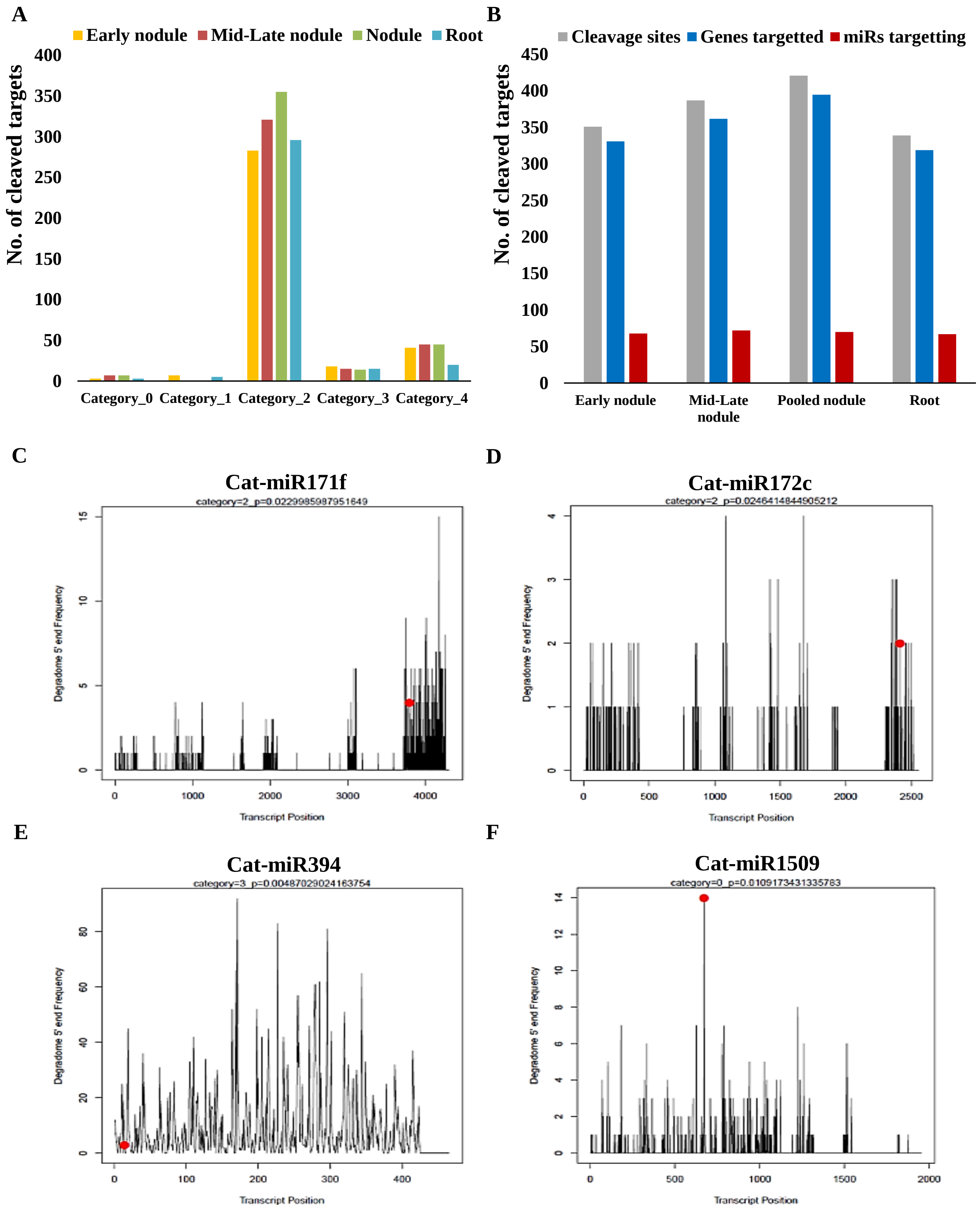


**Figure S5. Confirmed microRNA targets using degradome sequencing.** (**A**) Category wise distribution of cleaved targets. **(B)** Degardome stas representing the number of cleavage sites, genes cleaved and the miRs targetting the transcripts. The t-plots representing the category, p-value and cleavage position (red dot) for (**C**) cat-miR171f, (**D**) cat-miR172c, (**E**) cat-miR394 and (**F**) cat-miR1509

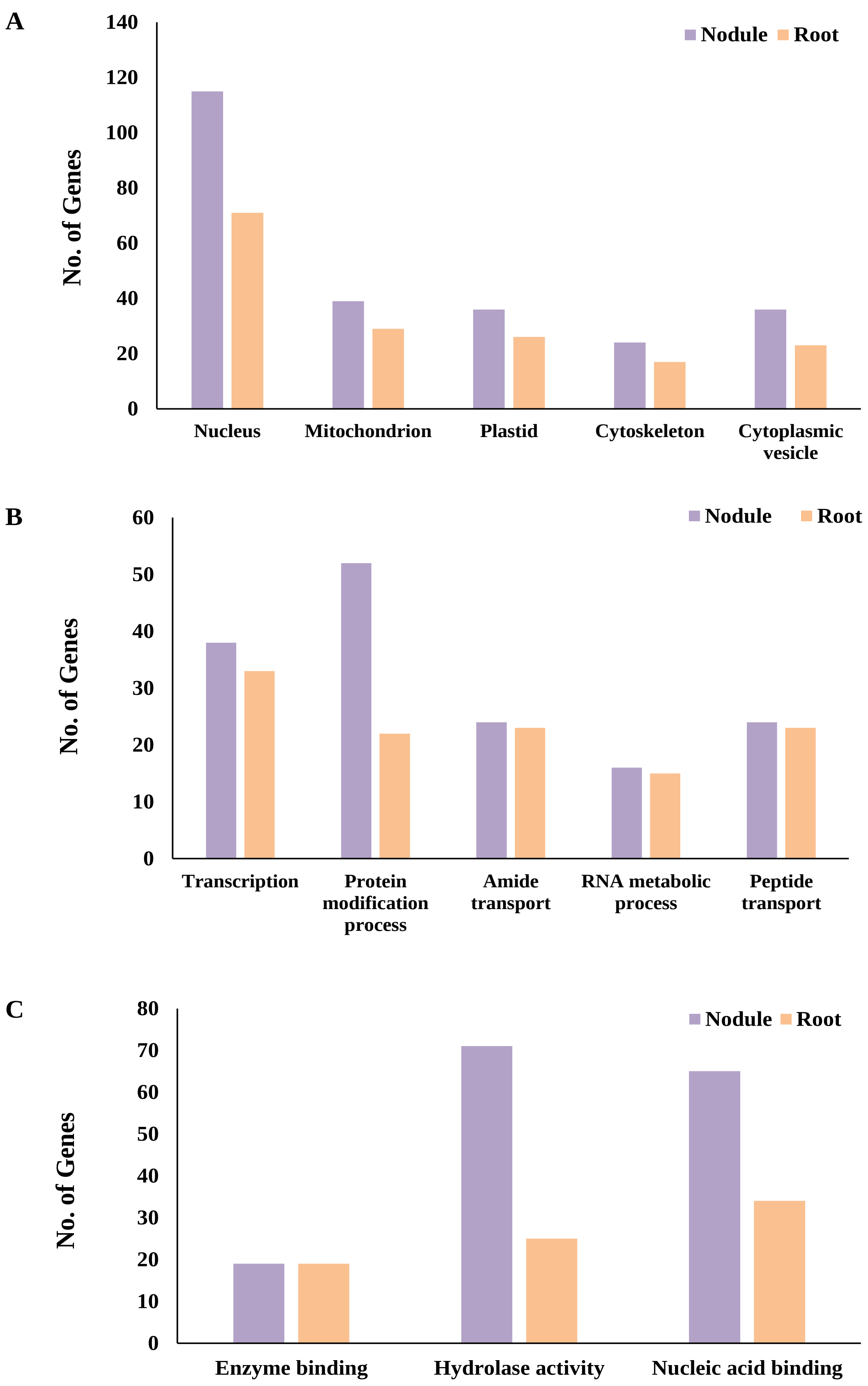


**Figure S6. Blast2GO based functional annotation of target transcripts. (A)** Cellular component **(B)** Biological process and **(C)** Molecular function.

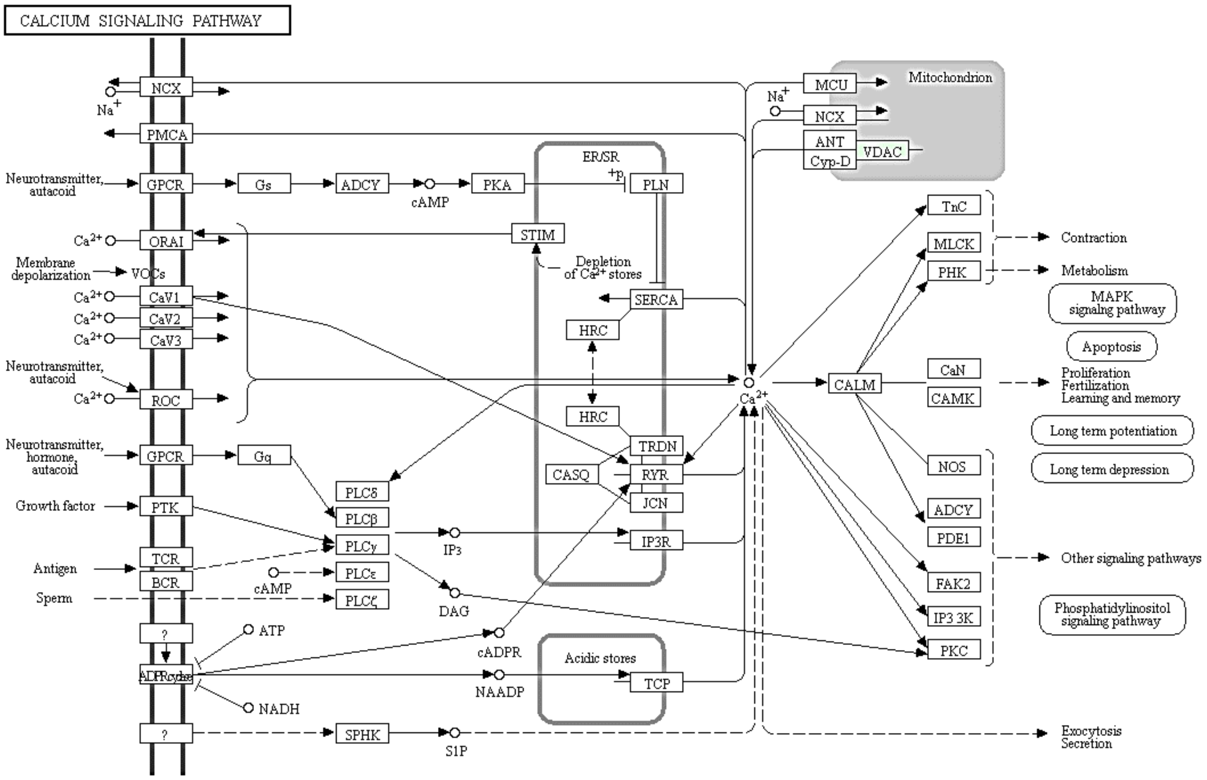


**Figure S7A. KEGG pathway analysis of targets.** Targets involved in calcium signaling pathway.

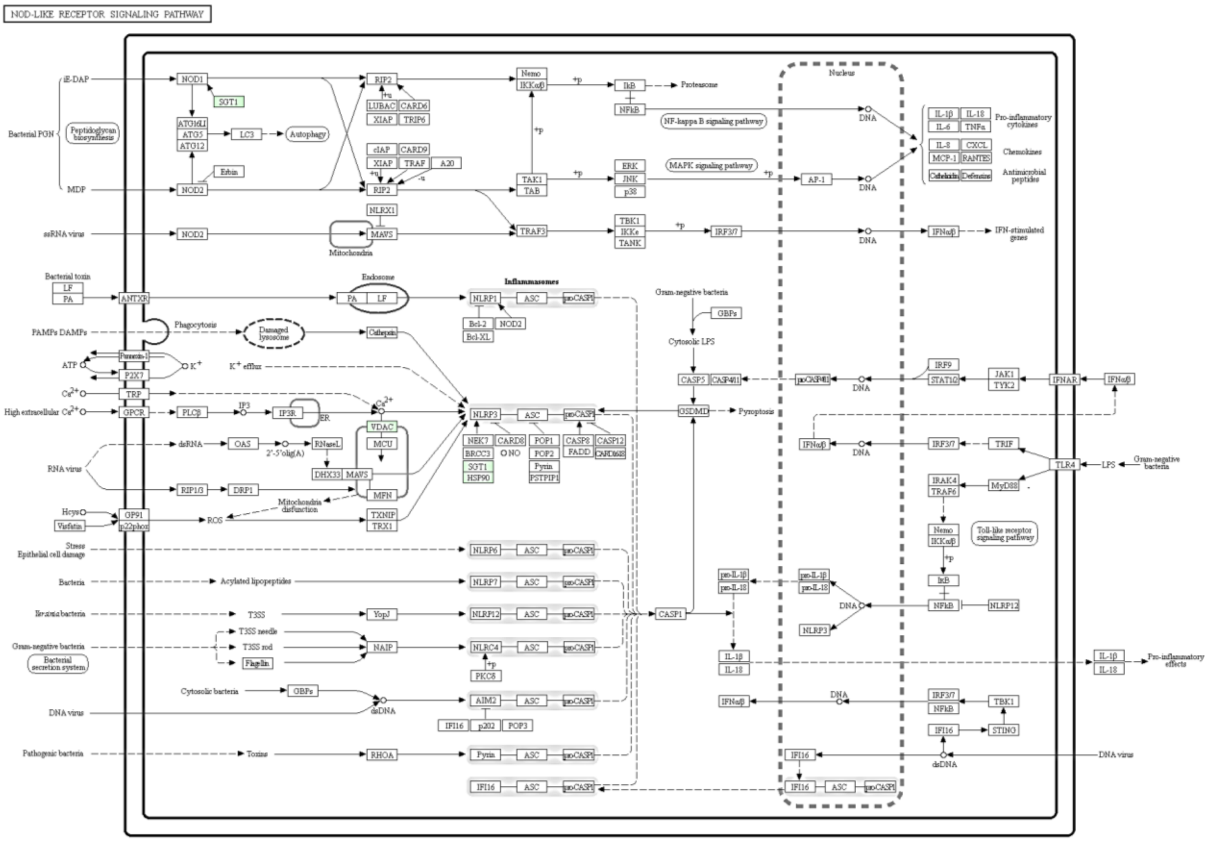


**Figure S7B. KEGG pathway analysis of targets.** Targets involved in Nod-like receptor signaling pathway.

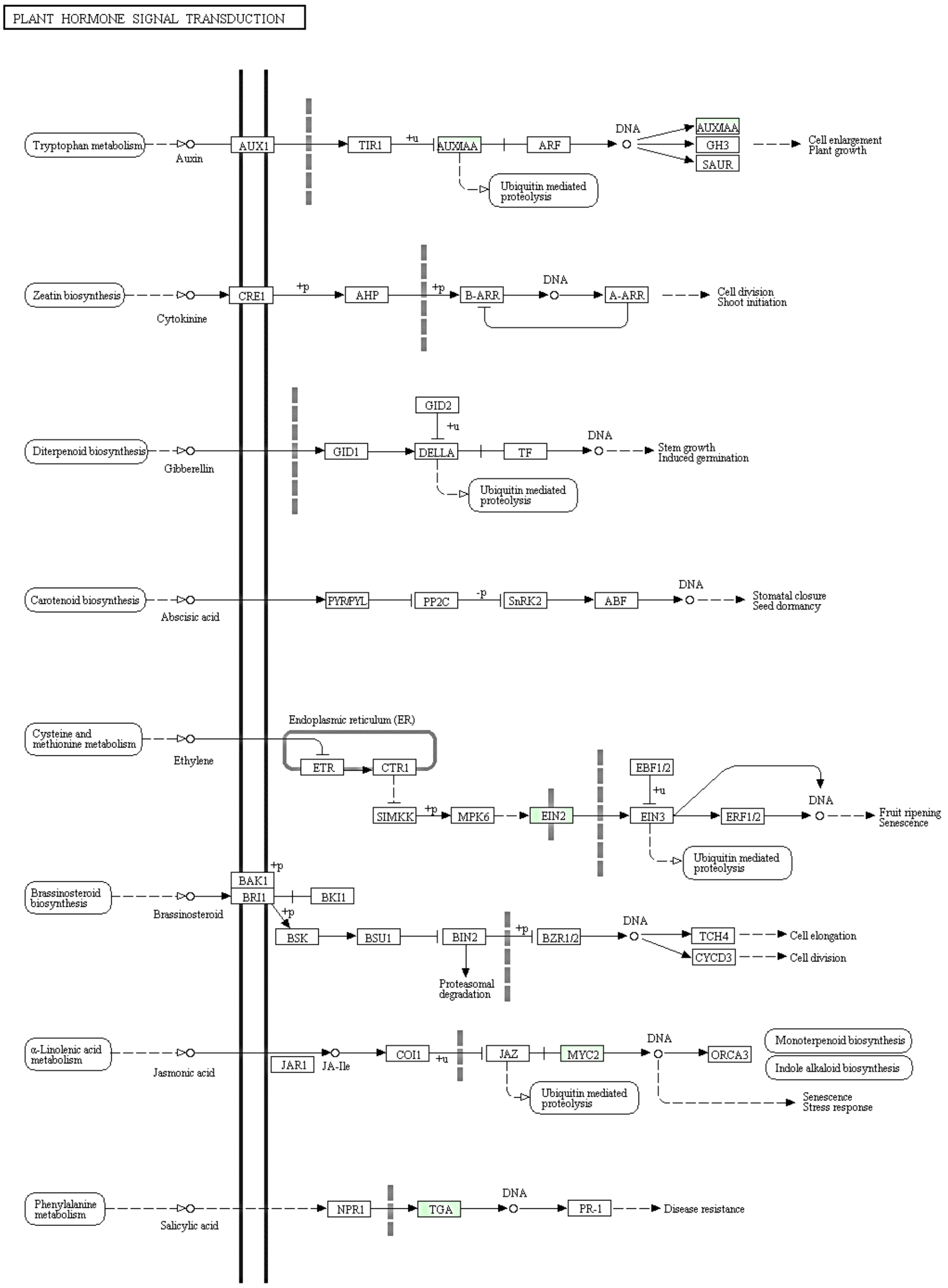


**Figure S7C. KEGG pathway analysis of targets.** Targets involved in plant hormone signal transduction pathway.

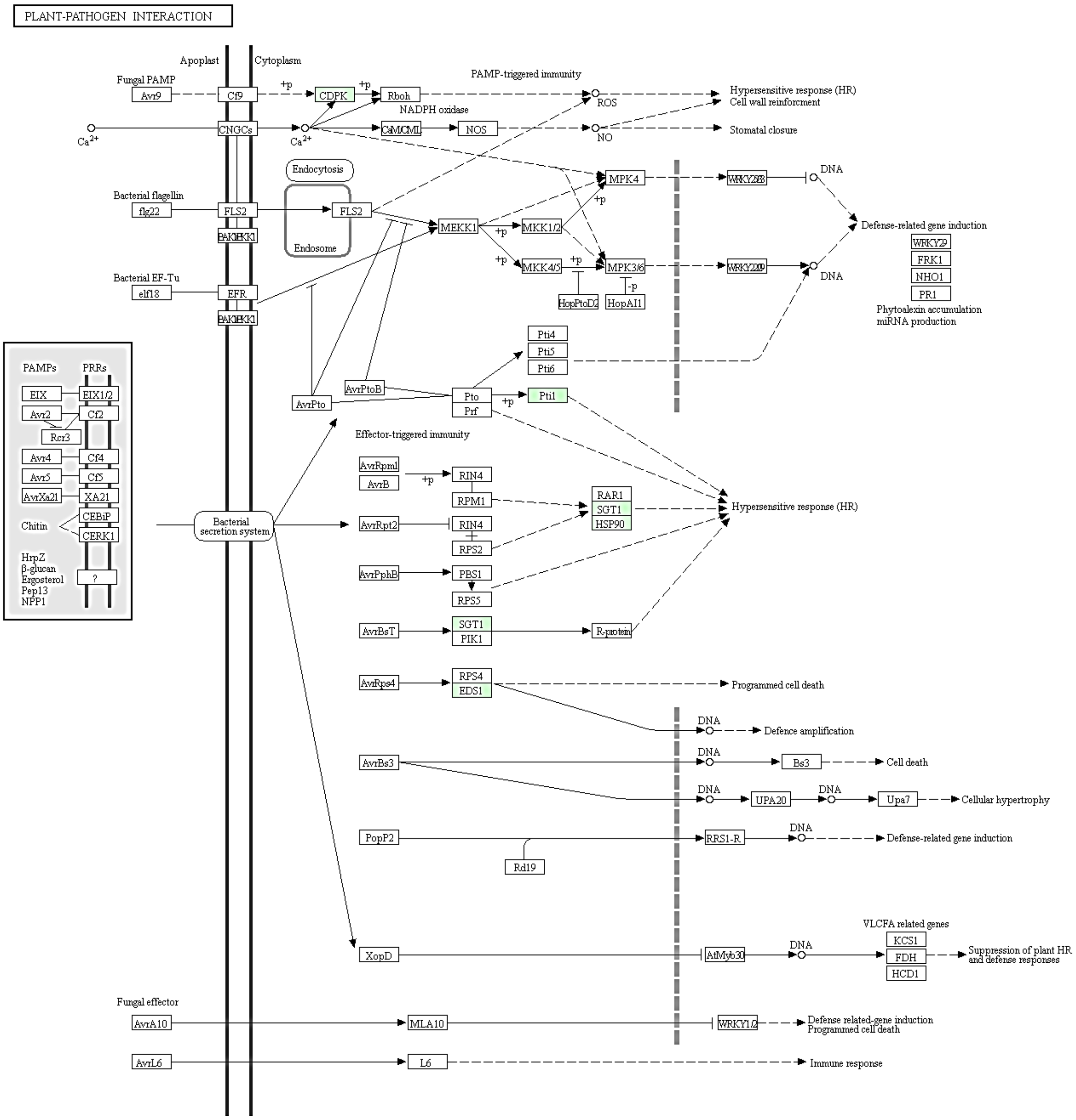


**Figure S7D. KEGG pathway analysis of targets.** Targets involved in plant pathogen interaction pathway.
