## Supplementary material for "Identification of miRNAs and their corresponding mRNA targets from chickpea nodules and functional characterization of candidate miRNAs by overexpression in chickpea roots": Table S1-S9

**Table S1. The length distribution of sequenced reads of microRNA in library**

| | |
|---|---|
| **Total Reads** | 21,760,971 |
| **Filtered Reads** | 4,445,569 |
| **Unique Tags** | 1,315,289 |

| | 20 nt | 21 nt | 22 nt | 23 nt | 24 nt |
|---|---|---|---|---|---|
| **Total Reads** | 730,049 | 1,028,108 | 696,535 | 665,149 | 1,325,728 |
| **%age of each Reads** | 16.42194734 | 23.12658 | 15.66807 | 14.96206672 | 29.82133 |
| **Unique Tags** | 134,181 | 179,316 | 175,292 | 172,229 | 654,271 |
| **%age of each Tags** | 10.20163629 | 13.6332 | 13.32726 | 13.09438458 | 49.74352 |

**Table S2. The detailed information about the miRNAs, chromosomal location, start and end position, length, sequence information of mature and precursors miRNAs with their corresponding RPM values from nodule library.**

| miR_name | Chr | Strand | Precursor Start | Precursor End | Mature_miRs | Length | Precursors | RPM | Specificity |
|---|---|---|---|---|---|---|---|---|---|
| cat-miR1507 | Ca7 | - | 45098706 | 45098787 | TTTCATTCCATACATCGTCTAA | 22 | AGAGTTGTATGGAGTGAAAGATGGTTTTTTTTTGGGATTTTCTAAATTGAACTTTCCCTCTTTCATTCCATACATCGTCTAA | 1.406124 | **Chickpea** |
| cat-miR1509 | Ca3 | + | 49536297 | 49536377 | TTAATCAGGGAAATCACAGTTG | 22 | TTAATCAGGGAAATCACAGTTGAGATGGTGTGAGATCTCTGTGTCTTTTTTTCTTTCTCTACTGTGTTTCCCTGTTTAAAG | 793.2881 | **Chickpea** |
| cat-miR1511 | Ca7 | + | 8736934 | 8737016 | AACCAGGCTCTGATACCATGA | 21 | ATGGTATCAGGTCCTGCTTCATCAAGTGTCTTGAGTTCTACTCCCAAATCAAGTACATGATTAACCAGGCTCTGATACCATGA | 2943.251 | **Chickpea** |
| cat-miR156f-2 | Ca6 | + | 2332750 | 2332851 | GCTCTCTAGACTTCTGTCATC | 21 | TGACAGAAGATAGAGAGCACAGACTATGATATACATATACATATATAGATACGCTATATAATTGTATCTCATCTCCTTTGTGCTCTCTAGACTTCTGTCATC | 63.74427 | **Legume** |
| cat-miR156f-1 | Ca6 | + | 2318065 | 2318166 | GCTCTCTAGACTTCTGTCATC | 21 | TGACAGAAGATAGAGAGCACAGAGTATGATATACATATACATATATAGATACGCTATATAATTGTATCTCATCTCCTTTGTGCTCTCTAGACTTCTGTCATC | 63.74427 | **Legume** |
| cat-miR156e-1 | Ca6 | + | 2318049 | 2318180 | TGCTCTCTAGACTTCTGTCATC | 22 | ATAATTAAATATAAGGTTGTTGACAGAAGATAGAGAGCACAGAGTATGATATACATATACATATATAGATACGCTATATAATTGTATCTCATCTCCTTTGTGCTCTCTAGACTTCTGTCATCTACCTCTAATTTAATTTCCA | 253.1023 | |
| cat-miR156e-2 | Ca8 | - | 11926199 | 11926349 | TGCTCTCTAGACTTCTGTCATC | 22 | ATAATTAAGTATAAGGTTGTTGACAGAAGATAGAGAGCACAGAGGATGATATACATATACATATATCTATACACTATATAGATGCAGGAGTTGAAGATAATTGTATCTCATCTCCTTTGTGCTCTCTAGACTTCTGTCATCTACCTCTAATTTAATTTCCA | 66.32217 | |
| cat-miR156a-3 | Ca8 | - | 11926313 | 11926539 | TTGACAGAAGATAGAGAGCAC | 21 | ATTCTCTATTTTCAATTCATAAGTCTCTTCTTATATTTAATATTATTTTACTCTATTTTCAATTCATAAGTCTCTTCTTATATTTAATATTATTTTAATTTGTTTTACTTTGTATGTACTTATATATATATTTTTTATATGAAGAAATGAATATTTATATTGGAGAAAATTAAGAGGGAGGGAGGAAATAATTAAGTATAAGGTTGTTGACAGAAGATAGAGAGCAC | 33.74697 | |
| cat-miR156a-1 | Ca3 | + | 33683372 | 33683457 | TTGACAGAAGATAGAGAGCAC | 21 | TTGACAGAAGATAGAGAGCACTGATGATGAAATGCATGAAAAACAATGCATTTCATTTATTTGTGCTTTCTATACTTCTGTCATCA | 42.88677 | |
| cat-miR156a-2 | Ca6 | - | 12698093 | 12698171 | TTGACAGAAGATAGAGAGCAC | 21 | TTGACAGAAGATAGAGAGCACAGATGATGACATGCATCGTGCATTTTATTCATTTGTGCTCTCTATGCTTCTGTCATCA | 34.45003 | |
| cat-miR156b | Ca4 | - | 54282273 | 54282352 | TGTGCTCATTCTATTCTGTCA | 21 | ACAGAAGAGAATGAGCACAGATAAGCATTTGATATATACTCTCTATACCATTTTTTATCTGTGCTCATTCTATTCTGTCA | 4.218371 | **Chickpea** |
| cat-miR156c-1 | Ca4 | - | 54809558 | 54809642 | TGACAGAAGAGAGTGAGCAC | 20 | TGACAGAAGAGAGTGAGCACACATGTATTTTCTTGTATGATGTTTCATGCTTGAAGCTATGCGTGCTCACTTTCTATCTGTCACC | 534.7957 | |
| cat-miR156c-2 | Ca5 | + | 17651771 | 17651855 | TGACAGAAGAGAGTGAGCAC | 20 | TGACAGAAGAGAGTGAGCACACAGAGGCATTCGATATAGACAATTATACCGTTACTTTTGCGTGCTCACTGCTCTTTCTGTCAAC | 780.8673 | |
| cat-miR156c-3 | Ca5 | - | 18239104 | 18239192 | TGACAGAAGAGAGTGAGCAC | 20 | TGACAGAAGAGAGTGAGCACACATAGTGTTTTCTTGCACGATGACGTTTCATGCTTGAAGTTATGTGTGCTTACTCTCTATCTGTCACC | 530.8117 | |
| cat-miR156c-4 | Ca7 | + | 45145189 | 45145273 | TGACAGAAGAGAGTGAGCAC | 20 | TGACAGAAGAGAGTGAGCACATACTGTTTGTTATTGTATCATGGCATACAATTCTGGGTGCGTGCTCACTTCTCTTTCTGTCATC | 771.9619 | |
| cat-miR156d | Ca5 | + | 47774204 | 47774292 | TTGACAGAAGAGAGAGAGCAC | 21 | TTGACAGAAGAGAGAGAGCACAACCCGAGAATGGTTGAAAGAGAATCTTTATCTTTGTTGGGAGTGTGCTCTCCCTTCTTCTGTCATCA | 8.436742 | |
| cat-miR159-2 | Ca8 | + | 10444080 | 10444244 | TTTGGATTGAAGGGAGCTCTA | 21 | GAGCTCCTTTTAGTCCAAATGAGGATCTTGCTGTGTTGATAGAGCTGCTTAGCTATGGATCCCTCAATTCTACCCATTTATGTTTTGTGGTAGTTTTGTGGCTTCCATATTTAGGGAGCTTTATCACCTATAGTCTTATCTTTCTTTGGATTGAAGGGAGCTCTA | 32815.41 | |
| cat-miR159-1 | Ca7 | + | 3643368 | 3643532 | TTTGGATTGAAGGGAGCTCTA | 21 | GAGCTCCTTTTAGTCCAAATGAGGATCTTGCTGTGTTGATAGAGCTGCTTAGCTATGGATCCCTCAATTCTACCCATTTATGTTTTGTGGTAGTTTTGTGGCTTCCATATTTAGGGAGCTTTATCACCTATAGTATTATCTTTCTTTGGATTGAAGGGAGCTCTA | 32815.41 | |
| cat-miR160-1 | Ca3 | - | 40858682 | 40858763 | TGCCTGGCTCCCTGTATGCCA | 21 | TGCCTGGCTCCCTGTATGCCATTTGTAGAGCCTATCAAGATAGTGATGACTTTAGTGAATGGCGTACAAGGAGCCATGCATG | 64.21298 | |
| cat-miR160-3 | Ca4 | - | 19285356 | 19285437 | TGCCTGGCTCCCTGTATGCCA | 21 | TGCCTGGCTCCCTGTATGCCATTTGTAGAGTTCATCAAAATAGTGATGGCTTTAATGAATGGCGTGCGAGGAGCCATGCATG | 63.97863 | |
| cat-miR160-2 | Ca4 | - | 9997140 | 9997221 | TGCCTGGCTCCCTGTATGCCA | 21 | TGCCTGGCTCCCTGTATGCCATATACAGAGCTCATCGAAACATCAATGACCTTTGTGGATGGCGTATGAGGAGCCAAGCATA | 64.21298 | |
| cat-miR162a | Ca4 | + | 1379511 | 1379599 | TCGGACCAGACTTCATTCCCC | 21 | GGAATATAGTCTGGTTCAAGATTATTCATAAATAGTCTCTTTCTAAAATATAACTTTTATGAATAATTTCGGACCAGACTTCATTCCCC | 7.030618 | **Chickpea** |
| cat-miR162b | Ca7 | + | 51425891 | 51425972 | TCGATAAACCTCTGCATCCAG | 21 | GGAGGCAGCGGTTCATCGATCTGTTCCTGAATTTGTTTGTGTAATACAAAACAATAATTGGTCGA | 126.5511 | |

| miR_name | Chr | Strand | Precursor Start | Precursor End | Mature_miRs | Length | Precursors | RPM | Specificity |
|---|---|---|---|---|---|---|---|---|---|
| | | | | | | | TAAACCTCTGCATCCAG | | |
| cat-miR164a-2 | Ca3 | + | 49640405 | 49640480 | TGGAGAAGCAGGGCACGTGCA | 21 | TGGAGAAGCAGGGCACGTGCAAACATAACTTTCTATTAGTTTGTCTCTCATTTTGCACGTGCTCCACTTTTCCAAC | 192.4046 | |
| cat-miR164a-1 | Ca3 | + | 29446839 | 29446979 | TGGAGAAGCAGGGCACGTGCA | 21 | TGGAGAAGCAGGGCACGTGCAATACTAACTCATGAAAACACACACACATGGAATTGAATTGCTAAGAGAATCCACATTCTTTTTTTTTCATTTCATTTTGTAGTGAATGAGTTAGTTCTTCATGTGCCCCTCTTCCCCATC | 193.8107 | |
| cat-miR164a-3 | Ca4 | + | 7468239 | 7468325 | TGGAGAAGCAGGGCACGTGCA | 21 | TGGAGAAGCAGGGCACGTGCAAATTCTCTCTTCTAATTCATTCTCTCAATGCATATCAATTTTTTGCACGCGCTCCCCTTCTCCAAC | 191.4672 | |
| cat-miR164b | Ca7 | + | 23774058 | 23774122 | CATGTGTTCTTGTTCTCCATC | 21 | TGGAGAAGCAGGGCACATGCTTGATGACTTACCAAAATTCTTAGCATGTGTTCTTGTTCTCCATC | 2.812247 | |
| cat-miR166a-5 | Ca7 | - | 31406488 | 31406565 | TCGGACCAGGCTTCATTCCCC | 21 | GGAATGTTGTCTGGCTCGAGGACTCTTCTTCTTGATCTATAAAGAAGTGTAGTATTCTCGGACCAGGCTTCATTCCCC | 6064.143 | |
| cat-miR166a-1 | Ca4 | - | 2867500 | 2867581 | TCGGACCAGGCTTCATTCCCC | 21 | GGAATTGTGTCTGGTTCGAGACCATTTACCTTAAGTAAATTGAATTGATTTAAGAATGATCTCGGACCAGGCTTCATTCCCC | 5916.5 | |
| cat-miR166a-2 | Ca5 | - | 29873124 | 29873214 | TCGGACCAGGCTTCATTCCCC | 21 | GGAATGTTGGCTGGCTCGAGGCTTTTCTTTTCACCCACTTACACCTTCCCTCGTTCTATAAATTAAAGCTTCGGACCAGGCTTCATTCCCC | 5197.267 | |
| cat-miR166a-3 | Ca6 | + | 16719642 | 16719742 | TCGGACCAGGCTTCATTCCCC | 21 | GGAATGTTGGCTGGCTCGAGGCTTTTGAGTTTGACAAAGGAGATAGATATTTCATTCCATCTTGGAAGAACTATAAGGCTTCGGACCAGGCTTCATTCCCC | 5197.267 | |
| cat-miR166a-4 | Ca6 | + | 16719799 | 16719892 | TCGGACCAGGCTTCATTCCCC | 21 | GGAACGTTGTCTGGCTCGAGGAGATAAAGATGGTTGAAGAGAATACTTTCTCATCATTCATTCACTCTCAACCTCGGACCAGGCTTCATTCCCC | 4477.332 | |
| cat-miR166b-3 | Ca8 | + | 764356 | 764440 | TCGGACCAGGCTTCATTCCTC | 21 | GGAATGTTGTCTGGTTCGAGACCATTCTTCTGAAGCAATTAAATTAACCTATCAATGAATGATTTCGGACCAGGCTTCATTCCTC | 459.5681 | |
| cat-miR166b-1 | Ca5 | + | 27903657 | 27903740 | TCGGACCAGGCTTCATTCCTC | 21 | GGAATGGTGTCTGGTCCAAGGAGATCCATCCATTACTCACTTTCACTTCAATCATTTCTCTCCTCGGACCAGGCTTCATTCCTC | 463.3177 | |
| cat-miR166b-2 | Ca5 | - | 29872976 | 29873054 | TCGGACCAGGCTTCATTCCTC | 21 | GGAATGTTGTCTGGCTCGAGGTCATATTACTTCTCATCGATCTGATCCGATCCCATCCTCGGACCAGGCTTCATTCCTC | 599.0087 | |
| cat-miR167c | Ca4 | + | 30476512 | 30476580 | TGAAGCTGCCAGCATGATCT | 20 | TGAAGCTGCCAGCATGATCTCAATTTTCCTTTTTATAAGGGAAAGATTAGATCATGCGGCTGTTTCACT | 65.6191 | |
| cat-miR167d | Ca6 | + | 60244647 | 60244717 | TGAAGCTGCCAGCATGATCTT | 21 | TGAAGCTGCCAGCATGATCTTAACTTCCCTCCTTTTGTAGGGGAAAGATCAGATCATGTGGCAGTTTCACC | 157.0171 | |
| cat-miR167a-1 | Ca3 | + | 39126024 | 39126149 | TGAAGCTGCCAGCATGATCTA | 21 | TGAAGCTGCCAGCATGATCTAAACTTAGGTTAGAGAGGTTTTCTTTTCTTTTCTTTAAAAGAAGAACAAGATGAAGATGAAAGACTAACCCTAACTTGGATTAGGTCATGCTGTGACAGCCTCACT | 246.0716 | |
| cat-miR167a-2 | Ca4 | - | 1611551 | 1611635 | TGAAGCTGCCAGCATGATCTA | 21 | TGAAGCTGCCAGCATGATCTAGGTTTGATTGGTTATACGATAGTATAACATAATTTTGATTAGGTCATGCTGTGCTAGCCTCACT | 242.5563 | |
| cat-miR167b-1 | Ca3 | + | 50353774 | 50353869 | TGAAGCTGCCAGCATGATCTGA | 22 | TGAAGCTGCCAGCATGATCTGAGCTTACCTAGCTTTGCTTCTTCAATTTCTGTATGGCGGTGCCAAGGCAATAACAGATCATGTCGGAGCTTCACC | 2722.958 | |
| cat-miR167b-2 | Ca4 | - | 4867904 | 4867971 | TGAAGCTGCCAGCATGATCTGA | 22 | TGAAGCTGCCAGCATGATCTGAATTTACTTTTGTAATGGGTAAGAATAGATCATGCGGCAACTTCACC | 2722.958 | |
| cat-miR168 | Ca8 | + | 7115132 | 7115236 | TCGCTTGGTGCAGGTCGGGAA | 21 | CTCACTTCGCGGTCTCTAATTCGCTTGGTGCAGGTCGGGAACCACTTCTTCGACAGTTTTCGTGCGGAAATGACGGAACGGTCACCCGCGGAGAATTGGATCCCGCCTTGCATCAACTGAATCGGAGGCAGCGGTGAATATT | 394.1833 | |
| cat-miR171a | Ca1 | - | 5436538 | 5436619 | TTGAGCCGTGCCAATATCACA | 21 | AGATATTGGTACAGTTCAATCAAAAGGACGTGTTTTATGAATCTAAAGCTTTCTTATTTGATTGAGCCGTGCCAATATCACA | 3.280955 | |
| cat-miR171b | Ca1 | - | 13770966 | 13771059 | TTGAGCCGCGTCAATATCTTG | 21 | AGGTATTGGCGCGCCTCAATTTGAAGACATGGCTATGAATCATGGAAAATTAATTAGCAGCCATGTAGTTTGATTGAGCCGCGTCAATATCTTG | 4.452725 | **Chickpea** |
| cat-miR171c | Ca5 | - | 27766156 | 27766228 | CGAGCCGAATCAATATCACCC | 21 | GTGATATTGTTTCTGCTCATTTTAATCACATACACATTGTTGTAGTATAAGACGAGCCGAATCAATATCACCC | 690.4067 | **Chickpea** |
| cat-miR171d | Ca5 | - | 27766327 | 27766441 | TGTGATATTGATACGGCTCATC | 22 | TGTGATATTGATACGGCTCATCTTCAATTTCAACCATATTCATATTTCCATTCTTCAATTTCAACTATATTTATATTCATATTTACATTTAGATGAGCCGAATCAATATCACTCT | 20.15444 | **Chickpea** |
| cat-miR171e | Ca6 | + | 28037698 | 28037781 | TTGAGCCGCGTCAATATCTCG | 21 | AGGTATTGGCGTGCCTCAATTTGAAGACATGGTTAACATAAAATTACCAACCATGTAATTTGATTGAGCCGCGTCAATATCTCG | 7.499326 | **Chickpea** |
| cat-miR171f | Ca7 | - | 50402576 | 50402661 | TTGAGCCGCGCCAATATCACT | 21 | CGATGTTGGTGAGGTTCAATCTGAAGACGGATTTACATGTAGAAGAAGTAAAATACGATCTCAGATTGAGCCGCGCCAATATCACT | 92.1011 | |
| cat-miR172a-5p | Ca1 | - | 10989804 | 10989881 | GCAGCATCATCAAGATTCACA | 21 | GCAGCATCATCAAGATTCACAAGGCTTATCAATTTGAAGTTTCATTATATAAATTTGAGAATCTTGATGATGCTGCAT | 0.468708 | |
| cat-miR172a-3p | Ca1 | - | 10989805 | 10989880 | AGAATCTTGATGATGCTGCA | 20 | CAGCATCATCAAGATTCACAAGGCTTATCAATTTGAAGTTTCATTATATAAATTTGAGAATCTTGATGATGCTGCA | 0.468708 | |
| cat-miR172b | Ca2 | + | 16100540 | 16100629 | CGAATCCTGATGATGCTGCAG | 21 | GGAGCATCATCAAGATTCACGATCTATATTTTGAAGGGTTTAATTTATTAGCCCTTTGATATATAAATACGAATCCTGATGATGCTGCAG | 1.874832 | **Chickpea** |
| cat-miR172c-2 | Scaffold0556 | + | 563 | 654 | AGAATCTTGATGATGCTGCAG | 21 | GGAGCATCATCAAGATTCACAATCTTTAAGGGGCTTTTTTGATTTGTTTGAGGTGTGGTCCCTCTATATTGAGAATCTTGATGATGCTGCAG | 2.812247 | |
| cat-miR172c-1 | Ca6 | - | 65531331 | 65531422 | AGAATCTTGATGATGCTGCAG | 21 | GGAGCATCATCAAGATTCACAATCTTTAAGGGGCTTTTTTGATTTGTTTGAGGTGTGGTCCCTCTATATTGAGAATCTTGATGATGCTGCAG | 2.812247 | |
| cat-miR2111-3 | Ca3 | - | 7647381 | 7647445 | TAATCTGCATCCTGAGGTTT | 20 | TAATCTGCATCCTGAGGTTTAGAACAATATTTGTATTGTATCTAGTCCTCGGAATGCAGATTATC | 19.68573 | |
| cat-miR2111-2 | Ca3 | - | 7647253 | 7647320 | TAATCTGCATCCTGAGGTTT | 20 | TAATCTGCATCCTGAGGTTTAGAACAATAGTTGTGTAATTGTTTCTAGTCCTTAGAATGTAGATTATC | 16.40478 | |

| miR_name | Chr | Strand | Precursor Start | Precursor End | Mature_miRs | Length | Precursors | RPM | Specificity |
|---|---|---|---|---|---|---|---|---|---|
| cat-miR2111-1 | Ca3 | - | 7625969 | 7626033 | TAATCTGCATCCTGAGGTTT | 20 | TAATCTGCATCCTGAGGTTTAAAATAACGAGTTTATTATATTTAATCTTTGGAATGCAGATTACC | 16.40478 | |
| cat-miR2111-4 | Ca6 | + | 52845756 | 52845820 | TAATCTGCATCCTGAGGTTT | 20 | TAATCTGCATCCTGAGGTTTAAAACAATATATTTATTGTATCTAGTCCTTAGGATGTAGATTATC | 16.40478 | |
| cat-miR2111-5 | Ca6 | + | 52864589 | 52864653 | TAATCTGCATCCTGAGGTTT | 20 | TAATCTGCATCCTGAGGTTTAAAACAATATATTTATTGTGTCTAGTCCTTAGGATACAGATTATC | 16.40478 | |
| cat-miR2118 | Ca4 | - | 15079567 | 15079659 | TTACCGATTCCACCCATTCCTA | 22 | GGATATGGGAGGGTCGGTAAAGTAGATCTGAATCTGATTAATCTTGTAGATTTTCAGTACTAAATATTTCTTTACCGATTCCACCCATTCCTA | 134.2848 | **Legume** |
| cat-miR319a | Ca1 | + | 33458275 | 33458338 | TTGGACTGAAGGGAGCTCCT | 20 | GAGTTCCTTGCAGCCCAAAGCAGAGAACTTCTTGCAATGTTGTTTTGGACTGAAGGGAGCTCCT | 325.9863 | |
| cat-miR319b | Ca2 | + | 11504015 | 11504147 | TGGACTGAAGGGAGCTCCTTC | 21 | AGGAGCTTCCTACAGCCCAACCATAGATATAAGAGCAGAATTAATTAGTACTAAGTGAGTGATATAATAAAAAAATGATATTCCATTTAGAATGAAATAATAGCAATGTGGTTGGACTGAAGGGAGCTCCTTC | 5802.135 | **Legume** |
| cat-miR319d | Ca5 | - | 28460150 | 28460324 | TGGACTGAAGGGGAGCTCCTTC | 22 | AGGAGCTTCCTTCAGCCCGTGCATGGATATAGTTGCTGAAGTATCTATATATCTTCTGATCTCAGTCATTCTTTCATCAATTAATTAATTAATTAATATATGAATGAATGAGTGAGAAGGGATATATTATATTTCGGATCGATATATCAATCTTGGACTGAAGGGGAGCTCCTTC | 3301.813 | **Legume** |
| cat-miR319c-4 | Ca8 | + | 1468937 | 1469106 | TTGGACTGAAGGGAGCTCCCT | 21 | GGAGCTTTCTTTAGTCCACTAATGGGTGACATGTAGGATTTAATTAGCTGCCGAATCATTCATCCAAATGGTGAGTAAATAGAATGAATGAATACTCATCAAATGAGTGAATGAAGCGGGAGACAAATTGAATCTTAAGTTTCCTATACTTGGACTGAAGGGAGCTCCCT | 1672.35 | |
| cat-miR319c-1 | Ca2 | + | 14183650 | 14183824 | TTGGACTGAAGGGAGCTCCCT | 21 | AGAGCTTCCTTCAGTCCACTCATGGAAGAGTAAGGGGTTTGAATTACCTGCTGACTCATTCATTCAAACACAATAGATAATGATTCATATATAAAAGCTATTGTGAATGTGTGAATGATGCAGGAGGTGAATTTCTTCCTTTTCTTCTCTTTGCTTGGACTGAAGGGAGCTCCCT | 1671.412 | |
| cat-miR319c-2 | Ca5 | + | 10324943 | 10325116 | TTGGACTGAAGGGAGCTCCCT | 21 | AGAGCTTCCTTCAGTCCACTCATGGATAGGAAAAAGTTTTGAATTAGCTGCTGACTCATTCATTCAAACACAATAGAAATAAAACATAATGATATGCTATTGTGAATGTGTGGATGATGCAGGAGCTGAATTTCATCTTTCTCTTGTCTTTGCTTGGACTGAAGGGAGCTCCCT | 1692.504 | |
| cat-miR319c-3 | Ca6 | + | 15008473 | 15008645 | TTGGACTGAAGGGAGCTCCCT | 21 | AGAGCTTTCTTCGGTCCACTCATGGGTGGTTGTAAGATTGAATTAGCTGCTGACTCATTCATCCAAATGTTGAGTATAGTAATCCAATCAATAAATACTCATCAAATGACTGAATGATGCGGGAGACAGATTCAATCTCATGCTTCCTGTATTTGGACTGAAGGGAGCTCCCT | 1615.402 | |
| cat-miR390a-2 | Ca5 | + | 9117902 | 9117974 | AAGCTCAGGAGGGATAGCGCC | 21 | AAGCTCAGGAGGGATAGCGCCGCTTATATCTATATATTTATATGGTAATTGGCGCTATCCATCCTGAGTTTTA | 70.07183 | |
| cat-miR390a-1 | Ca2 | + | 12992522 | 12992589 | AAGCTCAGGAGGGATAGCGCC | 21 | AAGCTCAGGAGGGATAGCGCCATGAGATATGATATACGTACGTTTGGCGCTATCCATCCTGAGTTTTA | 70.07183 | |
| cat-miR390b | Ca6 | + | 46146838 | 46146929 | CGCTATCCATCCTGAGTTTCA | 21 | AAGCTCGGGAGGGATAGCGCCATAGAATATCTTCAATTTCTCTTCTTTGTTGTTATATTGATTTTCTCTAGCGCTATCCATCCTGAGTTTCA | 558.9342 | |
| cat-miR394-1 | Ca4 | + | 58435248 | 58435317 | TTGGCATTCTGTCCACCTCC | 20 | TTGGCATTCTGTCCACCTCCCTTAACTAGATATTTTTCTAAATATATCGGAGGTGGACACACTGCCAATT | 12.88947 | |
| cat-miR394-2 | Ca5 | - | 14116931 | 14116999 | TTGGCATTCTGTCCACCTCC | 20 | TTGGCATTCTGTCCACCTCCCTTCTCTCTTTCTTTTTGCTTTGCTTTGGAGGTGGGCATACTGCCAACA | 12.88947 | |
| cat-miR396a | Ca7 | + | 50347278 | 50347414 | TTCCACAGCTTTCTTGAACTG | 21 | TCGTGCTCCTCTTTGTATTCTTCCACAGCTTTCTTGAACTGCATACAAAAGAGTTCCTTTCCATGCATGCCATGGCATTCATTCTTCTACACAATATTGTTTTGCGGTTCAATAAAGCTGTGGGAAGATACAGATAGGGTCAATTAC | 1280.979 | |
| cat-miR396b | Ca7 | + | 50347296 | 50347402 | TTCCACAGCTTTCTTGAACT | 20 | TTCCACAGCTTTCTTGAACTGCATACAAAAGAGTTCCTTTCCATGCATGCCATGGCATTCATTCTTCTACACAATATTGTTTTGCGGTTCAATAAAGCTGTGGGAAG | 1805.697 | |
| cat-miR396c | Ca7 | - | 50356311 | 50356402 | TTCCACAGCTTTCTTGAACTT | 21 | TTCCACAGCTTTCTTGAACTTCTTTTGCATCTTAAATCTTTCCAAATTCAAGATTAATAGCCCTAGAAGAAGCTCAAGAAAGCTGTGGGAGA | 2992.934 | |
| cat-miR398a-3p | Ca2 | + | 2467754 | 2467864 | TGTGTTCTCAGGTCACCCCTT | 21 | GGAGTGAATCTTAGAACACAAGATTGATTGGTTAGACAACAATATCATCTATTGTCATATGGACTATATTTGTTTCTTGCTAATTTATTTTGTGTTCTCAGGTCACCCCTT | 3.280955 | |
| cat-miR398a-5p | Ca2 | + | 2467755 | 2467863 | GAGTGAATCTTAGAACACAAGA | 22 | GAGTGAATCTTAGAACACAAGATTGATTGGTTAGACAACAATATCATCTATTGTCATATGGACTATATTTGTTTCTTGCTAATTTATTTTGTGTTCTCAGGTCACCCCT | 11.01464 | **Chickpea** |
| cat-miR398b-1 | Ca2 | + | 19357721 | 19357798 | TGTGTTCTCAGGTCGCCCCTG | 21 | GGGTCGTCCTTAGATCACATGAAGCTGCTGTCCTATGTATCTAAGCTATCCAACTCATGTGTTCTCAGGTCGCCCCTG | 20.38879 | |
| cat-miR398b-2 | Ca2 | - | 19441272 | 19441346 | TGTGTTCTCAGGTCGCCCCTG | 21 | GGGTCGTCCTAAGACCACATGAAGCTATCTTGTGTGCCTATGCTATCTAGCTCATGTGTTCTCAGGTCGCCCCTG | 20.38879 | |
| cat-miR399-3 | Ca8 | + | 18958632 | 18958718 | TGCCAAAGGAGAGTTGCCCTG | 21 | GGGCGCCTCTCACCTGACAAGTAAACAGTCGATCAGTATATTGGAGCGAACTCGATGTCAATGACTTGCCAAAGGAGAGTTGCCCTG | 9.139804 | |
| cat-miR399-2 | Ca8 | - | 18926087 | 18926170 | TGCCAAAGGAGAGTTGCCCTG | 21 | AGGCACCTCTCACCTGGCAAGTAAAGTGTCAGTATATTTGGAGCTAATTCAATGCCATTTACTTGCCAAAGGAGAGTTGCCCTG | 9.139804 | |
| cat-miR399-1 | Ca4 | + | 12492894 | 12492970 | TGCCAAAGGAGAGTTGCCCTG | 21 | GGGCTTCTCTTTGTTGGCAGGAAAGTAACTATCATATTGTAATATATGGTGTGTCTTGCCAAAGGAGAGTTGCCCTG | 9.139804 | |
| cat-miR408 | Ca4 | + | 6654506 | 6654594 | ATGCACTGCCTCTTCCCTGGC | 21 | CAGGGAACAGGCTGAGCATGGATGGAACTATCGACAGAATTGAGGAGACAACTGTCATGGTTCCACTCATGCACTGCCTCTTCCCTGGC | 9.608512 | |
| cat-miR482 | Ca3 | + | 33671732 | 33671821 | TTACCAATTCCGCCCATTCCTA | 22 | GGAATGGGTTGTTTGGTTAGAAACACTTTCAAATTCATAAGCTTAATCCGCTTTGCTTTGTGTGTTTCTTACCAATTCCGCCCATTCCTA | 42.18371 | **Chickpea** |
| cat-miR5213 | Ca1 | + | 28202359 | 28202444 | TACGGGTGTCTTCACCTCTGA | 21 | TACGGGTGTCTTCACCTCTGAACTATGATTCTTATGTTTCAATCTTTGATTCATTGTTTAGATTCGGATGTGTAGATACCCGCATC | 6.093203 | **Chickpea** |
| cat-NovmiR1 | Ca2 | + | 215573 | 215651 | TGTTTGGAGGAACTCTCGGAA | 21 | CCTAGAGTTCTCTTAAACATTTCACTAAAGATTCCATTTCTCGTCCACGTTCTTGAAGTGTTTGGAGGAACTCTCGGAA | 13.59253 | |
| cat-NovmiR2 | Ca3 | + | 33484522 | 33484585 | GCTGGATTCTTTTGAAGGAAC | 21 | TCCCTCAAAGGCTTCCAGTATTCGGTCATATAATTGCTTGAATGCTGGATTCTTTTGAAGGAAC | 3.515309 | |
| cat-NovmiR3 | Ca4 | + | 837664 | 837777 | TTTCGGGAGTGAGAATCAGTGA | 22 | ACTGATCTCACTCCTGAAATGCATTTAGTTAATTTCGAAGGCAACACTAGCTAGCTAGTTGATCG | 2.343539 | |

| miR_name | Chr | Strand | Precursor Start | Precursor End | Mature_miRs | Length | Precursors | RPM | Specificity |
|---|---|---|---|---|---|---|---|---|---|
| cat-NovmiR4 | Ca5 | - | 61737154 | 61737236 | AAGGGTCTGTTTGAGAGAAGTGGT | 24 | ATCACATGTGAATGGTTATGAGATACATTTCGGGAGTGAGAATCAGTGA<br>AAGGGTCTGTTTGAGAGAAGTGGTAAAGAGGGGAAAGACTCCCCATTTTGTTAATTTGCCACTT<br>TTTTCAAACAGTCCCCGAG | 134.5192 | |
| cat-NovmiR5 | Ca6 | + | 64879657 | 64879754 | AACCGGCACAAACTTGACTTG | 21 | AGTCTAGCTTGTGCCGTTTCTTCGTCATTCTTAAGTTCCACCAGTCAGGCCCTCCTTGAACAACA<br>GAGATGCTGAAGAACCGGCACAAACTTGACTTG | 206.2315 | |
| cat-NovmiR6 | Ca6 | + | 8531285 | 8531374 | CGGAATCGCCACCGCAACAGC | 21 | TGTTGCGTTGGCGATTACCATGGCTCTCTCGTCACCACTTCAGAACCTCTCTTCATTAATCTCTGC<br>TCACGGAATCGCCACCGCAACAGC | 5.390141 | |
| cat-NovmiR7 | Scaffold0567 | + | 1377 | 1458 | CTATCAATTACTTCATCTTGT | 21 | AACATCTATACAACAGAATGCTATCAATTACTTCATCTTGTCTATCCTTTTATATCCTTCATCTTG<br>TGAAAAAGGATAAACAAGATGAAGTAATGGATAACATTTATGTTTTTTTGGAGAAA | 2.109186 | |

### Highlighted rows are Polycistronic miRs :19 (20.87%)

**Table S3. The detailed information of length distribution of mature miRNAs and their first nucleotide distribution**

| miR_name | Mature_miRs | Length |
|---|---|---|
| cat-miR156c-1 | TGACAGAAGAGAGTGAGCAC | 20 |
| cat-miR156c-2 | TGACAGAAGAGAGTGAGCAC | 20 |
| cat-miR156c-3 | TGACAGAAGAGAGTGAGCAC | 20 |
| cat-miR156c-4 | TGACAGAAGAGAGTGAGCAC | 20 |
| cat-miR167c | TGAAGCTGCCAGCATGATCT | 20 |
| cat-miR172a-3p | AGAATCTTGATGATGCTGCA | 20 |
| cat-miR2111-3 | TAATCTGCATCCTGAGGTTT | 20 |
| cat-miR2111-2 | TAATCTGCATCCTGAGGTTT | 20 |
| cat-miR2111-1 | TAATCTGCATCCTGAGGTTT | 20 |
| cat-miR2111-4 | TAATCTGCATCCTGAGGTTT | 20 |
| cat-miR2111-5 | TAATCTGCATCCTGAGGTTT | 20 |
| cat-miR319a | TTGGACTGAAGGGAGCTCCT | 20 |
| cat-miR394-1 | TTGGCATTCTGTCCACCTCC | 20 |
| cat-miR394-2 | TTGGCATTCTGTCCACCTCC | 20 |
| cat-miR396b | TTCCACAGCTTTCTTGAACT | 20 |
| cat-miR1511 | AACCAGGCTCTGATACCATGA | 21 |
| cat-miR156f-2 | GCTCTCTAGACTTCTGTCATC | 21 |
| cat-miR156f-1 | GCTCTCTAGACTTCTGTCATC | 21 |
| cat-miR156a-3 | TTGACAGAAGATAGAGAGCAC | 21 |
| cat-miR156a-1 | TTGACAGAAGATAGAGAGCAC | 21 |
| cat-miR156a-2 | TTGACAGAAGATAGAGAGCAC | 21 |
| cat-miR156b | TGTGCTCATTCTATTCTGTCA | 21 |
| cat-miR156d | TTGACAGAAGAGAGAGAGCAC | 21 |
| cat-miR159-2 | TTTGGATTGAAGGGAGCTCTA | 21 |
| cat-miR159-1 | TTTGGATTGAAGGGAGCTCTA | 21 |
| cat-miR160-1 | TGCCTGGCTCCCTGTATGCCA | 21 |
| cat-miR160-3 | TGCCTGGCTCCCTGTATGCCA | 21 |
| cat-miR160-2 | TGCCTGGCTCCCTGTATGCCA | 21 |
| cat-miR162a | TCGGACCAGACTTCATTCCCC | 21 |
| cat-miR162b | TCGATAAACCTCTGCATCCAG | 21 |
| cat-miR164a-2 | TGGAGAAGCAGGGCACGTGCA | 21 |

| miR_name | Mature_miRs | Length |
|---|---|---|
| cat-miR164a-1 | TGGAGAAGCAGGGCACGTGCA | 21 |
| cat-miR164a-3 | TGGAGAAGCAGGGCACGTGCA | 21 |
| cat-miR164b | CATGTGTTCTTGTTCTCCATC | 21 |
| cat-miR166a-5 | TCGGACCAGGCTTCATTCCCC | 21 |
| cat-miR166a-1 | TCGGACCAGGCTTCATTCCCC | 21 |
| cat-miR166a-2 | TCGGACCAGGCTTCATTCCCC | 21 |
| cat-miR166a-3 | TCGGACCAGGCTTCATTCCCC | 21 |
| cat-miR166a-4 | TCGGACCAGGCTTCATTCCCC | 21 |
| cat-miR166b-3 | TCGGACCAGGCTTCATTCCTC | 21 |
| cat-miR166b-1 | TCGGACCAGGCTTCATTCCTC | 21 |
| cat-miR166b-2 | TCGGACCAGGCTTCATTCCTC | 21 |
| cat-miR167d | TGAAGCTGCCAGCATGATCTT | 21 |
| cat-miR167a-1 | TGAAGCTGCCAGCATGATCTA | 21 |
| cat-miR167a-2 | TGAAGCTGCCAGCATGATCTA | 21 |
| cat-miR168 | TCGCTTGGTGCAGGTCGGGAA | 21 |
| cat-miR171a | TTGAGCCGTGCCAATATCACA | 21 |
| cat-miR171b | TTGAGCCGCGTCAATATCTTG | 21 |
| cat-miR171c | CGAGCCGAATCAATATCACCC | 21 |
| cat-miR171e | TTGAGCCGCGTCAATATCTCG | 21 |
| cat-miR171f | TTGAGCCGCGCCAATATCACT | 21 |
| cat-miR172a-5p | GCAGCATCATCAAGATTCACA | 21 |
| cat-miR172b | CGAATCCTGATGATGCTGCAG | 21 |
| cat-miR172c-2 | AGAATCTTGATGATGCTGCAG | 21 |
| cat-miR172c-1 | AGAATCTTGATGATGCTGCAG | 21 |
| cat-miR319b | TGGACTGAAGGGAGCTCCTTC | 21 |
| cat-miR319c-4 | TTGGACTGAAGGGAGCTCCCT | 21 |
| cat-miR319c-1 | TTGGACTGAAGGGAGCTCCCT | 21 |
| cat-miR319c-2 | TTGGACTGAAGGGAGCTCCCT | 21 |
| cat-miR319c-3 | TTGGACTGAAGGGAGCTCCCT | 21 |
| cat-miR390a-2 | AAGCTCAGGAGGGATAGCGCC | 21 |
| cat-miR390a-1 | AAGCTCAGGAGGGATAGCGCC | 21 |

| miR_name | Mature_miRs | Length |
|---|---|---|
| cat-miR390b | CGCTATCCATCCTGAGTTTCA | 21 |
| cat-miR396a | TTCCACAGCTTTCTTGAACTG | 21 |
| cat-miR396c | TTCCACAGCTTTCTTGAACTT | 21 |
| cat-miR398a-3p | TGTGTTCTCAGGTCACCCCTT | 21 |
| cat-miR398b-1 | TGTGTTCTCAGGTCGCCCCTG | 21 |
| cat-miR398b-2 | TGTGTTCTCAGGTCGCCCCTG | 21 |
| cat-miR399-3 | TGCCAAAGGAGAGTTGCCCTG | 21 |
| cat-miR399-2 | TGCCAAAGGAGAGTTGCCCTG | 21 |
| cat-miR399-1 | TGCCAAAGGAGAGTTGCCCTG | 21 |
| cat-miR408 | ATGCACTGCCTCTTCCCTGGC | 21 |
| cat-miR5213 | TACGGGTGTCTTCACCTCTGA | 21 |
| cat-NovmiR1 | TGTTTGGAGGAACTCTCGGAA | 21 |
| cat-NovmiR2 | GCTGGATTCTTTTGAAGGAAC | 21 |
| cat-NovmiR5 | AACCGGCACAAACTTGACTTG | 21 |
| cat-NovmiR6 | CGGAATCGCCACCGCAACAGC | 21 |
| cat-NovmiR7 | CTATCAATTACTTCATCTTGT | 21 |
| cat-miR1507 | TTTCATTCCATACATCGTCTAA | 22 |
| cat-miR1509 | TTAATCAGGGAAATCACAGTTG | 22 |
| cat-miR156e-1 | TGCTCTCTAGACTTCTGTCATC | 22 |
| cat-miR156e-2 | TGCTCTCTAGACTTCTGTCATC | 22 |
| cat-miR167b-1 | TGAAGCTGCCAGCATGATCTGA | 22 |
| cat-miR167b-2 | TGAAGCTGCCAGCATGATCTGA | 22 |
| cat-miR171d | TGTGATATTGATACGGCTCATC | 22 |
| cat-miR2118 | TTACCGATTCCACCCATTCCTA | 22 |
| cat-miR319d | TGGACTGAAGGGGAGCTCCTTC | 22 |
| cat-miR398a-5p | GAGTGAATCTTAGAACACAAGA | 22 |
| cat-miR482 | TTACCAATTCCGCCCATTCCTA | 22 |
| cat-NovmiR3 | TTTCGGGAGTGAGAATCAGTGA | 22 |
| cat-NovmiR4 | AAGGGTCTGTTTGAGAGAAGTGGT | 24 |

| | 20nt | 21nt | 22nt | 24nt | Total |
|---|---|---|---|---|---|
| **No. of MiRs** | 15 | 63 | 12 | 1 | 91 |
| **%age of total** | 16.48352 | 69.23077 | 13.18681 | 1.098901 | 100 |

| | First nucleotide | | | | |
|---|---|---|---|---|---|
| **Length of miRs** | **A** | **G** | **C** | **T** | **Total** |
| **20 nt miRs** | 1 (6.66%) | 0 (0%) | 0 (0%) | 14 (93.33%) | 14 |
| **21 nt miRs** | 7 (11.11%) | 4 (6.34%) | 6 (9.52%) | 46 (73.01%) | 63 |
| **22 nt miRs** | 0 (0%) | 1 (8.33%) | 0 (0%) | 11 (91.66%) | 12 |
| **24 nt miRs** | 1 (100%) | 0 (0%) | 0 (0%) | 0 (0%) | 1 |

**Table S4. The detailed information about the other plant species miRNAs, chromosomal location, start and end position, length, sequence information of mature and precursors miRNAs with their corresponding RPM values from nodule library**

| Chr | Precursor start | Precursor end | Strand | Mature miRNA sequence | Mature miRNA | Precursor sequence |
|---|---|---|---|---|---|---|
| ***Arachis hypogea*** | | | | | | |
| arahy.Tifrunner.gnm1.Arahy.01 | 98830773 | 98830900 | - | TTGAGCCGCGCCAATATCACT | ahy-miR171-1 | GAGAAAGAAAGAAAGGAAAGCGATGTTGGTGAGGTTCAATCCGAAGACGGATTTACATGGTATAATCCAGTAAAATACGATCTCAGATTGAGCCGCGCCAATATCACTTTTTCATTTATATCCATATT |
| arahy.Tifrunner.gnm1.Arahy.01 | 108768469 | 108768595 | + | TCGATAAACCTCTGCATCCAG | ahy-miR162-1 | ATTTGTAAGGTGAAGTCACTGGAGGCAGCGGTTCATCGATCTCTTCCTGAATTTGGCTGTGGAATCATACACAAAGCAAGAATCGGTCGATAAACCTCTGCATCCAGCGCTCACTTTGCCTCTTTTT |
| arahy.Tifrunner.gnm1.Arahy.03 | 132995755 | 132995878 | - | AACCAGGCTCTGATACCATGA | ahy-miR1511 | TGGGTCGAGTGGCTGCTTTCGTGGTATCAGGTCCTGCTTCACCAAGTGATCTTGTGTTCAATTCCCATTCCAAGCACATGGTTAACCAGGCTCTGATACCATGATGCGTTCTCTTGCACTTCCC |
| arahy.Tifrunner.gnm1.Arahy.05 | 2723854 | 2723984 | - | TCGCTTGGTGCAGGTCGGGAA | ahy-miR168-1 | CTCACTGCGCGGTCTCTAATTCGCTTGGTGCAGGTCGGGAACCGATCTTCCCGCGCGTAATGGATGCCGTTGGCGGCGGCGAATTTGGATCCCGCCTTGCATCAACTGAATCGGAGGCCACGGTGAAGATT |
| arahy.Tifrunner.gnm1.Arahy.09 | 110335471 | 110335588 | - | TGAAGCTGCCAGCATGATCTT | ahy-miR167-1 | GATCATGCACCACTACAAGTTGAAGCTGCCAGCATGATCTTAACTTTCCCTCTCCTATGATTTGTTGGGGTGAGATCAGATCATGTGGCAGTTTCACCTAGTTGTTGGAAGCATGAAT |
| arahy.Tifrunner.gnm1.Arahy.10 | 6288066 | 6288186 | + | TCGGACCAGGCTTCATTCCCG | ahy-miR166-1 | ATGGGAGATTGAGGTTGATGGGAATGTTGTTTGGCTCGAGGTCACTTATTGAGTAGTTATATACCCTTTATAGTGAGATCTCGGACCAGGCTTCATTCCCGTCGACTTGATCTCTCTCGCT |
| arahy.Tifrunner.gnm1.Arahy.11 | 132159843 | 132159969 | - | TCGATAAACCTCTGCATCCAG | ahy-miR162-2 | ATTTGTAAGGTGAAGTCACTGGAGGCAGCGGTTCATCGATCTCTTCCTGAATTTGGCTGTGGAATCATACACAAAGCAAGAATCGGTCGATAAACCTCTGCATCCAGCGCTCACTTTGCCTCTTTCT |
| arahy.Tifrunner.gnm1.Arahy.11 | 145417047 | 145417174 | + | TTGAGCCGCGCCAATATCACT | ahy-miR171-2 | ATGGGAGAAAGAAAGGAAAGCGATGTTGGTGAGGTTCAATCCGAAGACGGATTTACATGGTATAATCCAGTAAAATACGATCTCAGATTGAGCCGCGCCAATATCACTTTTCATTCATATCCATATTC |
| arahy.Tifrunner.gnm1.Arahy.11 | 145486202 | 145486346 | + | TTCCACAGCTTTCTTGAACTT | ahy-miR396 | TCTGAGTCCTGGTCATGCTTTTCCACAGCTTTCTTGAACTTCTTGTATGTGCATCCATCTCAACTTCTTCTAAGTAAGTATGTAATTACAGTCTTAACCAAGAAGCTCAAGAAAGCTGTGGGAGAATATGGCAATTTCAGGCCTT |
| arahy.Tifrunner.gnm1.Arahy.15 | 2723854 | 2723984 | - | TCGCTTGGTGCAGGTCGGGAA | ahy-miR168-2 | CTCACTGCGCGGTCTCTAATTCGCTTGGTGCAGGTCGGGAACCGATCTTCCCGCGCGTAATGGATGCCGTTGGCGGCGGCGAATTTGGATCCCGCCTTGCATCAACTGAATCGGAGGCCACGGTGAAGATT |
| arahy.Tifrunner.gnm1.Arahy.19 | 157637786 | 157637903 | + | TGAAGCTGCCAGCATGATCTT | ahy-miR167-2 | GATCATGCACCACTACAAGTTGAAGCTGCCAGCATGATCTTAACTTTCCCTCTCCTATGATTTGTTGGGGTGAGATCAGATCATGTGGCAGTTTCACCTAGTTGTTGGAAGCATGAAT |
| arahy.Tifrunner.gnm1.Arahy.20 | 11507523 | 11507643 | + | TCGGACCAGGCTTCATTCCCG | ahy-miR166-2 | ATGGGAGATTGAGGTTGATGGGAATGTTGTTTGGCTCGAGGTCACTTATTGAGTAGTTGTATACCCTTTATAATGAGATCTCGGACCAGGCTTCATTCCCGTCGACTTGATCTCTCTCGTT |
| ***Arachis duranensis*** | | | | | | |
| aradu.V14167.gnm1.Aradu.A01 | 94030435 | 94030564 | + | TTCCACAGCTTTCTTGAACTG | adu-miR396 | ATATGGCCCTCTTTGTATTCTTCCACAGCTTTCTTGAACTGCATCTAATTGTTGCATGCATGCCATGGCACTCTTCTTCTTGCGTTGCGGTTCAATAAAGCTGTGGGAAGATACAGATAGGGTCAAACGC |
| aradu.V14167.gnm1.Aradu.A01 | 94090142 | 94090269 | - | TTGAGCCGCGCCAATATCACT | adu-miR171 | GAGAAAGAAAGAAAGGAAAGCGATGTTGGTGAGGTTCAATCCGAAGACGGATTTACATGGTATAATCCAGTAAAATACGATCTCAGATTGAGCCGCGCCAATATCACTTTTTCATTTATATCCATATT |
| aradu.V14167.gnm1.Aradu.A01 | 103703401 | 103703527 | + | TCGATAAACCTCTGCATCCAG | adu-miR162 | ATTTGTAAGGTGAAGTCACTGGAGGCAGCGGTTCATCGATCTCTTCCTGAATTTGGCTGTGGAATCATACACAAAGCAAGAATCGGTCGATAAACCTCTGCATCCAGCGCTCACTTTGCCTCTTTTT |
| aradu.V14167.gnm1.Aradu.A02 | 83264365 | 83264509 | + | CGCTATCCATCCTGAGTTTCA | adu-miR390a | ATAGTGTGGAAGAATCTGTAAAGCTCAGGAGGGATAGCGCCACGAAAGCATCGGCGATGATCCGTTGTTTGTGAAGAGTTTTTATGCTCTTTGATTTTCTGTAGCGCTATCCATCCTGAGTTTCATGGGTTCTTCACACTGCTTC |
| aradu.V14167.gnm1.Aradu.A03 | 109726686 | 109726786 | + | TCGGACCAGGCTTCATTCCTC | adu-miR166a | GGATGAGTTGGTGGTTGAGGGGAATGATGGCTGGCTCGAGGCTTCAATTTGGTTTGGGCTTCGGACCAGGCTTCATTCCTCTCAAACTAACTAAACTCCAT |
| aradu.V14167.gnm1.Aradu.A03 | 109726844 | 109726986 | + | TCGGACCAGGCTTCATTCCCC | adu-miR166b-1 | TGCAAAGGTTAAGGTTGAGAGGAATGTTGTCTGGCTCGAGAGGTCTTATTAGCTTCATGAATTGTTTGATTTGATGATGATTAGTCAAAGTGACTAATAATCTCGGACCAGGCTTCATTCCCCCCATCCAACCTTTGCTATAT |
| aradu.V14167.gnm1.Aradu.A05 | 2611261 | 2611391 | - | TCGCTTGGTGCAGGTCGGGAA | adu-miR168 | CTCACTGCGCGGTCTCTAATTCGCTTGGTGCAGGTCGGGAACCGATCTTCCCGCGCGTAATGGATGCCGTTGGCGGCGGCGAATTTGGATCCCGCCTTGCATCAACTGAATCGGAGGCCACGGTGAAGATT |
| aradu.V14167.gnm1.Aradu.A05 | 15261219 | 15261432 | - | TTGGACTGAAGGGAGCTCCCT | adu-miR319 | TGCATGGAAGGAGGTAAGAGAGAGCTTTCTTCAGTCCACTCATGGGTGACAATAAGATTTAATTAGCTGCCGACTCATTCATCCAAATGCTGAGTGAAGATATAGATAGAGAGATACACTCAGTAAATGAGTGAATGATGCGGGAGACAAATTGAATCTTATGTTTCCTATACTTGGACTGAAGGGAGCTCCCTTTTTTTGTTCTTCTTCATAT |
| aradu.V14167.gnm1.Aradu.A07 | 22352638 | 22352746 | - | CGCTATCTATCCTGAGTTTCA | adu-miR390b | GTAGTAAGGTAGAATCTGTTAAGCTCAGGAGGGATAGCGCCACATATAACACAATTCATTCGTATTGGCGCTATCTATCCTGAGTTTCACGGCTTCTTCTTACTTTAAA |
| aradu.V14167.gnm1.Aradu.A08 | 46224025 | 46224169 | - | TCGGACCAGGCTTCATTCCCC | adu-miR166b-2 | TGTTGAGGGGAATGTTGGCTGGCTCGAGGCTTTTCACAAAATGAAGGTTCTCACTCACTCACTCATCATATTTTGGGAGATGAAGTAGTAGAACTATTAAGGCTTCGGACCAGGCTTCATTCCCCTCAAAATATCTTTTATATGC |
| aradu.V14167.gnm1.Aradu.A09 | 94006421 | 94006553 | + | TCGGACCAGGCTTCATTCCCC | adu-miR166b-3 | TTTTATATTTCGAGTTGAGGGGAATGTCGTTTGGTTCGAGACCATTCTTTTGATGAAGCATAGGTCACTACGCACTCGAAAAAAGAATGATTTCGGACCAGGCTTCATTCCCCCTAACTCAAAATGATGTCTT |
| aradu.V14167.gnm1.Aradu.A09 | 110881205 | 110881301 | - | TGAAGCTGCCAGCATGATCTGA | adu-miR167 | GAAGCTGCCAGCATGATCTTAACTTTCCCTCTCCTATGATTTGTTGGGGTGAGATCAGATCATGTGGCAGTTTCACCTAGTTGTTGGAAGCATGAAT |
| aradu.V14167.gnm1.Aradu.A09 | 119039906 | 119040027 | + | TGCCTGGCTCCCTGTATGCCA | adu-miR160 | CATGCATACATGTGTGTATGTGCCTGGCTCCCTGTATGCCATTTGTAGAGCTCATCGAAATATCGATGACCTCTGTAGATGGCGTATGAGGAGCCAAGCATATTCCATATACATACATGGAG |
| aradu.V14167.gnm1.Aradu.A10 | 6589989 | 6590097 | + | TCGGACCAGGCTTCATTCCCT | adu-miR166c | ATGGGAGATTGAGGTTGATGGGAATGTTGTTTGGCTCGAGGTCACTTATATACCCTTTATAGTGAGATCTCGGACCAGGCTTCATTCCCGTCGACTTGATCTCTCTCGC |
| ***Arachis ipaensis*** | | | | | | |
| araip.K30076.gnm1.Araip.B01 | 121580643 | 121580769 | - | TCGATAAACCTCTGCATCCAG | aip-miR162 | ATTTGTAAGGTGAAGTCACTGGAGGCAGCGGTTCATCGATCTCTTCCTGAATTTGGCTGTGGAATCATACACAAAGCAAGAATCGGTCGATAAACCTCTGCATCCAGCGCTCACTTTGCCTCTTTCT |
| araip.K30076.gnm1.Araip.B01 | 133748674 | 133748801 | + | TTGAGCCGCGCCAATATCACT | aip-miR171 | ATGGGAGAAAGAAAGGAAAGCGATGTTGGTGAGGTTCAATCCGAAGACGGATTTACATGGTATAATCCAGTAAAA |

| Chr | Precursor start | Precursor end | Strand | Mature miRNA sequence | Mature miRNA | Precursor sequence |
|---|---|---|---|---|---|---|
| araip.K30076.gnm1.Araip.B01 | 133824954 | 133825098 | + | TTCCACAGCTTTCTTGAACTT | aip-miR396a | TACGATCTCAGATTGAGCCGCGCCAATATCACTTTTCATTCATATCCATATTCTCTGAGTCCTGGTCATGCTTTTCCACAGCTTTCTTGAACTTCTTGTATGTGCATCCATCTCAACTTCTTCTAAGTAAGTATGTAATTACAGTCTTAACCAAGAAGCTCAAGAAAGCTGTGGGAGAATATGGCAATTTCAGGCCTT |
| araip.K30076.gnm1.Araip.B01 | 133831168 | 133831297 | - | TTCCACAGCTTTCTTGAACTG | aip-miR396b | ACATGGCCCTCTTTGTATTCTTCCACAGCTTTCTTGAACTGCATCTAATTGTTGCATGCATGCCATGGCACTCTTCTTCTTGCGTTGCGGTTCAATAAAGCTGTGGGAAGATACAGATAGGGTCAAACGC |
| araip.K30076.gnm1.Araip.B02 | 95165530 | 95165674 | + | CGCTATCCATCCTGAGTTTCA | aip-miR390a | ATAGTGTGGAAGAATCTGTAAAGCTCAGGAGGGATAGCGCCACGGAAGCATCGACGATGATCCGTTGTTTGTGAAGAGTTTTTACGCTCTTTGATCTTCTGTAGCGCTATCCATCCTGAGTTTCATGGGTTCTTCACACTGCTTC |
| araip.K30076.gnm1.Araip.B03 | 111545162 | 111545262 | + | TCGGACCAGGCTTCATTCCTC | aip-miR166a | GGATGAGTTGGTGGTTGAGGGGAATGTTGGCTGGCTCGAGGCTTCAATTTGGTTTGGGCTTCGGACCAGGCTTCATTCCTCTCAAACTAACTAAACTCCAT |
| araip.K30076.gnm1.Araip.B03 | 111545317 | 111545459 | + | TCGGACCAGGCTTCATTCCCC | aip-miR166b | TGCAAAGGTTAAGGTTGAGAGGAATGTTGTCTGGCTCGAGAGGTCTTATTAGCTTCATGAATTGTTTGATTTGATGATGATTAGTCAAAGTGTCTAATAATCTCGGACCAGGCTTCATTCCCCCCATCCAACCTTTGCTATAT |
| araip.K30076.gnm1.Araip.B03 | 125055332 | 125055454 | - | ATGGTATCAGGTCCTGCTTCA | aip-miR1511 | GGGTCGAGCGGCTGCTTTCGTGGTATCAGGTCCTGCTTCACCAAGCGATCTTGTGTTCAATTCCCATCCCAAGCACATGGTTAACCAGGCTCTTATACCATGATGCTCTCTCTCGCACTTCCC |
| araip.K30076.gnm1.Araip.B04 | 132623136 | 132623269 | - | TTGACAGAAGATAGAGAGCAC | aip-miR156 | GGAGGTATTTAGGCGTGTTGTTGACAGAAGATAGAGAGCACAGATGATGACATGCATAAAATTAGAATGAATATGGCATTTTACTCATTTGTGCTCTCTATTCTTCTGTCATCACCCTTAACTTTGGTCTCTGT |
| araip.K30076.gnm1.Araip.B05 | 2423496 | 2423626 | - | TCGCTTGGTGCAGGTCGGGAA | aip-miR168 | CTCACTGCGCGGTCTCTAATTCGCTTGGTGCAGGTCGGGAACCGATCTTCCCGCGCGTAATGGATGCCGTTGGCGGCGGCGAATTTGGATCCCGCCTTGCATCAACTGAATCGGAGGCCACGGTGAAGATT |
| araip.K30076.gnm1.Araip.B05 | 16094990 | 16095203 | - | TTGGACTGAAGGGAGCTCCCT | aip-miR319 | TGCATGGAAGGAGGTAAGAGAGAGCTTTCTTCAGTCCACTCATGGGTGACAATAAGATTTAATTAGCTGCCGACTCATTCATCCAAATGCTGAGTGAAGAGATAGATAGAGAGATACACTCAGTAAATGAGTGAATGATGCGGGAGACAAATTGAATCTTATGTTTCCTATACTTGGACTGAAGGGAGCTCCCTTTTTTTGTTCTTCTTCATAT |
| araip.K30076.gnm1.Araip.B07 | 23109315 | 23109423 | - | CGCTATCTATCCTGAGTTTCA | aip-miR390b | GTAGTAAGGTAGAATCTGTTAAGCTCAGGAGGGATAGCGCCACATATAACACAATTCATTCGTATTGGCGCTATCTATCCTGAGTTTCACGGCTTCTTCTTACTTTAAG |
| araip.K30076.gnm1.Araip.B09 | 135868897 | 135869018 | - | TGCCTGGCTCCCTGTATGCCA | aip-miR160 | CATGCATACATGTGTGTATGTGCCTGGCTCCCTGTATGCCATTTGTAGAGCTCATCGAAATATCTATGACCTCTGTAGATGGCGTATGAGGAGCCAAGCATATTCCATATACATACATGGAG |
| araip.K30076.gnm1.Araip.B09 | 146076538 | 146076655 | + | TGAAGCTGCCAGCATGATCTGA | aip-miR167 | GATCATGCACCACTACAAGTTGAAGCTGCCAGCATGATCTTAACTTTCCCTCTCCTATGATTTGTTGGGGTGAGATCAGATCATGTGGCAGTTTCACCTAGTTGTTGGAAGCATGAAT |
| araip.K30076.gnm1.Araip.B10 | 10968772 | 10968892 | + | TCGGACCAGGCTTCATTCCCG | aip-miR166c | ATGGGAGATTGAGGTTGATGGGAATGTTGTTTGGCTCGAGGTCACTTATTGAGTAGTTGTATACCCTTTATAATGAGATCTCGGACCAGGCTTCATTCCCGTCGACTTGATCTCTCTCGTT |
| **Arabidopsis** | | | | | | |
| Chr1 | 78912 | 79050 | - | TCGGACCAGGCTTCATCCCCT | ath-miR166a | TTAAGCTATTTCAGTTGAGGGGAATGTTGTCTGGATCGAGGATATTATAGATATATACATGTGTATGTTAATGATTCAAGTGATCATAGAGAGTATCCTCGGACCAGGCTTCATCCCCCCCAACATGTTATTGCCTCTG |
| Chr1 | 24913185 | 24913315 | - | TGCTCTCTAGACTTCTGTCATC | ath-miR156a | TTGAGAGGCATTGATAGTGTTGACAGAAGATAGAGAGCACAGATGATGAGATACAATTCGGAGCATGTTCTTTGCATCTTACTCCTTTGTGCTCTCTAGCCTTCTGTCATCACCTTTTATTTGCTTTATTG |
| Chr2 | 16061951 | 16062063 | + | CGCTATCCATCCTGAGTTTCA | ath-miR390a | AAAGTAGAGAAGAATCTGTAAAGCTCAGGAGGGATAGCGCCATGATGATCACATTCGTTATCTATTTTTTGGCGCTATCCATCCTGAGTTTCATTGGCTCTTCTTACTACAAT |
| Chr2 | 16340262 | 16340382 | + | TGCCTGGCTCCCTGTATGCCA | ath-miR160 | GTATATGTATTATATATGTATGCCTGGCTCCCTGTATGCCATATGCTGAGCCCATCGAGTATCGATGACCTCCGTGGATGGCGTATGAGGAGCCATGCATATCCTCATACATATATAATTA |
| Chr3 | 3599771 | 3599911 | - | AGAATCTTGATGATGCTGCAG | ath-miR172 | ACCAGCTACTGTTCGCTGTTGGAGCATCATCAAGATTCACAAATCATCAAGTATTCGTGTAAATAAACCCATTTATGATTAGATTTTTGATGTATGTATGAGAATCTTGATGATGCTGCAGCTGCAATCAGTGGCTTACAC |
| Chr4 | 10578632 | 10578777 | + | TCGCTTGGTGCAGGTCGGGAA | ath-miR168 | TCTCACCATCGGGCTCGGATTCGCTTGGTGCAGGTCGGGAACCAATTCGGCTGACACAGCCTCGTGACTTTTAAACCTTTATTGGTTTGTGAGCAGGGATTGGATCCCGCCTTGCATCAACTGAATCGGATCCTCGAGGTGTAAAA |
| Chr5 | 2634916 | 2635038 | - | TCGATAAACCTCTGCATCCAG | ath-miR162a | GAAGAAGAGTGAGAGTCGCTGGAGGCAGCGGTTCATCGATCTCTTCCTGTGAACACATTAAAAATGTAAAAGCATGAATAGATCGATAAACCTCTGCATCCAGCGTTTGCCTCTTGTATCTTT |
| Chr5 | 2838632 | 2838777 | + | TCGGACCAGGCTTCATTCCCC | ath-miR166b-1 | TGTGCGATTTAGTGTTGAGAGGATTGTTGTCTGGCTCGAGGTCATGAAGAAGAGAATCACTCGAATTAATTTGGAAGAACAAATTAAGAAAACCCTAGATGATTCTCGGACCAGGCTTCATTCCCCCTAACCTACTTATCGCTTTT |
| Chr5 | 3867193 | 3867330 | + | TGACAGAAGAGAGTGAGCAC | ath-miR156b | TGTGAAAGGTAATTAGGAGGTGACAGAAGAGAGTGAGCACACATGGTGGTTTCTTGCATGCTTTTTTGATTAGGGTTTCATGCTTGAAGCTATGTGTGCTTACTCTCTCTCTGTCACCCCTTCTCTCTCTCTCTCTTT |
| Chr5 | 7740590 | 7740715 | - | TCGATAAACCTCTGCATCCAG | ath-miR162b | GAGGAAAGAGTGAAGTCGCTGGAGGCAGCGGTTCATCGATCAATTCCTGTGAATATTTATTTTTGTTTACAAAAGCAAGAATCGATCGATAAACCTCTGCATCCAGCGCTGCTTGCTCTTCATGAC |
| Chr5 | 13611788 | 13611944 | + | TTCCACAGCTTTCTTGAACTT | ath-miR396 | TGAAGATCCTGGTCATACTTTTCCACAGCTTTCTTGAACTTTCTTTTTCATTTCCATTGTTTTTTTCTTAAACAAAAGTAAGAAGAAAAAAAACTTTAAGATTAAGCATTTTGGAAGCTCAAGAAAGCTGTGGGAAAACATGACAATTCAGGGTTTT |
| Chr5 | 23636941 | 23637074 | + | CGCTATCCATCCTGAGTTCCA | ath-miR390b | GTAGTAGAGAAATAGCTATAAAGCTCAGGAGGGATAGCGCCATGGCTCACCAGTGCTGTATGTTTTGTATATGCGTACATGTATATCTGTTGGCGCTATCCATCCTGAGTTCCATAGCTTCTTCTTGCTTATAT |
| Chr5 | 25504795 | 25504922 | + | TCGGACCAGGCTTCATTCCCC | ath-miR166b-2 | GAAGCGATTTAGGGTTTAGAGGAATGTTGTTTGGCTCGAGGTCATGGAGAGTAATTCGTTAACCCAACTCAAAACTCTAAATGATTCTCGGACCAGGCTTCATTCCCCTCAACCTATTTTATCGCATT |
| ***Cajanus cajan*** | | | | | | |
| cajca.ICPL87119.gnm1.Cc10 | 3999620 | 3999748 | - | TGCTCACTTTCTATCTGTCACC | cca-miR156b | GAGATTAAGGGTAAGGGAGGTGACAGACGAGAGTGAGCACACATGGTACTTTCTTGGATGATAATGTTTCATGCTTGAAATTATGTGTGCTCACTCTCTATCTGTCACCAACTCTATTTCTATCACTTT |
| cajca.ICPL87119.gnm1.Cc10 | 8958556 | 8958674 | - | TCGGACCAGGCTTCATTCCCC | cca-miR166-3 | GGAAGCTTTGTCTTTTGAGGGGAATGTTGTCTGGCTCGAGGACCCTTCTTCATCTTCATCAAGAAATGTTTAGTGTTCTCGGACCAGGCTTCATTCCCCCAATTCCATGCTTCCAAAT |
| cajca.ICPL87119.gnm1.Cc06 | 928064 | 928187 | + | TCGATAAACCTCTGCATCCAG | cca-miR162-1 | GAAGGATTGGTGAAGTCACTGGAGGTAGCGGTTCATCGATCTCTTCCTGAACTTTGTTGTTGAACATAACAACCATGAATCGATCGATAAACCTCTGCATCCAGCCCTTACTATTCCCCTTCAT |
| cajca.ICPL87119.gnm1.Cc06 | 4651568 | 4651710 | - | TGCTCTCTAGACTTCTGTCATC | cca-miR156a | ACAATAGTTGATAAGGTTGTTGACAGAAGATAGAGAGCACTAATGATGATATGCACACATAGATACATGAAACTATAAAATAATTGCATCTCACTCCTTTGTGCTCTCTAGGCTTCTGTCATCCACCTTCACTAACCTCCTTA |

| Chr | Precursor start | Precursor end | Strand | Mature miRNA sequence | Mature miRNA | Precursor sequence |
|---|---|---|---|---|---|---|
| cajca.ICPL87119.gnm1.Cc08 | 6048584 | 6048720 | - | TCGGACCAGGCTTCATTCCCC | cca-miR166-1 | CTGCAAAGGTTCGGTTGAGAGGAATGTTGTCTGGCTCGAGGTCATGGAAATATATATACATCAAGTGTTTCAGTTTCCAGGGTGAATTAATGATTCTCGGACCAGGCTTCATTCCCCCCAGCCAACATTTGCTTCTC |
| cajca.ICPL87119.gnm1.Cc08 | 6048778 | 6048904 | - | TCGGACCAGGCTTCATTCCCC | cca-miR166-2 | GTTAATTAGTTGTGTTGAGGGGAATGTTGGCTGGCTCGAGGCTTTTCACTAAGGAGGTTCTCACTTGCAAGAACAACTATAAGGCTTCGGACCAGGCTTCATTCCCCTCAAAACTAAAGTTTTCAAT |
| cajca.ICPL87119.gnm1.Cc11 | 11675424 | 11675547 | + | AACCAGGCTCTGATACCATGA | cca-miR1511 | CTGGTTGAGTGGTTCCACTCGTGGTACCAGGTCCTGCTTCATCAAGTGGTCTTGAGTTCAAATCCAGTCCCAAGCACATGGTTAACCAGGCTCTGATACCATGATGTTTAAATAAACTCAACAG |
| cajca.ICPL87119.gnm1.Cc11 | 20746999 | 20747124 | + | TTGAGCCGCGCCAATATCACT | cca-miR171 | GATATGGGAAACAGAGAAAGCGATGTTGGTGAGGTTCAATCCGAAGACGGATTTACATGTAGAAGCAGTAAAGTACGATCTCAGATTGAGCCGCGCCAATATCACTTTTATTATTGATGGTGGTTG |
| cajca.ICPL87119.gnm1.Cc11 | 32360792 | 32360916 | - | TCGATAAACCTCTGCATCCAG | cca-miR162-2 | ATTTTCGTGGTGAAGTCACTGGAGGCAGTGGTTCATCGATCTCTTCCTGAGTTTGGCCGTGGAAGAACACGAATCAGGAATCGGTCGATAAACCTCTGCATCCAGTGCGAAACCAACACTTTCTT |
| cajca.ICPL87119.gnm1.Cc11 | 41360249 | 41360372 | + | TGACAGAAGAGAGTGAGCAC | cca-miR156c-1 | TGCAGGTTTGTTTGTGGAGGTGACAGAAGAGAGTGAGCACTTAGCTCTTTCCTAGCATGGACCTTTCATGCTAAAAGCTGTGTGCTTACTCTCTATCTGTCACCCACCTTCTTTAATTAGCTCC |
| cajca.ICPL87119.gnm1.Cc01 | 6891831 | 6891949 | - | CGCTATCTATCCTGAGTTTCA | cca-miR390a | AGAGTACAGAAAAATCTGTAAAGCTCAGGAGGGATAGCGCCGCAACACACCAGGATTTTTACATCATGATGTATTTGGCGCTATCTATCCTGAGTTTCACGGGTTCTTCTTACTATATA |
| cajca.ICPL87119.gnm1.Cc02 | 28518502 | 28518623 | - | TGCCTGGCTCCCTGTATGCCA | cca-miR160 | CATGCATACATATGTGTATGTGCCTGGCTCCCTGTATGCCATTTGCAGAGCTCATCGAAGTATCAATGACCTCCGTGGATGGCGTATGAGGAGCCAAGCATATTCCATGTACATACATGGAG |
| cajca.ICPL87119.gnm1.Cc02 | 31347044 | 31347154 | - | TGAAGCTGCCAGCATGATCTT | cca-miR167 | CATCATGCAGCACTACCGGTTGAAGCTGCCAGCATGATCTTAACTTCCCTCACTTGGTTGAGGAAAGATCAGATCATGTGGCAGTTTCACCTGGTAGTTGTTGGCCGCATG |
| cajca.ICPL87119.gnm1.Scaffold 130121 | 191758 | 191854 | - | TGCAAGCTTAGAATCCTCTCTC | cca-miR-Novel1 | GGGGTTATGTGAACTGAAGAGGGAGGGTACTGGTTTGTGGAGGCAGTGACACTTCTGCAAGCTTAGAATCCTCTCTCCGATTTCTTTTTTAAATGAA |
| cajca.ICPL87119.gnm1.Scaffold 134929 | 36809 | 36938 | + | TCGCTTGGTGCAGGTCGGGAA | cca-miR168 | CTCACCGTGCGGTCTCTAATTCGCTTGGTGCAGGTCGGGAACCGGTTTTCTCGCACAATGGAGGAGCGGTCTCTGGCGGCGAATTGGATCCCGCCTTGCATCAACTGAATCGGAGGCCGCCGTGAATGTT |
| cajca.ICPL87119.gnm1.Scaffold 000115 | 23340 | 23468 | + | TGACAGAAGAGAGTGAGCAC | cca-miR156c-2 | AGGTCAAAGGGTAAGGGAGGTGACAGAAGAGAGTGAGCACACATGGCACTTTCTTGCACGATGACGATTCATGCTTGAAGTTATGCGTGCTCACTCTCTATCTGTCACCCCACCACCATCTCTCAAATC |
| cajca.ICPL87119.gnm1.Scaffold 133489 | 279650 | 279772 | - | TGACAGAAGAGAGTGAGCAC | cca-miR156c-3 | ATTTCTTGGGACATAGAAATTGACAGAAGAGAGTGAGCACACAGAGGCACTTGGTATAGTTATATACTGTTGCTTTTGCGTGCTCACTTCTCTTTCTGTCAACTTCCAGTGCTGGAAATTACT |
| cajca.ICPL87119.gnm1.Scaffold 000045 | 273755 | 273897 | + | TTCCACAGCTTTCTTGAACT | cca-miR396a | TCATGGCCCTCTTTGTATTCTTCCACAGCTTTCTTGAACTGCATCCATAGAGTTCCTTTGCATGCATGCCATGGCACTCTTGCTCCCACACCTTGTTTTGCGGTTCAATAAAGCTGTGGGAAGATACAGATAGGGTCAACACT |
| cajca.ICPL87119.gnm1.Scaffold 000045 | 282375 | 282492 | - | TTCCACAGCTTTCTTGAACTT | cca-miR396b | CTCAAGTCCTGGTCATGCTTTTCCACAGCTTTCTTGAACTTCTTATGCCTCCACCTCCCGGATTCTAAGCCCTAGAAGCTCAAGAAAGCTGTGGGAGAATATGGCAATTCAGGCTTCT |
| cajca.ICPL87119.gnm1.Scaffold 133676 | 126183 | 126307 | + | CGCTATCCATCCTGAGTTTCA | cca-miR390b | TTAGTGTGGAAGAATCTGTAAAGCTCAGGAGGGATAGCGCCATGGATGGTCTTCTTCTCTCCCTCCATTCTTGATCCTCTCTCGCGCTATCCATCCTGAGTTTCATGGCTTCTATCCACACACAC |
| ***Glycine max*** | | | | | | |
| 1 | 7214748 | 7214878 | - | TGTGTTCTCAGGTCACCCCTT | gma-miR398 | TCCATATGGTTTATCTCGGAGGAGTGAATCTGAGAACACAAGGCTGGTTTGCACTGCTATATCATCTATTGGTATAAGGTGAATTTACTTTGTGTTCTCAGGTCACCCCTTTGAGCCAACCTGTTGACATA |
| 1 | 48944799 | 48944928 | - | TCGCTTGGTGCAGGTCGGGAA | gma-miR168 | CTCACTGTGCGGTCTCTAATTCGCTTGGTGCAGGTCGGGAACCGGTTTTCGCGCGGAATGGAGGAACGGTCGCCGGCGGCGAATTGGATCCCGCCTTGCATCAACTGAATCGGAGGCCGCGGTGAAGCTT |
| 2 | 1754553 | 1754680 | + | TGAAGTGTTTGGGGGAACTCC | gma-miR395 | ATTTGTTGGCTGTCTCTTGGAGTTCCTCTGAATGCTTCATATATTAGGGACTAGTCTAAGACTCATTAGAAAAAGTTAGTCCTTCAATGAAGTGTTTGGGGGAACTCCTGGATTCAACCACCATTGCA |
| 4 | 4306337 | 4306465 | + | TGACAGAAGAGAGTGAGCAC | gma-miR156a-1 | AAGTCAAAGGGTAAGGGAGGTGACAGAAGAGAGTGAGCACACATAATACTTTCTTGTAAGATAAAGATTCATGCTTGAAGTTATGTGTGCTCACTCTCTATCTGTCACCCCACCACCATCTCTCACCAC |
| 4 | 50116123 | 50116232 | + | CGAGCCGAATCAATATCACCC | gma-miR171a | TCTGCAAAAGTAGACACGGCGTGATATTGGTACGGCTCATCTTAATTCAACCATTATATCAACAATTGAGACGAGCCGAATCAATATCACTCTTGTTTGCTTCATTGCAT |
| 5 | 39506581 | 39506788 | - | TTGGACTGAAGGGAGCTCCCT | gma-miR319 | GTTGAAGACCCTAAGGTAAGAGAGCTTTCTTCAGTCCACTCATGGGTGACAGTAAGATTCAATTAGCTGCCGACTCATTCATCCAAATGTTGAGTGTAAGCGAATAAATATACTCAGCAGATGAGTGAATGATGCGGGAGACAAATTGAATCTTAAGTTTCCTGTACTTGGACTGAAGGGAGCTCCCTTTTCCTTTTGTCTCTTACTT |
| 9 | 40438035 | 40438173 | - | TGCTCTCTAGACTTCTGTCATC | gma-miR156b-1 | GATCTAGTTGATAGGGTTGTTGACAGAAGATAGAGAGCACAGATAGTGATATGCATAAAAATATGGAACGGGAAAGCAATTGCATCTCACTCCTTTGTGCTCTCTAGGCTTCTGTCATCCACACCTTCACTGCCCCTCC |
| 10 | 47144842 | 47144951 | - | TGAAGCTGCCAGCATGATCTT | gma-miR167 | CATCATGCACCACTACCAGTTGAAGCTGCCAGCATGATCTTAACTTCCCTCACTTGCTGTGGAAAGATCAGATCATGTGGCAGTTTCACCTAGTAGTTGTTGGCCGCATG |
| 10 | 49169702 | 49169828 | - | TTACCGATTCCACCCATTCCTA | gma-miR2118-1 | AAGAGCTTGAGGAAGTGATGGGAGATGGGAGGGTCGGTAAAGGATAACAGCGTCTCTATGATTAATTGTTGTGTTGTTTATTCTTTTGCCGATTCCACCCATTCCTATGATTTTCTTTGGTTCCTTT |
| 13 | 27458218 | 27458343 | + | TTGAGCCGCGCCAATATCACT | gma-miR171b-1 | GATATGGGAAGCAGAGAAAGCGATGTTGGTGAGGTTCAATCCGAAGACGGATTTACATGTAGAAGCAGTAAAATACGATCTCAGATTGAGCCGCGCCAATATCACTTTATCATTGATGAGCCCCCC |
| 14 | 24007968 | 24008111 | + | TTCCACAACTTTCTTGAACT | gma-miR396 | TCATGGCCCTCTTTGTATTCTTCAACAACTTTCTTGAACTGCATCCATAGAGTTCCTTTGCATGCATGCCAAGGAACTCTTGCTCCTACACCTTGTTTTGTGGTTCAATAAAGCTATGGGAAGATACAGATAGGGTCAACAACA |
| 17 | 8830879 | 8831004 | - | TTGAGCCGCGCCAATATCACT | gma-miR171b-2 | GATATGGGAAGCAGAGAAAGCGATGTTGGTGAGGTTCAATCCGAAGACGGATTTACATGTAGAAGCAGTAAAATACGATCTCAGATTGAGCCGCGCCAATATCACTTTATCATTGATGAGCTCCCC |
| 17 | 38139703 | 38139833 | - | TGACAGAAGAGAGTGAGCAC | gma-miR156a-2 | AATATTAAGGGTAAGGGAGGTGACAGAAGAGAGTGAGCACACATAGTACTTTCTTGAATGATATAAGGTTTCATGCTTGAAGTTATGCGTGCTTACTCTCTATCTGTCACCCATCACCATCTCTTTCTTTT |
| 18 | 20910821 | 20910943 | + | AACCAGGCTCTGATACCATGA | gma-miR1511 | TGGGTGGAATGGTTTCAGCCGTGGTATCAGGTCCTGCTTCATCAAGTGGTCTTGTGTTCAAATCCAGCCTCAAGCACATGGTTAACCAGGCTCTGATACCATGGTGAATATAATTATCTTCCA |
| 18 | 49001203 | 49001348 | + | AAGCTCGGGAGGGATAGCGCC | gma-miR390 | TCAGTGTGGAAGAATTTGTAAAGCTCAGGAGGGATAGCGCCATGGAAGGTCTTTCTACTAATTTGCTCTATCATGGGTGCTTTGGTTTCGATCGATCTTCTCTAGCGCTACCCATCCTGAGTTTCATGGTTTCTTCATACTTTTCA |
| 18 | 57144434 | 57144558 | - | TGCTCTCTAGACTTCTGTCATC | gma-miR156b-2 | GGAGGTGTTGGTGATGCTGCTGACAGAAGATAGAGAGCACTGATGATGAAATGCATGAAAGGGAATGGCATCTCA |

| Chr | Precursor start | Precursor end | Strand | Mature miRNA sequence | Mature miRNA | Precursor sequence |
|---|---|---|---|---|---|---|
| | | | | | | CTCCTTTGTGCTCTCTAGTCTTCTGTCATCATCCTTCTCCCTCCCCTCTC |
| 19 | 4558497 | 4558593 | - | TGCAAGCTTAGAATCCTCTCTC | gma-miR-Novel1 | AGGGTTATGTGAACTGAAGAGGGAGGGTACTGGTTTGTGGAGGCAGTGACACTTTTGCAAGCTTAGAATCCTCTATCCTAGTTTTTTGTGAAAACAA |
| 20 | 36444870 | 36444996 | + | TTACCGATTCCACCCATTCCTA | gma-miR2118-2 | AAGAGCTTGAGGAAGTGATGGGAGATGGGAGGGTCGGTAAAGAATATATCTGAGACTCGACTCAATCTCGATCTCTCTCAGTGTTGTGTTGTTTTGTTTATCCTTTTGCCGATTCCACCCATTCCTA |
| 20 | 41672359 | 41672479 | + | TGCCTGGCTCCCTGTATGCCA | gma-miR160 | CATGCATACATATGTGTATGTGCCTGGCTCCCTGTATGCCATTTGCATAGTTCATTGAGCATCAATGATCTTTGTGAATGGCGTATGAGGAGCCAAGCATATTCCATGTACATTCATGGAG |
| ***Glycine soja*** | | | | | | |
| glyso.W05.gnm1.Chr01 | 49466901 | 49467030 | - | TCGCTTGGTGCAGGTCGGGAA | gso-miR168 | CTCACTGTGCGGTCTCTAATTCGCTTGGTGCAGGTCGGGAACCGGTTTTCGCGCGGAATGGAGGAACGGTCGCCGGCGGCGAATTGGATCCCGCCTTGCATCAACTGAATCGGAGGCCGCGGTGAAGCTT |
| glyso.W05.gnm1.Chr02 | 1737153 | 1737280 | + | TGAAGTGTTTGGGGGAACTCC | gso-miR395 | ATTTGTTGGCTGTCTCTTGGAGTTCCTCTGAATGCTTCATATATTAGGGACTAGTCTAAGACTCATTAGAAAAAGTTAGTCCTTCAATGAAGTGTTTGGGGGAACTCCTGGATTCAACCACCATTGCA |
| glyso.W05.gnm1.Chr04 | 4299929 | 4300057 | + | TGACAGAAGAGAGTGAGCAC | gso-miR156a-1 | AAGTCAAAGGGTAAGGGAGGTGACAGAAGAGAGTGAGCACACATAATACTTTCTTGTAAGATAAAGATTCATGCTTGAAGTTATGTGTGCTCACTCTCTATCTGTCACCCCACCACCATCTCTCACCAC |
| glyso.W05.gnm1.Chr06 | 10960238 | 10960349 | - | CGAGCCGAATCAATATCACCC | gso-miR171a | TCTGCAAAAGTAGACACGGCGTGATATTGGTACGGCTCATCTTAATTCAACCACTATATCAACGATTGAGACGAGCCGAATCAATATCACTCTTGTTTGCTTCATTGCATAT |
| glyso.W05.gnm1.Chr07 | 4554465 | 4554609 | - | TCGGACCAGGCTTCATTCCCC | gso-miR166-1 | GGAAGCTTTGTGTTTTGAGGGGAATGTTGTCTGGCTCGAGGACCCTTCTTCTTCTTGATCTTGTGTAGACTGCTATACCTGTGGTCAAGGAATGCTTAGTGTTTTCGGACCAGGCTTCATTCCCCCCAATTATATGCTTCCAAAT |
| glyso.W05.gnm1.Chr09 | 38894253 | 38894391 | - | TGCTCTCTAGACTTCTGTCATC | gso-miR156b-1 | GATCTAGTTGATAGGGTTGTTGACAGAAGATAGAGAGCACAGATAGTGATATGCATAAAAATATGGAACGGGAAAGCAATTGCATCTCACTCCTTTGTGCTCTCTAGGCTTCTGTCATCCACACCTTCACTGCCCCTCC |
| glyso.W05.gnm1.Chr10 | 3015344 | 3015468 | - | TCGGACCAGGCTTCATTCCCC | gso-miR166-2 | ATATACTTAAAAAGTTGAGGGGAATGTCGTTTGGTTCGAGATCATTCATGCAAGTAGTCTCAGACATAACTCTTCTGAGTGATTTCGGACCAGGCTTCATTCCCCTCAGCTACCCAGCAGACACT |
| glyso.W05.gnm1.Chr10 | 45967368 | 45967489 | - | TGCCTGGCTCCCTGTATGCCA | gso-miR160 | CATGCATACATATGTGTATGTGCCTGGCTCCCTGTATGCCATTTGTAGAGCTCATCGAAGCATCAATGACCTTTGTGGATGGCGTATGAGGAGCCAAGCATATTTCATATACATACATGGAG |
| glyso.W05.gnm1.Chr10 | 50777203 | 50777329 | - | TTACCGATTCCACCCATTCCTA | gso-miR2118-1 | AAGAGCTTGAGGAAGTGATGGGAGATGGGAGGGTCGGTAAAGGATAACAGCGTCTCTATTATTAATTGTTGTGTTGTTTATTCTTTTGCCGATTCCACCCATTCCTATGATTTTCTTTGGTTCCTTT |
| glyso.W05.gnm1.Chr11 | 35127616 | 35127741 | + | TTGAGCCGCGCCAATATCACT | gso-miR171b-1 | GATATGGGAAGCAGAGAAAGCGATGTTGGTGAGGTTCAATCCGAAGACGGATTTACATGTAGAAGCAGTAAAATACGATCTCAGATTGAGCCGCGCCAATATCACTTTATCATTGATGAGCCCCCC |
| glyso.W05.gnm1.Chr11 | 35186082 | 35186210 | + | TTCCACAGCTTTCTTGAACTT | gso-miR396a-1 | CTCAAGTCCTGGTCATGCTTTTCCACAGCTTTCTTGAACTTCTTATGCATCTTATATCTCTCCACCTCCAGGATTTTAAGCCCTAGAAGCTCAAGAAAGCTGTGGGAGAATATGGCAATTCAGGCTTTT |
| glyso.W05.gnm1.Chr17 | 8800653 | 8800781 | - | TTCCACAGCTTTCTTGAACTT | gso-miR396a-2 | CTCAAGTCCTGGTCATGCTTTTCCACAGCTTTCTTGAACTTCTTATGCATCTTATATCTCTCCACTTCCAGCATTTTAAGCCCTAGAAGCTCAAGAAAGCTGTGGGAGAATATGGCAATTCAGGCTTTT |
| glyso.W05.gnm1.Chr17 | 8849411 | 8849536 | - | TTGAGCCGCGCCAATATCACT | gso-miR171b-2 | GATATGGGAAGCAGAGAAAGCGATGTTGGTGAGGTTCAATCCGAAGACGGATTTACATGTAGAAGCAGTAAAATACGATCTCAGATTGAGCCGCGCCAATATCACTTTATCATTGATGAGCTCCCC |
| glyso.W05.gnm1.Chr17 | 38310519 | 38310649 | - | TGACAGAAGAGAGTGAGCAC | gso-miR156a-2 | AATATTAAGGGTAAGGGAGGTGACAGAAGAGAGTGAGCACACATAGTACTTTCTTGAATGATATAAAGTTTCATGCTTGAAGTTATGCGTGCTTACTCTCTATCTGTCACCCCATCACCATCTCTTTCTTT |
| glyso.W05.gnm1.Chr18 | 48638064 | 48638208 | + | CGCTATCCATCCTGAGTTTCA | gso-miR390 | TCAGTGTGGAAGAATTTGTAAAGCTCAGGAGGGATAGCGCCATGGAAGGTCTTTCTACTAATTTGCTCTATCATGGGTGCTTTGGTTCCGATCATCTTCTCTAGCGCTATCCATCCTGAGTTTCATGGTTTCTTCATACTTTTCA |
| glyso.W05.gnm1.Chr18 | 56871643 | 56871767 | - | TGCTCTCTAGACTTCTGTCATC | gso-miR156b-2 | GGAGGTGTTGGTGATGCTGTTGACAGAAGATAGAGAGCACTGATGATGAAATGCATGAAAGGGAATGGCATCTCACTCCTTTGTGCTCTCTAGTCTTCTGTCATCATCCTTCTCCCTCCCCTCTC |
| glyso.W05.gnm1.Chr19 | 4573298 | 4573394 | - | TGCAAGCTTAGAATCCTCTCTC | gso-miR-Novel1 | AGGGTTATGTGAACTGAAGAGGGAGGGTACTGGTTTGTGGAGGCAGTGACACTTTTGCAAGCTTAGAATCCTCTATCCTAGTTTTTTGTGAAAACAA |
| glyso.W05.gnm1.Chr20 | 39442568 | 39442694 | + | TTACCGATTCCACCCATTCCTA | gso-miR2118-2 | AAGAGCTTGAGGAAGTGATGGGAGATGGGAGGGTCGGTAAAGAATATATCTGAGACTCAACTCAATCTCGATCTCTCTCAGTGTTGTGTTGTTTTGTTTATCCTTTTGCCGATTCCACCCATTCCTA |
| glyso.W05.gnm1.Chr20 | 42067591 | 42067700 | + | TGAAGCTGCCAGCATGATCTT | gso-miR167 | CATCATGCACCACTACCAGTTGAAGCTGCCAGCATGATCTTAACTTCCCTCACTTGCCGTGGAAAGATCAGATCATGTGGCAGTTTCACCTAGTAGTTGCTGGCCGCATG |
| glyso.W05.gnm1.tig00104511_1_pilon_84547_1256723 | 1162286 | 1162429 | + | TTCAACAGCTTTCTTGAACT | gso-miR396b | TCATGGCCCTCTTTGTATTCTTCAACAACTTTCTTGAACTGCATCCATAGAGTTCCTTTGCATGCATGCCAAGGAACTCTTGCTCCTACACCTTGTTTTGTGGTTCAATAAAGCTATGGGAAGATACAGATAGGGTCAACAACA |
| ***Lupinus angustifolius*** | | | | | | |
| LG01 | 1781448 | 1781573 | + | CGATGTTGGTGAGGTTCAATC | lan-miR171a-1 | GATAATGGAAGCAGAAAAAGCGATGTTGGTGAGGTTCAATCCGAATACGGATTTACATTGTATAAGCAGTAAATACGATCTTAGATTGAGCCGCGCCAATATCACTACTTGATGACTTTCCACCTC |
| LG01 | 1793877 | 1794005 | + | TTCCACAGCTTTCTTGAACTT | lan-miR396a-1 | CTCAAGTCCTGGTCATACTTTTCCACAGCTTTCTTGAACTTCTTATGCATCTTAGTGTTCTTCATGTCCAGAATTTGAAGCCCTAGAAGTTCAAGAAAGCTGTGGGAGAATATGGCAACTCAGGCTTGG |
| LG01 | 20975320 | 20975437 | - | TCGATAAACCTCTGCATCCAG | lan-miR162 | TCGATCGAGGTGAAGTCTCTGGAGGCAGCGGTTTATCGATCTCTTCTTGAATTTTGTTAACACAAACCATGAATCGATCGATAAACCTCTGCATCCAGCGCTCACTTTCATTCTCTCT |
| LG01 | 21390226 | 21390352 | + | CGATGTTGGTGAGGTTCAATC | lan-miR171a-2 | AATAAGGGAAGCAGAGAAAGCGATGTTGGTGAGGTTCAATCCGAAGACGGATTTACATGATAGAAGCAGTAAAATACGATCTCAGATTGAGCCGTGCCAATATCACTTTTATGGTTGAAAAACTCTC |
| LG01 | 34141970 | 34142095 | - | GGAATGTTGGCTGGCTCGAGG | lan-miR166a | GATGGGTTTTTTGTTGAGGGGAATGTTGGCTGGCTCGAGGCTTTTACAAATCAATGTTCGTAGTTTTGAAGAACAATAAGGCTTCGGACCAGGCTTCATTCCCCTCAAAACTTGTTTGCTTTTG |
| LG05 | 24390027 | 24390171 | + | TTGACAGAAGATAGAGAGCAC | lan-miR156a | GGCACTAGTTAGTGAGATTGTTGACAGAAGATAGAGAGCACAGATGATGTTATGCATATTATATATACATGGAGCTGAATAGTGTTGCATCTCACTCCTTTGTGCTCTCTATGCTTCTGTCATCATCTTCACCCTCCATTTCTCA |
| LG08 | 9166410 | 9166535 | - | TCGGACCAGGCTTCATTCCCC | lan-miR166b-1 | TGTTACATAAAAGGTTGAGGGGAATGTCGTCTGGCTCGAGATCATCCATGCAAGTAGCCTCAGGCATAACTCTTGATGAGTGATTTCGGACCAGGCTTCATTCCCCTCAACTACCAAACAGACACT |
| LG09 | 19501052 | 19501173 | + | AAGCTCGGGAGGGATAGCGCC | lan-miR390-1 | AGAGTATGGAAGAATCTGTTAAGCTCAGGAGGGATAGCGCCGCCACGATACTAATGATTTTATCATAGTACTATCCTTTGGCGCTGTCCATCCTGAGTTTCACGGCTTCTTCTTACTGTTTA |

| Chr | Precursor start | Precursor end | Strand | Mature miRNA sequence | Mature miRNA | Precursor sequence |
|---|---|---|---|---|---|---|
| LG12 | 3923031 | 3923165 | - | AAGCTCGGGAGGGATAGCGCC | lan-miR390-2 | GAAAGAATCTGTTAAGCTCAGGAGGGATAGCGCCATGGATGGTCTTCTTTTTTGCTCCATTTTCTGGTTTGCATTTTAATTTGATCTTCTATTGTGCTATCCATCCTGAGTTTCATGGCTTTCTCACACTCTCAT |
| LG12 | 17799192 | 17799320 | - | TTCCACAGCTTTCTTGAACTT | lan-miR396a-2 | CTCAAGCCTTGGTCATACTTTTCCACAGCTTTCTTGAACTTCTTATGCATTTTAAAGTTCTCCATCTTCAGAATTTAAAGCCCTAGAAGCTCAAGAAAGCTGTGGGAGAATATGGCAATTTAGGCTTTG |
| LG12 | 17811667 | 17811793 | - | TTGAGCCGCGCCAATATCACT | lan-miR171b | ACAAAGGGAAGTAAAGAAAGCGATGTTGGTGAGGTTCAATCCGAAGACGGATTTACATTGTGGAAACGGTAAAATACGATCTCAGATTGAGCCGCGCCAATATCACTTTAATGTTGATGATTTTCCT |
| LG14 | 4107954 | 4108084 | + | TTCCACAGCTTTCTTGAACTG | lan-miR396b | TCCTGACCCTCTTTGTATTCTTCCACAGCTTTCTTGAACTGCATCTAAGAGTTCCGTTGTATGCATGCCATGGCACTCTCAAACACTGCGGTTCAATAAAGTTGTGGGAAGATACAGAATAGGGTCAAGTG |
| LG15 | 19919302 | 19919430 | - | TGACAGAAGAGAGTGAGCAC | lan-miR156b | TTTGTTAGGGGTAAGGGAGGTGACAGAAGAGAGTGAGCACACATGGTGTTTTCTTGCATGATGATGTTTAATGCTTGAAGCTATGTGTGCTTACTCTCTATCTGTCACCCCAATCCTCATCTCTTTCTC |
| LG17 | 17019050 | 17019180 | - | TCGGACCAGGCTTCATTCCCC | lan-miR166b-2 | AGGAAGCTTTCTTTTTTAGGGGAATGTTGTCTGGCTCGAGGACTCTTTCATCTTTAATCAATATACTTGATTAAGAAATTTATAGTGTTGTCGGACCAGGCTTCATTCCCCTGAACTATAATCTTCCGGAT |
| LG17 | 17641733 | 17641856 | - | AACCAGGCTCTGATACCATGA | lan-miR1511 | TTGAAGAAGTGGTGTATTTCGTGGTATCAGGTCCTGCTTCACCAAGTGGTCTTGAGTTCAATTCCTGTCCCAAGCACATGGTTAACCAGGCTCTGATACCATGATGGATTAACCCAATCTCAAA |
| LG20 | 5701638 | 5701752 | + | TGAAGCTGCCAGCATGATCTGA | lan-miR167 | TATGTTGCCCTTGAAATAGTTGAAGCTGCCAGCATGATCTGATGTTACCTTGTATCTTTTGTTACGGTAAGAATAGATCATGTGGCAGTTTCACCTGTTGAATGGAAGCATAGAA |
| ***Lotus japonicus,*** | | | | | | |
| lotja.MG20.gnm3.Lj0 | 60412133 | 60412249 | + | TGGAGAAGCAGGGCACGTGCA | lja-miR164 | GATTGGTGAGTAGGTCTTGTTGGAGAAGCAGGGCACGTGCAATTTTCTCTCATCACAAGCTCAAACCTTGAATTTGCACGTGCTCCCCTTTTCCAACATGATTTTCCCACCATGTTC |
| lotja.MG20.gnm3.Lj0 | 130771215 | 130771339 | - | TCGATAAACCTCTGCATCCAG | lja-mir162 | AGAGAGGCGACAAAGTCGCTGGACGCAGCGGTTCATCGATCTCTTCCTGAATTTGGCTGTGGAAGAACACAAAACAAGAATCGGTCGATAAACCTCTGCATCCAGCGCTCATCTCGCCTCTCTCT |
| lotja.MG20.gnm3.Lj0 | 151870758 | 151870902 | + | TGCTCTCTAGACTTCTGTCATC | lja-mir156a | TAGCTAGTGAGCAAGGTTGTTGACAGAAGATAGAGAGCACAGATGATGATATGCTTATACATATATACATACACATGGAGATGGAGATGAAGCACCATTGCATGTCATTCCTTTGTGCTCTCTAGGCTTCTGTCATCACCTTGAG |
| lotja.MG20.gnm3.Lj1 | 1227992 | 1228100 | + | TGAAGCTGCCAGCATGATCTT | lja-mir167-1 | CATCATGCACCACTACCAGTTGAAGCTGCCAGCATGATCTTAACTTCCCTCCATGGTGGGGAAAGATCAGATCATGTGGCAGTTTCACCTTGTTGTTGGTTGCATGAAC |
| lotja.MG20.gnm3.Lj2 | 24691406 | 24691561 | - | TCGCTTGGTGCAGGTCGGGAA | lja-mir168 | CTCGCTTCGCGGTCTCTAATTCGCTTGGTGCAGGTCGGGAACCGCTGCTGCTTCTGCCGTTTTCGCGCGTAATCACTCACCGGAGGAGCGGTTACCGCCGCCGGTTGGATCGGATCCCGCCTTGCATCAACTGAATCGGAACCCGCCTTGAAGCTT |
| lotja.MG20.gnm3.Lj2 | 33780587 | 33780716 | - | CGCTATCCATCCTGAGTTTCA | lja-mir390 | AGAGTATGGGAGAATCTGTTAAGCTCAGGAGGGATAGCGCCATGATATATATTGATGCAGTTTCTATTGTCTTCAATATTGGTCTTTGGCGCTATCCATCCTGAGTTTCATGGGTTCTTCTTACTGTTTA |
| lotja.MG20.gnm3.Lj4 | 26573725 | 26573850 | + | TTGAGCCGCGCCAATATCACT | lja-mir171-1 | GATTTGGGAAGCAGAGAAAGCGATGTTGGTGAGGTTCAATCTGAAGACGGATTTACATGAAGAAGCAGTAAAATACGATCTCAGATTGAGCCGCGCCAATATCACTTTTCTTATTCATGAGCTCTG |
| lotja.MG20.gnm3.Lj4 | 26634655 | 26634785 | + | TTCCACAGCTTTCTTGAACTT | lja-mir396a | CACAAGTCCTGGTCATGCTTTTCCACAGCTTTCTTGAACTTCTTGTGTGCATATTATTCAAAGCTATTTCCAGAATTAACAGCCCTAGAAGCTCAAGAAAGCTGTGGGAGAATATGGCAGCTCAGACTTTT |
| lotja.MG20.gnm3.Lj4 | 26642888 | 26643035 | - | GTTCAATAAAGCTGTGGGAAG | lja-mir396b-1 | TCATGGCTCTCTTTGTATTCTTCCACAGCTTTCTTGAACTGCATCCAAAGAGTGATGTACTTTTCATGCATATATGCCATGACACCTCTAATTACACTTGTTTTGCGGTTCAATAAAGCTGTGGGAAGATACAGATAGGGTCAACTGC |
| lotja.MG20.gnm3.Lj4 | 38041447 | 38041579 | - | TCGGACCAGGCTTCATTCCCC | lja-mir166 | TTGCAAAAGTTAGGCTGAGAGGAACGTTGTCTGGCTCGAGGTGTCAATTCTTTTTCTTCTAGCCTTTCACTGAGAGAAGAATTGATTCATCCTCGGACCAGGCTTCATTCCCCCTAGCCAACTTTTGCTACTT |
| lotja.MG20.gnm3.Lj4 | 41835334 | 41835469 | + | TTGACAGAAGATAGAGAGCAC | lja-mir156b | AAATACTAGTTATGAGGTCGTTGACAGAAGATAGAGAGCACAGATGATGATATGCATGAATATATAAGATGGATTGCGCATTTCACTCCTTTGTGCTCTCTATGCTTCTGTCATCACCTTCAGCCTCTCACTCCCC |
| lotja.MG20.gnm3.Lj5 | 8569138 | 8569258 | - | TGCCTGGCTCCCTGTATGCCA | lja-mir160-1 | GTTTATTAGCATACTTTGTGTGCCTGGCTCCCTGTATGCCATTTGTAGAGCACATCACCACGGTGATGGCCTAATGAATGGCGTACAAGGAGCCAAGCATGCTGTGTGTTGCTATTTAGCT |
| lotja.MG20.gnm3.Lj5 | 26130935 | 26131056 | - | TGCCTGGCTCCCTGTATGCCA | lja-mir160-2 | CATGCATACATATGTGTATGTGCCTGGCTCCCTGTATGCCATTTGGAGAGCTCATCGAAACATCAATGATCTCTTTGGATGGCGTATGAGGAGCCAAGCATATTCCATGTTTCATACATGGA |
| lotja.MG20.gnm3.Lj5 | 28811357 | 28811477 | + | GTTCAATAAAGCTGTGGGAAG | lja-mir396b-2 | TCATGGTCCTCTTTGTATTCTTCCACAGCTTTCTTGAACTGCATCCAATTGCATGCATGCCATGACACTCTTATTTTGCGGTTCAATAAAGCTGTGGGAAGATATAGATAGGGTCAACTGC |
| lotja.MG20.gnm3.Lj5 | 29010588 | 29010696 | - | TGAAGCTGCCAGCATGATCTT | lja-mir167-2 | CATCATGCACCACTACCAGTTGAAGCTGCCAGCATGATCTTAACTTCCCTCCATGGTGGGGAAAGATCAGATCATGTGGCAGTTTCACCTTGTTGTTGGTTGCATGAAC |
| lotja.MG20.gnm3.Lj6 | 5464352 | 5464477 | + | TTGAGCCGCGCCAATATCACT | lja-mir171-2 | GATTTGGGAAGCAGAGAAAGCGATGTTGGTGAGGTTCAATCTGAAGACGGATTTACATGAAGAAGCAGTAAAATACGATCTCAGATTGAGCCGCGCCAATATCACTTTTCTTATTCATGAGCTCTG |
| ***Medicago truncatula*** | | | | | | |
| medtr.A17_HM341.gnm4.chr1 | 4043005 | 4043130 | - | TGACAGAAGAGAGTGAGCAC | mtr-miR156a-1 | TCAATCGAGAGTAAGGGAGGTGACAGAAGAGAGTGAGCACACATGGTACTTTCGTGTATGATGTTTCATTCTCGAAGCTATGTGTGCTCACTCTCTATCTGTCACCCCATCACCATCTTTTTCTTT |
| medtr.A17_HM341.gnm4.chr3 | 26834668 | 26834778 | + | CGCTATCCATCCTGAGTTTCA | mtr-miR390-1 | AGAGTATAGGAGGGTCGGTAAAGCTCAGGAGGGATAGCGCCATTGATAAATGTGTGTACGTGGTATTTGGCGCTATCCATCCTGAGTTTCACCGGTTCTTCTTACTAGCTA |
| medtr.A17_HM341.gnm4.chr3 | 48230211 | 48230339 | + | TGACAGAAGAGAGTGAGCAC | mtr-miR156a-2 | AAGGGTAAGGGTAAGGGTGGTGACAGAAGAGAGTGAGCACACATGGTGTTTTCTTGCACGATTATGTTTCCTGCTTGAAGCTATATGTGCTTACTCTCTATCTGTCACCCCACCACCATCTCTCTCTCT |
| medtr.A17_HM341.gnm4.chr4 | 4052253 | 4052384 | - | CGCTATCCATCCTGAGTTTCA | mtr-miR390-2 | TTTGTGTGAAAGAATCTGTTAAGCTCAGGAGGGATAGCGCCATAGAATGTCTTCTTTTTTTGGTTCCATTTTATTGATTGATCTTCTCTTGCGCTATCCATCCTGAGTTTCATGGCTTCTTCATACTCTCTT |
| medtr.A17_HM341.gnm4.chr4 | 46038008 | 46038132 | - | TCGATAAACCTCTGCATCCAG | mtr-miR162 | ATGAGGTGAAGTTCGTCACTGGATGCAGCGGTTCATCGATCTGTTCCTGAATTTTGTTTGTCTCGTAAAACAAACATGAATCGGTCGATAAACCTCTGCATCCAGCGCTCACTTTCCCCTCTCTT |
| medtr.A17_HM341.gnm4.chr4 | 47495940 | 47496066 | + | TTGAGCCGCGCCAATATCACT | mtr-miR171 | GTTTGGGAAGCCGAAAAAAGCGATGTTGGTGAGGTTCAATCCGAAGACGGATTTACATGTATAGAGTTGTAAAATACGATCTCAGATTGAGCCGCGCCAATATCACTTTTATTATTGATCCTCTTTC |
| medtr.A17_HM341.gnm4.chr4 | 47580901 | 47581025 | + | TTCCACAGCTTTCTTGAACTT | mtr-miR396a | TTCAAGTCCTGGTCATGCTTTTCCACAGCTTTCTTGAACTTCTTTCGTATCTTAAATCTGTTTTCAAGATTAAAAGTCCTAGAAGCTCAAGAAAGCTGTGGGAGAATATGGCAATTCAGGCTTCA |

| Chr | Precursor start | Precursor end | Strand | Mature miRNA sequence | Mature miRNA | Precursor sequence |
|---|---|---|---|---|---|---|
| medtr.A17_HM341.gnm4.chr4 | 47589540 | 47589666 | - | TTCCACAGCTTTCTTGAACTG | mtr-miR396b | ATATGGTCCTCTTTGTATTCTTCCACAGCTTTCTTGAACTGCATCCAAATTGAGTTCCTTTGCATTGCCATGGCCATTGTTTTGCGGTTCAATAAAGCTGTGGGAAGATACAGATAGGGTCAACTAC |
| medtr.A17_HM341.gnm4.chr5 | 10535753 | 10535878 | + | TCGCTTGGTGCAGGTCGGGAA | mtr-miR168 | CTCACTTCGCGGTCTCTAATTCGCTTGGTGCAGGTCGGGAACCACTACATCCGCTGTTTTCGTAGAAACCGGCGGTGAATTGGATCCCGCCTTGCATCAACTGAATCGGAGGCCGCGGTGAATCTA |
| medtr.A17_HM341.gnm4.chr6 | 30617114 | 30617264 | - | TGCTCTCTAGACTTCTGTCATC | mtr-miR156b | AGCTAGTTAAGTAAGGTTGTTGACAGAAGATAGAGGGCACTAAGGATGATATGCATACACATATATATACAACATGGAGGAGGAGCTTAATTGCATTTCATTTCCTTTGTGCTCTCTAGACTTCTGTCATCACCTCATCTTTCTCACCTTT |
| medtr.A17_HM341.gnm4.chr7 | 46417700 | 46417816 | - | TGGAGAAGCAGGGCACGTGCA | mtr-miR164a | CACTGTTGGGTAGGTCTTGTTGGAGAAGCAGGGCACGTGCAACCGTAGATCTCTCTCAAACTTCAATCTCATTTTGCACGTGCTCCACTTTTCCAACTTGATCTTCCCACCATCTTT |
| medtr.A17_HM341.gnm4.chr8 | 9298712 | 9298846 | - | TCGGACCAGGCTTCATTCCCC | mtr-miR166 | GGAAGCTTTATTTTTTGAGGGGAATGTTGTCTGGCTCGAGGACGCTTTCTTCTCGATCTAATGCAAATTTGTGGTCATGGATTGTAAAGTATTCTCGGACCAGGCTTCATTCCCCCCAATTATATGCTTCCATGG |
| medtr.A17_HM341.gnm4.chr8 | 27290673 | 27290795 | + | AACCAGGCTCTGATACCATGA | mtr-miR1511 | TGGGCCGGTTGGTTTCCTTCATGGTGTCAGGACCTGCTTCATCAAGTGGTCTTGAGTTCAACTCCTGTTCAAGCACATGGTTAACCAGGCTCTGATACCATGATTTAGCTAGCAAATGGTCCT |
| medtr.A17_HM341.gnm4.chr8 | 39663053 | 39663184 | - | TCGGACCAGGCTTCATTCCCC | mtr-miR164b | AAGCAAGAGTTAGGTTGAGAGGAACGTTGTCTGGCTCGAGGTGATGGAGATGGAAGAGTACTCTCTACTCACTCATCACTAACTTTCAATCTCGGACCAGGCTTCATTCCCCCCAGCAAACTTTTGCTAGCT |
| ***Oryza sativa*** | | | | | | |
| 2 | 26130059 | 26130189 | + | TCGGACCAGGCTTCATTCCCC | osa-miR166a | GAAGCTTTTTCACTTTGAGGGGAATGTTGTCTGGCTCGAGGTGCATGGAGAAACCTCTGATCGATCAGGTTTGATCTGTAGAGACTGATCTCGGACCAGGCTTCATTCCCCTCAAGTAAAGCTCCCATATT |
| 2 | 32435164 | 32435300 | - | TCGGACCAGGCTTCATTCCTC | osa-miR166b | TTGGCCATGGTGGCTTGTGGGGAATGTTGGCTGGCTCGAGGTATCCACATCTTAATTCCTCTCCGGCGATCGAGCCGGCTCGGGCGTGTGGAAGCGTCGGACCAGGCTTCATTCCTCGCAAGCCGATGCATCCATGG |
| 4 | 25026314 | 25026441 | - | TGACAGAAGAGAGTGAGCAC | osa-miR156 | GGCGGGCCGGTGGCGCGAGGTGACAGAAGAGAGTGAGCACACGGCCGGGCGTGACGGCACCGGCGGGCGTGCCGTCGCGGCCGCGTGCTCACTGCTCTTTCTGTCATCCGGTGCCGGCCGTTTTCTCC |
| 6 | 3669639 | 3669776 | + | TTCCACAGCTTTCTTGAACTG | osa-miR396 | AGATGGTCCTCTTTGTGGTCTTCCACAGCTTTCTTGAACTGCATCTTTGAGAGAGATTAGCATCCCTATGTGTGGATTTTGCTTGCACGAGTGTGCAGTTCAATAAAGCTGTGGGAAATTACAGAGAGAGGTCAATTG |
| 6 | 25891120 | 25891238 | + | TTGAGCCGCGCCAATATCACT | osa-miR171 | TGGTTGATCAATGGAAAGAGCGATATTGGTGAGGTTCAATCCGATGATTGGTTTTACAGCAGTGGTAAAATCAGTATCTGATTGAGCCGCGCCAATATCTCTTCCTCTCCTATATCCTG |
| ***Phaseolus vulgaris*** | | | | | | |
| phavu.G19833.gnm1.Chr01 | 19172736 | 19172866 | + | TGACAGAAGAGAGTGAGCAC | pvu-miR156a-1 | GGAGATTATGGTAAGGGAGGTGACAGAAGAGAGTGAGCACACATGGTACTTTCTTGGATGATATAAGGTTTCATGCTTGAAGCTATGCGTGCTTACTCTCTATCTGTCACCCCATCACCATCTCTTTCTTT |
| phavu.G19833.gnm1.Chr02 | 24876261 | 24876390 | + | TCGCTTGGTGCAGGTCGGGAA | pvu-miR168 | ATCACTGTGCGGTCTCTAATTCGCTTGGTGCAGGTCGGGAACCTGTTTTTGCGCGAAATGGTGGAGTGGTTGCCGGCGGCGAATTGGATCCCGCCTTGCATCAACTGAATCGGAGGCCGCAGTGAAGGTT |
| phavu.G19833.gnm1.Chr02 | 44066921 | 44067044 | - | TCGGACCAGGCTTCATTCCCC | pvu-miR166-1 | GAGATGGGTTGGTATTGAGGGGAATGTTGGCTGGCTCGAGGCTTTTCACAAAGGAGGTTCACAGTGGATGGAACTATAAGGCTTCGGACCAGGCTTCATTCCCCTCAAAGTACTTCCTTCATGG |
| phavu.G19833.gnm1.Chr02 | 48410409 | 48410545 | + | TGCTCTCTAGACTTCTGTCATC | pvu-miR156b-1 | ACTGGTTTTGGTAAGGTTGTTGACAGAAGATAGAGAGCACAGATGATGATATGCATATTACTATATATAGAGAGCATGAACTGATCAATGGTGCATCTTACTCCTTTGTGCTCTCTATACTTCTGTCATCACCTTCA |
| phavu.G19833.gnm1.Chr03 | 37749148 | 37749273 | + | TTGAGCCGCGCCAATATCACT | pvu-miR171 | GATATGGGAAGCAGAGAAAGCGATGTTGGTGAGGTTCAATCCGAAGACGGATTTACATGTAGAAGCAGTAAAATACGATCTCAGATTGAGCCGCGCCAATATCACTTATATCATTGCTGAGTCCCC |
| phavu.G19833.gnm1.Chr03 | 37833718 | 37833846 | + | TTCCACAGCTTTCTTGAACTT | pvu-miR396a | CTCAAGTCCTGGTCATGCTTTTCCACAGCTTTCTTGAACTTCTTCTGCATCTTATATCTCTCCGCCTCCAGGATTTTAAGCCCTAGAAGCTCAAGAAAGCTGTGGGAGAATATGGCAATTCAGGCCTTT |
| phavu.G19833.gnm1.Chr03 | 37841429 | 37841571 | - | TTCCACAGCTTTCTTGAACTG | pvu-miR396b | TCATGGCCCTCTTTGTATTCTTCCACAGCTTTCTTGAACTGCATCCATAGAGTTCCTTTGCATGCATGCCATGGCACTCTTGCTCCCACACCTTGTTTTGCGGTTCAATAAAGCTGTGGGAAGATACAGATAGGGTCAACAAC |
| phavu.G19833.gnm1.Chr03 | 39163873 | 39163997 | + | TCGATAAACCTCTGCATCCAG | pvu-miR162 | GGAGATGAGGTGAAGACACTGGAGGCAGCGGTTCATCGATCTCTTCCTGAATTTGGCTGTGGAAGAACACAAAACAAGAATCGGTCGATAAACCTCTGCATCCAGCGCTCACTTTGCCTCTCTTT |
| phavu.G19833.gnm1.Chr06 | 19301533 | 19301642 | + | CGCTATCTATCCTGAGTTTCA | pvu-miR390a | AGAGTACAGAAGAATCTGTAAAGCTCAGGAGGGATAGCGCCATGATTCTCACATCATAGTGTTCTTTGGCGCTATCTATCCTGAGTTTCACGGCTTCTTCTTACTGTATA |
| phavu.G19833.gnm1.Chr07 | 5470895 | 5471005 | + | TGAAGCTGCCAGCATGATCTT | pvu-miR167 | CATCGTGCACCACTACCAGTTGAAGCTGCCAGCATGATCTTAACTTCCCTCACTTGGTTGAGGAGAGATCAGATCATGTGGCAGTTTCACCTAGTTGTTGGCCGCATGAAC |
| phavu.G19833.gnm1.Chr09 | 12837439 | 12837567 | + | TGACAGAAGAGAGTGAGCAC | pvu-miR156a-2 | AAGTTTAAGGGTAAGGGAGGTGACAGAAGAGAGTGAGCACAGATGGTATTTTCTTGCAAGATAACGATTTATGCTTGAAGCTATGCGTGCTTACTCTCTATCTGTCACCCCACCACCATCTCCACCCTT |
| phavu.G19833.gnm1.Chr09 | 13696970 | 13697092 | - | TGACAGAAGAGAGTGAGCAC | pvu-miR156a-3 | CTTTCTTGGAACATAGAAATTGACAGAAGAGAGTGAGCACACAGAGGCACCTGGTATAGTTATATGCTGTTACTTTTGCGTGCTCACTTCTCTTTCTGTCAACTTCCAGTGCTGGAAATTACT |
| phavu.G19833.gnm1.Chr09 | 22103444 | 22103567 | + | TCGGACCAGGCTTCATTCCCC | pvu-miR166-2 | TGGGTTAGTAGTGTTTGAGGGGAATGTTGGCTGGCTCGAGGCTTTTCACATAGGAGGTTCTCACTGGCAAGAACTATAAGGCTTCGGACCAGGCTTCATTCCCCTCAAAACTTTTCTTTTTGTG |
| phavu.G19833.gnm1.Chr10 | 12464234 | 12464376 | - | CGCTATCCATCCTGAGTTTCA | pvu-miR390b | TTAGTGTAGAAGAATCTGTAAAGCTCAGGAGGGATAGCGCCATGGATGGTCATTGCTCATCTCTCTTCTCTTTCCTCCCTTTGCCTCTTTGATCTTCTCTTGCGCTATCCATCCTGAGTTTCATGGCTTCTTTCTACACACAT |
| phavu.G19833.gnm1.scaffold_24 | 150445 | 150583 | - | TGCTCTCTAGACTTCTGTCATC | pvu-miR156b-2 | GTTTGGGGAAAGAGGATTGTTGACAGAAGATAGAGAGCACAGATGATGATATGCACAAACATATGGAACGGGAAAGCAATTGTGTCTCACTCCTTTGTGCTCTCTAGGCTTCTGTCATCCACCTTCACTACCCTCCATT |
| ***Trifolium pratense*** | | | | | | |
| LG1 | 9523129 | 9523256 | - | TCGATAAACCTCTGCATCCAG | tpr-miR162 | AGAGATGAGGTGAAGTCACTGGAGGCAGCGGTTCATCGATCTGTTCCTGAATTTGTTTGTGTAATTATTTCACAAAACAAGAATCGGTCGATAAACCTCTGCATCCAGCGCTCACTTTCCCCTCTCTG |
| LG1 | 16688066 | 16688194 | - | TGACAGAAGAGAGTGAGCAC | tpr-miR156a | TCAATCGAGGGTAAGGGAGGTGACAGAAGAGAGTGAGCACACATGGTATTTTCTTGTATGATGATGTTTCATACTTGAAGCTATGCGTGCTCACTCTCTATCTGTCACCCCATCACCATCTTTTTCTTT |
| LG2 | 5731651 | 5731773 | - | CGAGCCGAATCAATATCACCC | tpr-miR171a | TTCTGCAGAAGTAGACATGAGTGATATTGTTTCTGTTCATCTTATTAATTAATGAATGAAACACATTTGCTTAATTATAAGACGAGCCGAATCAATATCACTCGAGTACACTTCACTGCATAT |
| LG2 | 25691458 | 25691587 | + | TCGGACCAGGCTTCATTCCCG | tpr-miR166a | AAGGGAGATTATGGTTGATGGGAATGTTGTTTGGCTCGAGGTAACTAAGAGAGGTTCAAAGTAAGGTTTTTGATTTTGAAGATGTGATCTCGGACCAGGCTTCATTCCCGTCAACTTGATCTCTCAGGAT |

| Chr | Precursor start | Precursor end | Strand | Mature miRNA sequence | Mature miRNA | Precursor sequence |
|---|---|---|---|---|---|---|
| LG2 | 26976934 | 26977059 | - | TTTCATTCCATACATCGTCTAA | tpr-miR1507 | ATCTTAATATGATGTTGGGTAGAGTTGTATGGAATGAAAGATGGGAATTTTCTATTTTGGATTCTGTTCATTCAATTTTCCCTCTTTCGTTCCATACATCGTCTAACCAACGTAGTGTTAAAAGTG |
| LG3 | 30140882 | 30141006 | + | AAGGGTCTGTTTGAGAGAAGTGGT | tpr-miR-Novel1 | AGTTGTTGTCTAAGAACACTAAGGGTCTGTTTGAGAAAAGTGGTAAAGAGGGGAAAAACTCTCCCCATTTTGTCAATTTGCCACTTTTTTCAAACAGTCCCCGAGTCTTCTTTGGTGGTAGACTG |
| LG4 | 3988808 | 3988934 | + | TTGAGCCGCGCCAATATCACT | tpr-miR171b | GAATTCGGAAAGTAGAAAAGCGATGTTGGTGAGGTTCAATCCGAAGACGGATTTACATATAGAAAAATGTAAAATACGATCTCAGATTGAGCCGCGCCAATATCACTTTTATTATTGGATTGCTTTC |
| LG4 | 4041678 | 4041801 | + | TTCCACAGCTTTCTTGAACTT | tpr-miR396a | ATCAAGTCCTGGTCATCTTTTTCCACAGCTTTCTTGAACTTCTTTTGTATCTTAAATCTCTTCAAAGATTAAAAGCCCTAGAAGCTCAAGAAAGCTGTGGGAGAATATGGCAATTCAGGCTTAA |
| LG4 | 4053226 | 4053376 | - | TTCCACAGCTTTCTTGAACTG | tpr-miR396b | ATATGGTCCTCTTTGTATTCTTCCACAGCTTTCTTGAACTGCATCAAAATTAAGAGTTAAATTGCATGCATGTAATTAATGGCACTTTGTGTTCTACTCATTGTTTTGCGGTTCAATAAAGCTGTGGGAAGATACAGATAGGATCAACTAC |
| LG5 | 12423096 | 12423182 | + | CATGTGTTCTTGTTCTCCATC | tpr-miR164a | TCGAGGAGATTGAACCATGCTGGAGAAGCAGGGCACATGCTTGATTTATCAAAAATCTGAGCATGTGCTCTTGTTCTCCATCATGGA |
| LG6 | 6076751 | 6076868 | + | TGGAGAAGCAGGGCACGTGCA | tpr-miR164b | CATTGTTGGGTAGGTCTTGTTGGAGAAGCAGGGCACGTGCAAACATAGATCTCTCTCAAAGTTTCGATCTCATTCTGCACGTGCTCCACTTTTCCAACTTGATTTCCCCACCATCTTT |
| LG7 | 2745274 | 2745411 | + | TGCTCTCTAGACTTCTGTCATC | tpr-miR156b | GAGGGAGAGGCTAAGGTTGTTGACAGAAGATAGAGAGCACTAAGGATGTTATGCATAAACATACATGGAGATGAAGGAGCTTATGATCAATTGCATTTCATTTCCTTTGTGCTCTCTATACTTCTGTCATCCACTCTA |
| LG7 | 6103531 | 6103665 | + | TCGGACCAGGCTTCATTCCCC | tpr-miR166b | TGGGAAGCTTTATTTTGCGGGGAATGTTGTCTGGCTCGAGGACTTTTTCATCTTGATCTATAGAAATATATCTGTCAAGAAATGTTTAGTGTTCTCGGACCAGGCTTCATTCCCCACAATTATATGCTTCCATTC |
| FKJA01000389.1 | 154335 | 154456 | + | TGCCTGGCTCCCTGTATGCCA | tpr-miR160 | AGATATATATAATAATCGCGTGCCTGGCTCCCTGTATGCCATTTGTAGAGTTCATCAAACTGGTGATGACTTTAATGAATGGCGTACGAGGAGCCATGCATGCTGTATATATTTTTTTATAT |
| FKJA01000881.1 | 21693 | 21797 | + | TAATCTGCATCCTGAGGTTT | tpr-miR2111 | AATATTGGGTAATGATTAGATAATCTGCATCCTGAGGTTTATAGCAATATTTTAGTTGTCTTTAATCCTTGGAATGCAGATTATCCCTTCCTTATCGCCAATCCA |
| FKJA01003145.1 | 2374 | 2496 | + | AACCAGGCTCTGATACCATGA | tpr-miR1511 | TTGGCCGAGTGGTTTGTTTCATGGTGTCAGGTCCTGCTTCATCAAGTGGTCTTGAGTTCTACTCCCAATCAAGCACATGGTTAACCAGGCTCTGATACCATGATTAAGGTCCTTAAAAGAGTT |
| ***Vigna angularis*** | | | | | | |
| vigan.Gyeongwon.gnm3.Va02 | 5699390 | 5699516 | + | TGCTCTCTAGACTTCTGTCATC | van-miR156b | GAAGTAGTTGGTAAGGTTGTTGACAGAAGATAGAGAGCACAGATGATGATATGCAGAAACATATGGAACAGGAAAGCAATTGCGTCTCACTCCTTTGTGCTCTCTAGGCTTCTGTCATCCACCTTCA |
| vigan.Gyeongwon.gnm3.Va01 | 27760176 | 27760304 | - | TGACAGAAGAGAGTGAGCAC | van-miR156a | GCAGATTAGGTTAAGGGAGGTGACAGAAGAGAGTGAGCACACATGGTACTTTCTTGGATGATAAGGTTTCATGCTTGAAGCTATGTGTGCTTACTCTCTATCTGTCACCCCATCTCCATCTCTTTCTTT |
| vigan.Gyeongwon.gnm3.Va11 | 12430488 | 12430612 | - | TCGATAAACCTCTGCATCCAG | van-miR162-2 | GGAGATGAGGTGAAGTCACTGGAGGCAGCGGTTCATCGATCTCTTCCTGAATTTGGCTGTGGAAGAACACAAAGCAAGAATCGGTCGATAAACCTCTGCATCCAGCGCTCACTTTGCCTCTCTTT |
| vigan.Gyeongwon.gnm3.Va11 | 13677988 | 13678113 | + | TTGAGCCGCGCCAATATCACT | van-miR171 | GATATGGGAGGCAGAGAAAGCGATGTTGGTGAGGTTCAATCCGAAGACGGATTTACATGTAGAAGCAGTAAAATACGATCTCAGATTGAGCCGCGCCAATATCACTTATCATTGCTGAGTCCCCCT |
| vigan.Gyeongwon.gnm3.Va11 | 13743098 | 13743226 | + | TTCCACAGCTTTCTTGAACTT | van-miR396a | CTCAAGTCCTCGTCATGCTTTTCCACAGCTTTCTTGAACTTCTTATGCATCTTATATCTCTCCACCTTCAGGATTTTAAGCCCTAGAAGCTCAAGAAAGCTGTGGGAGAATATGGCAATTCGGGCTTTA |
| vigan.Gyeongwon.gnm3.Va11 | 13750572 | 13750714 | - | TTCCACAGCTTTCTTGAACTG | van-miR396b | TCATGGTCCTCTTTGTATTCTTCCACAGCTTTCTTGAACTGCATCCATAGAGTTCCTTTGCATGCATGCCATGGCACTCTTGCTCCCACACCTTGTTTTGCGGTTCAATAAAGCTGTGGGAAGATACAGATCGGTCCAACAAC |
| vigan.Gyeongwon.gnm3.Va10 | 3925997 | 3926120 | + | TCGGACCAGGCTTCATTCCCC | van-miR166 | GAGATGGGTTGGTATTGAGGGGAATGTTGGCTGGCTCGAGGCTTTTCACAAACGAGGTTCACAGTGGATGGAACTATAAGGCTTCGGACCAGGCTTCATTCCCCTCAAAAGTACTTCCTTCATG |
| vigan.Gyeongwon.gnm3.Va06 | 5958556 | 5958666 | + | TGAAGCTGCCAGCATGATCTT | van-miR167 | CATCATGCACCACTACCGGTTGAAGCTGCCAGCATGATCTTAACTTTCCTCACTTGCTTGAAGAGAGATCAGATCATGTGGCAGTTTCACCTAGTAGTTGTTGGCCGCATG |
| vigan.Gyeongwon.gnm3.Va06 | 8545307 | 8545428 | + | TGCCTGGCTCCCTGTATGCCA | van-miR160 | CATGCATACATATGTGTATGTGCCTGGCTCCCTGTATGCCATTTGCAGAGCTCATCGAAGCGTCAATGACCTTCGTAGATGGCGTATGAGGAGCCAAGCATATTCCATGTACATACATGGAG |
| vigan.Gyeongwon.gnm3.SuperScaf_33 | 1683497 | 1683619 | + | AACCAGGCTCTGATACCATGA | van-miR1511 | TGGGTTGAGTGGTTCCATTCGTGGTATCAGGTCCTGCTTTATCAAATGATCTCGAGTTCAATCCCTGCTCAAGCACTTGGTTAACCAGGCTCTGATACCATGAACATTACGCCTCTCAACAAT |
| vigan.Gyeongwon.gnm3.Va09 | 6243353 | 6243476 | - | TCGATAAACCTCTGCATCCAG | van-miR162-1 | AGACGTTTTCATGTTACACTGGAGGCAGCGGTTCATCGATCTCTTCCTGAGCTTTGTTGTTTAACGTCACAATCATGAATCGATCGATAAACCTCTGCATCCAGTGTTCTCTGTCACATTTTCT |
| vigan.Gyeongwon.gnm3.scaffold_1871 | 2358 | 2488 | - | CGCTATCCATCCTGAGTTTCA | van-miR390 | TTAGTGTGGAAGAATCTGTAAAGCTCAGGAGGGATAGCGCCATGGATGGTCATTCTTCATCTCTCTTCTCTTTCCTTTGATCTTCTCTTGCGCTATCCATCCTGAGTTTCATGGCTTCTCTCCACACACAC |
| vigan.Gyeongwon.gnm3.SuperScaf_2 | 1930085 | 1930215 | + | TCGCTTGGTGCAGGTCGGGAA | van-miR168 | ATCACTGTGCGGTCTCTAATTCGCTTGGTGCAGGTCGGGAACCGGTTTTTGCGGCAAAATGGTGAAGTGGTAGCCGGCGGCGAATTGGATCCCGCCTTGCATCAACTGAATCGGAGGCCGCAGTGAAGGTT |
| ***Vigna unguiculata*** | | | | | | |
| vigun.IT97K-499-35.gnm1.Vu02 | 19512113 | 19512247 | + | TGTGTTCTCAGGTCACCCCTT | vun-miR398 | GTCATAGGGTTTATCTCAGAGGAGTGAATCTGAGAACACAAGGCTGGTTTGCAGTTCATATCATCATTCTGTTGGTATTTTGGTGAATTTACTTTGTGTTCTCAGGTCACCCCTTTGAGCCAACATGATGACATT |
| vigun.IT97K-499-35.gnm1.Vu02 | 26110301 | 26110430 | - | TCGCTTGGTGCAGGTCGGGAA | vun-miR168 | ATCACTGTGCGGTCTCTAATTCGCTTGGTGCAGGTCGGGAACCGGTTTCTGCGCGAAATGGTGAAGTGGTAGCCGGCGGCGAATTGGATCCCGCCTTGCATCAACTGAATCGGAGGCCGCAGTGAGGGTT |
| vigun.IT97K-499-35.gnm1.Vu03 | 508974 | 509142 | - | TGCTCTCTAGACTTCTGTCATC | vun-miR156a-1 | CACTAGTTTTCTAAGGTTGTTGACAGAAGATAGAGAGCACTGATGATGATATGCATGTTACTATATATATATATATAGAGAGAGAGAGAGCAGAAAATGATGATGAATTGTGCATTTCACTCCTTTGTGCTCTCTATACTTCTGTCATCACCTTCGGCCTCCATTTCTC |
| vigun.IT97K-499-35.gnm1.Vu03 | 39239351 | 39239473 | - | AACCAGGCTCTGATACCATGA | vun-miR1511 | TGGGTTGAATGGTTGCATTCGTGGTATCAGGTCCTGCTTTATCAAATGGTCTTGAGTTCAAGTCCTGCCCAAGCAGTTGGTCAACCAGGCTCTGATACCATGAAGATCACAATTCCCAATTCA |
| vigun.IT97K-499-35.gnm1.Vu03 | 54538004 | 54538128 | - | TCGATAAACCTCTGCATCCAG | vun-miR162 | GGAGATGAGGTGAAGTCACTGGAGGCAGCGGTTCATCGATCTCTTCCTGAATTTGGCTGTGGAAGAACACAAAGCAAGAATCGGTCGATAAACCTCTGCATCCAGCGCTCACTTTGCCTCTCTTT |
| vigun.IT97K-499-35.gnm1.Vu03 | 56141915 | 56142040 | + | TTGAGCCGCGCCAATATCACT | vun-miR171 | GATATGGGAGGCAGAGAAAGCGATGTTGGTGAGGTTCAATCCGAAGACGGATTTACATGTAGAAGCAGTAAAATACGATCTCAGATTGAGCCGCGCCAATATCACTTATCATTGTTGCTTTCCAGC |

| Chr | Precursor start | Precursor end | Strand | Mature miRNA sequence | Mature miRNA | Precursor sequence |
|---|---|---|---|---|---|---|
| vigun.IT97K-499-35.gnm1.Vu03 | 56207574 | 56207687 | + | TTCCACAGCTTTCTTGAACTT | vun-miR396a | CTCAAGTCCTGGTCATGCTTTTCCACAGCTTTCTTGAACTTCTTCTGCACCTTCGCCATTTTAAGCCCTAGAAGCTCA<br>AGAAAGCTGTGGGAGAATATGGCAATTCAGGCTTCT |
| vigun.IT97K-499-35.gnm1.Vu03 | 56215118 | 56215260 | - | TTCCACAGCTTTCTTGAACTG | vun-miR396b | TCATGGTCCTCTTTGTATTCTTCCACAGCTTTCTTGAACTGCATCCATAGAGTTCCTTTGCATGCATGCCATGGCACT<br>CTTGCTCCCACACCTTGTTTTGCGGTTCAATAAAGCTGTGGGAAGATACAGATAGGTCCAACAAC |
| vigun.IT97K-499-35.gnm1.Vu03 | 60021460 | 60021587 | + | CGACAGAAGAGAGTGAGCAC | vun-miR156b | TCGAGGTTTGTTCATGGAGGCGACAGAAGAGAGTGAGCACATCACTCTTTCCTAGCATGCACCTTAGTTTCGTGCT<br>AAAAGCTGTGTGCTTACTCTCTATCTGTCACCACTTTCTTTAACTTCTTATT |
| vigun.IT97K-499-35.gnm1.Vu04 | 2176259 | 2176386 | + | TGCTCTCTAGACTTCTGTCATC | vun-miR156a-2 | GAAGTAGTTGATAAGGTTGTTGACAGAAGATAGAGAGCACAGAAGATGATATGCAGAAACATATGGAACAACAA<br>AAGCAATTGCGTCTCACTCCTTTGTGCTCTCTAGGCTTCTGTCATCCACCTTCA |
| vigun.IT97K-499-35.gnm1.Vu06 | 21349467 | 21349577 | + | CGCTATCTATCCTGAGTTTCA | vun-miR390a | AGAGTACTGAAGAATCTGTAAAGCTCAGGAGGGATAGCGCCATGATTTGTCACATCATCGTGTTCTTTGGCGCTAT<br>CTATCCTGAGTTTCACGGTTTCTTCTTACTGTGTT |
| vigun.IT97K-499-35.gnm1.Vu07 | 32010417 | 32010535 | + | AGAATCTTGATGATGCTGCAG | vun-miR172 | TATAGTCGTTGTTTGCAGGTGCAGCAGCATCAAGATTCACACACAAATTCTACCTCCATGGGGGAATGTTTTGGAG<br>TTGAGAATCTTGATGATGCTGCATCAGCAATAGACGATTCCCG |
| vigun.IT97K-499-35.gnm1.Vu07 | 32453170 | 32453291 | - | TGCCTGGCTCCCTGTATGCCA | vun-miR160 | CATGCATACATATGTCTATGTGCCTGGCTCCCTGTATGCCATTTGCAGAGCTCATCGAACCGTCAATGACCTCCGTG<br>GATGGCGTATGAGGAGCCAAGCATATTCCATGTACATACATGGAG |
| vigun.IT97K-499-35.gnm1.Vu07 | 35590036 | 35590146 | - | TGAAGCTGCCAGCATGATCTT | vun-miR167 | CATCATGCACCACTACCAGTTGAAGCTGCCAGCATGATCTTAACTTTCCTCACCCAGTTGAAGAGAGATCAGATCA<br>TGTGGCAGTTTCACCCAGTAGTTGTTGGCCGCATG |
| vigun.IT97K-499-35.gnm1.Vu08 | 3923799 | 3923927 | + | TGACAGAAGAGAGTGAGCAC | vun-miR156c-1 | AGATTAGGGTAAAAGGGAGGTGACAGAAGAGAGTGAGCACACATGGTACTTTCTTGGATGATAAGGTTTCATGCT<br>TGAAGCTATGTGTGCTTACTCTCTATCTGTCACCCCATCTTCATCTCTTTCTTT |
| vigun.IT97K-499-35.gnm1.Vu09 | 27035706 | 27035829 | - | TCGGACCAGGCTTCATTCCCC | vun-miR166 | ATGGGTTCGTAGTGTTGAGGGGAATGTTGGCTGGCTCGAGGCTTTTCACATAGGAGGTTCTCACTGGCAAGAACTA<br>TAAGGCTTCGGACCAGGCTTCATTCCCCTCAAAACTTTTCTTTTTGTG |
| vigun.IT97K-499-35.gnm1.Vu09 | 38948334 | 38948456 | + | TGACAGAAGAGAGTGAGCAC | vun-miR156c-2 | GTTTCTTGGAACATAGAAATTGACAGAAGAGAGTGAGCACACAGAGGCACCTGGTATAGTCATACACTGTTACTTT<br>TGCGTGCTCACTTCTCTTTCTGTCAACTTCCAGTGCTGGAAATTACT |
| vigun.IT97K-499-35.gnm1.Vu10 | 25038234 | 25038373 | + | CGCTATCCATCCTGAGTTTCA | vun-miR390b | TTAGTGTGGAAGAATCTGTAAAGCTCAGGAGGGATAGCGCCATGGATGGTCATTCCTCACCTCTCTTTTTCTCTCTT<br>TCTCCCCTTTGATCTTCTCTTGCGCTATCCATCCTGAGTTTCATGGCTTCTCTCCACACACAA |

**Table S5. The detailed information about the bacterial small RNAs, chromosomal location, start and end position, length, sequence information of mature and precursors miRNAs with their corresponding RPM values from nodule library**

| Chr | Strand | Number of reads | Mature start | Mature end | Precursor start | Precursor end | Mature sequence | Precursor sequence |
|---|---|---|---|---|---|---|---|---|
| NZ_CP015062.1 | + | 9 | 2408577 | 2408594 | 2408554 | 24085942 | TCACGGCGGGCGACCTGT | GGGTTCCCTGCTATTGATGCCTTTCACGGCGGGCGACCTGT |
| NZ_CP015062.1 | + | 1 | 484916 | 484939 | 484916 | 484979 | CTGGGTGCGGCCGGTTCCTCGACC | CTGGGTGCGGCCGGTTCCTCGACCTGCCGTGTAACGCTCTGGGTGAGATTGCTGGCCTCCGGTC |
| NZ_CP015062.1 | + | 32 | 4284652 | 4284675 | 4284603 | 4284675 | CGGGTTTGAGGGGGTGGGTGGAGC | GACACCCTCCCTTCTGGACCTTTTGCCCTTGAGCATCGATCACTGCAGTCGGGTTTGAGGGGGTGGGTGGAGC |
| NZ_CP015062.1 | + | 221 | 4933790 | 4933813 | 4933790 | 4933862 | GCGGGTGTAGCTCAGGGGTAGAGC | GCGGGTGTAGCTCAGGGGTAGAGCACAACCTTGCCAAGGTTGGGGTCGAGCGTTCGAATCGCTTCACCCGCT<br>C |
| NZ_CP015062.1 | + | 1 | 5013737 | 5013759 | 5013737 | 5013810 | GCAGCTGGACCGTGGAAGGCATC | GCAGCTGGACCGTGGAAGGCATCGGCGAGGATTTCGTGCCGCCCAATGCCGATCTGTCGCTGGTCAAGAAGG<br>CC |
| NZ_CP015062.1 | + | 13 | 5020399 | 5020419 | 5020295 | 5020419 | AGCGGAGGCTGGGTGAGGGGG | CCCTCATCCGGCCGCTTCGCGGCCACCTTCTCCCCGCTGGGGAGAAGAGGGAAGCGCCGGCGCTGCAAGTCT<br>CCTCTCCCCTCGGGGAGAGGTCAGCCGAGCGAAGCGGAGGCTGGGTGAGGGGG |
| NZ_CP015062.1 | + | 1 | 5042623 | 5042646 | 5042590 | 5042646 | CGGCGTGCCGACGGCGATCCTGGA | CGGGCTGGCCGTCGCGCACGCCTTGATGAACGCCGGCGTGCCGACGGCGATCCTGGA |
| NZ_CP015062.1 | + | 3 | 5047641 | 5047658 | 5047641 | 5047684 | CGGTCGGCTGCTGGGACT | CGGTCGGCTGCTGGGACTGGAGATCCGAAAGCAGCCGGGCCGAT |
| NZ_CP015062.1 | + | 1 | 5264185 | 5264206 | 5264156 | 5264206 | GGCGATTGCCGTGCGAAGGGGC | CCCTCGCCCGTAACTCTGCGTTTCATGGAGGCGATTGCCGTGCGAAGGGGC |
| NZ_CP015062.1 | + | 101 | 5565022 | 5565043 | 5564974 | 5565043 | GACGGCTGGCAGGCGAACTGGA | AACTTCACGCTGGCGCCCAACACGACTGATCCTTTCTATATTCACGGTGACGGCTGGCAGGCGAACTGGA |
| NZ_CP015062.1 | + | 7 | 2553288 | 2553311 | 2553288 | 2553466 | GCTGATCGGGTTCATGTAGACGGT | GCTGATCGGGTTCATGTAGACGGTGCGCGTGTTCAGCCGGTGGTCGGTGACATTGTCGGCCAGCCCGATATG<br>CTGCAGCAGGCCGGTCAGCACCATGTGCCAGCTGCCGTAGAGGCGCGGCAGGCCGATCAGCACCAGCGGCA<br>GGAACGACCACAGATGAAGCGCTAGCCCGATCGCCC |
| NZ_CP015062.1 | + | 2 | 6225301 | 6225318 | 6225221 | 6225318 | ATGTCGCGCTGGAAAAGT | TTTTCTGGCGATCGACGTTCCGCTAGATCGCGCGGCCCAGGCGCTGGAGACCTTCGCCGCCGACGCCGGTCTG<br>TCACTCGATGTCGCGCTGGAAAAGT |
| NZ_CP015062.1 | - | 9 | 6166158 | 6166181 | 6166114 | 6166181 | ATCAACGATCGGTCAAAGGAGCTC | ATCAACGATCGGTCAAAGGAGCTCGCCATGTCCGTGGACAGTTTTTCCTCACATCGGCACGATGATGC |
| NZ_CP015062.1 | + | 2 | 646888 | 646906 | 646844 | 646906 | ACATCGTCGGTGTCAAGGA | CATGATGCCGGACACGATGGGCCGGCTCGTGCACGACTTCAAGAACATCGTCGGTGTCAAGGA |
| NZ_CP015062.1 | - | 2 | 6036001 | 6036024 | 6036001 | 6036067 | CAGGTACTGCTTCGGGAAGGCGTC | TGTGCGCGGGACCAGGACAGAAGGTGCCGTAGGCCATGGCATACAGGTACTGCTTCGGGAAGGCGTC |
| NZ_CP015062.1 | - | 1 | 5893135 | 5893155 | 5893135 | 5893189 | ACCTCGACGCCGGTGGGAGAT | TTCCACGCCGGTCGACTACATCAACACCCATGGCACCTCGACGCCGGTGGGAGAT |
| NZ_CP015062.1 | - | 13 | 5853754 | 5853776 | 5853728 | 5853776 | ATTCGGCGAGGCCCTGCACAAGG | ATTCGGCGAGGCCCTGCACAAGGAGATGGACAAGGTGCTGGCCGACCTG |
| NZ_CP015062.1 | - | 1 | 5697345 | 5697368 | 5697298 | 5697368 | TCCAGCGCGAGATCGCCGAACGGC | TCCAGCGCGAGATCGCCGAACGGCTCGGCTACACCACGCACCCCGGCCTGTCGGCGGTCGAGCGCTTCATG |
| NZ_CP015062.1 | + | 3 | 675549 | 675567 | 675549 | 675774 | CTTCTGGGTCGGCGGCACC | CTTCTGGGTCGGCGGCACCCGCGAGAATGCCGGGGATGTCGGCGAGCTTGAGCGGCTGCCGGGTGCGGCCGG<br>GATCAAGGTTTTCATGGGCTCGTCCACTGGCGACCTGCTGGTCGAGGATGACGAGGGCGTGGCGTCGATCCT<br>GAGAAACACGCGCCGGCGCGCCGCCTTTCATTCGGAAGACGAATTCCGGCTGCGCGAGCGCCTTGGCGAACG<br>CATCGAAGGC |
| NZ_CP015062.1 | + | 14 | 2631416 | 2631437 | 2631390 | 2631437 | AGGGCGGGATCATGGAGGGACC | TCCCTCGTCTCCGGCAAGCTCTTTTTAGGGCGGGATCATGGAGGGACC |
| NZ_CP015062.1 | + | 3 | 675550 | 675568 | 675550 | 675773 | TTCTGGGTCGGCGGCACCC | TTCTGGGTCGGCGGCACCCGCGAGAATGCCGGGGATGTCGGCGAGCTTGAGCGGCTGCCGGGTGCGGCCGG<br>GATCAAGGTTTTCATGGGCTCGTCCACTGGCGACCTGCTGGTCGAGGATGACGAGGGCGTGGCGTCGATCCT<br>GAGAAACACGCGCCGGCGCGCCGCCTTTCATTCGGAAGACGAATTCCGGCTGCGCGAGCGCCTTGGCGAACG<br>CATCGAAGG |
| NZ_CP015062.1 | - | 5 | 5586772 | 5586793 | 5586772 | 5586878 | ATGGGCGGGCTTGGTGGCGGGT | CGGCGGCGGAAGCCTTGTCGGTCTTGACGGTCGAAACGGTCATTTCACTGCTCCTTGGGATTTCGATTCAGGC |

| Chr | Strand | Number of reads | Mature start | Mature end | Precursor start | Precursor end | Mature sequence | Precursor sequence |
|---|---|---|---|---|---|---|---|---|
| | | | | | | | | AACGGCGTGGAGATGGGCGGGCTTGGTGGCGGGT |
| NZ_CP015062.1 | - | 52 | 4988927 | 4988948 | 4988872 | 4988948 | GATCGCAATCGTCATCCTGGTG | GATCGCAATCGTCATCCTGGTGGTGGTCAAGCCGTTCTGATGCATGTCGCTCAAAAACGGCCAGGAGACTGCGACAG |
| NZ_CP015062.1 | - | 13 | 4954408 | 4954428 | 4954313 | 4954428 | AGCGGAGGCTGGGTGAGGGGG | AGCGGAGGCTGGGTGAGGGGGCCGATGCCCCAATACAGTTCTCCGCAGGAGCAGCATCAACACAAGCTATCCGCCGAAGGAGAGAGATATGGCCGCCGTCACCGACAGTCTGCTGA |
| NZ_CP015062.1 | - | 2 | 4803176 | 4803193 | 4803134 | 4803193 | ATTTCGCGACCGCCCGCC | ATTTCGCGACCGCCCGCCGCGAGGCCGAGGCGGCGATCCGGATGGACCGTCGCGAGGGCG |
| NZ_CP015062.1 | - | 1 | 4790206 | 4790227 | 4790206 | 4790273 | ATTGTGGCCCGGTGACGTCGAG | AGAATGACGGGCTGCACCGTAGCTTCGCGCGGCGGACGGTGGAGCAATTGTGGCCCGGTGACGTCGAG |
| NZ_CP015062.1 | - | 9 | 4141410 | 4141430 | 4141410 | 4141458 | ACACGGTCGAGGAGCTGGAGC | TTCGGCATTCCGGTACGGCTCAATTTCTACACGGTCGAGGAGCTGGAGC |
| NZ_CP015062.1 | - | 2 | 4135697 | 4135719 | 4135697 | 4135828 | TCCGGCGGCGGCGCCAAGCGCTC | GCGCAAGGACGCCGACAAGCCGGTCAAGGAAGCGTCGTCGAAGCCGAAATCCGATGACAAGCAATTCAACGCCGACGAGATCTCCGCGCTCCTCGACAAGCAGAAACCATCCGGCGGCGGCGCCAAGCGCTC |
| NZ_CP015062.1 | - | 48 | 3524778 | 3524795 | 3524704 | 3524795 | TCCAGTCGGTGGCGTGGC | TCCAGTCGGTGGCGTGGCCCTCGCATCCCATCATCGCCGGCGGCCAGCATGTCGTCGTCGTCGGCGGCGGCGATACCGCGTCCGACTGCGTC |
| NZ_CP015062.1 | + | 1 | 2904429 | 2904446 | 2904357 | 2904446 | AGCCGATCTCCCCCCTTG | AGGGGGGAGATTGGCAGCTTTGCCGACTGCTTTCGCTTTGCGACGTTGTGAATTGGCGAAGACGGCCATGACAGCCGATCTCCCCCCTTG |
| NZ_CP015062.1 | - | 2 | 3524130 | 3524149 | 3524094 | 3524149 | CCGCGCTGCGAGACTAACGA | CCGCGCTGCGAGACTAACGATTGATCGCGGTCGTGGTGTCGGTCGCGGCTGCGGCC |
| NZ_CP015062.1 | - | 2 | 3499947 | 3499965 | 3499947 | 3500024 | CTCCGGAGCCTGAACAGGA | CATCGTGGCGCCGGAGAATGCGCGGCGCGCCGAGATTCCGCAGGTGACCGAACAGCAGGCTCCGGAGCCTGAACAGGA |
| NZ_CP015062.1 | + | 89 | 920916 | 920934 | 920916 | 920966 | ATCTGGTCGGCGAGGACGT | ATCTGGTCGGCGAGGACGTGCTGGTCACCGGCGCCGGGCCGATCGGCATCA |
| NZ_CP015062.1 | - | 1 | 2623060 | 2623081 | 2623018 | 2623081 | GCGGGCGCGGCGCTGCAGTACC | GCGGGCGCGGCGCTGCAGTACCAGGTCTCGGAAGGCTATCTGCCTTTGCGGCGCTGGCTCGCCG |
| NZ_CP015062.1 | - | 512 | 2563828 | 2563849 | 2563778 | 2563849 | GATGATCATGTGGGCTTTCCTC | GATGATCATGTGGGCTTTCCTCAACTTCACCGATAGCTGGTGGGGGTTGGCGCTGATCTACGGCATCGTCAC |
| NZ_CP015062.1 | - | 22 | 2454544 | 2454564 | 2454517 | 2454564 | CCATCGGTTCGGGCCTGCATG | CCATCGGTTCGGGCCTGCATGGCGACCATGAGTTCGACGCCACCGGCT |
| NZ_CP015062.1 | - | 3 | 2284913 | 2284934 | 2284913 | 2285052 | TTTCGGCGACGCTGGGCGAGGC | TTCGCGCGCATCTACTGGATCGCCGGCGGCCTGCCCAAGGAAGGCGGCATCGAACCCTTGCGCGGTTTCTACCCGCGCATCGCCAAGGCCTATCTGATCGGCGAGGCGGCGCCGGCGTTTTCGGCGACGCTGGGCGAGGC |
| NZ_CP015062.1 | + | 3 | 961351 | 961370 | 961351 | 961399 | CCGGCTGGGCGAGGGGCTCA | CCGGCTGGGCGAGGGGCTCACCGCCAATGGCCGGCTCGGCCAGCCGGCG |
| NZ_CP015062.1 | - | 39 | 1914501 | 1914521 | 1914471 | 1914521 | ATGGGAAGGCGGAGGCGGGTT | ATGGGAAGGCGGAGGCGGGTTTCGGTTCTATCGCCTTGCGCCGTCGCTTCT |
| NZ_CP015062.1 | - | 195 | 1901631 | 1901651 | 1901631 | 1901706 | CGATTCCGGGCGAGGCCTCCA | GAGGCCTCGTGGCGGAGTGGTTACGCAGAGGACTGCAAATCCTTGCACCCCGGTTCGATTCCGGGCGAGGCCTCCA |
| NZ_CP015062.1 | - | 83 | 1882833 | 1882855 | 1882791 | 1882855 | TCAAGTCGGACGGTTCGCTGGTG | TCAAGTCGGACGGTTCGCTGGTGCCGCGTGAAGATCCGGCGGCACCCGCCGCTCCGCAAGTGGCC |
| NZ_CP015062.1 | - | 2 | 1731602 | 1731622 | 1731602 | 1731645 | CAACCGGCCGGCGCGCAAGCT | ATTGCCGCCGGCCTGATCATCGTCAACCGGCCGGCGCGCAAGCT |
| NZ_CP015062.1 | - | 25 | 1710984 | 1711003 | 1710984 | 1711076 | TCGGGTGGGCGGCGAGGGTT | CCGGCGCCTTTGACAGGAATCTGATGAGAGCCGATCGCCGAGGGGCGATCGGCTTTTCGACATTTGGCAGGAATCGGGTGGGCGGCGAGGGTT |
| NZ_CP015062.1 | + | 9 | 1057917 | 1057936 | 1057861 | 1057936 | TGTGCAGCAGGCGCTCTTGA | AAGATTTCGCGCAAGTCGGACAATCCCTCGATCACCCGCGACGTGTCGGGCGAGGGTGTGCAGCAGGCGCTCTTGA |
| NZ_CP015062.1 | - | 2 | 563175 | 563198 | 563175 | 563248 | TCAGCGACGGCGGCTTCGATTTGC | AACCTATGTCGAGCCGCCTCTGGCGGACTATGTGACGATGTATGCAGCAGTCAGCGACGGCGGCTTCGATTTGC |
| NZ_CP015062.1 | - | 2 | 126833 | 126854 | 126781 | 126854 | ACACCGCGGCGACCGATCCGAC | ACACCGCGGCGACCGATCCGACCACCTTGCTGAACACGGCGGCAAAGTCGTTCCTGAATGCGGTCGGCGCCTCC |
| NZ_CP015062.1 | + | 2 | 1474076 | 1474097 | 1474037 | 1474097 | CTGGTGGTCGGCCGTGAAGGCG | CTTGGGCCGGCTCTTCTGGCAGTTGTCGGCGTCGCCGTGCTGGTGGTCGGCCGTGAAGGCG |
| NZ_CP015062.1 | + | 12 | 2236467 | 2236487 | 2236467 | 2236527 | AGCATCACCGTGGAGTAGACC | AGCATCACCGTGGAGTAGACCGAGTCATGCCCTTCGTCTTTCCGCCATGGATGCGCATTGC |
| NZ_CP015062.1 | + | 11 | 3296549 | 3296568 | 3296549 | 3296611 | ATCCGACGCCGTGCGAGAAC | ATCCGACGCCGTGCGAGAACTGCTTGTCATGTTGGACGGCATTTTCGGCAGCGGCAAGGATGA |
| NZ_CP015062.1 | + | 3 | 2317252 | 2317272 | 2317252 | 2317325 | TTGTAGACGCTGGTGCTCTGT | TTGTAGACGCTGGTGCTCTGTGTCGTGGTGCCGTTTTCGGTGCAAGAGACAGTCGAAACCAGTGTCCATCCGGC |
| NZ_CP015062.1 | + | 7 | 3644310 | 3644329 | 3644265 | 3644329 | AGGACAGGAAAGTCGGGATT | TGCCGAACTCCTGCTTGTGCTTCGACAGCTTCTTTTCGATGGCATAGGACAGGAAAGTCGGGATT |
| NZ_CP015062.1 | + | 16 | 3761408 | 3761427 | 3761408 | 3761459 | AGTCTGCGATCTTCTGTTCG | AGTCTGCGATCTTCTGTTCGGCGGCGTGCTGCCGGCGACGGAGGATCGTTTC |
| NZ_CP015062.1 | + | 107 | 3943077 | 3943097 | 3943040 | 3943097 | ACATGCTGGAGATCGCGATCC | CTGGCGACCACCGGCAGCGACGGCGCGCCGGAGCCGGACATGCTGGAGATCGCGATCC |

**Table S6. The distribution of 91 microRNA in different tissues of chickpea**

| miRs exclusive to nodule library |
|---|
| Cat-miR156f-2 |
| Cat-miR171c |
| Cat-miR156f-1 |
| Cat-miR162a |
| Cat-miR168 |
| Cat-miR162b |
| Cat-NovmiR7 |

**miRs common between nodule and other tissue libraries**

| Fbud | Flower | Leaf | Pod | Root | Shoot | Stem |
|---|---|---|---|---|---|---|
| Cat-NovmiR6 | Cat-NovmiR6 | Cat-NovmiR6 | Cat-NovmiR6 | Cat-NovmiR6 | Cat-NovmiR6 | Cat-NovmiR6 |
| Cat-miR2118 | Cat-miR2118 | Cat-miR2118 | Cat-miR2118 | Cat-miR2118 | Cat-miR2118 | Cat-miR2118 |
| Cat-miR164a-3 | Cat-miR164a-3 | Cat-miR164a-3 | Cat-miR164a-3 | Cat-miR164a-3 | Cat-miR164a-3 | Cat-miR164a-3 |

| Fbud | Flower | Leaf | Pod | Root | Shoot | Stem |
|---|---|---|---|---|---|---|
| Cat-miR167b-2 | Cat-miR167b-2 | Cat-miR167b-2 | Cat-miR167b-2 | Cat-miR167b-2 | Cat-miR167b-2 | Cat-miR167b-2 |
| Cat-miR156a-1 | Cat-miR156a-1 | Cat-miR156a-1 | Cat-miR156a-1 | Cat-miR156a-1 | Cat-miR156a-1 | Cat-miR156a-1 |
| Cat-miR172c-1 | Cat-miR172c-1 | Cat-miR172c-1 | Cat-miR172c-1 | Cat-miR172c-1 | Cat-miR172c-1 | Cat-miR172c-1 |

| Fbud | Flower | Leaf | Pod | Root | Shoot | Stem |
|---|---|---|---|---|---|---|
| Cat-miR156c-4 | Cat-miR156c-4 | Cat-miR156c-4 | Cat-miR156c-4 | Cat-miR156c-4 | Cat-miR156c-4 | Cat-miR156c-4 |
| Cat-miR156c-2 | Cat-miR156c-2 | Cat-miR156c-2 | Cat-miR156c-2 | Cat-miR156c-2 | Cat-miR156c-2 | Cat-miR156c-2 |
| Cat-miR399-1 | Cat-miR399-1 | Cat-miR399-1 | Cat-miR399-1 | Cat-miR399-1 | Cat-miR399-1 | Cat-miR399-1 |
| Cat-miR167c | Cat-miR167c | Cat-miR167c | Cat-miR167c | Cat-miR167c | Cat-miR167c | Cat-miR167c |
| Cat-miR482 | Cat-miR482 | Cat-miR482 | Cat-miR482 | Cat-miR482 | Cat-miR482 | Cat-miR482 |
| Cat-miR156c-1 | Cat-miR156c-1 | Cat-miR156c-1 | Cat-miR156c-1 | Cat-miR156c-1 | Cat-miR156c-1 | Cat-miR156c-1 |
| Cat-miR166a-5 | Cat-miR166a-5 | Cat-miR166a-5 | Cat-miR166a-5 | Cat-miR166a-5 | Cat-miR166a-5 | Cat-miR166a-5 |
| Cat-miR319c-1 | Cat-miR319c-1 | Cat-miR319c-1 | Cat-miR319c-1 | Cat-miR319c-1 | Cat-miR319c-1 | Cat-miR319c-1 |
| Cat-miR171f | Cat-miR171f | Cat-miR171f | Cat-miR171f | Cat-miR171f | Cat-miR171f | Cat-miR171f |
| Cat-miR166b-1 | Cat-miR166b-1 | Cat-miR166b-1 | Cat-miR166b-1 | Cat-miR166b-1 | Cat-miR166b-1 | Cat-miR166b-1 |
| Cat-miR167a-2 | Cat-miR167a-2 | Cat-miR167a-2 | Cat-miR167a-2 | Cat-miR167a-2 | Cat-miR167a-2 | Cat-miR167a-2 |
| Cat-miR319c-4 | Cat-miR319c-4 | Cat-miR319c-4 | Cat-miR319c-4 | Cat-miR319c-4 | Cat-miR319c-4 | Cat-miR319c-4 |
| Cat-miR167a-1 | Cat-miR167a-1 | Cat-miR167a-1 | Cat-miR167a-1 | Cat-miR167a-1 | Cat-miR167a-1 | Cat-miR167a-1 |
| Cat-miR156a-2 | Cat-miR156a-2 | Cat-miR156a-2 | Cat-miR156a-2 | Cat-miR156a-2 | Cat-miR156a-2 | Cat-miR156a-2 |
| Cat-miR164a-1 | Cat-miR164a-1 | Cat-miR164a-1 | Cat-miR164a-1 | Cat-miR164a-1 | Cat-miR164a-1 | Cat-miR164a-1 |
| Cat-miR1509 | Cat-miR1509 | Cat-miR1509 | Cat-miR1509 | Cat-miR1509 | Cat-miR1509 | Cat-miR1509 |
| Cat-miR167d | Cat-miR167d | Cat-miR167d | Cat-miR167d | Cat-miR167d | Cat-miR167d | Cat-miR167d |
| Cat-miR408 | Cat-miR408 | Cat-miR408 | Cat-miR408 | Cat-miR408 | Cat-miR408 | Cat-miR408 |
| Cat-miR319d | Cat-miR319d | Cat-miR319d | Cat-miR319d | Cat-miR319d | Cat-miR319d | Cat-miR319d |
| Cat-miR160-3 | Cat-miR160-3 | Cat-miR160-3 | Cat-miR160-3 | Cat-miR160-3 | Cat-miR160-3 | Cat-miR160-3 |
| Cat-miR172c-2 | Cat-miR172c-2 | Cat-miR172c-2 | Cat-miR172c-2 | Cat-miR172c-2 | Cat-miR172c-2 | Cat-miR172c-2 |
| Cat-miR319b | Cat-miR319b | Cat-miR319b | Cat-miR319b | Cat-miR319b | Cat-miR319b | Cat-miR319b |
| Cat-miR160-1 | Cat-miR160-1 | Cat-miR160-1 | Cat-miR160-1 | Cat-miR160-1 | Cat-miR160-1 | Cat-miR160-1 |
| Cat-miR156d | Cat-miR156d | Cat-miR156d | Cat-miR156d | Cat-miR156d | Cat-miR156d | Cat-miR156d |
| Cat-miR159-1 | Cat-miR159-1 | Cat-miR159-1 | Cat-miR159-1 | Cat-miR159-1 | Cat-miR159-1 | Cat-miR159-1 |
| Cat-miR160-2 | Cat-miR160-2 | Cat-miR160-2 | Cat-miR160-2 | Cat-miR160-2 | Cat-miR160-2 | Cat-miR160-2 |
| Cat-miR171e | Cat-miR171e | Cat-miR171e | Cat-miR171e | Cat-miR171e | Cat-miR171e | Cat-miR171e |
| Cat-miR172b | Cat-miR172b | Cat-miR172b | Cat-miR172b | Cat-miR172b | Cat-miR172b | Cat-miR172b |
| Cat-miR398a-3p | Cat-miR398a-3p | Cat-miR398a-3p | Cat-miR398a-3p | Cat-miR398a-3p | Cat-miR398a-3p | Cat-miR398a-3p |
| Cat-miR156c-3 | Cat-miR156c-3 | Cat-miR156c-3 | Cat-miR156c-3 | Cat-miR156c-3 | Cat-miR156c-3 | Cat-miR156c-3 |
| Cat-miR164a-2 | Cat-miR164a-2 | Cat-miR164a-2 | Cat-miR164a-2 | Cat-miR164a-2 | Cat-miR164a-2 | Cat-miR164a-2 |
| Cat-miR390b | Cat-miR390b | Cat-miR390b | Cat-miR390b | Cat-miR390b | Cat-miR390b | Cat-miR390b |
| Cat-miR319c-2 | Cat-miR319c-2 | Cat-miR319c-2 | Cat-miR319c-2 | Cat-miR319c-2 | Cat-miR319c-2 | Cat-miR319c-2 |
| Cat-miR2111-5 | Cat-miR2111-5 | Cat-miR2111-5 | Cat-miR2111-5 | Cat-miR2111-5 | Cat-miR2111-5 | Cat-miR2111-5 |
| Cat-miR399-3 | Cat-miR399-3 | Cat-miR399-3 | Cat-miR399-3 | Cat-miR399-3 | Cat-miR399-3 | Cat-miR399-3 |
| Cat-miR399-2 | Cat-miR399-2 | Cat-miR399-2 | Cat-miR399-2 | Cat-miR399-2 | Cat-miR399-2 | Cat-miR399-2 |
| Cat-miR5213 | Cat-miR5213 | Cat-miR5213 | Cat-miR5213 | Cat-miR5213 | Cat-miR5213 | Cat-miR5213 |
| Cat-miR398b-2 | Cat-miR398b-2 | Cat-miR398b-2 | Cat-miR398b-2 | Cat-miR398b-2 | Cat-miR398b-2 | Cat-miR398b-2 |
| Cat-miR390a-2 | Cat-miR390a-2 | Cat-miR390a-2 | Cat-miR390a-2 | Cat-miR390a-2 | Cat-miR390a-2 | Cat-miR390a-2 |
| Cat-miR319a | Cat-miR319a | Cat-miR319a | Cat-miR319a | Cat-miR319a | Cat-miR319a | Cat-miR319a |
| Cat-miR390a-1 | Cat-miR390a-1 | Cat-miR390a-1 | Cat-miR390a-1 | Cat-miR390a-1 | Cat-miR390a-1 | Cat-miR390a-1 |

| Fbud | Flower | Leaf | Pod | Root | Shoot | Stem |
|---|---|---|---|---|---|---|
| Cat-miR398b-1 | Cat-miR398b-1 | Cat-miR398b-1 | Cat-miR398b-1 | Cat-miR398b-1 | Cat-miR398b-1 | Cat-miR398b-1 |
| Cat-miR319c-3 | Cat-miR319c-3 | Cat-miR319c-3 | Cat-miR319c-3 | Cat-miR319c-3 | Cat-miR319c-3 | Cat-miR319c-3 |
| Cat-miR166a-4 | Cat-miR166a-4 | Cat-miR166a-4 | Cat-miR166a-4 | Cat-miR166a-4 | Cat-miR166a-4 | Cat-miR166a-4 |
| Cat-miR156a-3 | Cat-miR156a-3 | Cat-miR156a-3 | Cat-miR156a-3 | Cat-miR156a-3 | Cat-miR156a-3 | Cat-miR156a-3 |
| Cat-miR1511 | Cat-miR1511 | Cat-miR1511 | Cat-miR1511 | Cat-miR1511 | Cat-miR1511 | Cat-miR1511 |
| Cat-miR159-2 | Cat-miR159-2 | Cat-miR159-2 | Cat-miR159-2 | Cat-miR159-2 | Cat-miR159-2 | Cat-miR159-2 |
| Cat-miR166b-3 | Cat-miR166b-3 | Cat-miR166b-3 | Cat-miR166b-3 | Cat-miR166b-3 | Cat-miR166b-3 | Cat-miR166b-3 |
| Cat-miR166b-2 | Cat-miR166b-2 | Cat-miR166b-2 | Cat-miR166b-2 | Cat-miR166b-2 | Cat-miR166b-2 | Cat-miR166b-2 |
| Cat-miR171b | Cat-miR171b | Cat-miR171b | Cat-miR171b | Cat-miR171b | Cat-miR171b | Cat-miR171b |
| Cat-miR2111-3 | Cat-miR2111-3 | Cat-miR2111-3 | Cat-miR2111-3 | Cat-miR2111-3 | Cat-miR2111-3 | Cat-miR2111-3 |
| Cat-miR2111-4 | Cat-miR2111-4 | Cat-miR2111-4 | Cat-miR2111-4 | Cat-miR2111-4 | Cat-miR2111-4 | Cat-miR2111-4 |
| Cat-miR171a | Cat-miR171a | Cat-miR171a | Cat-miR171a | Cat-miR171a | Cat-miR171a | Cat-miR171a |
| Cat-miR166a-1 | Cat-miR166a-1 | Cat-miR166a-1 | Cat-miR166a-1 | Cat-miR166a-1 | Cat-miR166a-1 | Cat-miR166a-1 |
| Cat-miR394-1 | Cat-miR394-1 | Cat-miR394-1 | Cat-miR394-1 | Cat-miR394-1 | Cat-miR394-1 | Cat-miR394-1 |
| Cat-miR167b-1 | Cat-miR167b-1 | Cat-miR167b-1 | Cat-miR167b-1 | Cat-miR167b-1 | Cat-miR167b-1 | Cat-miR167b-1 |
| Cat-miR166a-2 | Cat-miR166a-2 | Cat-miR166a-2 | Cat-miR166a-2 | Cat-miR166a-2 | Cat-miR166a-2 | Cat-miR166a-2 |
| Cat-miR1507 | Cat-miR1507 | Cat-miR1507 | Cat-miR1507 | Cat-miR1507 | Cat-miR1507 | Cat-miR1507 |
| Cat-miR396a | Cat-miR396a | Cat-miR396a | Cat-miR396a | Cat-miR396a | Cat-miR396a | Cat-miR396a |
| Cat-NovmiR4 | Cat-NovmiR4 | Cat-NovmiR4 | Cat-NovmiR4 | Cat-NovmiR4 | Cat-NovmiR4 | Cat-NovmiR4 |
| Cat-miR394-2 | Cat-miR394-2 | Cat-miR394-2 | Cat-miR394-2 | Cat-miR394-2 | Cat-miR394-2 | Cat-miR394-2 |
| Cat-miR2111-1 | Cat-miR2111-1 | Cat-miR2111-1 | Cat-miR2111-1 | Cat-miR2111-1 | Cat-miR2111-1 | Cat-miR2111-1 |
| Cat-miR396c | Cat-miR396c | Cat-miR396c | Cat-miR396c | Cat-miR396c | Cat-miR396c | Cat-miR396c |
| Cat-miR396b | Cat-miR396b | Cat-miR396b | Cat-miR396b | Cat-miR396b | Cat-miR396b | Cat-miR396b |
| Cat-miR166a-3 | Cat-miR166a-3 | Cat-miR166a-3 | Cat-miR166a-3 | Cat-miR166a-3 | Cat-miR166a-3 | Cat-miR166a-3 |
| Cat-miR398a-5p | Cat-miR398a-5p | Cat-miR398a-5p | Cat-miR398a-5p | Cat-miR398a-5p | Cat-miR398a-5p | Cat-miR398a-5p |
| Cat-miR2111-2 | Cat-miR2111-2 | Cat-miR2111-2 | Cat-miR2111-2 | Cat-miR2111-2 | Cat-miR2111-2 | Cat-miR2111-2 |
| Cat-NovmiR5 | Cat-NovmiR5 | Cat-NovmiR5 | Cat-NovmiR5 | Cat-NovmiR5 | Cat-NovmiR5 | Cat-NovmiR5 |
| Cat-miR171d | Cat-NovmiR2 | Cat-miR171d | Cat-miR171d | Cat-miR171d | Cat-miR171d | Cat-miR171d |
| Cat-miR156e-1 | Cat-miR156e-1 | Cat-NovmiR2 | Cat-NovmiR2 | Cat-NovmiR2 | Cat-NovmiR2 | Cat-NovmiR2 |
| Cat-miR156e-2 | Cat-miR156e-2 | Cat-miR156e-1 | Cat-miR172a-3p | Cat-miR156e-1 | Cat-miR156e-1 | Cat-miR156e-1 |
| Cat-miR172a-3p | Cat-miR172a-3p | Cat-miR156e-2 | Cat-miR156b | Cat-miR156e-2 | Cat-miR156e-2 | Cat-miR156e-2 |
| Cat-miR156b | Cat-NovmiR3 | Cat-miR172a-3p | Cat-miR164b | Cat-miR164b | Cat-miR172a-3p | Cat-miR172a-3p |
| | | Cat-miR156b | | Cat-NovmiR1 | Cat-miR156b | Cat-miR156b |
| | | Cat-NovmiR1 | | Cat-miR172a-5p | Cat-miR164b | Cat-miR164b |
| | | Cat-miR172a-5p | | | Cat-NovmiR3 | Cat-NovmiR3 |
| | | Cat-NovmiR3 | | | Cat-NovmiR1 | |
| | | | | | Cat-miR172a-5p | |

**Table S7. *In silico* tissue-wise expression in different tissues of chickpea**

| miRNA | Nodule | Root | Leaf | Young Pod | Flower bud | Shoot | Flower | Stem |
|---|---|---|---|---|---|---|---|---|
| cat-miR1507 | **1.406124** | 3.82749 | 5.110889 | 2.66011 | 5.511492 | 3.11211 | 6.58272 | 5.255575 |
| cat-miR399-1 | **9.139804** | 34.5634 | 28.42153 | 4.433517 | 35.6488 | 3.11211 | 47.23102 | 13.08054 |
| cat-NovmiR2 | **3.515309** | 12.17838 | 1.49587 | 1.568783 | 0 | 3.556697 | 1.069692 | 3.970879 |
| cat-miR399-2 | **9.139804** | 35.37529 | 44.00351 | 6.411548 | 37.64232 | 4.112431 | 50.60466 | 15.18277 |
| cat-miR162b | **126.5511*** | 0 | 0 | 0 | 0 | 0 | 0 | 0 |
| cat-miR399-3 | **9.139804** | 38.15892 | 27.29963 | 10.02657 | 39.7531 | 5.668486 | 49.3704 | 18.10253 |
| cat-NovmiR3 | **2.343539** | 0 | 13.33817 | 0 | 0 | 9.780916 | 1.64568 | 1.167905 |
| cat-miR171c | **690.4067*** | 0 | 0 | 0 | 0 | 0 | 0 | 0 |
| cat-miR398a-5p | **11.01464** | 7.423011 | 457.9856 | 33.14907 | 54.52859 | 19.67298 | 63.93467 | 364.7369 |
| cat-NovmiR1 | **13.59253** | 80.60926 | 7.354695 | 0 | 0 | 21.78477 | 0 | 0 |
| cat-miR171d | **20.15444** | 86.17652 | 3.490364 | 2.046239 | 2.814379 | 27.67555 | 0 | 4.20446 |
| cat-NovmiR5 | **206.2315** | 40.24664 | 21.31615 | 19.30285 | 16.76901 | 40.34628 | 24.6852 | 50.33672 |
| cat-miR2111-1 | **16.40478** | 48.24957 | 33.90639 | 5.320221 | 2.22805 | 40.90201 | 6.665004 | 11.67905 |
| cat-miR2111-5 | **16.40478** | 48.71351 | 37.14744 | 5.865884 | 2.579847 | 42.90266 | 7.899264 | 12.84696 |
| cat-miR2111-2 | **16.40478** | 49.75737 | 42.50764 | 6.138716 | 2.931645 | 55.1288 | 8.63982 | 15.88351 |

| miRNA | Nodule | Root | Leaf | Young Pod | Flower bud | Shoot | Flower | Stem |
|---|---|---|---|---|---|---|---|---|
| cat-miR2111-4 | **16.40478** | 49.98934 | 42.50764 | 6.34334 | 2.814379 | 55.46224 | 8.310684 | 16.35068 |
| cat-miR2111-3 | **19.68573** | 67.85096 | 57.21703 | 12.8231 | 3.63524 | 105.8117 | 14.97569 | 26.27787 |
| cat-miR162a | **7.030618*** | 0 | 0 | 0 | 0 | 0 | 0 | 0 |
| cat-miR166b-1 | **463.3177** | 326.2645 | 147.3432 | 99.4472 | 278.7408 | 124.0398 | 216.5715 | 194.4563 |
| cat-miR166a-1 | **5916.5** | 335.0794 | 161.0553 | 107.1547 | 291.64 | 136.2659 | 232.1232 | 207.0696 |
| cat-miR172a-3p | **0.468708** | 0 | 212.6629 | 17.05199 | 81.38246 | 147.0472 | 65.99177 | 406.8983 |
| cat-miR398b-1 | **20.38879** | 651.1373 | 1074.783 | 63.56982 | 143.6506 | 168.832 | 128.1985 | 539.4555 |
| cat-miR398b-2 | **20.38879** | 651.4852 | 1074.783 | 63.22878 | 143.6506 | 168.832 | 128.1162 | 539.3387 |
| cat-miR398a-3p | **3.280955** | 636.6392 | 1288.069 | 94.05877 | 183.7555 | 181.5027 | 183.1642 | 875.1115 |
| cat-miR171b | **4.452725** | 181.8638 | 92.74395 | 175.2945 | 370.4426 | 185.2817 | 537.3145 | 44.96436 |
| cat-miR171e | **7.499326** | 186.1552 | 92.8686 | 175.5673 | 370.6772 | 186.282 | 538.3842 | 45.54831 |
| cat-NovmiR7 | **2.109186*** | 0 | 0 | 0 | 0 | 0 | 0 | 0 |
| cat-miR390b | **558.9342** | 229.6494 | 186.9838 | 65.1386 | 178.1267 | 204.3989 | 1194.023 | 58.27848 |
| cat-miR390a-2 | **70.07183** | 247.859 | 188.2303 | 72.98251 | 240.8639 | 225.2945 | 1253.268 | 62.48294 |
| cat-miR390a-1 | **70.07183** | 248.091 | 188.4796 | 73.11893 | 241.2157 | 225.628 | 1254.913 | 62.59973 |

| miRNA | Nodule | Root | Leaf | Young Pod | Flower bud | Shoot | Flower | Stem |
|---|---|---|---|---|---|---|---|---|
| cat-NovmiR6 | **5.390141** | 12.64232 | 174.2689 | 44.06234 | 44.9128 | 230.8519 | 73.64418 | 85.02352 |
| cat-miR394-2 | **12.88947** | 163.0743 | 426.1983 | 118.8183 | 325.5298 | 231.741 | 822.0172 | 255.0705 |
| cat-miR168 | **394.1833*** | 0 | 0 | 0 | 0 | 0 | 0 | 0 |
| cat-miR172b | **1.874832** | 140.9212 | 209.6711 | 228.4967 | 415.4727 | 311.8779 | 491.2355 | 1190.796 |
| cat-NovmiR4 | **134.5192** | 365.2353 | 283.2181 | 281.6989 | 200.8763 | 353.2245 | 250.0611 | 211.858 |
| cat-miR482 | **42.18371** | 619.5895 | 982.288 | 186.3441 | 183.521 | 395.3491 | 264.5431 | 397.4382 |
| cat-miR2118 | **134.2848** | 673.1743 | 1039.256 | 215.6736 | 204.5115 | 450.4779 | 299.9252 | 480.7099 |
| cat-miR394-1 | **12.88947** | 323.3649 | 837.5626 | 232.9984 | 639.8022 | 455.0349 | 1597.708 | 500.4475 |
| cat-miR5213 | **6.093203** | 191.4905 | 475.562 | 283.5405 | 288.9429 | 470.8177 | 386.8171 | 595.2814 |
| cat-miR1511 | **2943.251** | 440.5093 | 386.5578 | 397.516 | 517.3767 | 494.3809 | 508.5974 | 689.181 |
| cat-miR164b | **2.812247** | 747.2885 | 0 | 1143.37 | 0 | 575.7403 | 0 | 1330.945 |
| cat-miR164a-3 | **191.4672** | 766.1939 | 118.9217 | 1192.548 | 154.439 | 596.0802 | 366.8221 | 1375.676 |
| cat-miR164a-2 | **192.4046** | 768.1657 | 119.4203 | 1187.569 | 154.9081 | 600.3037 | 367.2335 | 1381.399 |
| cat-miR164a-1 | **193.8107** | 808.4123 | 122.412 | 1255.572 | 157.3707 | 615.4197 | 379.7407 | 1486.276 |
| cat-miR171f | **92.1011** | 223.8502 | 229.8654 | 188.254 | 730.8004 | 617.8649 | 757.3419 | 194.573 |
| cat-miR171a | **3.280955** | 208.6562 | 238.2173 | 170.3153 | 730.6832 | 636.5376 | 776.5964 | 183.4779 |
| cat-miR160-3 | **63.97863** | 1475.092 | 414.6053 | 321.6005 | 400.8145 | 805.3695 | 455.0305 | 477.323 |
| cat-miR160-1 | **64.21298** | 1476.715 | 415.1039 | 321.4641 | 401.4008 | 806.3699 | 455.6888 | 477.323 |
| cat-miR160-2 | **64.21298** | 1499.1 | 426.323 | 326.1022 | 404.8015 | 827.1543 | 459.5561 | 488.4181 |
| cat-miR172c-1 | **2.812247** | 276.2752 | 4216.858 | 293.226 | 618.4598 | 887.9516 | 666.1713 | 2509.595 |
| cat-miR172c-2 | **2.812247** | 276.2752 | 4216.858 | 293.226 | 618.4598 | 887.9516 | 666.1713 | 2509.595 |
| cat-miR172a-5p | **0.468708** | 274.3035 | 4220.597 | 0 | 0 | 890.3968 | 0 | 0 |
| cat-miR319d | **3301.813** | 545.2434 | 181.6236 | 141.1223 | 61.44728 | 1178.712 | 70.35282 | 2322.964 |
| cat-miR166b-3 | **459.5681** | 2611.74 | 864.9869 | 616.941 | 660.2064 | 1203.72 | 764.8298 | 1348.931 |
| cat-miR1509 | **793.2881** | 1034.234 | 1160.047 | 608.0057 | 777.3549 | 1389.224 | 633.5045 | 1038.151 |
| cat-miR166a-4 | **4477.332** | 3027.893 | 927.8134 | 1041.945 | 1217.571 | 1829.254 | 1154.033 | 2052.243 |
| cat-miR396b | **1805.697** | 3440.914 | 2477.036 | 830.2273 | 490.9919 | 1922.395 | 748.0438 | 3246.076 |
| cat-miR166a-3 | **5197.267** | 3040.767 | 928.5614 | 1209.873 | 1410.942 | 2001.865 | 1292.682 | 2274.262 |
| cat-miR166a-2 | **5197.267** | 3040.187 | 928.8107 | 1209.736 | 1410.707 | 2002.087 | 1292.353 | 2274.145 |
| cat-miR166b-2 | **599.0087** | 3066.863 | 932.301 | 1208.645 | 1408.949 | 2012.424 | 1294.739 | 2274.145 |
| cat-miR166a-5 | **6064.143** | 3072.431 | 935.2928 | 1212.942 | 1415.281 | 2021.76 | 1307.657 | 2276.131 |
| cat-miR396a | **1280.979** | 3483.828 | 2546.594 | 876.6087 | 510.458 | 2026.539 | 775.1153 | 3343.713 |

| miRNA | Nodule | Root | Leaf | Young Pod | Flower bud | Shoot | Flower | Stem |
|---|---|---|---|---|---|---|---|---|
| cat-miR408 | **9.608512** | 1824.785 | 1122.277 | 847.4157 | 578.2376 | 2101.008 | 176.5815 | 1973.76 |
| cat-miR319a | **325.9863** | 1484.602 | 588.5002 | 538.229 | 1149.674 | 2106.787 | 2182.336 | 2873.631 |
| cat-miR319c-4 | **1672.35** | 1772.824 | 187.1084 | 477.7967 | 1698.478 | 2336.972 | 2740.386 | 7340.753 |
| cat-miR319c-1 | **1671.412** | 2166.475 | 236.5968 | 532.7724 | 1795.808 | 2866.031 | 2878.294 | 8859.964 |
| cat-miR319c-2 | **1692.504** | 2181.901 | 238.5913 | 534.8186 | 1798.857 | 2888.816 | 2886.44 | 8903.644 |
| cat-miR319b | **5802.135** | 2637.953 | 262.5252 | 587.5433 | 2067.513 | 2906.933 | 3023.608 | 7667.766 |
| cat-miR319c-3 | **1615.402** | 2100.828 | 700.3165 | 633.0381 | 1826.532 | 3105.663 | 2976.13 | 7561.721 |
| cat-miR396c | **2992.934** | 3981.75 | 4272.454 | 1481.34 | 880.3143 | 3125.781 | 1416.93 | 5111.221 |
| cat-miR156b | **4.218371** | 0 | 3577.623 | 54.22533 | 29.08192 | 3195.136 | 0 | 184.6459 |
| cat-miR156f-2 | **63.74427*** | 0 | 0 | 0 | 0 | 0 | 0 | 0 |
| cat-miR156f-1 | **63.74427*** | 0 | 0 | 0 | 0 | 0 | 0 | 0 |
| cat-miR156c-2 | **780.8673** | 892.1532 | 3697.168 | 59.54555 | 32.83442 | 3335.292 | 46.32589 | 195.9745 |
| cat-miR156c-4 | **771.9619** | 892.0372 | 3698.663 | 57.4311 | 31.8963 | 3335.737 | 45.58534 | 194.6898 |
| cat-miR167c | **65.6191** | 4265.912 | 5868.672 | 3290.898 | 1645.474 | 3344.407 | 1251.951 | 175.1858 |
| cat-miR167a-1 | **246.0716** | 4266.492 | 5868.922 | 3288.101 | 1642.425 | 3347.296 | 1251.622 | 174.2515 |
| cat-miR167d | **157.0171** | 4274.611 | 5880.016 | 3307.063 | 1648.523 | 3352.409 | 1257.546 | 173.6675 |
| cat-miR167a-2 | **242.5563** | 4262.432 | 5866.297 | 3233.262 | 1616.978 | 3359.078 | 1232.203 | 172.4996 |
| cat-miR167b-1 | **2722.958** | 4307.086 | 5927.884 | 3288.647 | 1641.956 | 3403.425 | 1262.319 | 175.5362 |
| cat-miR167b-2 | **2722.958** | 4314.625 | 5929.006 | 3293.558 | 1648.288 | 3409.316 | 1266.351 | 177.4048 |
| cat-miR156c-1 | **534.7957** | 1254.953 | 5600.164 | 69.09466 | 40.1049 | 3562.476 | 52.00349 | 197.4928 |
| cat-miR156c-3 | **530.8117** | 1258.2 | 5629.956 | 71.55015 | 38.93224 | 3565.477 | 53.40232 | 193.4051 |
| cat-miR156a-3 | **33.74697** | 1867.119 | 3627.36 | 188.595 | 280.148 | 4606.478 | 265.7773 | 336.24 |
| cat-miR156a-1 | **42.88677** | 1950.28 | 3624.369 | 175.9083 | 260.0955 | 4614.814 | 263.7202 | 328.8822 |
| cat-miR156a-2 | **34.45003** | 1965.59 | 3654.535 | 179.0459 | 264.4344 | 4670.499 | 268.1636 | 333.437 |
| cat-miR156e-1 | **253.1023** | 1987.395 | 3679.092 | 0 | 314.2723 | 4715.291 | 304.2039 | 369.4085 |
| cat-miR156e-2 | **66.32217** | 1990.527 | 3679.466 | 0 | 314.0378 | 4716.847 | 303.7925 | 370.5764 |
| cat-miR156d | **8.436742** | 1905.626 | 4690.051 | 174.4077 | 259.6265 | 5388.84 | 261.0048 | 346.1672 |
| cat-miR159-1 | **32815.41** | 30492.22 | 41974.74 | 29411.07 | 13492.48 | 78033.15 | 19847.07 | 50284.75 |
| cat-miR159-2 | **32815.41** | 30492.22 | 41974.74 | 29411.07 | 13492.48 | 78033.15 | 19847.07 | 50284.75 |

Highlighted rows are high expressing in nodule; asterisks are only expressing in nodule

**Table S8. The list of expression of microRNA in nodule tissue in comparison to root**

| miRs | Nodule | Root | Fold Change Nodule vs Root |
|---|---|---|---|
| cat-miR172a-5p | **0.468708** | 274.3035 | 0.001709 |
| cat-miR164b | **2.812247** | 747.2885 | 0.003763 |
| cat-miR156d | **8.436742** | 1905.626 | 0.004427 |
| cat-miR398a-3p | **3.280955** | 636.6392 | 0.005154 |
| cat-miR408 | **9.608512** | 1824.785 | 0.005266 |
| cat-miR172c-1 | **2.812247** | 276.2752 | 0.010179 |
| cat-miR172c-2 | **2.812247** | 276.2752 | 0.010179 |
| cat-miR172b | **1.874832** | 140.9212 | 0.013304 |
| cat-miR167c | **65.6191** | 4265.912 | 0.015382 |
| cat-miR171a | **3.280955** | 208.6562 | 0.015724 |
| cat-miR156a-2 | **34.45003** | 1965.59 | 0.017527 |
| cat-miR156a-3 | **33.74697** | 1867.119 | 0.018074 |
| cat-miR156a-1 | **42.88677** | 1950.28 | 0.02199 |
| cat-miR171b | **4.452725** | 181.8638 | 0.024484 |
| cat-miR398b-2 | **20.38879** | 651.4852 | 0.031296 |
| cat-miR398b-1 | **20.38879** | 651.1373 | 0.031313 |
| cat-miR5213 | **6.093203** | 191.4905 | 0.03182 |
| cat-miR156e-2 | **66.32217** | 1990.527 | 0.033319 |
| cat-miR167d | **157.0171** | 4274.611 | 0.036733 |
| cat-miR394-1 | **12.88947** | 323.3649 | 0.03986 |
| cat-miR171e | **7.499326** | 186.1552 | 0.040285 |
| cat-miR160-2 | **64.21298** | 1499.1 | 0.042834 |
| cat-miR160-3 | **63.97863** | 1475.092 | 0.043373 |
| cat-miR160-1 | **64.21298** | 1476.715 | 0.043484 |

| miRs | Nodule | Root | Fold Change Nodule vs Root |
|---|---|---|---|
| cat-miR167a-2 | **242.5563** | 4262.432 | 0.056906 |
| cat-miR167a-1 | **246.0716** | 4266.492 | 0.057675 |
| cat-miR482 | **42.18371** | 619.5895 | 0.068083 |
| cat-miR394-2 | **12.88947** | 163.0743 | 0.07904 |
| cat-miR156e-1 | **253.1023** | 1987.395 | 0.127354 |
| cat-NovmiR1 | **13.59253** | 80.60926 | 0.168622 |
| cat-miR166b-3 | **459.5681** | 2611.74 | 0.175962 |
| cat-miR166b-2 | **599.0087** | 3066.863 | 0.195316 |
| cat-miR2118 | **134.2848** | 673.1743 | 0.19948 |
| cat-miR319a | **325.9863** | 1484.602 | 0.219578 |
| cat-miR171d | **20.15444** | 86.17652 | 0.233874 |
| cat-miR399-3 | **9.139804** | 38.15892 | 0.239519 |
| cat-miR164a-1 | **193.8107** | 808.4123 | 0.239742 |
| cat-miR164a-3 | **191.4672** | 766.1939 | 0.249894 |
| cat-miR164a-2 | **192.4046** | 768.1657 | 0.250473 |
| cat-miR399-2 | **9.139804** | 35.37529 | 0.258367 |
| cat-miR399-1 | **9.139804** | 34.5634 | 0.264436 |
| cat-miR390a-1 | **70.07183** | 248.091 | 0.282444 |
| cat-miR390a-2 | **70.07183** | 247.859 | 0.282708 |
| cat-NovmiR2 | **3.515309** | 12.17838 | 0.288652 |
| cat-miR2111-3 | **19.68573** | 67.85096 | 0.290132 |
| cat-miR2111-4 | **16.40478** | 49.98934 | 0.328165 |
| cat-miR2111-2 | **16.40478** | 49.75737 | 0.329695 |
| cat-miR2111-5 | **16.40478** | 48.71351 | 0.33676 |

| miRs | Nodule | Root | Fold Change Nodule vs Root |
|---|---|---|---|
| cat-miR2111-1 | **16.40478** | 48.24957 | 0.339998 |
| cat-miR1507 | **1.406124** | 3.82749 | 0.367375 |
| cat-miR396a | **1280.979** | 3483.828 | 0.367693 |
| cat-NovmiR4 | **134.5192** | 365.2353 | 0.368308 |
| cat-miR171f | **92.1011** | 223.8502 | 0.411441 |
| cat-miR156c-3 | **530.8117** | 1258.2 | 0.421882 |
| cat-miR156c-1 | **534.7957** | 1254.953 | 0.426148 |
| cat-NovmiR6 | **5.390141** | 12.64232 | 0.426357 |
| cat-miR396b | **1805.697** | 3440.914 | 0.524773 |
| cat-miR167b-2 | **2722.958** | 4314.625 | 0.6311 |
| cat-miR167b-1 | **2722.958** | 4307.086 | 0.632204 |
| cat-miR396c | **2992.934** | 3981.75 | 0.751663 |
| cat-miR1509 | **793.2881** | 1034.234 | 0.767029 |
| cat-miR319c-3 | **1615.402** | 2100.828 | 0.768936 |
| cat-miR319c-1 | **1671.412** | 2166.475 | 0.771489 |
| cat-miR319c-2 | **1692.504** | 2181.901 | 0.775702 |
| cat-miR156c-4 | **771.9619** | 892.0372 | 0.865392 |
| cat-miR156c-2 | **780.8673** | 892.1532 | 0.875262 |
| cat-miR319c-4 | **1672.35** | 1772.824 | 0.943325 |
| cat-miR159-1 | **32815.41** | 30492.22 | 1.07619 |
| cat-miR159-2 | **32815.41** | 30492.22 | 1.07619 |
| cat-miR166b-1 | **463.3177** | 326.2645 | 1.420068 |
| cat-miR166a-4 | **4477.332** | 3027.893 | 1.478696 |
| cat-miR398a-5p | **11.01464** | 7.423011 | 1.48385 |

| miRs | Nodule | Root | Fold Change Nodule vs Root |
|---|---|---|---|
| cat-miR166a-3 | **5197.267** | 3040.767 | 1.709196 |
| cat-miR166a-2 | **5197.267** | 3040.187 | 1.709522 |
| cat-miR166a-5 | **6064.143** | 3072.431 | 1.973728 |
| cat-miR319b | **5802.135** | 2637.953 | 2.199484 |
| cat-miR390b | **558.9342** | 229.6494 | 2.433858 |
| cat-NovmiR5 | **206.2315** | 40.24664 | 5.124191 |
| cat-miR319d | **3301.813** | 545.2434 | 6.055668 |
| cat-miR1511 | **2943.251** | 440.5093 | 6.681473 |
| cat-miR166a-1 | **5916.5** | 335.0794 | 17.65701 |
| cat-miR162b | **126.5511** | 0 | |
| cat-NovmiR3 | **2.343539** | 0 | |
| cat-miR171c | **690.4067** | 0 | |
| cat-miR162a | **7.030618** | 0 | |
| cat-miR172a-3p | **0.468708** | 0 | |
| cat-NovmiR7 | **2.109186** | 0 | |
| cat-miR168 | **394.1833** | 0 | |
| cat-miR156b | **4.218371** | 0 | |
| cat-miR156f-2 | **63.74427** | 0 | |
| cat-miR156f-1 | **63.74427** | 0 | |

(green) > 10 fold down regulated in nodule tissue
(yellow) > 10 fold up regulated in nodule tissue

**Table S9. The information of miRNA, target, annotation and category**

| miR | Target | Target Annotation | Category |
|---|---|---|---|
| **Early nodule** | | | |
| cat-miR319d | Ca1_48624443-48627841 | PREDICTED: transcription factor TCP4-like | 1 |
| cat-miR319b | Ca1_3936259-3937645 | PREDICTED: transcription factor TCP2 | 1 |
| cat-miR1507 | Ca4_12620685-12623805 | #N/A | 2 |
| cat-miR1507 | Ca4_56032289-56034688 | subtilisin-like protease SBT1.7 | 2 |
| cat-miR1509 | Ca4_848277-850064 | PREDICTED: protein JASON | 1 |
| cat-miR1509 | Ca1_12076586-12079921 | PREDICTED: adenylate kinase isoenzyme 6 homolog | 1 |
| cat-miR1509 | Ca7_48336341-48337063 | 0 | 2 |
| cat-miR1509 | Ca4_4297171-4301616 | PREDICTED: U11/U12 smalluclear ribonucleoprotein 48 kDa protein | 2 |
| cat-miR1509 | Ca6_885909-888498 | PREDICTED: adenylate kinase isoenzyme 6 homolog | 2 |
| cat-miR1511 | Ca5_16064453-16073091 | ER membrane protein complex subunit-like protein | 2 |
| cat-miR156a-1 | Ca2_17579240-17596608 | PREDICTED: probable leucine-rich repeat receptor-like serine/threonine-protein kinase At3g14840 | 2 |
| cat-miR156a-1 | Ca6_49578548-49588910 | PREDICTED: protein ROS1-like | 2 |
| cat-miR156a-1 | Ca3_32941384-32944363 | Acyl-CoA thioesterase, putative | 2 |
| cat-miR156a-1 | Ca3_40920821-40927671 | PREDICTED: ethylene-insensitive protein 2 | 4 |
| cat-miR156a-2 | Ca8_10742165-10752979 | AGC family Serine/Threonine kinase family protein | 2 |
| cat-miR156a-2 | Ca4_7628103-7654987 | PREDICTED: uncharacterized protein LOC101497938 | 2 |
| cat-miR156a-2 | Ca4_28830181-28838574 | PREDICTED: protein EMSY-LIKE 1 | 2 |
| cat-miR156a-3 | Ca7_47840061-47910402 | Calcium-dependent lipid-binding-like protein | 2 |
| cat-miR156a-3 | Ca5_21732086-21737394 | PREDICTED: uncharacterized protein LOC101494695 isoform X1 | 2 |
| cat-miR156a-3 | Ca2_3794978-3800049 | PREDICTED: LOW QUALITY PROTEIN: squamosa promoter-binding-like protein 6 | 4 |
| cat-miR156b | Ca7_54854284-54857859 | PREDICTED: uncharacterized protein LOC101499362 | 2 |
| cat-miR156b | Ca8_2794259-2803059 | Lon protease homolog 2, peroxisomal | 2 |
| cat-miR156b | Ca7_51830319-51831891 | PREDICTED: uncharacterized protein LOC101514397 | 4 |
| cat-miR156c-2 | Ca8_1066045-1068026 | PREDICTED: uncharacterized protein LOC101490125 | 2 |
| cat-miR156c-2 | Ca8_12188165-12275518 | Plant synaptotagmin | 4 |
| cat-miR156c-3 | Ca6_20650286-20653460 | PREDICTED: ATPase ASNA1 homolog | 4 |
| cat-miR156c-3 | Ca3_32989731-32993382 | Uncharacterized protein | 3 |
| cat-miR156c-4 | Ca5_16160180-16163354 | Coiled-coil domain-containing protein | 3 |
| cat-miR156c-4 | Ca8_3496746-3501741 | PREDICTED: BTB/POZ and TAZ domain-containing protein 4 | 4 |
| cat-miR156d | Ca6_20650286-20653460 | PREDICTED: ATPase ASNA1 homolog | 2 |
| cat-miR156d | Ca2_14022347-14025926 | PREDICTED: protein NETWORKED 4A-like | 2 |
| cat-miR156d | Ca8_12188165-12275518 | Plant synaptotagmin | 2 |
| cat-miR156d | Ca6_5704788-5709534 | PREDICTED: protein IQ-DOMAIN 14 | 2 |
| cat-miR156d | Ca4_15483995-15488401 | PREDICTED: cysteine--tRNA ligase, cytoplasmic | 2 |
| cat-miR156d | Ca6_23638344-23655362 | Serine/threonine-protein phosphatase | 2 |
| cat-miR156d | Ca7_12558028-12561380 | PREDICTED: uncharacterized protein LOC101498996 | 2 |
| cat-miR156d | Ca7_50193176-50199734 | Probable sucrose-phosphate synthase | 2 |
| cat-miR156d | Ca3_58041311-58055198 | Actin-97 | 2 |
| cat-miR156d | Ca3_58041311-58055198 | Actin-97 | 2 |
| cat-miR156d | Ca4_41546915-41550573 | PREDICTED: high mobility group B protein 6-like | 2 |
| cat-miR156d | Ca6_63314526-63322921 | Uncharacterized protein | 2 |
| cat-miR156d | Ca4_42250206-42251142 | PREDICTED: uncharacterized protein LOC101509491 | 2 |
| cat-miR156d | Ca4_54328033-54332303 | PREDICTED: probable protein phosphatase 2C 59 | 2 |
| cat-miR156d | Ca1_3908972-3925137 | Uncharacterized protein | 2 |
| cat-miR156d | Ca5_11061294-11065235 | PREDICTED: DEAD-box ATP-dependent RNA helicase 18 | 2 |
| cat-miR156d | Ca5_45301553-45334438 | PREDICTED: formin-like protein 20 isoform X1 | 4 |
| cat-miR156d | Ca8_704872-710512 | PREDICTED: putative 12-oxophytodienoate reductase 11 | 3 |
| cat-miR156d | Ca1_9464244-9467765 | PREDICTED: uncharacterized protein LOC101506888 | 4 |
| cat-miR156d | Ca4_3948179-3958105 | Serine/Threonine kinase family protein | 4 |
| cat-miR156e-1 | Ca5_18544803-18546508 | #N/A | 0 |
| cat-miR156e-1 | Ca8_4552808-4554903 | #N/A | 2 |
| cat-miR156e-1 | Ca7_51171571-51188309 | #N/A | 2 |
| cat-miR156e-1 | Ca6_57031140-57042957 | PREDICTED: protein transport protein SEC31 homolog B | 2 |
| cat-miR156e-1 | Ca5_20201398-20203182 | #N/A | 2 |
| cat-miR156e-1 | Ca4_4231486-4243578 | #N/A | 2 |
| cat-miR156e-1 | Ca5_20823051-20831241 | PREDICTED: telomere length regulation protein TEL2 homolog | 2 |
| cat-miR156e-2 | Ca1_11776043-11780075 | #N/A | 2 |
| cat-miR156e-2 | Ca2_19636676-19639510 | #N/A | 2 |
| cat-miR156e-2 | Ca5_17944777-17949876 | Eukaryotic translation initiation factor 3 subunit B | 2 |
| cat-miR156e-2 | Ca6_29033826-29037684 | #N/A | 2 |
| cat-miR156f-1 | Ca4_4145536-4155030 | #N/A | 4 |
| cat-miR156f-2 | Ca4_16113171-16120762 | PREDICTED: putative transcription elongation factor SPT5 homolog 1 | 2 |
| cat-miR156f-2 | Ca5_20201398-20203182 | #N/A | 3 |
| cat-miR159-2 | Ca1_15593646-15596122 | 50S ribosomal L24-like protein | 2 |
| cat-miR162b | Scaffold0592_921962-925549 | Uncharacterized protein | 2 |
| cat-miR162b | Ca5_27769382-27784627 | PREDICTED: MAG2-interacting protein 2 | 4 |
| cat-miR162b | Ca4_13381070-13385617 | PREDICTED: primary amine oxidase-like | 2 |
| cat-miR162b | Ca2_29208613-29238954 | Uncharacterized protein | 3 |
| cat-miR162b | Ca5_27125082-27127900 | PREDICTED: probable inactive receptor kinase At4g23740 | 4 |
| cat-miR164a-2 | Ca2_10486844-10507678 | PREDICTED: MADS-box transcription factor 23-like isoform X3 | 2 |
| cat-miR164a-2 | Ca6_9175135-9178533 | Impaired sucrose induction protein, putative | 2 |
| cat-miR164a-2 | Ca4_8115048-8127026 | GroEL-like chaperone, ATPase | 2 |
| cat-miR164a-2 | Ca7_51381378-51385717 | PREDICTED: protein Dr1 homolog isoform X1 | 2 |
| cat-miR164a-2 | Ca8_12928494-12931984 | PREDICTED: proline-rich receptor-like protein kinase PERK1 | 2 |
| cat-miR164a-3 | Ca1_39628189-39632859 | NAC transcription factor | 3 |
| cat-miR164b | Ca6_15230790-15233212 | 1-aminocyclopropane-1-carboxylate synthase | 3 |
| cat-miR164b | Ca1_9876547-9880271 | ATP-dependent 6-phosphofructokinase | 4 |
| cat-miR164b | Ca4_36083974-36091304 | PREDICTED: uncharacterized protein LOC101491241 isoform X1 | 0 |
| cat-miR164b | Ca6_39786200-39790820 | Splicing factor-like protein | 2 |
| cat-miR164b | Ca4_12071292-12077676 | PREDICTED: titin-like | 2 |
| cat-miR164b | Ca5_65215170-65217442 | PREDICTED: NAD(P)H-dependent 6""-deoxychalcone synthase | 2 |
| cat-miR164b | Ca6_12344042-12349655 | Sel1 repeat protein | 2 |
| cat-miR164b | Scaffold3523_21890-26350 | PREDICTED: purple acid phosphatase 18-like isoform X1 | 2 |
| cat-miR164b | Ca4_12071292-12077676 | PREDICTED: titin-like | 2 |
| cat-miR164b | Ca7_50532288-50533892 | PREDICTED: blue copper protein-like | 2 |
| cat-miR164b | Ca7_50113541-50115041 | PREDICTED:ucleolin-like | 2 |
| cat-miR164b | Ca1_27442502-27449511 | PREDICTED: DEAD-box ATP-dependent RNA helicase 51 | 2 |
| cat-miR164b | Ca2_13066142-13068074 | PREDICTED: exocyst complex component EXO70B1-like | 2 |
| cat-miR164b | Ca6_7219977-7224962 | Uncharacterized protein | 2 |
| cat-miR164b | Ca4_15973579-15984001 | PREDICTED: serine/threonine-protein kinase prpf4B | 2 |
| cat-miR164b | Ca4_18391305-18393071 | 0 | 2 |
| cat-miR164b | Ca4_18391220-18395966 | PREDICTED: extensin-2 | 2 |
| cat-miR164b | Ca6_2731520-2734216 | PREDICTED: uncharacterized protein LOC101491587 | 2 |
| cat-miR164b | Ca4_11360318-11362309 | 0 | 2 |
| cat-miR164b | Ca1_14649449-14663687 | PREDICTED: protein TIC 100 | 2 |
| cat-miR164b | Ca4_57482015-57488353 | PREDICTED: auxilin-like protein 1 isoform X1 | 2 |
| cat-miR164b | Ca1_6530616-6539298 | PREDICTED: enhancer of mRNA-decapping protein 4-like isoform X1 | 2 |
| cat-miR164b | Ca5_22508111-22512378 | PREDICTED: flowering time control protein FPA isoform X1 | 2 |
| cat-miR164b | Ca3_39736284-39739002 | PREDICTED: rho GDP-dissociation inhibitor 1-like | 2 |
| cat-miR164b | Ca3_58667250-58671227 | TPR protein | 2 |
| cat-miR164b | Ca1_10230562-10235575 | Casein kinase I-like protein | 2 |
| cat-miR164b | Scaffold0590_52293-60863 | Beta-amyrin synthase | 2 |
| cat-miR164b | Ca7_55602199-55605689 | BTB/POZ and TAZ domain protein | 2 |
| cat-miR164b | Ca6_3567998-3572811 | PREDICTED: uncharacterized protein LOC101502776 isoform X1 | 2 |
| cat-miR164b | Ca8_13921235-13925887 | PREDICTED: thioredoxin-like 2, chloroplastic | 2 |
| cat-miR164b | Ca1_16066254-16071720 | PREDICTED: probable ADP-ribosylation factor GTPase-activating protein AGD5 isoform X1 | 2 |
| cat-miR164b | Ca6_14387825-14388800 | PREDICTED: uncharacterized protein LOC101506109 | 2 |

| miR | Target | Target Annotation | Category |
|---|---|---|---|
| cat-miR164b | Ca1_9247901-9254902 | Drug resistance transporter-like ABC domain protein | 2 |
| cat-miR164b | Ca6_15728084-15732728 | PREDICTED: myosin-11-like | 2 |
| cat-miR164b | Ca4_16070652-16075435 | PREDICTED: glycine-rich RNA-binding protein RZ1B isoform X1 | 2 |
| cat-miR164b | Ca5_13861235-14094850 | Niemann-Pick C1 protein | 2 |
| cat-miR164b | Ca8_7907544-7912538 | PREDICTED: uncharacterized protein LOC101507671 | 2 |
| cat-miR164b | Ca7_44786154-44790831 | PREDICTED: aspartic proteinase-like | 2 |
| cat-miR164b | Ca6_48150860-48156205 | PREDICTED: uncharacterized protein LOC101504845 | 2 |
| cat-miR164b | Ca1_17810654-17814971 | Uncharacterized protein | 2 |
| cat-miR164b | Ca7_21948765-21949374 | PREDICTED: dof zinc finger protein DOF1.7-like | 2 |
| cat-miR164b | Ca4_5699940-5701446 | Ribosomal L22e family protein | 2 |
| cat-miR164b | Ca6_65079202-65085741 | NADH-ubiquinone oxidoreductase 75 kDa subunit | 2 |
| cat-miR164b | Ca4_40696172-40697926 | PREDICTED: tankyrase | 2 |
| cat-miR164b | Ca4_6655508-6656242 | PREDICTED: protein RADIALIS-like 3 | 4 |
| cat-miR164b | Ca7_33354243-33359696 | PREDICTED: uncharacterized protein LOC101494421 | 4 |
| cat-miR166a-4 | Ca2_10846940-10850162 | PREDICTED: auxin-responsive protein IAA8-like | 2 |
| cat-miR166a-4 | Ca3_33420766-33427344 | COP9 signalosome complex subunit-like protein | 2 |
| cat-miR166a-4 | Ca8_7351796-7363853 | PREDICTED: probable glucan 1,3-alpha-glucosidase isoform X1 | 2 |
| cat-miR166a-4 | Ca7_53730248-53735086 | PREDICTED: putative dual specificity protein phosphatase DSP8 | 2 |
| cat-miR166a-5 | Ca1_7078032-7086500 | PREDICTED: 30-kDa cleavage and polyadenylation specificity factor 30 | 2 |
| cat-miR166a-5 | Ca4_38849643-38852342 | PREDICTED: glucan endo-1,3-beta-glucosidase 5 | 2 |
| cat-miR166b-1 | Ca6_7391482-7402641 | Pyruvate kinase | 2 |
| cat-miR166b-1 | Ca2_36592665-36595595 | PREDICTED: uncharacterized protein LOC101489688 | 2 |
| cat-miR166b-1 | Ca7_31017195-31020568 | Uncharacterized protein | 2 |
| cat-miR166b-2 | Ca5_7142949-7143897 | PREDICTED: uncharacterized protein LOC101502143 | 2 |
| cat-miR166b-2 | Ca7_48670948-48678394 | Kinesin-like protein | 2 |
| cat-miR166b-2 | Ca2_30205339-30209864 | PREDICTED: KH domain-containing protein At4g18375 isoform X2 | 2 |
| cat-miR166b-2 | Ca1_8841297-8852892 | Uncharacterized protein | 2 |
| cat-miR166b-3 | Ca5_27571200-27578359 | PP2A regulatory subunit TAP46-like protein | 2 |
| cat-miR166b-3 | Ca3_45664654-45668033 | General substrate transporter | 2 |
| cat-miR166b-3 | Scaffold4867_464840-474215 | PREDICTED: U1 smalluclear ribonucleoprotein A | 4 |
| cat-miR167a-2 | Ca6_60959805-60962198 | PREDICTED: probable cysteine proteinase A494 | 2 |
| cat-miR167a-2 | Ca3_49651605-49656657 | PREDICTED: uncharacterized serine-rich protein C215.13-like | 2 |
| cat-miR167a-2 | Ca6_10350393-10367492 | PREDICTED: ENHANCER OF AG-4 protein 2, partial | 2 |
| cat-miR167a-2 | Ca1_6955157-6967502 | PREDICTED: tubulin-folding cofactor E-like | 4 |
| cat-miR167b-1 | Ca5_38602945-38605542 | PREDICTED: uncharacterized endoplasmic reticulum membrane protein C16E8.02-like | 2 |
| cat-miR167b-1 | Ca3_51465255-51472649 | PREDICTED: protein STRUBBELIG-RECEPTOR FAMILY 6 | 2 |
| cat-miR167b-2 | Ca6_63308083-63310409 | Tubulin alpha-5 | 2 |
| cat-miR167b-2 | Ca2_10364324-10370128 | PREDICTED: E3 ubiquitin-protein ligase UPL1-like | 4 |
| cat-miR167c | Ca1_11493121-11497391 | Uncharacterized protein | 2 |
| cat-miR167c | Ca3_47670916-47674120 | PREDICTED: proton-coupled amino acid transporter 3 isoform X1 | 2 |
| cat-miR167c | Ca6_63719084-63723495 | PREDICTED: probable magnesium transporter NIPA1 | 2 |
| cat-miR167c | Ca3_35838616-35848681 | #N/A | 2 |
| cat-miR167c | Ca2_33616402-33617797 | #N/A | 2 |
| cat-miR167c | Ca4_9357210-9365840 | #N/A | 2 |
| cat-miR167c | Ca5_19773896-19787589 | NADPH--cytochrome P450 reductase | 4 |
| cat-miR167d | Ca7_51269864-51282369 | PREDICTED: transcription initiation factor TFIID subunit 2 isoform X1 | 2 |
| cat-miR167d | Ca4_55408064-55412024 | #N/A | 2 |
| cat-miR168 | Ca6_28321742-28324304 | #N/A | 2 |
| cat-miR168 | Ca1_757386-761417 | #N/A | 2 |
| cat-miR168 | Ca4_16248874-16251793 | #N/A | 2 |
| cat-miR168 | Ca4_17219946-17227299 | #N/A | 2 |
| cat-miR168 | Ca4_14251980-14254921 | #N/A | 2 |
| cat-miR171a | Ca3_58041311-58055198 | Actin-97 | 2 |
| cat-miR171a | Ca7_47326242-47331530 | Sucrose synthase | 2 |
| cat-miR171a | Ca6_12344042-12349655 | Sel1 repeat protein | 2 |
| cat-miR171a | Ca1_4577000-4583210 | PREDICTED: squamosa promoter-binding-like protein 7 | 2 |

| miR | Target | Target Annotation | Category |
|---|---|---|---|
| cat-miR171a | Ca7_41378626-41381714 | PREDICTED: MLP-like protein 34 | 2 |
| cat-miR171a | Ca7_50532288-50533892 | PREDICTED: blue copper protein-like | 2 |
| cat-miR171a | Scaffold0539_951-1631 | PREDICTED: protein YIPF1 homolog | 2 |
| cat-miR171a | Ca3_49880382-49888862 | Protein YIPF | 2 |
| cat-miR171b | Ca1_11929778-11933751 | PREDICTED: protein disulfide isomerase-like 1-4 | 2 |
| cat-miR171b | Ca2_12720759-12723193 | Uncharacterized protein | 2 |
| cat-miR171b | Scaffold0592_98689-99652 | Putative mitochondrial dicarboxylate carrier protein | 2 |
| cat-miR171b | Ca1_8874995-8876253 | PREDICTED: protein LURP-one-related 5-like | 2 |
| cat-miR171b | Ca5_13250526-13262167 | Kinesin-like protein | 2 |
| cat-miR171b | Ca6_5272901-5275964 | PREDICTED: cellulose synthase-like protein G2 | 4 |
| cat-miR171c | Ca4_56157637-56159879 | PREDICTED: F-box protein At1g47056-like | 2 |
| cat-miR171c | Ca4_55615010-55616659 | PREDICTED: U-box domain-containing protein 30-like | 2 |
| cat-miR171c | Ca2_21771031-21774610 | PREDICTED: cytochrome c oxidase assembly protein COX15 | 2 |
| cat-miR171c | Ca2_19048159-19050842 | PREDICTED: UBP1-associated protein 2B-like | 2 |
| cat-miR171c | Ca4_8311023-8318521 | Uncharacterized protein | 4 |
| cat-miR171d | Ca6_9003418-9008245 | PREDICTED: switch 2 | 2 |
| cat-miR171d | Ca7_9465647-9470859 | Putative uncharacterized protein (Fragment) | 2 |
| cat-miR171d | Ca2_16538767-16542277 | Uncharacterized protein | 2 |
| cat-miR171d | Ca2_22157858-22160693 | Emp24/gp25L/p24 family protein | 2 |
| cat-miR171d | Ca8_15447348-15448844 | PREDICTED: heat stress transcription factor B-2a-like | 2 |
| cat-miR171e | Ca5_3849972-3852598 | PREDICTED: serine/threonine-protein phosphatase 7 long form homolog isoform X1 | 2 |
| cat-miR171e | Ca5_9952938-9963653 | Glutathione S-transferase/chloride channel, C-terminal | 2 |
| cat-miR171e | Ca1_2889341-2889854 | PREDICTED: pathogenesis-related genes transcriptional activator PTI5-like | 2 |
| cat-miR171e | Ca6_23768274-23771391 | Putative uncharacterized protein | 2 |
| cat-miR171e | Ca5_29055856-29059960 | PREDICTED: uncharacterized protein LOC101498857 isoform X1 | 2 |
| cat-miR171e | Ca2_8028916-8034293 | PREDICTED: thylakoid lumenal 29 kDa protein, chloroplastic isoform X1 | 2 |
| cat-miR171e | Ca5_12422255-12435106 | P-loopucleoside triphosphate hydrolase superfamily protein, putative | 2 |
| cat-miR171f | Ca6_33178001-33180911 | PREDICTED: scarecrow-like protein 6 | 2 |
| cat-miR171f | Ca8_8846184-8853412 | Nodulation receptor kinase | 2 |
| cat-miR171f | Ca3_36867207-36872512 | Ubiquitin-associated/TS-N domain protein | 2 |
| cat-miR171f | Ca1_38017772-38019712 | PREDICTED: cytochrome P450 CYP736A12-like | 2 |
| cat-miR171f | Ca7_5299299-5302601 | PREDICTED: uncharacterized protein LOC101503997 | 2 |
| cat-miR172a-3p | Ca8_3898204-3903405 | PREDICTED: CBS domain-containing protein CBSX1, chloroplastic | 2 |
| cat-miR172a-3p | Ca4_546779-563816 | CLIP-associated protein | 2 |
| cat-miR172a-3p | Ca4_7969482-8002552 | DUF3595 family protein (Fragment) | 2 |
| cat-miR172a-3p | Ca7_39277891-39279802 | PREDICTED: uncharacterized protein LOC101513554 | 2 |
| cat-miR172a-3p | Ca2_22044305-22048524 | RNA-binding (RRM/RBD/RNP motif) family protein | 2 |
| cat-miR172a-3p | Ca2_30988838-30993899 | PREDICTED: E3 ubiquitin-protein ligase RGLG2-like | 2 |
| cat-miR172a-3p | Ca3_48484897-48491490 | Glutamyl endopeptidase, putative | 2 |
| cat-miR172a-3p | Ca5_26717065-26723528 | ENTH/VHS/GAT family protein | 4 |
| cat-miR172a-3p | Ca4_860991-866920 | PREDICTED: tetratricopeptide repeat protein 7A | 4 |
| cat-miR172a-3p | Ca2_3398346-3401287 | Alpha-1,4-glucan-protein synthase [UDP-forming] | 4 |
| cat-miR172a-3p | Ca5_54194811-54199144 | PREDICTED: probable disease resistance protein At4g27220 | 4 |
| cat-miR172a-3p | Ca7_44835890-44849459 | Uncharacterized protein (Fragment) | 4 |
| cat-miR172a-5p | Ca7_55259767-55262607 | #N/A | 2 |
| cat-miR172a-5p | Scaffold0591_1314926-1319102 | #N/A | 2 |
| cat-miR172a-5p | Ca4_11533055-11536940 | PREDICTED: uncharacterized protein LOC101505330 | 2 |
| cat-miR172a-5p | Ca5_48732758-48737368 | #N/A | 2 |
| cat-miR172a-5p | Ca2_5116299-5118157 | #N/A | 3 |
| cat-miR172b | Ca5_24111827-24114120 | PREDICTED: serine/arginine-rich SC35-like splicing factor SCL30A isoform X2 | 2 |
| cat-miR172b | Ca1_19242992-19251306 | Subtilisin-like serine protease | 2 |
| cat-miR172b | Ca4_13468756-13481359 | Long chain acyl-CoA synthetase 7, peroxisomal protein | 3 |
| cat-miR172b | Ca3_30142083-30145590 | Uncharacterized protein | 3 |
| cat-miR172c-2 | Ca3_49158156-49169453 | Protein transporter Sec23-like protein | 2 |
| cat-miR2111-1 | Ca8_16873473-16884993 | PREDICTED: uncharacterized protein LOC101502546 isoform X1 | 2 |

| miR | Target | Target Annotation | Category |
|---|---|---|---|
| cat-miR2111-1 | Ca6_9906419-9914289 | Beta-adaptin-like protein | 2 |
| cat-miR2111-3 | Ca2_30769414-30776053 | Disease resistance protein (TIR-NBS-LRR class) | 2 |
| cat-miR2111-3 | Ca6_60920158-60924027 | PREDICTED: probableucleolar protein 5-1 | 2 |
| cat-miR2111-4 | Ca2_26940540-26945259 | Phosphatidylinositol-4-phosphate 5-kinase family protein | 2 |
| cat-miR2111-4 | Ca1_16524045-16525988 | Uncharacterized protein | 2 |
| cat-miR2111-4 | Ca7_53850511-53852814 | RAB GTPase-like protein A2D | 2 |
| cat-miR2111-5 | Ca4_124070-125393 | Lung seven transmembrane receptor family protein | 2 |
| cat-miR2111-5 | Ca7_55918746-55931978 | PREDICTED: galactinol--sucrose galactosyltransferase-like | 2 |
| cat-miR2111-5 | Ca6_15457474-15467723 | Peptidase M1 family aminopeptidase N | 3 |
| cat-miR2118 | Ca2_6884865-6893616 | Aminoacylase-1 | 2 |
| cat-miR2118 | Ca5_13031414-13036085 | PREDICTED: translocase of chloroplast 120, chloroplastic | 2 |
| cat-miR2118 | Ca6_14607759-14612073 | Protein kish | 2 |
| cat-miR319a | Ca7_54021813-54027912 | PREDICTED: potassium transporter 5 | 2 |
| cat-miR319a | Ca1_4115513-4408222 | PREDICTED: DNA-directed RNA polymerase I subunit RPA2-like | 4 |
| cat-miR319a | Ca1_4182016-4187149 | PREDICTED: la-related protein 6A | 4 |
| cat-miR319a | Ca4_3568757-3569348 | PREDICTED: 22.7 kDa class IV heat shock protein | 4 |
| cat-miR319b | Ca5_14983733-14987921 | PREDICTED: DNA-3-methyladenine glycosylase 1 | 0 |
| cat-miR319b | Scaffold5165_362219-376629 | Glycyl-tRNA synthetase alpha chain/beta chain | 1 |
| cat-miR319b | Ca1_13639768-13651895 | Protein DETOXIFICATION | 2 |
| cat-miR319b | Ca7_8080890-8082864 | PREDICTED: transcription factor MYC2-like | 2 |
| cat-miR319b | Ca3_63003816-63016414 | PREDICTED: mediator of RNA polymerase II transcription subunit 21 | 2 |
| cat-miR319b | Ca1_11611872-11616950 | Cell division cycle protein 48 homolog | 2 |
| cat-miR319b | Ca1_8470025-8476073 | Uncharacterized protein | 2 |
| cat-miR319b | Ca6_18805835-18811743 | CRAL/TRIO domain protein | 2 |
| cat-miR319b | Ca6_62917846-62919893 | PREDICTED: disease resistance protein RPS5-like isoform X1 | 2 |
| cat-miR319b | Ca2_19464837-19470581 | PREDICTED: putative disease resistance protein At3g14460 isoform X2 | 2 |
| cat-miR319b | Ca4_51501484-51510833 | L-galactono-1,4-lactone dehydrogenase | 2 |
| cat-miR319b | Ca5_18707298-18713407 | PREDICTED: DNA excision repair protein ERCC-8 | 2 |
| cat-miR319b | Ca4_27438016-27445232 | Ser/Thr protein kinase | 2 |
| cat-miR319d | Ca7_3592401-3593184 | PREDICTED: cold shock domain-containing protein 3 | 2 |
| cat-miR319d | Ca8_7856616-7859608 | Uncharacterized protein | 2 |
| cat-miR319d | Ca5_28788211-28793732 | PREDICTED: probable inactive receptor kinase At5g10020 | 2 |
| cat-miR319d | Ca8_2287662-2305919 | PREDICTED: chaperone protein dnaJ 13-like | 2 |
| cat-miR319d | Ca3_39148259-39153665 | PREDICTED: peptidyl-prolyl cis-trans isomerase CYP63 | 2 |
| cat-miR390a-1 | Ca4_18273370-18277558 | Pti1-like kinase | 2 |
| cat-miR390a-1 | Ca1_1022793-1034531 | PREDICTED: calmodulin-binding transcription activator 2-like isoform X1 | 4 |
| cat-miR390a-2 | Ca5_6598811-6601730 | Heat shock protein 90-2 | 3 |
| cat-miR390a-2 | Ca1_370423-371986 | PREDICTED: transcription factor TCP8 | 2 |
| cat-miR390a-2 | Ca1_2565745-2567233 | PREDICTED: auxin-repressed 12.5 kDa protein | 4 |
| cat-miR390b | Ca4_13988336-13991009 | Tubulin alpha-6 chain | 2 |
| cat-miR390b | Ca6_50406217-50412544 | PREDICTED: uncharacterized protein LOC101496135 | 2 |
| cat-miR390b | Ca4_11360318-11362309 | 0 | 2 |
| cat-miR390b | Ca1_14414398-14419065 | Vacuolar protein sorting-associated protein 4 | 2 |
| cat-miR390b | Ca5_24797040-24800711 | Uncharacterized protein | 3 |
| cat-miR394-1 | Ca7_45851918-45856242 | Uncharacterized protein | 2 |
| cat-miR394-1 | Ca4_18391220-18395966 | PREDICTED: extensin-2 | 2 |
| cat-miR394-1 | Ca4_18391220-18395966 | PREDICTED: extensin-2 | 2 |
| cat-miR394-1 | Ca4_18391305-18393071 | 0 | 2 |
| cat-miR394-1 | Ca6_2927678-2933079 | PREDICTED: polyadenylate-binding protein 2-like isoform X1 | 2 |
| cat-miR394-1 | Ca6_3090272-3109695 | PREDICTED: protein EXECUTER 1, chloroplastic | 2 |
| cat-miR394-1 | Ca5_64553658-64648791 | PREDICTED: ABC transporter C family member 12-like isoform X1 | 2 |
| cat-miR394-1 | Ca4_7710504-7714878 | PREDICTED: zinc finger CCCH domain-containing protein 32 isoform X1 | 2 |
| cat-miR394-1 | Ca4_19076720-19080079 | PREDICTED: protein IQ-DOMAIN 1 | 2 |
| cat-miR394-1 | Ca3_40235920-40238827 | Heat shock protein 70 | 2 |
| cat-miR394-1 | Ca1_39238900-39268403 | Elongation factor 2 | 2 |
| cat-miR394-1 | Ca6_5319822-5353332 | PREDICTED: isoflavone 4""-O-methyltransferase | 2 |

| miR | Target | Target Annotation | Category |
|---|---|---|---|
| cat-miR394-1 | Ca1_5487604-5490382 | 5'-adenylylsulfate reductase | 4 |
| cat-miR394-2 | Ca4_2036642-2049631 | PREDICTED: mitochondrial Rho GTPase 2-like | 1 |
| cat-miR394-2 | Ca4_18391305-18393071 | 0 | 2 |
| cat-miR394-2 | Ca4_18391305-18393071 | 0 | 2 |
| cat-miR394-2 | Ca4_18391220-18395966 | PREDICTED: extensin-2 | 2 |
| cat-miR394-2 | Ca6_15050184-15052289 | PREDICTED: loricrin isoform X2 | 2 |
| cat-miR394-2 | Ca1_28133226-28138762 | PREDICTED: plant cysteine oxidase 3 | 2 |
| cat-miR394-2 | Ca7_50532288-50533892 | PREDICTED: blue copper protein-like | 2 |
| cat-miR394-2 | Ca4_55458026-55462700 | TCP-1/cpn60 chaperonin family protein | 4 |
| cat-miR396b | Ca4_50439612-50444040 | PREDICTED: uncharacterized protein LOC101494882 | 2 |
| cat-miR396b | Ca5_12839788-12843991 | PREDICTED: MLO-like protein 1 | 2 |
| cat-miR396b | Ca1_12488795-12493660 | RhoGAP domain-containing protein | 2 |
| cat-miR398a-3p | Ca6_13854616-13859598 | #N/A | 1 |
| cat-miR398a-3p | Ca7_54676569-54680715 | PREDICTED: copper chaperone for superoxide dismutase, chloroplastic/cytosolic | 2 |
| cat-miR398a-3p | Ca1_47822038-47834873 | PREDICTED: BES1/BZR1 homolog protein 4-like isoform X2 | 2 |
| cat-miR398a-3p | Ca3_49574045-49616098 | PREDICTED: ATP-dependent 6-phosphofructokinase 6-like | 2 |
| cat-miR398a-3p | Ca5_12802655-12809975 | PREDICTED: chromo domain protein LHP1-like | 2 |
| cat-miR398a-3p | Ca5_27876977-27881480 | #N/A | 2 |
| cat-miR398a-3p | Ca8_134400-141438 | Presequence protease | 2 |
| cat-miR398a-3p | Ca4_724504-727359 | S-(hydroxymethyl)glutathione dehydrogenase | 2 |
| cat-miR398a-3p | Ca7_49896355-49909071 | #N/A | 2 |
| cat-miR398a-3p | Ca4_10604838-10635751 | PREDICTED: putative 1-phosphatidylinositol-3-phosphate 5-kinase FAB1D isoform X1 | 4 |
| cat-miR398b-1 | Ca7_54935055-54940470 | PREDICTED: FAM10 family protein At4g22670 | 2 |
| cat-miR398b-2 | Ca1_46430145-46432252 | PREDICTED: cucumber peeling cupredoxin-like | 3 |
| cat-miR398b-2 | Ca6_47621471-47626541 | PREDICTED: cytochrome c oxidase subunit 5b-2, mitochondrial-like | 2 |
| cat-miR398b-2 | Ca6_46206753-46208202 | PREDICTED: uncharacterized protein LOC101502166 | 2 |
| cat-miR399-1 | Ca7_3901069-3905879 | Serine/threonine phosphatase family, 2C domain protein | 2 |
| cat-miR408 | Ca6_62198400-62199252 | Basic blue copper protein | 4 |
| cat-miR408 | Ca8_14689512-14691283 | 0 | 3 |
| cat-miR408 | Ca7_53661305-53664808 | PREDICTED: transcription factor MYB1R1 | 2 |
| cat-miR408 | Ca7_36431558-36445178 | PREDICTED: MATE efflux family protein FRD3-like | 2 |
| cat-miR408 | Ca7_1013376-1016393 | PREDICTED: probable inactive receptor kinase At5g67200 | 2 |
| cat-miR408 | Ca5_21577685-21598302 | PREDICTED: protein MODIFIER OF SNC1 1-like isoform X1 | 2 |
| cat-miR408 | Ca7_51849608-51852123 | PREDICTED: dnaJ homolog subfamily C member 21 | 2 |
| cat-miR408 | Ca4_6535078-6538929 | PREDICTED: pectin acetylesterase 6-like | 2 |
| cat-miR408 | Ca5_20132599-20133510 | PREDICTED: blue copper protein-like | 4 |
| cat-miR482 | Ca3_46000144-46006977 | Putative FAD synthase | 2 |
| cat-miR482 | Ca5_13031414-13036085 | PREDICTED: translocase of chloroplast 120, chloroplastic | 2 |
| cat-miR482 | Ca5_8502860-8503460 | Clathrin heavy chain | 2 |
| cat-miR482 | Ca3_62098022-62101425 | Kinase 1B | 2 |
| cat-miR482 | Ca5_38041345-38041591 | 0 | 2 |
| cat-miR5213 | Ca4_15336820-15340954 | PREDICTED: uncharacterized protein LOC101515007 | 2 |
| cat-miR5213 | Ca6_46206753-46208202 | PREDICTED: uncharacterized protein LOC101502166 | 2 |
| cat-miR5213 | Ca2_30769414-30776053 | Disease resistance protein (TIR-NBS-LRR class) | 4 |
| cat-NovmiR1 | Scaffold4730_239146-242260 | PREDICTED: annexin D5-like | 3 |
| cat-NovmiR1 | Ca3_40611229-40613412 | P-loopucleoside triphosphate hydrolase superfamily protein | 2 |
| cat-NovmiR1 | Ca6_65693359-65706527 | Uncharacterized protein | 2 |
| cat-NovmiR1 | Ca5_33770931-33795513 | Uncharacterized protein | 2 |
| cat-NovmiR1 | Ca1_493585-501429 | Importin beta-4 | 2 |
| cat-NovmiR1 | Ca1_13531449-13535357 | Uncharacterized protein | 2 |
| cat-NovmiR1 | Ca3_32369219-32372317 | PREDICTED: CBS domain-containing protein CBSX6 | 2 |
| cat-NovmiR1 | Ca7_52636757-52641226 | Hipl2 protein | 2 |
| cat-NovmiR1 | Ca8_8648977-8654820 | PREDICTED: protein SPT2 homolog | 2 |
| cat-NovmiR1 | Ca4_35821825-35825849 | Nucleotide/sugar transporter family protein | 2 |
| cat-NovmiR1 | Ca7_32904863-32909252 | NBS-LRR protein | 4 |
| cat-NovmiR1 | Ca4_18878309-18881682 | Adenylate kinase | 4 |
| cat-NovmiR1 | Ca6_32935683-32940967 | Ubiquinone biosynthesis protein UbiB | 4 |
| cat-miR398a-5p | Ca4_28039768-28047455 | PREDICTED: uncharacterized protein LOC101509839 | 2 |

| miR | Target | Target Annotation | Category |
|---|---|---|---|
| cat-NovmiR2 | Ca7_49490390-49494954 | Amine oxidase | 2 |
| cat-NovmiR2 | Ca3_61410219-61413086 | PREDICTED: desumoylating isopeptidase 1 | 2 |
| cat-NovmiR2 | Ca5_3768744-3777872 | Serine/threonine-protein phosphatase 5 | 2 |
| cat-NovmiR2 | Ca7_52218004-52226931 | Geranylgeranyl hydrogenase | 2 |
| cat-NovmiR2 | Ca4_10430997-10446943 | Exocyst complex component sec5 | 2 |
| cat-NovmiR4 | Ca1_18680660-18684596 | PREDICTED: metal tolerance protein 1-like | 2 |
| cat-NovmiR6 | Ca7_46620916-46627159 | PREDICTED: protein trichome birefringence-like 5 | 4 |
| cat-NovmiR6 | Ca3_47272688-47277573 | PREDICTED: protein YLS9 | 3 |
| cat-NovmiR6 | Ca4_15962867-15967945 | PREDICTED: dihydrolipoyllysine-residue acetyltransferase component 4 of pyruvate dehydrogenase complex, chloroplastic | 2 |
| cat-NovmiR6 | Ca1_4797478-4808344 | PREDICTED: F-box/kelch-repeat protein At3g23880-like | 2 |
| cat-NovmiR6 | Ca6_3567998-3572811 | PREDICTED: uncharacterized protein LOC101502776 isoform X1 | 2 |
| cat-NovmiR6 | Ca6_47569511-47574675 | PREDICTED: uncharacterized protein LOC101501411 | 2 |
| cat-NovmiR6 | Ca8_5836454-5845794 | Vacuolar protein sorting-associated protein 35 | 2 |
| cat-NovmiR6 | Ca8_12830062-12831541 | PREDICTED: heat stress transcription factor C-1 | 2 |
| cat-NovmiR7 | Ca3_31814617-31817847 | PREDICTED: O-acyltransferase WSD1-like | 2 |
| cat-NovmiR7 | Ca6_11818450-11821061 | #N/A | 2 |
| cat-NovmiR7 | Ca3_46141134-46146044 | #N/A | 2 |
| cat-NovmiR7 | Ca4_57258537-57264573 | #N/A | 2 |
| cat-NovmiR7 | Ca7_3009749-3012053 | #N/A | 2 |
| cat-miR394 | Ca1_35237403-35238547 | Histidine phosphotransferase | 3 |

**Mid-Late nodule**

| miR | Target | Target Annotation | Category |
|---|---|---|---|
| Cat-miR156b | Ca3:38170417-38173423 | PREDICTED: squamosa promoter-binding-like protein 9 | 0 |
| Cat-miR319b | Ca1:3936260-3937645 | PREDICTED: transcription factor TCP2 | 0 |
| Cat-miR1507 | Ca4:36280496-36289841 | #N/A | 2 |
| Cat-miR1507 | Ca5:9916501-9923271 | #N/A | 2 |
| Cat-miR1507 | Ca6:60399878-60402185 | #N/A | 2 |
| Cat-miR1507 | Ca7:45954575-45961528 | Calmodulin-binding protein | 2 |
| Cat-miR1507 | Ca4:16705196-16718676 | #N/A | 2 |
| Cat-miR1507 | Ca6:47027344-47030772 | #N/A | 2 |
| Cat-miR1507 | Ca2:3102138-3115063 | PREDICTED: serine/threonine-protein kinase dst1 | 2 |
| Cat-miR1507 | Ca6:37118141-37119983 | #N/A | 2 |
| Cat-miR1507 | Scaffold0583:8101-11346 | #N/A | 4 |
| Cat-miR1509 | Ca4:848278-850064 | PREDICTED: protein JASON | 0 |
| Cat-miR1509 | Ca4:4297172-4301616 | PREDICTED: U11/U12 smalluclear ribonucleoprotein 48 kDa protein | 2 |
| Cat-miR1509 | Ca3:50200282-50202136 | PREDICTED: uncharacterized protein LOC101492518 | 2 |
| Cat-miR1509 | Ca6:885910-888498 | PREDICTED: adenylate kinase isoenzyme 6 homolog | 2 |
| Cat-miR1509 | Ca1:12076587-12079921 | PREDICTED: adenylate kinase isoenzyme 6 homolog | 2 |
| Cat-miR1511 | Ca4:37659490-37665064 | PREDICTED: pentatricopeptide repeat-containing protein At3g22470, mitochondrial-like | 4 |
| Cat-miR156a-1 | Ca6:49578549-49588910 | PREDICTED: protein ROS1-like | 2 |
| Cat-miR156a-2 | Ca4:28830182-28838574 | PREDICTED: protein EMSY-LIKE 1 | 2 |
| Cat-miR156a-2 | Ca2:17579241-17596608 | PREDICTED: probable leucine-rich repeat receptor-like serine/threonine-protein kinase At3g14840 | 2 |
| Cat-miR156a-2 | Ca3:20743615-20748677 | PREDICTED: transcription initiation factor TFIID subunit 11 isoform X1 | 2 |
| Cat-miR156a-2 | Ca3:32941385-32944363 | Acyl-CoA thioesterase, putative | 4 |
| Cat-miR156a-2 | Ca7:650930-653206 | PREDICTED: subtilisin-like protease SBT1.7 | 3 |
| Cat-miR156a-3 | Ca7:47840062-47910402 | Calcium-dependent lipid-binding-like protein | 2 |
| Cat-miR156a-3 | Ca8:10742166-10752979 | AGC family Serine/Threonine kinase family protein | 2 |
| Cat-miR156a-3 | Ca4:7628104-7654987 | PREDICTED: uncharacterized protein LOC101497938 | 2 |
| Cat-miR156a-3 | Ca6:14931925-14954984 | PREDICTED: centromere-associated protein E isoform X4 | 2 |
| Cat-miR156a-3 | Ca4:14419435-14421848 | PREDICTED: methylthioribose kinase-like isoform X1 | 2 |
| Cat-miR156a-3 | Ca2:3794979-3800049 | PREDICTED: LOW QUALITY PROTEIN: squamosa promoter-binding-like protein 6 | 4 |
| Cat-miR156a-3 | Ca3:40920822-40927671 | PREDICTED: ethylene-insensitive protein 2 | 4 |
| Cat-miR156b | Ca7:45249967-45254466 | Cytosol aminopeptidase family protein | 2 |
| Cat-miR156c-1 | Ca3:32989732-32993382 | Uncharacterized protein | 2 |
| Cat-miR156c-1 | Ca1:3241226-3245890 | Uncharacterized protein | 4 |
| Cat-miR156c-4 | Ca6:20650287-20653460 | PREDICTED: ATPase ASNA1 homolog | 4 |
| Cat-miR156d | Ca8:704873-710512 | PREDICTED: putative 12-oxophytodienoate reductase 11 | 2 |

| miR | Target | Target Annotation | Category |
|---|---|---|---|
| Cat-miR156d | Ca8:12188166-12275518 | Plant synaptotagmin | 2 |
| Cat-miR156d | Ca6:5704789-5709534 | PREDICTED: protein IQ-DOMAIN 14 | 2 |
| Cat-miR156d | Ca4:15483996-15488401 | PREDICTED: cysteine--tRNA ligase, cytoplasmic | 2 |
| Cat-miR156d | Ca6:1687903-1697332 | PREDICTED: LOW QUALITY PROTEIN: E3 ubiquitin-protein ligase UPL1-like | 2 |
| Cat-miR156d | Ca7:50193177-50199734 | Probable sucrose-phosphate synthase | 2 |
| Cat-miR156d | Ca7:31644338-31648917 | Endoplasmic reticulum-type calcium-transporting ATPase | 2 |
| Cat-miR156d | Ca3:58041312-58055198 | Actin-97 | 2 |
| Cat-miR156d | Ca3:58041312-58055198 | Actin-97 | 2 |
| Cat-miR156d | Ca4:42250207-42251142 | PREDICTED: uncharacterized protein LOC101509491 | 2 |
| Cat-miR156d | Ca6:4306107-4308806 | PREDICTED: armadillo repeat-containing protein 7 | 2 |
| Cat-miR156d | Ca1:3908973-3925137 | Uncharacterized protein | 2 |
| Cat-miR156d | Ca6:20650287-20653460 | PREDICTED: ATPase ASNA1 homolog | 4 |
| Cat-miR156d | Ca1:9464245-9467765 | PREDICTED: uncharacterized protein LOC101506888 | 4 |
| Cat-miR156d | Ca4:3948180-3958105 | Serine/Threonine kinase family protein | 4 |
| Cat-miR156e-1 | Ca7:51171572-51188309 | #N/A | 2 |
| Cat-miR156e-1 | Ca1:11776044-11780075 | #N/A | 2 |
| Cat-miR156e-1 | Ca5:20201399-20203182 | #N/A | 2 |
| Cat-miR156e-1 | Ca5:18544804-18546508 | #N/A | 2 |
| Cat-miR156e-1 | Ca5:20823052-20831241 | PREDICTED: telomere length regulation protein TEL2 homolog | 2 |
| Cat-miR156e-1 | Ca4:4231487-4243578 | #N/A | 2 |
| Cat-miR156e-2 | Ca8:4552809-4554903 | #N/A | 2 |
| Cat-miR156e-2 | Ca7:40816965-40822688 | #N/A | 2 |
| Cat-miR156e-2 | Ca4:39272539-39278624 | #N/A | 2 |
| Cat-miR156e-2 | Ca2:19636677-19639510 | #N/A | 2 |
| Cat-miR156e-2 | Ca6:29033827-29037684 | #N/A | 2 |
| Cat-miR156e-2 | Ca5:17944778-17949876 | Eukaryotic translation initiation factor 3 subunit B | 2 |
| Cat-miR156e-2 | Ca6:1283052-1285929 | #N/A | 2 |
| Cat-miR156f-1 | Ca1:27335753-27339610 | #N/A | 4 |
| Cat-miR156f-2 | Ca7:51171572-51188309 | #N/A | 2 |
| Cat-miR156f-2 | Ca5:18544804-18546508 | #N/A | 2 |
| Cat-miR156f-2 | Ca1:12701923-12713471 | #N/A | 2 |
| Cat-miR156f-2 | Ca5:20201399-20203182 | #N/A | 3 |
| Cat-miR160-1 | Ca2:20871772-20878030 | 2-oxoacid dehydrogenase acyltransferase family protein | 2 |
| Cat-miR162b | Scaffold0592:921963-925549 | Uncharacterized protein | 3 |
| Cat-miR162b | Ca5:27769383-27784627 | PREDICTED: MAG2-interacting protein 2 | 2 |
| Cat-miR162b | Ca6:6805928-6807310 | PREDICTED: serine carboxypeptidase-like 50 | 2 |
| Cat-miR162b | Ca4:9020986-9025907 | PREDICTED: uncharacterized protein LOC101488857 | 2 |
| Cat-miR162b | Ca2:29208614-29238954 | Uncharacterized protein | 3 |
| Cat-miR162b | Ca7:34044020-34054244 | PREDICTED: vacuolar protein sorting-associated protein 8 homolog | 4 |
| Cat-miR162b | Ca8:6258849-6260126 | PREDICTED: phosphoserine aminotransferase 2, chloroplastic | 4 |
| Cat-miR164a-1 | Ca7:55487062-55491962 | PREDICTED: uncharacterized protein LOC101493304 | 2 |
| Cat-miR164a-2 | Ca1:39628190-39632859 | NAC transcription factor | 4 |
| Cat-miR164a-2 | Ca7:51381379-51385717 | PREDICTED: protein Dr1 homolog isoform X1 | 2 |
| Cat-miR164a-3 | Scaffold0628:550854-554827 | Eukaryotic peptide chain release factor subunit 1-3 | 2 |
| Cat-miR164a-3 | Ca6:12063827-12067769 | Uncharacterized protein | 2 |
| Cat-miR164b | Ca4:36083975-36091304 | PREDICTED: uncharacterized protein LOC101491241 isoform X1 | 0 |
| Cat-miR164b | Ca4:17963476-17966111 | Serine/threonine phosphatase family, 2C domain protein | 2 |
| Cat-miR164b | Ca8:2571562-2577096 | PREDICTED: GATA transcription factor 26-like | 2 |
| Cat-miR164b | Ca6:63962722-63967929 | PREDICTED: pentatricopeptide repeat-containing protein At5g27270 isoform X1 | 2 |
| Cat-miR164b | Ca6:39786201-39790820 | Splicing factor-like protein | 2 |
| Cat-miR164b | Ca4:12071293-12077676 | PREDICTED: titin-like | 2 |
| Cat-miR164b | Ca5:65215171-65217442 | PREDICTED: NAD(P)H-dependent 6""-deoxychalcone synthase | 2 |
| Cat-miR164b | Ca7:50532289-50533892 | PREDICTED: blue copper protein-like | 2 |
| Cat-miR164b | Ca7:50113542-50115041 | PREDICTED:ucleolin-like | 2 |
| Cat-miR164b | Ca6:56585310-56590849 | PREDICTED: zinc finger CCCH domain-containing protein 13-like isoform X1 | 2 |
| Cat-miR164b | Ca2:9501287-9509470 | Glucose-6-phosphate isomerase | 2 |
| Cat-miR164b | Ca4:15973580-15984001 | PREDICTED: serine/threonine-protein kinase prpf4B | 2 |
| Cat-miR164b | Ca3:39922687-39930067 | ABC transporter-like family-protein | 2 |

| miR | Target | Target Annotation | Category |
|---|---|---|---|
| Cat-miR164b | Ca2:1466470-1475501 | PREDICTED: uncharacterized protein LOC101507145 isoform X1 | 2 |
| Cat-miR164b | Ca2:32259138-32321077 | PREDICTED: isoprenylcysteine alpha-carbonyl methylesterase ICME | 2 |
| Cat-miR164b | Ca4:18391221-18395966 | PREDICTED: extensin-2 | 2 |
| Cat-miR164b | Ca4:18391306-18393071 | 0 | 2 |
| Cat-miR164b | Ca6:2731521-2734216 | PREDICTED: uncharacterized protein LOC101491587 | 2 |
| Cat-miR164b | Ca4:9426488-9432327 | Peptidyl-prolyl cis-trans isomerase | 2 |
| Cat-miR164b | Ca2:13485828-13493999 | Translocon at the inner envelope membrane ofs 110 protein | 2 |
| Cat-miR164b | Ca4:57945566-57949624 | PREDICTED: transcription factor bHLH140 | 2 |
| Cat-miR164b | Ca4:11360319-11362309 | 0 | 2 |
| Cat-miR164b | Ca4:21055545-21059252 | Aspartate/glutamate/uridylate kinase family protein, putative | 2 |
| Cat-miR164b | Ca1:6530617-6539298 | PREDICTED: enhancer of mRNA-decapping protein 4-like isoform X1 | 2 |
| Cat-miR164b | Ca5:67434414-67441696 | Protein DETOXIFICATION | 2 |
| Cat-miR164b | Ca6:60737772-60743432 | PREDICTED: putative rRNA methyltransferase | 2 |
| Cat-miR164b | Ca6:9677683-9681003 | PREDICTED: protein TRAUCO | 2 |
| Cat-miR164b | Ca3:58667251-58671227 | TPR protein | 2 |
| Cat-miR164b | Ca1:10230563-10235575 | Casein kinase I-like protein | 2 |
| Cat-miR164b | Ca6:52242301-52247990 | PREDICTED: external alternative NAD(P)H-ubiquinone oxidoreductase B1, mitochondrial | 2 |
| Cat-miR164b | Scaffold0590:52294-60863 | Beta-amyrin synthase | 2 |
| Cat-miR164b | Ca3:41806985-41809898 | PREDICTED: S-adenosylmethionine decarboxylase proenzyme-like isoform X3 | 2 |
| Cat-miR164b | Ca8:10992136-10996143 | Endosomal targeting BRO1-like domain protein | 2 |
| Cat-miR164b | Ca6:13348470-13351975 | PREDICTED: putative N6-adenosine-methyltransferase MT-A70-like | 2 |
| Cat-miR164b | Ca7:53536623-53541682 | GDP-fucose O-fucosyltransferase-like protein | 2 |
| Cat-miR164b | Ca7:38756261-38764073 | PREDICTED: 11-oxo-beta-amyrin 30-oxidase-like | 2 |
| Cat-miR164b | Ca5:13861236-14094850 | Niemann-Pick C1 protein | 2 |
| Cat-miR164b | Ca5:15279345-15283738 | PREDICTED: protein odr-4 homolog | 2 |
| Cat-miR164b | Ca6:48150861-48156205 | PREDICTED: uncharacterized protein LOC101504845 | 2 |
| Cat-miR164b | Ca6:2873375-2888428 | PREDICTED: LOW QUALITY PROTEIN: outer envelope protein 64, mitochondrial | 2 |
| Cat-miR164b | Ca7:5955565-5957856 | PREDICTED: histone-lysine N-methyltransferase family member SUVH9-like | 2 |
| Cat-miR164b | Ca4:16113172-16120762 | PREDICTED: putative transcription elongation factor SPT5 homolog 1 | 2 |
| Cat-miR164b | Ca1:17810655-17814971 | Uncharacterized protein | 2 |
| Cat-miR164b | Ca7:21948766-21949374 | PREDICTED: dof zinc finger protein DOF1.7-like | 2 |
| Cat-miR164b | Ca4:5699941-5701446 | Ribosomal L22e family protein | 2 |
| Cat-miR164b | Ca4:40696173-40697926 | PREDICTED: tankyrase | 2 |
| Cat-miR164b | Ca7:2500248-2505805 | U1 smalluclear ribonucleoprotein 70 kDa protein, putative | 2 |
| Cat-miR164b | Ca3:33137425-33264074 | PREDICTED: subtilisin-like protease Glyma18g48580 | 2 |
| Cat-miR164b | Ca6:65079203-65085741 | NADH-ubiquinone oxidoreductase 75 kDa subunit | 2 |
| Cat-miR164b | Ca1:14649450-14663687 | PREDICTED: protein TIC 100 | 4 |
| Cat-miR166a-1 | Ca5:29055857-29059960 | PREDICTED: uncharacterized protein LOC101498857 isoform X1 | 2 |
| Cat-miR166a-5 | Ca2:10846941-10850162 | PREDICTED: auxin-responsive protein IAA8-like | 2 |
| Cat-miR166a-5 | Ca4:38849644-38852342 | PREDICTED: glucan endo-1,3-beta-glucosidase 5 | 2 |
| Cat-miR166a-5 | Ca5:13861236-14094850 | Niemann-Pick C1 protein | 2 |
| Cat-miR166a-5 | Ca8:7351797-7363853 | PREDICTED: probable glucan 1,3-alpha-glucosidase isoform X1 | 2 |
| Cat-miR166b-1 | Ca7:31017196-31020568 | Uncharacterized protein | 2 |
| Cat-miR166b-1 | Ca2:30205340-30209864 | PREDICTED: KH domain-containing protein At4g18375 isoform X2 | 2 |
| Cat-miR166b-1 | Ca3:45664655-45668033 | General substrate transporter | 2 |
| Cat-miR166b-2 | Ca5:27571201-27578359 | PP2A regulatory subunit TAP46-like protein | 2 |
| Cat-miR166b-2 | Ca1:8841298-8852892 | Uncharacterized protein | 2 |
| Cat-miR166b-3 | Ca7:54043773-54060442 | PREDICTED: pentatricopeptide repeat-containing protein At4g17616 | 2 |
| Cat-miR167a-1 | Ca1:6806903-6814467 | PREDICTED: uncharacterized protein LOC101513958 isoform X1 | 2 |
| Cat-miR167a-2 | Ca6:60959806-60962198 | PREDICTED: probable cysteine proteinase A494 | 4 |

| miR | Target | Target Annotation | Category |
|---|---|---|---|
| Cat-miR167b-1 | Ca6:63308084-63310409 | Tubulin alpha-5 | 2 |
| Cat-miR167b-1 | Ca8:9669877-9675673 | T-complex protein 1 subunit gamma | 2 |
| Cat-miR167b-1 | Ca5:19987741-20004113 | PREDICTED: E3 ubiquitin-protein ligase UPL1-like isoform X1 | 4 |
| Cat-miR167b-2 | Ca3:39563116-39567033 | PREDICTED: target of Myb protein 1 | 2 |
| Cat-miR167b-2 | Ca2:10364325-10370128 | PREDICTED: E3 ubiquitin-protein ligase UPL1-like | 4 |
| Cat-miR167c | Ca5:23552052-23560338 | PREDICTED: uncharacterized protein LOC101498628 | 2 |
| Cat-miR167c | Ca4:10064502-10067509 | PREDICTED: serine carboxypeptidase-like 27 | 2 |
| Cat-miR167c | Ca1:8576418-8591212 | PREDICTED: ABC transporter C family member 13 isoform X1 | 2 |
| Cat-miR167c | Ca4:1184794-1187058 | PREDICTED: uncharacterized protein LOC101503717 | 2 |
| Cat-miR167c | Ca3:35838617-35848681 | #N/A | 2 |
| Cat-miR167c | Scaffold0598:59111-59922 | #N/A | 2 |
| Cat-miR167c | Ca4:15973580-15984001 | PREDICTED: serine/threonine-protein kinase prpf4B | 2 |
| Cat-miR167c | Ca4:2210922-2223908 | #N/A | 2 |
| Cat-miR167c | Ca5:7515538-7519470 | #N/A | 2 |
| Cat-miR167c | Ca6:4545902-4555815 | #N/A | 2 |
| Cat-miR167c | Ca7:49718851-49725133 | #N/A | 2 |
| Cat-miR167d | Ca6:530903-536562 | Auxin response factor | 2 |
| Cat-miR167d | Ca6:2632042-2638142 | Dynamin-related protein 5B-2 | 2 |
| Cat-miR167d | Ca1:6971933-6975337 | PREDICTED: uncharacterized CRM domain-containing protein At3g25440, chloroplastic | 2 |
| Cat-miR167d | Ca8:13808270-13812637 | Uncharacterized protein | 2 |
| Cat-miR167d | Ca6:11538081-11541016 | PREDICTED: probable WRKY transcription factor 4 isoform X1 | 2 |
| Cat-miR167d | Scaffold0591:353979-356460 | PREDICTED: LOB domain-containing protein 41 | 2 |
| Cat-miR167d | Ca1:273124-277844 | PREDICTED: eukaryotic translation initiation factor 3 subunit C | 2 |
| Cat-miR167d | Ca7:50460763-50470321 | Topless-like protein | 2 |
| Cat-miR167d | Ca4:55408065-55412024 | #N/A | 2 |
| Cat-miR167d | Ca7:49182431-49189768 | Acyl-CoA-binding domain protein | 4 |
| Cat-miR168 | Scaffold0628:491140-497352 | PREDICTED: protein argonaute 1 | 3 |
| Cat-miR168 | Ca4:14251981-14254921 | #N/A | 2 |
| Cat-miR168 | Ca7:5926352-5929814 | #N/A | 2 |
| Cat-miR171a | Ca3:58041312-58055198 | Actin-97 | 3 |
| Cat-miR171a | Ca3:58041312-58055198 | Actin-97 | 3 |
| Cat-miR171a | Scaffold0592:786495-787619 | PREDICTED: F-box/kelch-repeat protein At1g23390 | 2 |
| Cat-miR171a | Ca7:47326243-47331530 | Sucrose synthase | 2 |
| Cat-miR171a | Ca1:4577001-4583210 | PREDICTED: squamosa promoter-binding-like protein 7 | 2 |
| Cat-miR171a | Ca7:41378627-41381714 | PREDICTED: MLP-like protein 34 | 2 |
| Cat-miR171a | Ca4:40048975-40055520 | PREDICTED: transcription initiation factor TFIID subunit 12 | 2 |
| Cat-miR171a | Ca7:50532289-50533892 | PREDICTED: blue copper protein-like | 2 |
| Cat-miR171a | Ca7:28588962-28595371 | PREDICTED: probable LRR receptor-like serine/threonine-protein kinase At1g56140 | 2 |
| Cat-miR171a | Ca3:49880383-49888862 | Protein YIPF | 2 |
| Cat-miR171a | Scaffold0539:952-1631 | PREDICTED: protein YIPF1 homolog | 2 |
| Cat-miR171b | Ca1:11929779-11933751 | PREDICTED: protein disulfide isomerase-like 1-4 | 2 |
| Cat-miR171b | Scaffold0592:98690-99652 | Putative mitochondrial dicarboxylate carrier protein | 2 |
| Cat-miR171b | Ca1:8874996-8876253 | PREDICTED: protein LURP-one-related 5-like | 2 |
| Cat-miR171b | Ca5:13250527-13262167 | Kinesin-like protein | 2 |
| Cat-miR171b | Ca6:46546994-46558783 | PREDICTED: malonate--CoA ligase-like | 2 |
| Cat-miR171b | Ca7:21740041-21744452 | Myo-inositol oxygenase | 4 |
| Cat-miR171c | Ca4:56157638-56159879 | PREDICTED: F-box protein At1g47056-like | 0 |
| Cat-miR171c | Ca2:19820125-19824906 | PREDICTED: DEAD-box ATP-dependent RNA helicase 53-like | 3 |
| Cat-miR171c | Ca4:8311024-8318521 | Uncharacterized protein | 2 |
| Cat-miR171c | Ca4:55615011-55616659 | PREDICTED: U-box domain-containing protein 30-like | 2 |
| Cat-miR171c | Ca7:46860844-46873793 | PREDICTED: uncharacterized protein LOC101507127 | 2 |
| Cat-miR171c | Ca2:21771032-21774610 | PREDICTED: cytochrome c oxidase assembly protein COX15 | 2 |
| Cat-miR171c | Ca2:19048160-19050842 | PREDICTED: UBP1-associated protein 2B-like | 2 |
| Cat-miR171d | Ca4:28563719-28566424 | PREDICTED: uncharacterized protein LOC101514582 | 2 |
| Cat-miR171d | Ca7:9465648-9470859 | Putative uncharacterized protein (Fragment) | 2 |
| Cat-miR171d | Ca2:16538768-16542277 | Uncharacterized protein | 2 |
| Cat-miR171d | Ca2:22157859-22160693 | Emp24/gp25L/p24 family protein | 2 |

| miR | Target | Target Annotation | Category |
|---|---|---|---|
| Cat-miR171e | Ca5:29055857-29059960 | PREDICTED: uncharacterized protein LOC101498857 isoform X1 | 0 |
| Cat-miR171e | Ca5:9952939-9963653 | Glutathione S-transferase/chloride channel, C-terminal | 2 |
| Cat-miR171e | Ca5:9952939-9963653 | Glutathione S-transferase/chloride channel, C-terminal | 2 |
| Cat-miR171e | Ca6:62350912-62354290 | PREDICTED: magnesium transporter MRS2-4 | 2 |
| Cat-miR171e | Ca6:23768275-23771391 | Putative uncharacterized protein | 2 |
| Cat-miR171e | Ca5:12422256-12435106 | P-loopucleoside triphosphate hydrolase superfamily protein, putative | 2 |
| Cat-miR171f | Ca6:15588397-15595765 | PREDICTED: kinesin KP1 | 2 |
| Cat-miR171f | Ca8:8846185-8853412 | Nodulation receptor kinase | 2 |
| Cat-miR171f | Ca3:36867208-36872512 | Ubiquitin-associated/TS-N domain protein | 2 |
| Cat-miR172a-3p | Ca6:933553-936728 | PREDICTED: myosin-binding protein 3 | 2 |
| Cat-miR172a-3p | Ca8:3898205-3903405 | PREDICTED: CBS domain-containing protein CBSX1, chloroplastic | 2 |
| Cat-miR172a-3p | Ca4:5034002-5059905 | Armadillo/beta-catenin-like repeat protein | 2 |
| Cat-miR172a-3p | Ca4:546780-563816 | CLIP-associated protein | 2 |
| Cat-miR172a-3p | Ca4:7969483-8002552 | DUF3595 family protein (Fragment) | 2 |
| Cat-miR172a-3p | Scaffold0591:750710-753404 | PREDICTED: uncharacterized protein LOC101504964 | 2 |
| Cat-miR172a-3p | Ca4:38186428-38194286 | PREDICTED: basic leucine zipper 9 | 2 |
| Cat-miR172a-3p | Ca2:22044306-22048524 | RNA-binding (RRM/RBD/RNP motif) family protein | 2 |
| Cat-miR172a-3p | Ca2:30988839-30993899 | PREDICTED: E3 ubiquitin-protein ligase RGLG2-like | 2 |
| Cat-miR172a-3p | Ca2:3398347-3401287 | Alpha-1,4-glucan-protein synthase [UDP-forming] | 2 |
| Cat-miR172a-3p | Ca5:5749833-5752712 | Uncharacterized protein | 2 |
| Cat-miR172a-3p | Ca5:8883186-8891249 | PREDICTED: ubiquitin-activating enzyme E1 1-like | 2 |
| Cat-miR172a-3p | Ca6:1707938-1712655 | PREDICTED: 6,7-dimethyl-8-ribityllumazine synthase, chloroplastic | 2 |
| Cat-miR172a-3p | Ca5:48738321-48742258 | SEC14 cytosolic factor-like protein | 2 |
| Cat-miR172a-3p | Ca1:1750428-1752891 | PREDICTED: protein PLANT CADMIUM RESISTANCE 8-like | 2 |
| Cat-miR172a-3p | Ca3:8527238-8541326 | PREDICTED: chromatin modification-related protein EAF1 B-like isoform X1 | 2 |
| Cat-miR172a-3p | Ca5:26717066-26723528 | ENTH/VHS/GAT family protein | 4 |
| Cat-miR172a-3p | Ca4:860992-866920 | PREDICTED: tetratricopeptide repeat protein 7A | 4 |
| Cat-miR172a-3p | Ca6:12220046-12224613 | 1-aminocyclopropane-1-carboxylate deaminase, putative | 4 |
| Cat-miR172a-3p | Ca7:41038401-41040818 | PREDICTED: cyclic dof factor 1-like | 4 |
| Cat-miR172a-5p | Ca1:2726067-2729458 | #N/A | 2 |
| Cat-miR172a-5p | Ca5:24235486-24236588 | #N/A | 2 |
| Cat-miR172a-5p | Ca2:33138434-33139531 | PREDICTED: proline-rich protein 4 | 2 |
| Cat-miR172a-5p | Scaffold0591:1314927-1319102 | #N/A | 2 |
| Cat-miR172a-5p | Ca1:12215227-12219389 | #N/A | 4 |
| Cat-miR172b | Ca7:46399050-46403793 | Subtilisin-like serine protease | 2 |
| Cat-miR172b | Ca1:19242993-19251306 | Subtilisin-like serine protease | 2 |
| Cat-miR172b | Ca8:1662291-1687210 | PREDICTED: uncharacterized protein LOC101496163 | 2 |
| Cat-miR172b | Ca3:30142084-30145590 | Uncharacterized protein | 3 |
| Cat-miR172b | Ca6:62515654-62518257 | PREDICTED: uncharacterized protein LOC101499502 | 4 |
| Cat-miR172c-1 | Ca5:12802656-12809975 | PREDICTED: chromo domain protein LHP1-like | 3 |
| Cat-miR172c-1 | Ca3:36097598-36100393 | Uncharacterized protein | 4 |
| Cat-miR172c-2 | Ca3:40632160-40634715 | PREDICTED: ethylene-responsive transcription factor RAP2-7-like | 2 |
| Cat-miR172c-2 | Ca4:39183200-39186576 | Uncharacterized protein | 2 |
| Cat-miR2111-1 | Ca1:2185737-2193039 | PREDICTED: allantoinase isoform X1 | 2 |
| Cat-miR2111-2 | Ca7:53850512-53852814 | RAB GTPase-like protein A2D | 2 |
| Cat-miR2111-2 | Ca1:16524046-16525988 | Uncharacterized protein | 2 |
| Cat-miR2111-4 | Ca6:15457475-15467723 | Peptidase M1 family aminopeptidase N | 4 |
| Cat-miR2111-5 | Scaffold0628:491140-497352 | PREDICTED: protein argonaute 1 | 2 |
| Cat-miR2111-5 | Ca7:5030078-5036364 | Nucleotide-diphospho-sugar transferase family protein | 2 |
| Cat-miR2111-5 | Ca6:29680283-29685859 | PREDICTED: ubiquitin carboxyl-terminal hydrolase 24 | 2 |
| Cat-miR2111-5 | Ca6:60920159-60924027 | PREDICTED: probableucleolar protein 5-1 | 2 |
| Cat-miR2118 | Ca6:14510327-14515093 | PREDICTED: trehalase isoform X2 | 4 |
| Cat-miR2118 | Ca5:6247007-6251330 | PREDICTED: beta-(1,2)-xylosyltransferase | 2 |
| Cat-miR2118 | Ca8:3556410-3556976 | PREDICTED: casein kinase II subunit alpha-like isoform X1 | 2 |
| Cat-miR2118 | Ca4:2514238-2517791 | Uncharacterized protein | 2 |
| Cat-miR2118 | Ca3:40822769-40828052 | DnaJ heat shock amino-terminal domain protein | 2 |
| Cat-miR2118 | Ca6:14607760-14612073 | Protein kish | 2 |
| Cat-miR319a | Ca7:54021814-54027912 | PREDICTED: potassium transporter 5 | 2 |

| miR | Target | Target Annotation | Category |
|---|---|---|---|
| Cat-miR319a | Ca1:40974393-40977412 | MOB kinase activator-like 1 | 2 |
| Cat-miR319a | Ca1:4115514-4408222 | PREDICTED: DNA-directed RNA polymerase I subunit RPA2-like | 4 |
| Cat-miR319a | Ca7:3592402-3593184 | PREDICTED: cold shock domain-containing protein 3 | 4 |
| Cat-miR319b | Ca4:22037567-22044080 | PREDICTED: uncharacterized protein LOC101497525 | 2 |
| Cat-miR319b | Ca6:667919-669328 | PREDICTED: transcription factor TCP2-like | 2 |
| Cat-miR319b | Ca7:8080891-8082864 | PREDICTED: transcription factor MYC2-like | 2 |
| Cat-miR319b | Ca5:14983734-14987921 | PREDICTED: DNA-3-methyladenine glycosylase 1 | 2 |
| Cat-miR319b | Ca1:11611873-11616950 | Cell division cycle protein 48 homolog | 2 |
| Cat-miR319b | Ca3:63003817-63016414 | PREDICTED: mediator of RNA polymerase II transcription subunit 21 | 2 |
| Cat-miR319b | Scaffold0628:168494-177214 | PREDICTED: protein NRDE2 homolog isoform X2 | 2 |
| Cat-miR319b | Scaffold5165:362220-376629 | Glycyl-tRNA synthetase alpha chain/beta chain | 2 |
| Cat-miR319b | Ca2:19464838-19470581 | PREDICTED: putative disease resistance protein At3g14460 isoform X2 | 2 |
| Cat-miR319b | Ca4:51501485-51510833 | L-galactono-1,4-lactone dehydrogenase | 2 |
| Cat-miR319b | Ca5:13568159-13589577 | Phosphatidylinositol 4-kinase alpha | 2 |
| Cat-miR319b | Ca1:13639769-13651895 | Protein DETOXIFICATION | 2 |
| Cat-miR319b | Ca7:45001932-45016449 | PREDICTED: beta-glucosidase 46-like | 2 |
| Cat-miR319b | Ca7:24743128-24747804 | PREDICTED: transcription factor GAMYB-like | 4 |
| Cat-miR319c-3 | Ca6:18805836-18811743 | CRAL/TRIO domain protein | 2 |
| Cat-miR319c-3 | Ca6:63845031-63848050 | Microfibrillar-associated-like protein | 2 |
| Cat-miR319d | Ca7:3592402-3593184 | PREDICTED: cold shock domain-containing protein 3 | 2 |
| Cat-miR319d | Ca4:58128028-58131242 | PREDICTED: uncharacterized protein LOC101498745 | 4 |
| Cat-miR319d | Ca1:48624444-48627841 | PREDICTED: transcription factor TCP4-like | 2 |
| Cat-miR319d | Ca3:50903102-50906853 | PREDICTED: BTB/POZ domain-containing protein POB1-like | 2 |
| Cat-miR319d | Ca8:7856617-7859608 | Uncharacterized protein | 2 |
| Cat-miR319d | Ca2:19401208-19405683 | PREDICTED: crt homolog 1 | 2 |
| Cat-miR319d | Ca5:28788212-28793732 | PREDICTED: probable inactive receptor kinase At5g10020 | 2 |
| Cat-miR319d | Ca8:2287663-2305919 | PREDICTED: chaperone protein dnaJ 13-like | 2 |
| Cat-miR319d | Ca3:39148260-39153665 | PREDICTED: peptidyl-prolyl cis-trans isomerase CYP63 | 2 |
| Cat-miR390a-1 | Ca5:6598812-6601730 | Heat shock protein 90-2 | 3 |
| Cat-miR390a-1 | Ca3:50828831-50831109 | PREDICTED: U-box domain-containing protein 9-like | 2 |
| Cat-miR390a-1 | Ca4:57986020-57991257 | PREDICTED: uncharacterized protein LOC101513058 | 2 |
| Cat-miR390a-1 | Ca7:22514430-22592040 | PREDICTED: uncharacterized protein LOC106773082 | 2 |
| Cat-miR390a-1 | Ca8:13404600-13646584 | PREDICTED: arginine--tRNA ligase, cytoplasmic-like | 2 |
| Cat-miR390a-1 | Ca6:64176250-64177894 | PREDICTED: F-box protein At1g67340-like | 2 |
| Cat-miR390a-2 | Ca4:18273371-18277558 | Pti1-like kinase | 2 |
| Cat-miR390a-2 | Ca1:370424-371986 | PREDICTED: transcription factor TCP8 | 2 |
| Cat-miR390a-2 | Ca4:16407430-16414461 | PREDICTED: protein indeterminate-domain 5, chloroplastic | 2 |
| Cat-miR390b | Ca4:13988337-13991009 | Tubulin alpha-6 chain | 2 |
| Cat-miR390b | Ca6:50406218-50412544 | PREDICTED: uncharacterized protein LOC101496135 | 2 |
| Cat-miR390b | Ca1:16975697-16977547 | PREDICTED: hybrid signal transduction histidine kinase M | 2 |
| Cat-miR390b | Ca1:14414399-14419065 | Vacuolar protein sorting-associated protein 4 | 2 |
| Cat-miR390b | Ca4:11360319-11362309 | 0 | 2 |
| Cat-miR390b | Ca4:3480880-3486645 | PREDICTED: AT-hook motifuclear-localized protein 6-like | 2 |
| Cat-miR390b | Ca6:1424592-1426150 | PREDICTED: peptidyl-prolyl cis-trans isomerase FKBP13, chloroplastic | 4 |
| Cat-miR390b | Ca5:24797041-24800711 | Uncharacterized protein | 4 |
| Cat-miR394-1 | Ca4:18391221-18395966 | PREDICTED: extensin-2 | 2 |
| Cat-miR394-1 | Ca4:18391306-18393071 | 0 | 2 |
| Cat-miR394-1 | Ca4:18391221-18395966 | PREDICTED: extensin-2 | 2 |
| Cat-miR394-1 | Ca4:18391306-18393071 | 0 | 2 |
| Cat-miR394-1 | Ca3:49651606-49656657 | PREDICTED: uncharacterized serine-rich protein C215.13-like | 2 |
| Cat-miR394-1 | Ca4:58809489-58811737 | PREDICTED: UPF0202 protein At1g10490 | 2 |
| Cat-miR394-1 | Ca4:19076721-19080079 | PREDICTED: protein IQ-DOMAIN 1 | 2 |
| Cat-miR394-1 | Ca3:40235921-40238827 | Heat shock protein 70 | 2 |
| Cat-miR394-1 | Ca1:39238901-39268403 | Elongation factor 2 | 2 |
| Cat-miR394-2 | Ca4:7710505-7714878 | PREDICTED: zinc finger CCCH domain-containing protein 32 isoform X1 | 0 |

| miR | Target | Target Annotation | Category |
|---|---|---|---|
| Cat-miR394-2 | Ca1:17600793-17602199 | PREDICTED: protein FAM179B isoform X1 | 2 |
| Cat-miR394-2 | Ca4:18391306-18393071 | 0 | 2 |
| Cat-miR394-2 | Ca4:18391221-18395966 | PREDICTED: extensin-2 | 2 |
| Cat-miR394-2 | Ca6:15050185-15052289 | PREDICTED: loricrin isoform X2 | 2 |
| Cat-miR394-2 | Ca6:2927679-2933079 | PREDICTED: polyadenylate-binding protein 2-like isoform X1 | 2 |
| Cat-miR394-2 | Ca6:12933370-12936084 | Flavin-binding kelch repeat F-box protein, putative | 2 |
| Cat-miR394-2 | Ca7:50532289-50533892 | PREDICTED: blue copper protein-like | 2 |
| Cat-miR394-2 | Ca6:5319823-5353332 | PREDICTED: isoflavone 4'''-O-methyltransferase | 2 |
| Cat-miR396a | Ca3:46653612-46657159 | PREDICTED: growth-regulating factor 4 | 4 |
| Cat-miR396b | Ca7:10338976-10345911 | PREDICTED: uncharacterized protein LOC101503342 isoform X1 | 2 |
| Cat-miR396c | Ca5:21800787-21814765 | Pantothenate kinase | 2 |
| Cat-miR396c | Ca6:23706025-23710160 | PREDICTED: protein EDS1L-like | 2 |
| Cat-miR396c | Ca1:46232634-46236488 | PREDICTED: palmitoyl-acyl carrier protein thioesterase, chloroplastic-like | 2 |
| Cat-miR396c | Ca1:7022482-7033437 | PREDICTED: protein ROOT HAIR DEFECTIVE 3 homolog 2 | 4 |
| Cat-miR398a-5p | Ca4:33186248-33196347 | P-loopucleoside triphosphate hydrolase superfamily protein | 2 |
| Cat-miR398a-5p | Ca3:49574046-49616098 | PREDICTED: ATP-dependent 6-phosphofructokinase 6-like | 2 |
| Cat-miR398a-5p | Ca7:53215261-53219795 | #N/A | 2 |
| Cat-miR398a-5p | Ca8:2165057-2166028 | #N/A | 2 |
| Cat-miR398a-5p | Ca5:27876978-27881480 | #N/A | 2 |
| Cat-miR398a-5p | Ca4:724505-727359 | S-(hydroxymethyl)glutathione dehydrogenase | 2 |
| Cat-miR398a-5p | Ca7:19818791-19826012 | #N/A | 2 |
| Cat-miR398b-1 | Ca6:47621472-47626541 | PREDICTED: cytochrome c oxidase subunit 5b-2, mitochondrial-like | 2 |
| Cat-miR398b-1 | Ca7:54935056-54940470 | PREDICTED: FAM10 family protein At4g22670 | 2 |
| Cat-miR398b-2 | Ca1:46430146-46432252 | PREDICTED: cucumber peeling cupredoxin-like | 3 |
| Cat-miR398b-2 | Ca4:393962-400992 | Arginine biosynthesis bifunctional protein ArgJ, chloroplastic | 2 |
| Cat-miR399-1 | Ca7:3901070-3905879 | Serine/threonine phosphatase family, 2C domain protein | 2 |
| Cat-miR408 | Ca6:62198401-62199252 | Basic blue copper protein | 3 |
| Cat-miR408 | Ca8:14689513-14691283 | 0 | 2 |
| Cat-miR408 | Ca1:11878534-11882430 | PREDICTED: double-stranded RNA-binding protein 2-like | 2 |
| Cat-miR408 | Ca7:53661306-53664808 | PREDICTED: transcription factor MYB1R1 | 2 |
| Cat-miR408 | Ca3:49039531-49042337 | PREDICTED: pectin acetylesterase 12-like | 2 |
| Cat-miR408 | Ca7:45940810-45951912 | Adenosylmethionine-8-amino-7-oxononanoate transaminase | 2 |
| Cat-miR408 | Ca1:10230563-10235575 | Casein kinase I-like protein | 2 |
| Cat-miR408 | Ca4:6535079-6538929 | PREDICTED: pectin acetylesterase 6-like | 2 |
| Cat-miR408 | Ca7:40952007-40958748 | PREDICTED: vacuolar protein sorting-associated protein 36 | 2 |
| Cat-miR408 | Ca7:46907215-46910735 | Transducin/WD-like repeat-protein | 4 |
| Cat-miR408 | Ca5:20132600-20133510 | PREDICTED: blue copper protein-like | 4 |
| Cat-miR408 | Ca5:62187399-62223614 | PREDICTED: U-box domain-containing protein 35-like isoform X1 | 4 |
| Cat-miR482 | Ca8:13928027-13929535 | PREDICTED: ectonucleotide pyrophosphatase/phosphodiesterase family member 1-like | 2 |
| Cat-miR482 | Ca3:62098023-62101425 | Kinase 1B | 2 |
| Cat-miR482 | Ca2:20768105-20772404 | PREDICTED: calcium-dependent protein kinase 2-like | 2 |
| Cat-miR482 | Ca4:11360319-11362309 | 0 | 2 |
| Cat-miR5213 | Ca6:65110104-65115782 | RNA-binding (RRM/RBD/RNP motif) family protein | 2 |
| Cat-miR5213 | Ca8:9664899-9668014 | Eukaryotic translation initiation factor 6 | 2 |
| Cat-miR5213 | Ca5:28688007-28692668 | PREDICTED: alpha-galactosidase 3 | 2 |
| Cat-NovmiR1 | Ca8:13720618-13725363 | ATP-dependent 6-phosphofructokinase | 3 |
| Cat-NovmiR1 | Scaffold4730:239147-242260 | PREDICTED: annexin D5-like | 4 |
| Cat-NovmiR1 | Ca4:18878310-18881682 | Adenylate kinase | 2 |
| Cat-NovmiR1 | Ca5:11919959-11932915 | Uncharacterized protein | 2 |
| Cat-NovmiR1 | Ca6:60583788-60586026 | PREDICTED: BTB/POZ domain-containing protein At3g05675-like | 2 |
| Cat-NovmiR1 | Ca1:493586-501429 | Importin beta-4 | 2 |
| Cat-NovmiR1 | Ca3:32369220-32372317 | PREDICTED: CBS domain-containing protein CBSX6 | 2 |

| miR | Target | Target Annotation | Category |
|---|---|---|---|
| Cat-NovmiR1 | Ca1:13531450-13535357 | Uncharacterized protein | 2 |
| Cat-NovmiR1 | Ca1:34617439-34624233 | PREDICTED: uncharacterized protein LOC101493362 | 2 |
| Cat-NovmiR1 | Ca8:8648978-8654820 | PREDICTED: protein SPT2 homolog | 2 |
| Cat-NovmiR1 | Ca7:45796269-45803660 | PREDICTED: diacylglycerol kinase 3 | 2 |
| Cat-NovmiR1 | Ca4:35821826-35825849 | Nucleotide/sugar transporter family protein | 2 |
| Cat-NovmiR1 | Ca3:1759826-1768541 | Uncharacterized protein | 2 |
| Cat-NovmiR1 | Ca3:40611230-40613412 | P-loopucleoside triphosphate hydrolase superfamily protein | 3 |
| Cat-NovmiR1 | Ca3:8578788-8589977 | PREDICTED: phosphatidylinositol/phosphatidylcholine transfer protein SFH9 isoform X1 | 4 |
| Cat-NovmiR2 | Ca3:61410220-61413086 | PREDICTED: desumoylating isopeptidase 1 | 2 |
| Cat-NovmiR2 | Ca5:3768745-3777872 | Serine/threonine-protein phosphatase 5 | 2 |
| Cat-NovmiR3 | Ca7:2992649-2996090 | PREDICTED: uncharacterized protein LOC101505599 | 2 |
| Cat-NovmiR3 | Ca1:3541926-3547106 | MACPF domain protein | 4 |
| Cat-NovmiR6 | Ca4:53787429-53788729 | PREDICTED: uncharacterized protein LOC101491982 | 2 |
| Cat-NovmiR6 | Ca2:21606499-21613803 | PREDICTED: uncharacterized protein LOC101489896 isoform X2 | 2 |
| Cat-NovmiR6 | Ca7:2484485-2493757 | PREDICTED: proline-rich receptor-like protein kinase PERK4 isoform X1 | 2 |
| Cat-NovmiR6 | Ca8:5836455-5845794 | Vacuolar protein sorting-associated protein 35 | 2 |
| Cat-NovmiR6 | Ca6:41242052-41245088 | Expansin-like protein B1 | 2 |
| Cat-NovmiR6 | Ca6:17729177-17738485 | ABC transporter-like protein | 4 |
| Cat-NovmiR7 | Ca6:11818451-11821061 | #N/A | 2 |
| Cat-NovmiR7 | Ca4:17179261-17180721 | #N/A | 2 |
| Cat-NovmiR7 | Ca7:50113542-50115041 | PREDICTED:ucleolin-like | 2 |
| Cat-NovmiR7 | Ca4:57258538-57264573 | #N/A | 2 |
| cat-miR394 | Ca1_35237403-35238547 | Histidine phosphotransferase | 4 |
| **Nodule** | | | |
| Cat-miR156b | Ca3:38170417-38173423 | PREDICTED: squamosa promoter-binding-like protein 9 | 0 |
| Cat-miR319b | Ca1:3936260-3937645 | PREDICTED: transcription factor TCP2 | 0 |
| Cat-miR1507 | Ca4:36280496-36289841 | #N/A | 2 |
| Cat-miR1507 | Ca5:9916501-9923271 | #N/A | 2 |
| Cat-miR1507 | Ca6:60399878-60402185 | #N/A | 2 |
| Cat-miR1507 | Ca7:45954575-45961528 | Calmodulin-binding protein | 2 |
| Cat-miR1507 | Ca4:16705196-16718676 | #N/A | 2 |
| Cat-miR1507 | Ca6:47027344-47030772 | #N/A | 2 |
| Cat-miR1507 | Ca6:47027344-47030772 | #N/A | 2 |
| Cat-miR1507 | Ca2:3102138-3115063 | PREDICTED: serine/threonine-protein kinase dst1 | 2 |
| Cat-miR1507 | Ca4:27081334-27086120 | #N/A | 2 |
| Cat-miR1507 | Ca4:54037436-54039424 | #N/A | 2 |
| Cat-miR1507 | Ca6:37118141-37119983 | #N/A | 2 |
| Cat-miR1507 | Scaffold0583:8101-11346 | #N/A | 4 |
| Cat-miR1509 | Ca4:848278-850064 | PREDICTED: protein JASON | 0 |
| Cat-miR1509 | Ca8:15984324-15987997 | PREDICTED: protein SGT1 homolog | 2 |
| Cat-miR1509 | Ca4:4297172-4301616 | PREDICTED: U11/U12 smalluclear ribonucleoprotein 48 kDa protein | 2 |
| Cat-miR1509 | Ca3:50200282-50202136 | PREDICTED: uncharacterized protein LOC101492518 | 2 |
| Cat-miR1509 | Ca6:885910-888498 | PREDICTED: adenylate kinase isoenzyme 6 homolog | 2 |
| Cat-miR1509 | Ca1:12076587-12079921 | PREDICTED: adenylate kinase isoenzyme 6 homolog | 2 |
| Cat-miR1511 | Ca4:37659490-37665064 | PREDICTED: pentatricopeptide repeat-containing protein At3g22470, mitochondrial-like | 4 |
| Cat-miR156a-1 | Ca2:17579241-17596608 | PREDICTED: probable leucine-rich repeat receptor-like serine/threonine-protein kinase At3g14840 | 2 |
| Cat-miR156a-1 | Ca6:49578549-49588910 | PREDICTED: protein ROS1-like | 2 |
| Cat-miR156a-1 | Ca3:20743615-20748677 | PREDICTED: transcription initiation factor TFIID subunit 11 isoform X1 | 2 |
| Cat-miR156a-2 | Ca7:47840062-47910402 | Calcium-dependent lipid-binding-like protein | 2 |
| Cat-miR156a-2 | Ca8:10742166-10752979 | AGC family Serine/Threonine kinase family protein | 2 |
| Cat-miR156a-2 | Ca4:14419435-14421848 | PREDICTED: methylthioribose kinase-like isoform X1 | 2 |
| Cat-miR156a-2 | Ca3:40920822-40927671 | PREDICTED: ethylene-insensitive protein 2 | 4 |
| Cat-miR156a-2 | Ca3:32941385-32944363 | Acyl-CoA thioesterase, putative | 4 |
| Cat-miR156a-3 | Ca4:7628104-7654987 | PREDICTED: uncharacterized protein LOC101497938 | 2 |
| Cat-miR156a-3 | Ca4:28830182-28838574 | PREDICTED: protein EMSY-LIKE 1 | 2 |
| Cat-miR156a-3 | Ca6:14931925-14954984 | PREDICTED: centromere-associated protein E isoform X4 | 2 |

| miR | Target | Target Annotation | Category |
|---|---|---|---|
| Cat-miR156a-3 | Ca7:650930-653206 | PREDICTED: subtilisin-like protease SBT1.7 | 3 |
| Cat-miR156b | Ca7:45249967-45254466 | Cytosol aminopeptidase family protein | 2 |
| Cat-miR156b | Ca3:35158876-35169610 | Importin beta-3, putative | 2 |
| Cat-miR156c-3 | Ca3:32989732-32993382 | Uncharacterized protein | 2 |
| Cat-miR156c-3 | Ca6:20650287-20653460 | PREDICTED: ATPase ASNA1 homolog | 4 |
| Cat-miR156c-4 | Ca1:3241226-3245890 | Uncharacterized protein | 4 |
| Cat-miR156d | Ca2:3794979-3800049 | PREDICTED: LOW QUALITY PROTEIN: squamosa promoter-binding-like protein 6 | 4 |
| Cat-miR156d | Ca8:704873-710512 | PREDICTED: putative 12-oxophytodienoate reductase 11 | 2 |
| Cat-miR156d | Ca8:12188166-12275518 | Plant synaptotagmin | 2 |
| Cat-miR156d | Ca6:5704789-5709534 | PREDICTED: protein IQ-DOMAIN 14 | 2 |
| Cat-miR156d | Ca4:15483996-15488401 | PREDICTED: cysteine--tRNA ligase, cytoplasmic | 2 |
| Cat-miR156d | Ca7:50193177-50199734 | Probable sucrose-phosphate synthase | 2 |
| Cat-miR156d | Ca7:31644338-31648917 | Endoplasmic reticulum-type calcium-transporting ATPase | 2 |
| Cat-miR156d | Ca3:58041312-58055198 | Actin-97 | 2 |
| Cat-miR156d | Ca3:58041312-58055198 | Actin-97 | 2 |
| Cat-miR156d | Ca4:42250207-42251142 | PREDICTED: uncharacterized protein LOC101509491 | 2 |
| Cat-miR156d | Ca1:3908973-3925137 | Uncharacterized protein | 2 |
| Cat-miR156d | Ca6:4306107-4308806 | PREDICTED: armadillo repeat-containing protein 7 | 2 |
| Cat-miR156d | Ca6:20650287-20653460 | PREDICTED: ATPase ASNA1 homolog | 4 |
| Cat-miR156d | Ca1:9464245-9467765 | PREDICTED: uncharacterized protein LOC101506888 | 4 |
| Cat-miR156d | Ca4:3948180-3958105 | Serine/Threonine kinase family protein | 3 |
| Cat-miR156e-1 | Ca1:11776044-11780075 | #N/A | 2 |
| Cat-miR156e-1 | Ca2:19636677-19639510 | #N/A | 2 |
| Cat-miR156e-1 | Ca4:4231487-4243578 | #N/A | 2 |
| Cat-miR156e-1 | Ca5:20823052-20831241 | PREDICTED: telomere length regulation protein TEL2 homolog | 2 |
| Cat-miR156e-2 | Ca5:47774132-47774620 | 0 | 0 |
| Cat-miR156e-2 | Ca8:4552809-4554903 | #N/A | 2 |
| Cat-miR156e-2 | Ca7:51171572-51188309 | #N/A | 2 |
| Cat-miR156e-2 | Ca7:40816965-40822688 | #N/A | 2 |
| Cat-miR156e-2 | Ca5:20201399-20203182 | #N/A | 2 |
| Cat-miR156e-2 | Ca4:39272539-39278624 | #N/A | 2 |
| Cat-miR156e-2 | Ca5:18544804-18546508 | #N/A | 2 |
| Cat-miR156e-2 | Ca6:29033827-29037684 | #N/A | 2 |
| Cat-miR156e-2 | Ca5:17944778-17949876 | Eukaryotic translation initiation factor 3 subunit B | 2 |
| Cat-miR156e-2 | Ca6:1283052-1285929 | #N/A | 2 |
| Cat-miR156f-1 | Ca7:51171572-51188309 | #N/A | 2 |
| Cat-miR156f-1 | Ca1:12701923-12713471 | #N/A | 2 |
| Cat-miR156f-2 | Ca5:20201399-20203182 | #N/A | 3 |
| Cat-miR156f-2 | Ca1:27335753-27339610 | #N/A | 4 |
| Cat-miR156f-2 | Ca4:4145537-4155030 | #N/A | 4 |
| Cat-miR162b | Ca5:27769383-27784627 | PREDICTED: MAG2-interacting protein 2 | 2 |
| Cat-miR162b | Scaffold0592:921963-925549 | Uncharacterized protein | 2 |
| Cat-miR162b | Ca6:6805928-6807310 | PREDICTED: serine carboxypeptidase-like 50 | 2 |
| Cat-miR162b | Ca4:9020986-9025907 | PREDICTED: uncharacterized protein LOC101488857 | 2 |
| Cat-miR162b | Ca2:29208614-29238954 | Uncharacterized protein | 3 |
| Cat-miR162b | Ca7:34044020-34054244 | PREDICTED: vacuolar protein sorting-associated protein 8 homolog | 4 |
| Cat-miR162b | Ca8:6258849-6260126 | PREDICTED: phosphoserine aminotransferase 2, chloroplastic | 4 |
| Cat-miR164a-2 | Ca7:55487062-55491962 | PREDICTED: uncharacterized protein LOC101493304 | 2 |
| Cat-miR164a-2 | Ca6:12063827-12067769 | Uncharacterized protein | 2 |
| Cat-miR164a-2 | Ca7:51381379-51385717 | PREDICTED: protein Dr1 homolog isoform X1 | 2 |
| Cat-miR164a-2 | Scaffold0594:292764-300788 | PREDICTED: cylicin-2-like isoform X1 | 4 |
| Cat-miR164a-3 | Ca1:39628190-39632859 | NAC transcription factor | 3 |
| Cat-miR164a-3 | Scaffold0628:550854-554827 | Eukaryotic peptide chain release factor subunit 1-3 | 2 |
| Cat-miR164b | Ca4:36083975-36091304 | PREDICTED: uncharacterized protein LOC101491241 isoform X1 | 0 |
| Cat-miR164b | Ca4:17963476-17966111 | Serine/threonine phosphatase family, 2C domain protein | 2 |
| Cat-miR164b | Ca8:2571562-2577096 | PREDICTED: GATA transcription factor 26-like | 2 |
| Cat-miR164b | Ca6:63962722-63967929 | PREDICTED: pentatricopeptide repeat-containing protein At5g27270 isoform X1 | 2 |
| Cat-miR164b | Ca6:39786201-39790820 | Splicing factor-like protein | 2 |
| Cat-miR164b | Ca3:35425864-35430610 | PREDICTED: uncharacterized protein LOC101495418 isoform X1 | 2 |
| Cat-miR164b | Ca4:12071293-12077676 | PREDICTED: titin-like | 2 |
| Cat-miR164b | Ca6:12344043-12349655 | Sel1 repeat protein | 2 |
| Cat-miR164b | Ca5:65215171-65217442 | PREDICTED: NAD(P)H-dependent 6""-deoxychalcone synthase | 2 |
| Cat-miR164b | Ca7:50532289-50533892 | PREDICTED: blue copper protein-like | 2 |
| Cat-miR164b | Ca7:50113542-50115041 | PREDICTED:ucleolin-like | 2 |
| Cat-miR164b | Ca6:56585310-56590849 | PREDICTED: zinc finger CCCH domain-containing protein 13-like isoform X1 | 2 |
| Cat-miR164b | Ca2:13066143-13068074 | PREDICTED: exocyst complex component EXO70B1-like | 2 |
| Cat-miR164b | Ca2:9501287-9509470 | Glucose-6-phosphate isomerase | 2 |
| Cat-miR164b | Ca3:39922687-39930067 | ABC transporter-like family-protein | 2 |
| Cat-miR164b | Ca4:15973580-15984001 | PREDICTED: serine/threonine-protein kinase prpf4B | 2 |
| Cat-miR164b | Ca2:32259138-32321077 | PREDICTED: isoprenylcysteine alpha-carbonyl methylesterase ICME | 2 |
| Cat-miR164b | Ca4:18391221-18395966 | PREDICTED: extensin-2 | 2 |
| Cat-miR164b | Ca4:18391306-18393071 | 0 | 2 |
| Cat-miR164b | Ca6:2731521-2734216 | PREDICTED: uncharacterized protein LOC101491587 | 2 |
| Cat-miR164b | Ca2:13485828-13493999 | Translocon at the inner envelope membrane ofs 110 protein | 2 |
| Cat-miR164b | Ca4:9426488-9432327 | Peptidyl-prolyl cis-trans isomerase | 2 |
| Cat-miR164b | Ca4:21055545-21059252 | Aspartate/glutamate/uridylate kinase family protein, putative | 2 |
| Cat-miR164b | Ca4:57945566-57949624 | PREDICTED: transcription factor bHLH140 | 2 |
| Cat-miR164b | Ca4:11360319-11362309 | 0 | 2 |
| Cat-miR164b | Ca7:24096813-24106471 | 1-phosphatidylinositol-3-phosphate 5-kinase | 2 |
| Cat-miR164b | Ca1:6530617-6539298 | PREDICTED: enhancer of mRNA-decapping protein 4-like isoform X1 | 2 |
| Cat-miR164b | Ca5:67434414-67441696 | Protein DETOXIFICATION | 2 |
| Cat-miR164b | Ca6:9677683-9681003 | PREDICTED: protein TRAUCO | 2 |
| Cat-miR164b | Ca6:60737772-60743432 | PREDICTED: putative rRNA methyltransferase | 2 |
| Cat-miR164b | Ca6:52242301-52247990 | PREDICTED: external alternative NAD(P)H-ubiquinone oxidoreductase B1, mitochondrial | 2 |
| Cat-miR164b | Ca3:58667251-58671227 | TPR protein | 2 |
| Cat-miR164b | Ca1:10230563-10235575 | Casein kinase I-like protein | 2 |
| Cat-miR164b | Scaffold0590:52294-60863 | Beta-amyrin synthase | 2 |
| Cat-miR164b | Ca6:13348470-13351975 | PREDICTED: putative N6-adenosine-methyltransferase MT-A70-like | 2 |
| Cat-miR164b | Ca8:10992136-10996143 | Endosomal targeting BRO1-like domain protein | 2 |
| Cat-miR164b | Ca7:53536623-53541682 | GDP-fucose O-fucosyltransferase-like protein | 2 |
| Cat-miR164b | Ca7:38756261-38764073 | PREDICTED: 11-oxo-beta-amyrin 30-oxidase-like | 2 |
| Cat-miR164b | Ca5:15279345-15283738 | PREDICTED: protein odr-4 homolog | 2 |
| Cat-miR164b | Ca6:37475010-37478844 | Structural constituent of cell wall protein, putative | 2 |
| Cat-miR164b | Ca5:13861236-14094850 | Niemann-Pick C1 protein | 2 |
| Cat-miR164b | Ca4:55837218-55848912 | PREDICTED: uncharacterized protein LOC101514792 isoform X1 | 2 |
| Cat-miR164b | Ca6:2873375-2888428 | PREDICTED: LOW QUALITY PROTEIN: outer envelope protein 64, mitochondrial | 2 |
| Cat-miR164b | Ca4:16113172-16120762 | PREDICTED: putative transcription elongation factor SPT5 homolog 1 | 2 |
| Cat-miR164b | Ca7:5955565-5957856 | PREDICTED: histone-lysine N-methyltransferase family member SUVH9-like | 2 |
| Cat-miR164b | Ca1:17810655-17814971 | Uncharacterized protein | 2 |
| Cat-miR164b | Ca6:48150861-48156205 | PREDICTED: uncharacterized protein LOC101504845 | 2 |
| Cat-miR164b | Ca7:2500248-2505805 | U1 smallnuclear ribonucleoprotein 70 kDa protein, putative | 2 |
| Cat-miR164b | Ca7:21948766-21949374 | PREDICTED: dof zinc finger protein DOF1.7-like | 2 |
| Cat-miR164b | Ca4:5699941-5701446 | Ribosomal L22e family protein | 2 |
| Cat-miR164b | Ca6:65079203-65085741 | NADH-ubiquinone oxidoreductase 75 kDa subunit | 2 |
| Cat-miR164b | Ca4:40696173-40697926 | PREDICTED: tankyrase | 2 |
| Cat-miR164b | Ca1:14649450-14663687 | PREDICTED: protein TIC 100 | 4 |
| Cat-miR166a-3 | Ca8:7351797-7363853 | PREDICTED: probable glucan 1,3-alpha-glucosidase isoform X1 | 2 |
| Cat-miR166a-4 | Ca5:29055857-29059960 | PREDICTED: uncharacterized protein LOC101498857 isoform X1 | 2 |
| Cat-miR166a-5 | Ca2:10846941-10850162 | PREDICTED: auxin-responsive protein IAA8-like | 2 |
| Cat-miR166a-5 | Ca4:38849644-38852342 | PREDICTED: glucan endo-1,3-beta-glucosidase 5 | 2 |

| miR | Target | Target Annotation | Category |
|---|---|---|---|
| Cat-miR166a-5 | Ca5:13861236-14094850 | Niemann-Pick C1 protein | 2 |
| Cat-miR166b-2 | Ca7:31017196-31020568 | Uncharacterized protein | 2 |
| Cat-miR166b-2 | Ca3:45664655-45668033 | General substrate transporter | 2 |
| Cat-miR166b-3 | Ca7:54043773-54060442 | PREDICTED: pentatricopeptide repeat-containing protein At4g17616 | 2 |
| Cat-miR166b-3 | Ca5:27571201-27578359 | PP2A regulatory subunit TAP46-like protein | 2 |
| Cat-miR166b-3 | Ca2:30205340-30209864 | PREDICTED: KH domain-containing protein At4g18375 isoform X2 | 2 |
| Cat-miR166b-3 | Ca1:8841298-8852892 | Uncharacterized protein | 2 |
| Cat-miR166b-3 | Scaffold4867:464841-474215 | PREDICTED: U1 smalluclear ribonucleoprotein A | 4 |
| Cat-miR167a-1 | Ca1:6806903-6814467 | PREDICTED: uncharacterized protein LOC101513958 isoform X1 | 2 |
| Cat-miR167a-2 | Ca6:60959806-60962198 | PREDICTED: probable cysteine proteinase A494 | 4 |
| Cat-miR167b-1 | Ca5:38602946-38605542 | PREDICTED: uncharacterized endoplasmic reticulum membrane protein C16E8.02-like | 2 |
| Cat-miR167b-1 | Ca3:39563116-39567033 | PREDICTED: target of Myb protein 1 | 2 |
| Cat-miR167b-1 | Ca2:10364325-10370128 | PREDICTED: E3 ubiquitin-protein ligase UPL1-like | 2 |
| Cat-miR167b-1 | Ca8:9669877-9675673 | T-complex protein 1 subunit gamma | 2 |
| Cat-miR167b-2 | Ca6:63308084-63310409 | Tubulin alpha-5 | 2 |
| Cat-miR167c | Ca8:13808270-13812637 | Uncharacterized protein | 2 |
| Cat-miR167c | Ca5:23552052-23560338 | PREDICTED: uncharacterized protein LOC101498628 | 2 |
| Cat-miR167c | Ca4:10064502-10067509 | PREDICTED: serine carboxypeptidase-like 27 | 2 |
| Cat-miR167c | Ca1:8576418-8591212 | PREDICTED: ABC transporter C family member 13 isoform X1 | 2 |
| Cat-miR167c | Ca4:1184794-1187058 | PREDICTED: uncharacterized protein LOC101503717 | 2 |
| Cat-miR167c | Ca3:35838617-35848681 | #N/A | 2 |
| Cat-miR167c | Scaffold0598:59111-59922 | #N/A | 2 |
| Cat-miR167c | Ca4:15973580-15984001 | PREDICTED: serine/threonine-protein kinase prpf4B | 2 |
| Cat-miR167c | Ca4:2210922-2223908 | #N/A | 2 |
| Cat-miR167c | Ca5:7515538-7519470 | #N/A | 2 |
| Cat-miR167c | Ca6:4545902-4555815 | #N/A | 2 |
| Cat-miR167c | Ca7:49718851-49725133 | #N/A | 2 |
| Cat-miR167c | Ca5:19773897-19787589 | NADPH--cytochrome P450 reductase | 3 |
| Cat-miR167d | Ca6:530903-536562 | Auxin response factor | 2 |
| Cat-miR167d | Ca6:2632042-2638142 | Dynamin-related protein 5B-2 | 2 |
| Cat-miR167d | Ca1:6971933-6975337 | PREDICTED: uncharacterized CRM domain-containing protein At3g25440, chloroplastic | 2 |
| Cat-miR167d | Ca6:11538081-11541016 | PREDICTED: probable WRKY transcription factor 4 isoform X1 | 2 |
| Cat-miR167d | Scaffold0591:353979-356460 | PREDICTED: LOB domain-containing protein 41 | 2 |
| Cat-miR167d | Ca7:50460763-50470321 | Topless-like protein | 2 |
| Cat-miR167d | Ca4:55408065-55412024 | #N/A | 2 |
| Cat-miR167d | Ca7:49182431-49189768 | Acyl-CoA-binding domain protein | 4 |
| Cat-miR168 | Scaffold0628:491140-497352 | PREDICTED: protein argonaute 1 | 3 |
| Cat-miR168 | Ca4:14251981-14254921 | #N/A | 2 |
| Cat-miR168 | Ca7:5926352-5929814 | #N/A | 2 |
| Cat-miR168 | Ca8:43221-56661 | #N/A | 2 |
| Cat-miR168 | Ca1:3409612-3409902 | #N/A | 2 |
| Cat-miR171a | Ca3:58041312-58055198 | Actin-97 | 3 |
| Cat-miR171a | Scaffold0592:786495-787619 | PREDICTED: F-box/kelch-repeat protein At1g23390 | 2 |
| Cat-miR171a | Ca7:47326243-47331530 | Sucrose synthase | 2 |
| Cat-miR171a | Ca1:4577001-4583210 | PREDICTED: squamosa promoter-binding-like protein 7 | 2 |
| Cat-miR171a | Ca7:41378627-41381714 | PREDICTED: MLP-like protein 34 | 2 |
| Cat-miR171a | Ca7:50532289-50533892 | PREDICTED: blue copper protein-like | 2 |
| Cat-miR171a | Ca4:40048975-40055520 | PREDICTED: transcription initiation factor TFIID subunit 12 | 2 |
| Cat-miR171a | Ca7:28588962-28595371 | PREDICTED: probable LRR receptor-like serine/threonine-protein kinase At1g56140 | 2 |
| Cat-miR171a | Ca3:49880383-49888862 | Protein YIPF | 2 |
| Cat-miR171a | Scaffold0539:952-1631 | PREDICTED: protein YIPF1 homolog | 2 |
| Cat-miR171a | Ca4:4832553-4839011 | PREDICTED: transcription factor UNE12-like isoform X1 | 2 |
| Cat-miR171b | Ca1:11929779-11933751 | PREDICTED: protein disulfide isomerase-like 1-4 | 2 |
| Cat-miR171b | Scaffold0592:98690-99652 | Putative mitochondrial dicarboxylate carrier protein | 2 |
| Cat-miR171b | Ca7:51993849-52003844 | PREDICTED: calponin homology domain-containing protein DDB_G0272472-like | 2 |
| Cat-miR171b | Ca1:8874996-8876253 | PREDICTED: protein LURP-one-related 5-like | 2 |

| miR | Target | Target Annotation | Category |
|---|---|---|---|
| Cat-miR171b | Ca5:13250527-13262167 | Kinesin-like protein | 2 |
| Cat-miR171b | Ca6:46546994-46558783 | PREDICTED: malonate--CoA ligase-like | 2 |
| Cat-miR171b | Ca5:60459208-60464708 | PREDICTED: reticulon-like protein B8 | 2 |
| Cat-miR171b | Ca7:21740041-21744452 | Myo-inositol oxygenase | 4 |
| Cat-miR171c | Ca4:56157638-56159879 | PREDICTED: F-box protein At1g47056-like | 0 |
| Cat-miR171c | Ca2:19820125-19824906 | PREDICTED: DEAD-box ATP-dependent RNA helicase 53-like | 2 |
| Cat-miR171c | Ca4:8311024-8318521 | Uncharacterized protein | 2 |
| Cat-miR171c | Ca4:55615011-55616659 | PREDICTED: U-box domain-containing protein 30-like | 2 |
| Cat-miR171c | Ca7:46860844-46873793 | PREDICTED: uncharacterized protein LOC101507127 | 2 |
| Cat-miR171c | Ca2:21771032-21774610 | PREDICTED: cytochrome c oxidase assembly protein COX15 | 2 |
| Cat-miR171c | Ca2:19048160-19050842 | PREDICTED: UBP1-associated protein 2B-like | 2 |
| Cat-miR171d | Ca5:5086418-5090333 | Purple acid phosphatase | 2 |
| Cat-miR171d | Ca4:28563719-28566424 | PREDICTED: uncharacterized protein LOC101514582 | 2 |
| Cat-miR171d | Ca7:9465648-9470859 | Putative uncharacterized protein (Fragment) | 2 |
| Cat-miR171d | Ca6:42127717-42129767 | PREDICTED: peroxidase 3-like | 2 |
| Cat-miR171d | Ca2:16538768-16542277 | Uncharacterized protein | 2 |
| Cat-miR171d | Ca2:22157859-22160693 | Emp24/gp25L/p24 family protein | 2 |
| Cat-miR171d | Ca8:15447349-15448844 | PREDICTED: heat stress transcription factor B-2a-like | 2 |
| Cat-miR171e | Ca5:29055857-29059960 | PREDICTED: uncharacterized protein LOC101498857 isoform X1 | 0 |
| Cat-miR171e | Ca5:9952939-9963653 | Glutathione S-transferase/chloride channel, C-terminal | 2 |
| Cat-miR171e | Ca5:9952939-9963653 | Glutathione S-transferase/chloride channel, C-terminal | 2 |
| Cat-miR171e | Ca6:62350912-62354290 | PREDICTED: magnesium transporter MRS2-4 | 2 |
| Cat-miR171e | Ca6:23768275-23771391 | Putative uncharacterized protein | 2 |
| Cat-miR171e | Ca5:12422256-12435106 | P-loopucleoside triphosphate hydrolase superfamily protein, putative | 2 |
| Cat-miR171f | Ca6:15588397-15595765 | PREDICTED: kinesin KP1 | 2 |
| Cat-miR171f | Ca8:8846185-8853412 | Nodulation receptor kinase | 2 |
| Cat-miR171f | Ca3:36867208-36872512 | Ubiquitin-associated/TS-N domain protein | 2 |
| Cat-miR172a-3p | Ca6:933553-936728 | PREDICTED: myosin-binding protein 3 | 2 |
| Cat-miR172a-3p | Ca8:3898205-3903405 | PREDICTED: CBS domain-containing protein CBSX1, chloroplastic | 2 |
| Cat-miR172a-3p | Ca5:59536233-59538671 | 0 | 2 |
| Cat-miR172a-3p | Ca4:5034002-5059905 | Armadillo/beta-catenin-like repeat protein | 2 |
| Cat-miR172a-3p | Ca4:546780-563816 | CLIP-associated protein | 2 |
| Cat-miR172a-3p | Ca4:7969483-8002552 | DUF3595 family protein (Fragment) | 2 |
| Cat-miR172a-3p | Scaffold0591:750710-753404 | PREDICTED: uncharacterized protein LOC101504964 | 2 |
| Cat-miR172a-3p | Ca4:38186428-38194286 | PREDICTED: basic leucine zipper 9 | 2 |
| Cat-miR172a-3p | Ca2:22044306-22048524 | RNA-binding (RRM/RBD/RNP motif) family protein | 2 |
| Cat-miR172a-3p | Ca2:3398347-3401287 | Alpha-1,4-glucan-protein synthase [UDP-forming] | 2 |
| Cat-miR172a-3p | Ca2:30988839-30993899 | PREDICTED: E3 ubiquitin-protein ligase RGLG2-like | 2 |
| Cat-miR172a-3p | Ca5:8883186-8891249 | PREDICTED: ubiquitin-activating enzyme E1 1-like | 2 |
| Cat-miR172a-3p | Ca4:860992-866920 | PREDICTED: tetratricopeptide repeat protein 7A | 2 |
| Cat-miR172a-3p | Ca5:5749833-5752712 | Uncharacterized protein | 2 |
| Cat-miR172a-3p | Ca6:1707938-1712655 | PREDICTED: 6,7-dimethyl-8-ribityllumazine synthase, chloroplastic | 2 |
| Cat-miR172a-3p | Ca5:48738321-48742258 | SEC14 cytosolic factor-like protein | 2 |
| Cat-miR172a-3p | Ca7:31815224-31835275 | PREDICTED: uncharacterized protein LOC101490119 isoform X1 | 2 |
| Cat-miR172a-3p | Ca1:1750428-1752891 | PREDICTED: protein PLANT CADMIUM RESISTANCE 8-like | 2 |
| Cat-miR172a-3p | Ca3:8527238-8541326 | PREDICTED: chromatin modification-related protein EAF1 B-like isoform X1 | 2 |
| Cat-miR172a-3p | Ca5:26717066-26723528 | ENTH/VHS/GAT family protein | 4 |
| Cat-miR172a-3p | Ca5:170508-173476 | PREDICTED: pre-mRNA-splicing factor cwf23 | 4 |
| Cat-miR172a-3p | Ca6:12220046-12224613 | 1-aminocyclopropane-1-carboxylate deaminase, putative | 4 |
| Cat-miR172a-3p | Ca7:41038401-41040818 | PREDICTED: cyclic dof factor 1-like | 4 |
| Cat-miR172a-5p | Ca1:2726067-2729458 | #N/A | 2 |
| Cat-miR172a-5p | Ca5:24235486-24236588 | #N/A | 2 |
| Cat-miR172a-5p | Scaffold0591:1314927-1319102 | #N/A | 2 |
| Cat-miR172a-5p | Ca3:536368-542955 | #N/A | 2 |
| Cat-miR172a-5p | Ca1:12215227-12219389 | #N/A | 4 |
| Cat-miR172b | Ca3:18827373-18836333 | PREDICTED: RNA-directed DNA methylation 4 | 2 |
| Cat-miR172b | Ca1:19242993-19251306 | Subtilisin-like serine protease | 2 |

| miR | Target | Target Annotation | Category |
| --- | --- | --- | --- |
| Cat-miR172b | Ca7:46399050-46403793 | Subtilisin-like serine protease | 2 |
| Cat-miR172b | Ca8:1662291-1687210 | PREDICTED: uncharacterized protein LOC101496163 | 2 |
| Cat-miR172b | Ca3:30142084-30145590 | Uncharacterized protein | 3 |
| Cat-miR172b | Ca6:62515654-62518257 | PREDICTED: uncharacterized protein LOC101499502 | 4 |
| Cat-miR172c-1 | Ca4:39183200-39186576 | Uncharacterized protein | 2 |
| Cat-miR172c-1 | Ca5:12802656-12809975 | PREDICTED: chromo domain protein LHP1-like | 3 |
| Cat-miR172c-1 | Ca3:36097598-36100393 | Uncharacterized protein | 4 |
| Cat-miR172c-2 | Ca3:40632160-40634715 | PREDICTED: ethylene-responsive transcription factor RAP2-7-like | 2 |
| Cat-miR2111-1 | Scaffold0628:491140-497352 | PREDICTED: protein argonaute 1 | 2 |
| Cat-miR2111-1 | Ca7:5030078-5036364 | Nucleotide-diphospho-sugar transferase family protein | 2 |
| Cat-miR2111-1 | Ca1:2185737-2193039 | PREDICTED: allantoinase isoform X1 | 2 |
| Cat-miR2111-1 | Ca6:60920159-60924027 | PREDICTED: probableucleolar protein 5-1 | 2 |
| Cat-miR2111-1 | Ca6:15457475-15467723 | Peptidase M1 family aminopeptidase N | 4 |
| Cat-miR2111-4 | Scaffold5165:341696-343910 | RAB GTPase-like protein A5D | 2 |
| Cat-miR2111-4 | Ca7:53850512-53852814 | RAB GTPase-like protein A2D | 2 |
| Cat-miR2111-5 | Ca1:16524046-16525988 | Uncharacterized protein | 2 |
| Cat-miR2118 | Ca6:14510327-14515093 | PREDICTED: trehalase isoform X2 | 4 |
| Cat-miR2118 | Ca8:6972263-6974587 | 0 | 2 |
| Cat-miR2118 | Ca5:6247007-6251330 | PREDICTED: beta-(1,2)-xylosyltransferase | 2 |
| Cat-miR2118 | Ca8:3556410-3556976 | PREDICTED: casein kinase II subunit alpha-like isoform X1 | 2 |
| Cat-miR2118 | Ca4:2514238-2517791 | Uncharacterized protein | 2 |
| Cat-miR2118 | Ca3:40822769-40828052 | DnaJ heat shock amino-terminal domain protein | 2 |
| Cat-miR2118 | Ca1:16796865-16799966 | Glucan endo-1,3-beta-glucosidase | 2 |
| Cat-miR2118 | Ca8:6258849-6260126 | PREDICTED: phosphoserine aminotransferase 2, chloroplastic | 2 |
| Cat-miR2118 | Ca6:14607760-14612073 | Protein kish | 2 |
| Cat-miR319a | Ca7:54021814-54027912 | PREDICTED: potassium transporter 5 | 2 |
| Cat-miR319a | Ca1:40974393-40977412 | MOB kinase activator-like 1 | 2 |
| Cat-miR319a | Ca1:4115514-4408222 | PREDICTED: DNA-directed RNA polymerase I subunit RPA2-like | 4 |
| Cat-miR319a | Ca7:3592402-3593184 | PREDICTED: cold shock domain-containing protein 3 | 4 |
| Cat-miR319b | Ca4:22037567-22044080 | PREDICTED: uncharacterized protein LOC101497525 | 2 |
| Cat-miR319b | Ca6:667919-669328 | PREDICTED: transcription factor TCP2-like | 2 |
| Cat-miR319b | Ca7:8080891-8082864 | PREDICTED: transcription factor MYC2-like | 2 |
| Cat-miR319b | Ca5:14983734-14987921 | PREDICTED: DNA-3-methyladenine glycosylase 1 | 2 |
| Cat-miR319b | Scaffold0628:168494-177214 | PREDICTED: protein NRDE2 homolog isoform X2 | 2 |
| Cat-miR319b | Ca3:63003817-63016414 | PREDICTED: mediator of RNA polymerase II transcription subunit 21 | 2 |
| Cat-miR319b | Ca1:11611873-11616950 | Cell division cycle protein 48 homolog | 2 |
| Cat-miR319b | Scaffold5165:362220-376629 | Glycyl-tRNA synthetase alpha chain/beta chain | 2 |
| Cat-miR319b | Ca2:19464838-19470581 | PREDICTED: putative disease resistance protein At3g14460 isoform X2 | 2 |
| Cat-miR319b | Ca4:51501485-51510833 | L-galactono-1,4-lactone dehydrogenase | 2 |
| Cat-miR319b | Ca5:13568159-13589577 | Phosphatidylinositol 4-kinase alpha | 2 |
| Cat-miR319b | Ca1:13639769-13651895 | Protein DETOXIFICATION | 2 |
| Cat-miR319b | Ca7:45001932-45016449 | PREDICTED: beta-glucosidase 46-like | 2 |
| Cat-miR319b | Ca3:49625528-49627551 | PREDICTED: uncharacterized protein LOC101493722 isoform X2 | 4 |
| Cat-miR319b | Ca7:24743128-24747804 | PREDICTED: transcription factor GAMYB-like | 4 |
| Cat-miR319c-1 | Ca6:18805836-18811743 | CRAL/TRIO domain protein | 2 |
| Cat-miR319c-2 | Ca6:63845031-63848050 | Microfibrillar-associated-like protein | 2 |
| Cat-miR319d | Ca7:3592402-3593184 | PREDICTED: cold shock domain-containing protein 3 | 2 |
| Cat-miR319d | Ca4:58128028-58131242 | PREDICTED: uncharacterized protein LOC101498745 | 4 |
| Cat-miR319d | Ca1:48624444-48627841 | PREDICTED: transcription factor TCP4-like | 2 |
| Cat-miR319d | Ca3:50903102-50906853 | PREDICTED: BTB/POZ domain-containing protein POB1-like | 2 |
| Cat-miR319d | Ca8:7856617-7859608 | Uncharacterized protein | 2 |
| Cat-miR319d | Ca2:19401208-19405683 | PREDICTED: crt homolog 1 | 2 |
| Cat-miR319d | Ca5:28788212-28793732 | PREDICTED: probable inactive receptor kinase At5g10020 | 2 |
| Cat-miR319d | Ca8:2287663-2305919 | PREDICTED: chaperone protein dnaJ 13-like | 2 |
| Cat-miR319d | Ca3:39148260-39153665 | PREDICTED: peptidyl-prolyl cis-trans isomerase CYP63 | 2 |
| Cat-miR390a-1 | Ca1:370424-371986 | PREDICTED: transcription factor TCP8 | 2 |
| Cat-miR390a-1 | Ca3:50828831-50831109 | PREDICTED: U-box domain-containing protein 9-like | 2 |

| miR | Target | Target Annotation | Category |
| --- | --- | --- | --- |
| Cat-miR390a-1 | Ca4:57986020-57991257 | PREDICTED: uncharacterized protein LOC101513058 | 2 |
| Cat-miR390a-1 | Ca8:13404600-13646584 | PREDICTED: arginine--tRNA ligase, cytoplasmic-like | 2 |
| Cat-miR390a-2 | Ca4:18273371-18277558 | Pti1-like kinase | 2 |
| Cat-miR390a-2 | Ca5:6598812-6601730 | Heat shock protein 90-2 | 3 |
| Cat-miR390a-2 | Ca4:16407430-16414461 | PREDICTED: protein indeterminate-domain 5, chloroplastic | 2 |
| Cat-miR390a-2 | Ca7:22514430-22592040 | PREDICTED: uncharacterized protein LOC106773082 | 2 |
| Cat-miR390a-2 | Ca6:64176250-64177894 | PREDICTED: F-box protein At1g67340-like | 2 |
| Cat-miR390b | Ca4:13988337-13991009 | Tubulin alpha-6 chain | 2 |
| Cat-miR390b | Ca6:50406218-50412544 | PREDICTED: uncharacterized protein LOC101496135 | 2 |
| Cat-miR390b | Ca1:16975697-16977547 | PREDICTED: hybrid signal transduction histidine kinase M | 2 |
| Cat-miR390b | Ca1:12270401-12287183 | MAP kinase kinase kinase-like protein | 2 |
| Cat-miR390b | Ca1:14414399-14419065 | Vacuolar protein sorting-associated protein 4 | 2 |
| Cat-miR390b | Ca4:11360319-11362309 | 0 | 2 |
| Cat-miR390b | Ca4:3480880-3486645 | PREDICTED: AT-hook motifuclear-localized protein 6-like | 2 |
| Cat-miR390b | Ca6:1424592-1426150 | PREDICTED: peptidyl-prolyl cis-trans isomerase FKBP13, chloroplastic | 4 |
| Cat-miR390b | Ca5:24797041-24800711 | Uncharacterized protein | 4 |
| Cat-miR394-1 | Ca4:18391221-18395966 | PREDICTED: extensin-2 | 2 |
| Cat-miR394-1 | Ca4:18391306-18393071 | 0 | 2 |
| Cat-miR394-1 | Ca4:18391221-18395966 | PREDICTED: extensin-2 | 2 |
| Cat-miR394-1 | Ca4:18391306-18393071 | 0 | 2 |
| Cat-miR394-1 | Ca3:49651606-49656657 | PREDICTED: uncharacterized serine-rich protein C215.13-like | 2 |
| Cat-miR394-1 | Ca6:15050185-15052289 | PREDICTED: loricrin isoform X2 | 2 |
| Cat-miR394-1 | Ca6:2927679-2933079 | PREDICTED: polyadenylate-binding protein 2-like isoform X1 | 2 |
| Cat-miR394-1 | Ca6:12933370-12936084 | Flavin-binding kelch repeat F-box protein, putative | 2 |
| Cat-miR394-1 | Ca1:39238901-39268403 | Elongation factor 2 | 2 |
| Cat-miR394-1 | Ca6:5319823-5353332 | PREDICTED: isoflavone 4'''-O-methyltransferase | 2 |
| Cat-miR394-2 | Ca1:17600793-17602199 | PREDICTED: protein FAM179B isoform X1 | 2 |
| Cat-miR394-2 | Ca4:18391306-18393071 | 0 | 2 |
| Cat-miR394-2 | Ca4:18391221-18395966 | PREDICTED: extensin-2 | 2 |
| Cat-miR394-2 | Ca4:58809489-58811737 | PREDICTED: UPF0202 protein At1g10490 | 2 |
| Cat-miR394-2 | Ca4:28750257-28755572 | PREDICTED: zinc finger CCCH domain-containing protein 44-like | 2 |
| Cat-miR394-2 | Ca4:7710505-7714878 | PREDICTED: zinc finger CCCH domain-containing protein 32 isoform X1 | 2 |
| Cat-miR394-2 | Ca4:19076721-19080079 | PREDICTED: protein IQ-DOMAIN 1 | 2 |
| Cat-miR394-2 | Ca2:32727608-32731518 | Uncharacterized protein | 2 |
| Cat-miR394-2 | Ca3:40235921-40238827 | Heat shock protein 70 | 2 |
| Cat-miR394-2 | Ca2:20250634-20256023 | Transport complex-like protein | 2 |
| Cat-miR394-2 | Ca7:50532289-50533892 | PREDICTED: blue copper protein-like | 2 |
| Cat-miR396b | Ca3:46653612-46657159 | PREDICTED: growth-regulating factor 4 | 4 |
| Cat-miR396b | Ca7:10338976-10345911 | PREDICTED: uncharacterized protein LOC101503342 isoform X1 | 2 |
| Cat-miR396c | Ca5:21800787-21814765 | Pantothenate kinase | 2 |
| Cat-miR396c | Ca6:23706025-23710160 | PREDICTED: protein EDS1L-like | 2 |
| Cat-miR396c | Ca4:55013239-55021508 | Glycine dehydrogenase (decarboxylating), mitochondrial | 2 |
| Cat-miR396c | Ca1:46232634-46236488 | PREDICTED: palmitoyl-acyl carrier protein thioesterase, chloroplastic-like | 2 |
| Cat-miR396c | Ca1:7022482-7033437 | PREDICTED: protein ROOT HAIR DEFECTIVE 3 homolog 2 | 4 |
| Cat-miR398a-3p | Ca7:54676570-54680715 | PREDICTED: copper chaperone for superoxide dismutase, chloroplastic/cytosolic | 3 |
| Cat-miR398a-3p | Ca4:33186248-33196347 | P-loopucleoside triphosphate hydrolase superfamily protein | 2 |
| Cat-miR398a-3p | Ca3:49574046-49616098 | PREDICTED: ATP-dependent 6-phosphofructokinase 6-like | 2 |
| Cat-miR398a-3p | Ca8:2165057-2166028 | #N/A | 2 |
| Cat-miR398a-3p | Ca7:53215261-53219795 | #N/A | 2 |
| Cat-miR398a-3p | Ca5:12802656-12809975 | PREDICTED: chromo domain protein LHP1-like | 2 |
| Cat-miR398a-3p | Ca5:27876978-27881480 | #N/A | 2 |
| Cat-miR398a-3p | Ca4:724505-727359 | S-(hydroxymethyl)glutathione dehydrogenase | 2 |

| miR | Target | Target Annotation | Category |
|---|---|---|---|
| Cat-miR398a-3p | Ca7:19818791-19826012 | #N/A | 2 |
| Cat-miR398b-1 | Ca6:47621472-47626541 | PREDICTED: cytochrome c oxidase subunit 5b-2, mitochondrial-like | 2 |
| Cat-miR398b-1 | Ca7:54935056-54940470 | PREDICTED: FAM10 family protein At4g22670 | 2 |
| Cat-miR398b-1 | Ca4:393962-400992 | Arginine biosynthesis bifunctional protein ArgJ, chloroplastic | 2 |
| Cat-miR398b-2 | Ca1:46430146-46432252 | PREDICTED: cucumber peeling cupredoxin-like | 3 |
| Cat-miR399-1 | Ca7:3901070-3905879 | Serine/threonine phosphatase family, 2C domain protein | 2 |
| Cat-miR399-2 | Ca5:69143850-69174772 | Alpha-N-acetylglucosaminidase family protein | 4 |
| Cat-miR408 | Ca6:62198401-62199252 | Basic blue copper protein | 3 |
| Cat-miR408 | Ca8:14689513-14691283 | 0 | 2 |
| Cat-miR408 | Ca1:11878534-11882430 | PREDICTED: double-stranded RNA-binding protein 2-like | 2 |
| Cat-miR408 | Ca7:53661306-53664808 | PREDICTED: transcription factor MYB1R1 | 2 |
| Cat-miR408 | Ca3:49039531-49042337 | PREDICTED: pectin acetylesterase 12-like | 2 |
| Cat-miR408 | Ca7:45940810-45951912 | Adenosylmethionine-8-amino-7-oxononanoate transaminase | 2 |
| Cat-miR408 | Ca1:10230563-10235575 | Casein kinase I-like protein | 2 |
| Cat-miR408 | Ca4:6535079-6538929 | PREDICTED: pectin acetylesterase 6-like | 2 |
| Cat-miR408 | Ca7:40952007-40958748 | PREDICTED: vacuolar protein sorting-associated protein 36 | 2 |
| Cat-miR408 | Ca7:46907215-46910735 | Transducin/WD-like repeat-protein | 4 |
| Cat-miR408 | Ca5:20132600-20133510 | PREDICTED: blue copper protein-like | 4 |
| Cat-miR408 | Ca5:62187399-62223614 | PREDICTED: U-box domain-containing protein 35-like isoform X1 | 4 |
| Cat-miR482 | Ca8:13928027-13929535 | PREDICTED: ectonucleotide pyrophosphatase/phosphodiesterase family member 1-like | 2 |
| Cat-miR482 | Ca3:62098023-62101425 | Kinase 1B | 2 |
| Cat-miR482 | Ca2:20768105-20772404 | PREDICTED: calcium-dependent protein kinase 2-like | 2 |
| Cat-miR482 | Ca4:11360319-11362309 | 0 | 2 |
| Cat-miR5213 | Ca6:65110104-65115782 | RNA-binding (RRM/RBD/RNP motif) family protein | 2 |
| Cat-miR5213 | Ca8:9664899-9668014 | Eukaryotic translation initiation factor 6 | 2 |
| Cat-miR5213 | Ca5:28688007-28692668 | PREDICTED: alpha-galactosidase 3 | 2 |
| Cat-NovmiR1 | Scaffold4730:239147-242260 | PREDICTED: annexin D5-like | 4 |
| Cat-NovmiR1 | Ca3:40611230-40613412 | P-loopucleoside triphosphate hydrolase superfamily protein | 2 |
| Cat-NovmiR1 | Ca4:18878310-18881682 | Adenylate kinase | 2 |
| Cat-NovmiR1 | Ca5:11919959-11932915 | Uncharacterized protein | 2 |
| Cat-NovmiR1 | Ca4:54393758-54396846 | PREDICTED: E3 ubiquitin-protein ligase AIP2 | 2 |
| Cat-NovmiR1 | Ca6:60583788-60586026 | PREDICTED: BTB/POZ domain-containing protein At3g05675-like | 2 |
| Cat-NovmiR1 | Ca1:493586-501429 | Importin beta-4 | 2 |
| Cat-NovmiR1 | Ca1:13531450-13535357 | Uncharacterized protein | 2 |
| Cat-NovmiR1 | Ca3:32369220-32372317 | PREDICTED: CBS domain-containing protein CBSX6 | 2 |
| Cat-NovmiR1 | Ca7:52636758-52641226 | Hipl2 protein | 2 |
| Cat-NovmiR1 | Ca1:34617439-34624233 | PREDICTED: uncharacterized protein LOC101493362 | 2 |
| Cat-NovmiR1 | Ca8:8648978-8654820 | PREDICTED: protein SPT2 homolog | 2 |
| Cat-NovmiR1 | Ca3:1759826-1768541 | Uncharacterized protein | 2 |
| Cat-NovmiR1 | Ca4:35821826-35825849 | Nucleotide/sugar transporter family protein | 2 |
| Cat-NovmiR1 | Ca3:8578788-8589977 | PREDICTED: phosphatidylinositol/phosphatidylcholine transfer protein SFH9 isoform X1 | 4 |
| Cat-NovmiR2 | Ca7:49490391-49494954 | Amine oxidase | 2 |
| Cat-NovmiR2 | Ca3:61410220-61413086 | PREDICTED: desumoylating isopeptidase 1 | 2 |
| Cat-NovmiR2 | Ca5:3768745-3777872 | Serine/threonine-protein phosphatase 5 | 2 |
| Cat-NovmiR3 | Ca7:2992649-2996090 | PREDICTED: uncharacterized protein LOC101505599 | 2 |
| Cat-NovmiR3 | Ca1:3541926-3547106 | MACPF domain protein | 4 |
| Cat-NovmiR6 | Ca4:53787429-53788729 | PREDICTED: uncharacterized protein LOC101491982 | 2 |
| Cat-NovmiR6 | Ca2:21606499-21613803 | PREDICTED: uncharacterized protein LOC101489896 isoform X2 | 2 |
| Cat-NovmiR6 | Ca8:8512685-8513353 | PREDICTED: uncharacterized protein LOC101498349 | 2 |
| Cat-NovmiR6 | Ca7:2484485-2493757 | PREDICTED: proline-rich receptor-like protein kinase PERK4 isoform X1 | 2 |
| Cat-NovmiR6 | Ca6:41242052-41245088 | Expansin-like protein B1 | 2 |
| Cat-NovmiR6 | Ca8:5836455-5845794 | Vacuolar protein sorting-associated protein 35 | 2 |
| Cat-NovmiR6 | Ca6:17729177-17738485 | ABC transporter-like protein | 4 |
| Cat-NovmiR7 | Ca6:11818451-11821061 | #N/A | 2 |

| miR | Target | Target Annotation | Category |
|---|---|---|---|
| Cat-NovmiR7 | Ca4:17179261-17180721 | #N/A | 2 |
| Cat-NovmiR7 | Ca7:50113542-50115041 | PREDICTED:ucleolin-like | 2 |
| Cat-NovmiR7 | Ca4:57258538-57264573 | #N/A | 2 |
| cat-miR394 | Ca1_35237403-35238547 | Histidine phosphotransferase | 4 |

| | | | |
|---|---|---|---|
| **Root** | | | |
| Cat-miR319d | Ca1:48624444-48627841 | PREDICTED: transcription factor TCP4-like | 1 |
| Cat-miR1507 | Ca4:12620686-12623805 | carboxylesterase 1-like (LOC101490909) | 2 |
| Cat-miR1507 | Ca6:60399878-60402185 | #N/A | 2 |
| Cat-miR1507 | Ca4:56032290-56034688 | #N/A | 2 |
| Cat-miR1509 | Ca4:848278-850064 | PREDICTED: protein JASON | 0 |
| Cat-miR1509 | Ca7:48336342-48337063 | 0 | 2 |
| Cat-miR1509 | Ca4:4297172-4301616 | PREDICTED: U11/U12 smalluclear ribonucleoprotein 48 kDa protein | 2 |
| Cat-miR1509 | Ca3:50200282-50202136 | PREDICTED: uncharacterized protein LOC101492518 | 2 |
| Cat-miR1509 | Ca1:12076587-12079921 | PREDICTED: adenylate kinase isoenzyme 6 homolog | 2 |
| Cat-miR1509 | Ca6:885910-888498 | PREDICTED: adenylate kinase isoenzyme 6 homolog | 2 |
| Cat-miR1511 | Ca5:16064454-16073091 | ER membrane protein complex subunit-like protein | 2 |
| Cat-miR156a-2 | Ca7:47840062-47910402 | Calcium-dependent lipid-binding-like protein | 2 |
| Cat-miR156a-2 | Ca4:28830182-28838574 | PREDICTED: protein EMSY-LIKE 1 | 2 |
| Cat-miR156a-2 | Ca6:49578549-49588910 | PREDICTED: protein ROS1-like | 2 |
| Cat-miR156a-2 | Ca3:32941385-32944363 | Acyl-CoA thioesterase, putative | 2 |
| Cat-miR156a-2 | Ca5:21732087-21737394 | PREDICTED: uncharacterized protein LOC101494695 isoform X1 | 2 |
| Cat-miR156a-3 | Ca8:10742166-10752979 | AGC family Serine/Threonine kinase family protein | 2 |
| Cat-miR156a-3 | Ca4:7628104-7654987 | PREDICTED: uncharacterized protein LOC101497938 | 2 |
| Cat-miR156a-3 | Ca2:17579241-17596608 | PREDICTED: probable leucine-rich repeat receptor-like serine/threonine-protein kinase At3g14840 | 2 |
| Cat-miR156a-3 | Ca3:40920822-40927671 | PREDICTED: ethylene-insensitive protein 2 | 4 |
| Cat-miR156b | Ca7:54854285-54857859 | PREDICTED: uncharacterized protein LOC101499362 | 2 |
| Cat-miR156b | Ca8:2794260-2803059 | Lon protease homolog 2, peroxisomal | 2 |
| Cat-miR156c-2 | Ca8:1066046-1068026 | PREDICTED: uncharacterized protein LOC101490125 | 2 |
| Cat-miR156c-4 | Ca6:20650287-20653460 | PREDICTED: ATPase ASNA1 homolog | 4 |
| Cat-miR156d | Ca2:3794979-3800049 | PREDICTED: LOW QUALITY PROTEIN: squamosa promoter-binding-like protein 6 | 4 |
| Cat-miR156d | Ca6:20650287-20653460 | PREDICTED: ATPase ASNA1 homolog | 2 |
| Cat-miR156d | Ca2:14022348-14025926 | PREDICTED: protein NETWORKED 4A-like | 2 |
| Cat-miR156d | Ca8:12188166-12275518 | Plant synaptotagmin | 2 |
| Cat-miR156d | Ca6:5704789-5709534 | PREDICTED: protein IQ-DOMAIN 14 | 2 |
| Cat-miR156d | Ca4:15483996-15488401 | PREDICTED: cysteine--tRNA ligase, cytoplasmic | 2 |
| Cat-miR156d | Ca6:23638345-23655362 | Serine/threonine-protein phosphatase | 2 |
| Cat-miR156d | Ca7:12558029-12561380 | PREDICTED: uncharacterized protein LOC101498996 | 2 |
| Cat-miR156d | Ca7:50193177-50199734 | Probable sucrose-phosphate synthase | 2 |
| Cat-miR156d | Ca3:58041312-58055198 | Actin-97 | 2 |
| Cat-miR156d | Ca4:41546916-41550573 | PREDICTED: high mobility group B protein 6-like | 2 |
| Cat-miR156d | Ca6:63314527-63322921 | Uncharacterized protein | 2 |
| Cat-miR156d | Ca4:42250207-42251142 | PREDICTED: uncharacterized protein LOC101509491 | 2 |
| Cat-miR156d | Ca1:3908973-3925137 | Uncharacterized protein | 2 |
| Cat-miR156d | Ca5:11061295-11065235 | PREDICTED: DEAD-box ATP-dependent RNA helicase 18 | 2 |
| Cat-miR156d | Ca4:54328034-54332303 | PREDICTED: probable protein phosphatase 2C 59 | 2 |
| Cat-miR156d | Ca5:45301554-45334438 | PREDICTED: formin-like protein 20 isoform X1 | 4 |
| Cat-miR156d | Ca8:704873-710512 | PREDICTED: putative 12-oxophytodienoate reductase 11 | 3 |
| Cat-miR156e-1 | Ca2:19636677-19639510 | #N/A | 2 |
| Cat-miR156e-1 | Ca5:18544804-18546508 | #N/A | 2 |
| Cat-miR156e-1 | Ca4:4231487-4243578 | #N/A | 2 |
| Cat-miR156e-1 | Ca6:29033827-29037684 | #N/A | 2 |
| Cat-miR156e-1 | Ca5:17944778-17949876 | Eukaryotic translation initiation factor 3 subunit B | 2 |
| Cat-miR156e-2 | Ca8:4552809-4554903 | #N/A | 2 |
| Cat-miR156e-2 | Ca7:51171572-51188309 | #N/A | 2 |
| Cat-miR156e-2 | Ca6:57031141-57042957 | PREDICTED: protein transport protein SEC31 homolog B | 2 |
| Cat-miR156e-2 | Ca1:11776044-11780075 | #N/A | 2 |
| Cat-miR156e-2 | Ca5:20201399-20203182 | #N/A | 2 |
| Cat-miR156e-2 | Ca5:20823052-20831241 | PREDICTED: telomere length regulation protein TEL2 homolog | 2 |
| Cat-miR156f-1 | Ca4:16113172-16120762 | PREDICTED: putative transcription elongation factor | 2 |

| miR | Target | Target Annotation | Category |
|---|---|---|---|
| | | SPT5 homolog 1 | |
| Cat-miR156f-1 | Ca5:20201399-20203182 | #N/A | 3 |
| Cat-miR160-2 | Scaffold5162:112110-115707 | PREDICTED: heme oxygenase 1, chloroplastic-like isoform X1 | 2 |
| Cat-miR162b | Scaffold0592:921963-925549 | Uncharacterized protein | 2 |
| Cat-miR162b | Ca5:27769383-27784627 | PREDICTED: MAG2-interacting protein 2 | 4 |
| Cat-miR162b | Ca4:13381071-13385617 | PREDICTED: primary amine oxidase-like | 2 |
| Cat-miR162b | Ca2:29208614-29238954 | Uncharacterized protein | 3 |
| Cat-miR164a-1 | Ca7:55487062-55491962 | PREDICTED: uncharacterized protein LOC101493304 | 2 |
| Cat-miR164a-2 | Ca1:39628190-39632859 | NAC transcription factor | 3 |
| Cat-miR164a-2 | Ca7:51381379-51385717 | PREDICTED: protein Dr1 homolog isoform X1 | 2 |
| Cat-miR164a-3 | Ca2:10486845-10507678 | PREDICTED: MADS-box transcription factor 23-like isoform X3 | 2 |
| Cat-miR164a-3 | Ca6:9175136-9178533 | Impaired sucrose induction protein, putative | 2 |
| Cat-miR164a-3 | Ca8:12928495-12931984 | PREDICTED: proline-rich receptor-like protein kinase PERK1 | 2 |
| Cat-miR164b | Ca1:9876548-9880271 | ATP-dependent 6-phosphofructokinase | 4 |
| Cat-miR164b | Ca6:15230791-15233212 | 1-aminocyclopropane-1-carboxylate synthase | 3 |
| Cat-miR164b | Ca4:36083975-36091304 | PREDICTED: uncharacterized protein LOC101491241 isoform X1 | 0 |
| Cat-miR164b | Ca6:39786201-39790820 | Splicing factor-like protein | 2 |
| Cat-miR164b | Ca4:12071293-12077676 | PREDICTED: titin-like | 2 |
| Cat-miR164b | Ca5:65215171-65217442 | PREDICTED: NAD(P)H-dependent 6""-deoxychalcone synthase | 2 |
| Cat-miR164b | Ca6:12344043-12349655 | Sel1 repeat protein | 2 |
| Cat-miR164b | Scaffold3523:21891-26350 | PREDICTED: purple acid phosphatase 18-like isoform X1 | 2 |
| Cat-miR164b | Ca4:12071293-12077676 | PREDICTED: titin-like | 2 |
| Cat-miR164b | Ca7:50532289-50533892 | PREDICTED: blue copper protein-like | 2 |
| Cat-miR164b | Ca7:50113542-50115041 | PREDICTED:ucleolin-like | 2 |
| Cat-miR164b | Ca1:27442503-27449511 | PREDICTED: DEAD-box ATP-dependent RNA helicase 51 | 2 |
| Cat-miR164b | Ca2:13066143-13068074 | PREDICTED: exocyst complex component EXO70B1-like | 2 |
| Cat-miR164b | Ca6:7219978-7224962 | Uncharacterized protein | 2 |
| Cat-miR164b | Ca4:15973580-15984001 | PREDICTED: serine/threonine-protein kinase prpf4B | 2 |
| Cat-miR164b | Ca4:18391306-18393071 | 0 | 2 |
| Cat-miR164b | Ca4:18391221-18395966 | PREDICTED: extensin-2 | 2 |
| Cat-miR164b | Ca6:2731521-2734216 | PREDICTED: uncharacterized protein LOC101491587 | 2 |
| Cat-miR164b | Ca4:11360319-11362309 | 0 | 2 |
| Cat-miR164b | Ca1:14649450-14663687 | PREDICTED: protein TIC 100 | 2 |
| Cat-miR164b | Ca4:57482016-57488353 | PREDICTED: auxilin-like protein 1 isoform X1 | 2 |
| Cat-miR164b | Ca1:6530617-6539298 | PREDICTED: enhancer of mRNA-decapping protein 4-like isoform X1 | 2 |
| Cat-miR164b | Ca7:33354244-33359696 | PREDICTED: uncharacterized protein LOC101494421 | 2 |
| Cat-miR164b | Ca5:22508112-22512378 | PREDICTED: flowering time control protein FPA isoform X1 | 2 |
| Cat-miR164b | Ca1:10230563-10235575 | Casein kinase I-like protein | 2 |
| Cat-miR164b | Ca3:58667251-58671227 | TPR protein | 2 |
| Cat-miR164b | Ca3:39736285-39739002 | PREDICTED: rho GDP-dissociation inhibitor 1-like | 2 |
| Cat-miR164b | Scaffold0590:52294-60863 | Beta-amyrin synthase | 2 |
| Cat-miR164b | Ca7:55602200-55605689 | BTB/POZ and TAZ domain protein | 2 |
| Cat-miR164b | Ca8:13921236-13925887 | PREDICTED: thioredoxin-like 2, chloroplastic | 2 |
| Cat-miR164b | Ca1:16066255-16071720 | PREDICTED: probable ADP-ribosylation factor GTPase-activating protein AGD5 isoform X1 | 2 |
| Cat-miR164b | Ca6:14387826-14388800 | PREDICTED: uncharacterized protein LOC101506109 | 2 |
| Cat-miR164b | Ca6:16081295-16085003 | PREDICTED: probable serine/threonine-protein kinase At1g54610 | 2 |
| Cat-miR164b | Ca6:15728085-15732728 | PREDICTED: myosin-11-like | 2 |
| Cat-miR164b | Ca1:9247902-9254902 | Drug resistance transporter-like ABC domain protein | 2 |
| Cat-miR164b | Ca4:16070653-16075435 | PREDICTED: glycine-rich RNA-binding protein RZ1B isoform X1 | 2 |
| Cat-miR164b | Ca6:37475010-37478844 | Structural constituent of cell wall protein, putative | 2 |
| Cat-miR164b | Ca5:13861236-14094850 | Niemann-Pick C1 protein | 2 |
| Cat-miR164b | Ca8:7907545-7912538 | PREDICTED: uncharacterized protein LOC101507671 | 2 |
| Cat-miR164b | Ca7:44786155-44790831 | PREDICTED: aspartic proteinase-like | 2 |
| Cat-miR164b | Ca1:17810655-17814971 | Uncharacterized protein | 2 |

| miR | Target | Target Annotation | Category |
|---|---|---|---|
| Cat-miR164b | Ca6:48150861-48156205 | PREDICTED: uncharacterized protein LOC101504845 | 2 |
| Cat-miR164b | Ca7:21948766-21949374 | PREDICTED: dof zinc finger protein DOF1.7-like | 2 |
| Cat-miR164b | Ca4:40696173-40697926 | PREDICTED: tankyrase | 2 |
| Cat-miR164b | Ca4:5699941-5701446 | Ribosomal L22e family protein | 2 |
| Cat-miR164b | Ca6:7677724-7679338 | PREDICTED: NAC transcription factor 29-like | 4 |
| Cat-miR166a-2 | Ca7:53730249-53735086 | PREDICTED: putative dual specificity protein phosphatase DSP8 | 2 |
| Cat-miR166a-4 | Ca2:10846941-10850162 | PREDICTED: auxin-responsive protein IAA8-like | 2 |
| Cat-miR166a-5 | Ca5:7142950-7143897 | PREDICTED: uncharacterized protein LOC101502143 | 2 |
| Cat-miR166a-5 | Ca1:7078033-7086500 | PREDICTED: 30-kDa cleavage and polyadenylation specificity factor 30 | 2 |
| Cat-miR166a-5 | Ca4:38849644-38852342 | PREDICTED: glucan endo-1,3-beta-glucosidase 5 | 2 |
| Cat-miR166a-5 | Ca8:7351797-7363853 | PREDICTED: probable glucan 1,3-alpha-glucosidase isoform X1 | 2 |
| Cat-miR166b-1 | Ca1:8841298-8852892 | Uncharacterized protein | 2 |
| Cat-miR166b-1 | Scaffold4867:464841-474215 | PREDICTED: U1 smalluclear ribonucleoprotein A | 4 |
| Cat-miR166b-2 | Ca6:7391483-7402641 | Pyruvate kinase | 2 |
| Cat-miR166b-3 | Ca7:48670949-48678394 | Kinesin-like protein | 2 |
| Cat-miR166b-3 | Ca7:31017196-31020568 | Uncharacterized protein | 2 |
| Cat-miR166b-3 | Ca2:36592666-36595595 | PREDICTED: uncharacterized protein LOC101489688 | 2 |
| Cat-miR166b-3 | Ca6:6418145-6421698 | PLC-like phosphodiesterase superfamily protein | 2 |
| Cat-miR166b-3 | Ca5:27571201-27578359 | PP2A regulatory subunit TAP46-like protein | 2 |
| Cat-miR166b-3 | Ca2:30205340-30209864 | PREDICTED: KH domain-containing protein At4g18375 isoform X2 | 2 |
| Cat-miR166b-3 | Ca3:45664655-45668033 | General substrate transporter | 2 |
| Cat-miR167a-1 | Ca3:49651606-49656657 | PREDICTED: uncharacterized serine-rich protein C215.13-like | 2 |
| Cat-miR167a-1 | Ca6:10350394-10367492 | PREDICTED: ENHANCER OF AG-4 protein 2, partial | 2 |
| Cat-miR167a-2 | Ca6:60959806-60962198 | PREDICTED: probable cysteine proteinase A494 | 2 |
| Cat-miR167a-2 | Ca1:6955158-6967502 | PREDICTED: tubulin-folding cofactor E-like | 4 |
| Cat-miR167b-1 | Ca5:38602946-38605542 | PREDICTED: uncharacterized endoplasmic reticulum membrane protein C16E8.02-like | 2 |
| Cat-miR167b-1 | Ca2:10364325-10370128 | PREDICTED: E3 ubiquitin-protein ligase UPL1-like | 4 |
| Cat-miR167b-2 | Ca6:63308084-63310409 | Tubulin alpha-5 | 2 |
| Cat-miR167b-2 | Ca3:51465256-51472649 | PREDICTED: protein STRUBBELIG-RECEPTOR FAMILY 6 | 2 |
| Cat-miR167c | Ca3:47670917-47674120 | PREDICTED: proton-coupled amino acid transporter 3 isoform X1 | 1 |
| Cat-miR167c | Ca1:11493122-11497391 | Uncharacterized protein | 2 |
| Cat-miR167c | Ca6:63719085-63723495 | PREDICTED: probable magnesium transporter NIPA1 | 2 |
| Cat-miR167c | Ca3:35838617-35848681 | #N/A | 2 |
| Cat-miR167c | Ca2:33616403-33617797 | #N/A | 2 |
| Cat-miR167c | Ca4:9357211-9365840 | #N/A | 2 |
| Cat-miR167d | Ca7:51269865-51282369 | PREDICTED: transcription initiation factor TFIID subunit 2 isoform X1 | 2 |
| Cat-miR167d | Ca4:55408065-55412024 | #N/A | 2 |
| Cat-miR168 | Ca6:28321743-28324304 | #N/A | 2 |
| Cat-miR168 | Ca4:16248875-16251793 | #N/A | 2 |
| Cat-miR168 | Ca4:14251981-14254921 | #N/A | 2 |
| Cat-miR168 | Ca4:17219947-17227299 | #N/A | 2 |
| Cat-miR171a | Ca3:58041312-58055198 | Actin-97 | 2 |
| Cat-miR171a | Ca7:47326243-47331530 | Sucrose synthase | 2 |
| Cat-miR171a | Ca6:12344043-12349655 | Sel1 repeat protein | 2 |
| Cat-miR171a | Ca1:4577001-4583210 | PREDICTED: squamosa promoter-binding-like protein 7 | 2 |
| Cat-miR171a | Ca7:41378627-41381714 | PREDICTED: MLP-like protein 34 | 2 |
| Cat-miR171a | Ca7:50532289-50533892 | PREDICTED: blue copper protein-like | 2 |
| Cat-miR171a | Scaffold0539:952-1631 | PREDICTED: protein YIPF1 homolog | 2 |
| Cat-miR171a | Ca3:49880383-49888862 | Protein YIPF | 2 |
| Cat-miR171a | Ca4:4832553-4839011 | PREDICTED: transcription factor UNE12-like isoform X1 | 2 |
| Cat-miR171b | Ca6:33178002-33180911 | PREDICTED: scarecrow-like protein 6 | 2 |
| Cat-miR171b | Ca1:11929779-11933751 | PREDICTED: protein disulfide isomerase-like 1-4 | 2 |
| Cat-miR171b | Ca7:10375560-10378656 | PREDICTED: phosphoribosylformylglycinamidine cyclo-ligase, chloroplastic/mitochondrial-like | 2 |
| Cat-miR171b | Ca2:12720760-12723193 | Uncharacterized protein | 2 |
| Cat-miR171b | Scaffold0592:98690-99652 | Putative mitochondrial dicarboxylate carrier protein | 2 |
| Cat-miR171b | Ca1:8874996-8876253 | PREDICTED: protein LURP-one-related 5-like | 2 |

| miR | Target | Target Annotation | Category |
|---|---|---|---|
| Cat-miR171b | Ca5:13250527-13262167 | Kinesin-like protein | 2 |
| Cat-miR171b | Ca6:5272902-5275964 | PREDICTED: cellulose synthase-like protein G2 | 4 |
| Cat-miR171c | Ca4:56157638-56159879 | PREDICTED: F-box protein At1g47056-like | 2 |
| Cat-miR171c | Ca4:55615011-55616659 | PREDICTED: U-box domain-containing protein 30-like | 2 |
| Cat-miR171c | Ca2:21771032-21774610 | PREDICTED: cytochrome c oxidase assembly protein COX15 | 2 |
| Cat-miR171c | Ca2:19048160-19050842 | PREDICTED: UBP1-associated protein 2B-like | 2 |
| Cat-miR171d | Ca6:9003419-9008245 | PREDICTED: switch 2 | 2 |
| Cat-miR171d | Ca7:9465648-9470859 | Putative uncharacterized protein (Fragment) | 2 |
| Cat-miR171d | Ca2:16538768-16542277 | Uncharacterized protein | 2 |
| Cat-miR171d | Ca2:22157859-22160693 | Emp24/gp25L/p24 family protein | 2 |
| Cat-miR171d | Ca8:15447349-15448844 | PREDICTED: heat stress transcription factor B-2a-like | 2 |
| Cat-miR171e | Ca5:3849973-3852598 | PREDICTED: serine/threonine-protein phosphatase 7 long form homolog isoform X1 | 2 |
| Cat-miR171e | Ca5:9952939-9963653 | Glutathione S-transferase/chloride channel, C-terminal | 2 |
| Cat-miR171e | Ca5:9952939-9963653 | Glutathione S-transferase/chloride channel, C-terminal | 2 |
| Cat-miR171e | Ca1:2889342-2889854 | PREDICTED: pathogenesis-related genes transcriptional activator PTI5-like | 2 |
| Cat-miR171e | Ca6:23768275-23771391 | Putative uncharacterized protein | 2 |
| Cat-miR171e | Ca5:29055857-29059960 | PREDICTED: uncharacterized protein LOC101498857 isoform X1 | 2 |
| Cat-miR171e | Ca2:8028917-8034293 | PREDICTED: thylakoid lumenal 29 kDa protein, chloroplastic isoform X1 | 2 |
| Cat-miR171e | Ca5:12422256-12435106 | P-loopucleoside triphosphate hydrolase superfamily protein, putative | 2 |
| Cat-miR171f | Ca8:8846185-8853412 | Nodulation receptor kinase | 2 |
| Cat-miR171f | Ca3:36867208-36872512 | Ubiquitin-associated/TS-N domain protein | 2 |
| Cat-miR171f | Ca1:38017773-38019712 | PREDICTED: cytochrome P450 CYP736A12-like | 2 |
| Cat-miR171f | Ca7:5299300-5302601 | PREDICTED: uncharacterized protein LOC101503997 | 2 |
| Cat-miR172a-3p | Ca8:3898205-3903405 | PREDICTED: CBS domain-containing protein CBSX1, chloroplastic | 2 |
| Cat-miR172a-3p | Ca4:546780-563816 | CLIP-associated protein | 2 |
| Cat-miR172a-3p | Ca7:39277892-39279802 | PREDICTED: uncharacterized protein LOC101513554 | 2 |
| Cat-miR172a-3p | Ca2:22044306-22048524 | RNA-binding (RRM/RBD/RNP motif) family protein | 2 |
| Cat-miR172a-3p | Ca2:30988839-30993899 | PREDICTED: E3 ubiquitin-protein ligase RGLG2-like | 2 |
| Cat-miR172a-3p | Ca5:8883186-8891249 | PREDICTED: ubiquitin-activating enzyme E1 1-like | 2 |
| Cat-miR172a-3p | Ca1:1750428-1752891 | PREDICTED: protein PLANT CADMIUM RESISTANCE 8-like | 2 |
| Cat-miR172a-3p | Ca5:26717066-26723528 | ENTH/VHS/GAT family protein | 4 |
| Cat-miR172a-3p | Ca4:860992-866920 | PREDICTED: tetratricopeptide repeat protein 7A | 4 |
| Cat-miR172a-5p | Scaffold0591:1314927-1319102 | #N/A | 2 |
| Cat-miR172a-5p | Ca7:51288702-51298435 | #N/A | 2 |
| Cat-miR172a-5p | Ca4:11533056-11536940 | PREDICTED: uncharacterized protein LOC101505330 | 2 |
| Cat-miR172a-5p | Ca5:48732759-48737368 | #N/A | 2 |
| Cat-miR172a-5p | Ca2:5116300-5118157 | #N/A | 3 |
| Cat-miR172b | Ca5:24111828-24114120 | PREDICTED: serine/arginine-rich SC35-like splicing factor SCL30A isoform X2 | 2 |
| Cat-miR172b | Ca1:19242993-19251306 | Subtilisin-like serine protease | 2 |
| Cat-miR172b | Ca4:13468757-13481359 | Long chain acyl-CoA synthetase 7, peroxisomal protein | 3 |
| Cat-miR2111-1 | Ca6:29680283-29685859 | PREDICTED: ubiquitin carboxyl-terminal hydrolase 24 | 2 |
| Cat-miR2111-1 | Ca6:60920159-60924027 | PREDICTED: probableucleolar protein 5-1 | 2 |
| Cat-miR2111-2 | Ca6:9906420-9914289 | Beta-adaptin-like protein | 2 |
| Cat-miR2111-3 | Ca7:53850512-53852814 | RAB GTPase-like protein A2D | 2 |
| Cat-miR2111-3 | Ca1:16524046-16525988 | Uncharacterized protein | 2 |
| Cat-miR2111-4 | Ca4:124071-125393 | Lung seven transmembrane receptor family protein | 2 |
| Cat-miR2111-4 | Ca7:55918747-55931978 | PREDICTED: galactinol--sucrose galactosyltransferase-like | 2 |
| Cat-miR2111-5 | Ca2:26940541-26945259 | Phosphatidylinositol-4-phosphate 5-kinase family protein | 2 |
| Cat-miR2111-5 | Ca2:30769415-30776053 | Disease resistance protein (TIR-NBS-LRR class) | 2 |
| Cat-miR2111-5 | Ca6:15457475-15467723 | Peptidase M1 family aminopeptidase N | 3 |
| Cat-miR2118 | Ca2:6884866-6893616 | Aminoacylase-1 | 2 |
| Cat-miR2118 | Ca5:13031415-13036085 | PREDICTED: translocase of chloroplast 120, chloroplastic | 2 |
| Cat-miR2118 | Ca6:14607760-14612073 | Protein kish | 2 |
| Cat-miR2118 | Ca1:12196867-12208484 | PREDICTED: uncharacterized protein LOC101497329 isoform X1 | 2 |
| Cat-miR319a | Ca7:54021814-54027912 | PREDICTED: potassium transporter 5 | 2 |

| miR | Target | Target Annotation | Category |
|---|---|---|---|
| Cat-miR319b | Ca5:14983734-14987921 | PREDICTED: DNA-3-methyladenine glycosylase 1 | 0 |
| Cat-miR319b | Scaffold5165:362220-376629 | Glycyl-tRNA synthetase alpha chain/beta chain | 1 |
| Cat-miR319b | Ca1:13639769-13651895 | Protein DETOXIFICATION | 2 |
| Cat-miR319b | Ca1:3936260-3937645 | PREDICTED: transcription factor TCP2 | 2 |
| Cat-miR319b | Ca7:8080891-8082864 | PREDICTED: transcription factor MYC2-like | 2 |
| Cat-miR319b | Ca3:63003817-63016414 | PREDICTED: mediator of RNA polymerase II transcription subunit 21 | 2 |
| Cat-miR319b | Ca1:11611873-11616950 | Cell division cycle protein 48 homolog | 2 |
| Cat-miR319b | Ca1:8470026-8476073 | Uncharacterized protein | 2 |
| Cat-miR319b | Ca6:18805836-18811743 | CRAL/TRIO domain protein | 2 |
| Cat-miR319b | Ca6:62917847-62919893 | PREDICTED: disease resistance protein RPS5-like isoform X1 | 2 |
| Cat-miR319b | Ca2:19464838-19470581 | PREDICTED: putative disease resistance protein At3g14460 isoform X2 | 2 |
| Cat-miR319b | Ca4:51501485-51510833 | L-galactono-1,4-lactone dehydrogenase | 2 |
| Cat-miR319b | Ca5:18707299-18713407 | PREDICTED: DNA excision repair protein ERCC-8 | 2 |
| Cat-miR319b | Ca4:27438017-27445232 | Ser/Thr protein kinase | 2 |
| Cat-miR319b | Ca7:45001932-45016449 | PREDICTED: beta-glucosidase 46-like | 2 |
| Cat-miR319b | Ca6:667919-669328 | PREDICTED: transcription factor TCP2-like | 4 |
| Cat-miR319d | Ca7:3592402-3593184 | PREDICTED: cold shock domain-containing protein 3 | 3 |
| Cat-miR319d | Ca8:7856617-7859608 | Uncharacterized protein | 2 |
| Cat-miR319d | Ca2:19401208-19405683 | PREDICTED: crt homolog 1 | 2 |
| Cat-miR319d | Ca5:28788212-28793732 | PREDICTED: probable inactive receptor kinase At5g10020 | 2 |
| Cat-miR319d | Ca8:2287663-2305919 | PREDICTED: chaperone protein dnaJ 13-like | 2 |
| Cat-miR319d | Ca3:39148260-39153665 | PREDICTED: peptidyl-prolyl cis-trans isomerase CYP63 | 2 |
| Cat-miR390a-1 | Ca5:6598812-6601730 | Heat shock protein 90-2 | 3 |
| Cat-miR390a-1 | Ca8:13404600-13646584 | PREDICTED: arginine--tRNA ligase, cytoplasmic-like | 2 |
| Cat-miR390a-2 | Ca4:18273371-18277558 | Pti1-like kinase | 2 |
| Cat-miR390a-2 | Ca1:370424-371986 | PREDICTED: transcription factor TCP8 | 2 |
| Cat-miR390a-2 | Ca1:1022794-1034531 | PREDICTED: calmodulin-binding transcription activator 2-like isoform X1 | 4 |
| Cat-miR390b | Ca4:13988337-13991009 | Tubulin alpha-6 chain | 2 |
| Cat-miR390b | Ca6:50406218-50412544 | PREDICTED: uncharacterized protein LOC101496135 | 2 |
| Cat-miR390b | Ca3:39239546-39243272 | PREDICTED: auxin-induced protein AUX28-like | 2 |
| Cat-miR390b | Ca1:14414399-14419065 | Vacuolar protein sorting-associated protein 4 | 2 |
| Cat-miR390b | Ca4:11360319-11362309 | 0 | 2 |
| Cat-miR394-1 | Ca4:2036643-2049631 | PREDICTED: mitochondrial Rho GTPase 2-like | 1 |
| Cat-miR394-1 | Ca7:45851919-45856242 | Uncharacterized protein | 2 |
| Cat-miR394-1 | Ca4:18391221-18395966 | PREDICTED: extensin-2 | 2 |
| Cat-miR394-1 | Ca4:18391306-18393071 | 0 | 2 |
| Cat-miR394-1 | Ca6:2927679-2933079 | PREDICTED: polyadenylate-binding protein 2-like isoform X1 | 2 |
| Cat-miR394-1 | Ca4:19076721-19080079 | PREDICTED: protein IQ-DOMAIN 1 | 2 |
| Cat-miR394-1 | Ca6:9906420-9914289 | Beta-adaptin-like protein | 2 |
| Cat-miR394-1 | Ca1:39238901-39268403 | Elongation factor 2 | 2 |
| Cat-miR394-1 | Ca1:28133227-28138762 | PREDICTED: plant cysteine oxidase 3 | 2 |
| Cat-miR394-1 | Ca6:5319823-5353332 | PREDICTED: isoflavone 4'''-O-methyltransferase | 2 |
| Cat-miR394-1 | Ca7:50532289-50533892 | PREDICTED: blue copper protein-like | 2 |
| Cat-miR394-2 | Ca4:18391221-18395966 | PREDICTED: extensin-2 | 2 |
| Cat-miR394-2 | Ca4:18391306-18393071 | 0 | 2 |
| Cat-miR394-2 | Ca4:18391306-18393071 | 0 | 2 |
| Cat-miR394-2 | Ca4:18391221-18395966 | PREDICTED: extensin-2 | 2 |
| Cat-miR394-2 | Ca6:15050185-15052289 | PREDICTED: loricrin isoform X2 | 2 |
| Cat-miR394-2 | Ca6:3090273-3109695 | PREDICTED: protein EXECUTER 1, chloroplastic | 2 |
| Cat-miR394-2 | Ca5:64553659-64648791 | PREDICTED: ABC transporter C family member 12-like isoform X1 | 2 |
| Cat-miR394-2 | Ca4:7710505-7714878 | PREDICTED: zinc finger CCCH domain-containing protein 32 isoform X1 | 2 |
| Cat-miR394-2 | Ca3:40235921-40238827 | Heat shock protein 70 | 2 |
| Cat-miR396b | Ca4:50439613-50444040 | PREDICTED: uncharacterized protein LOC101494882 | 2 |
| Cat-miR396c | Ca1:46232634-46236488 | PREDICTED: palmitoyl-acyl carrier protein thioesterase, chloroplastic-like | 2 |
| Cat-miR398a-3p | Ca6:13854617-13859598 | #N/A | 1 |
| Cat-miR398a-3p | Ca7:54676570-54680715 | PREDICTED: copper chaperone for superoxide dismutase, chloroplastic/cytosolic | 2 |

| miR | Target | Target Annotation | Category |
|---|---|---|---|
| Cat-miR398a-3p | Scaffold0605:293309-297939 | PREDICTED: chloroplastic group IIA intron splicing facilitator CRS1, chloroplastic-like | 4 |
| Cat-miR398a-3p | Ca1:47822039-47834873 | PREDICTED: BES1/BZR1 homolog protein 4-like isoform X2 | 2 |
| Cat-miR398a-3p | Ca3:49574046-49616098 | PREDICTED: ATP-dependent 6-phosphofructokinase 6-like | 2 |
| Cat-miR398a-3p | Ca8:2165057-2166028 | #N/A | 2 |
| Cat-miR398a-3p | Ca5:12802656-12809975 | PREDICTED: chromo domain protein LHP1-like | 2 |
| Cat-miR398a-3p | Ca5:27876978-27881480 | #N/A | 2 |
| Cat-miR398a-3p | Ca8:134401-141438 | Presequence protease | 2 |
| Cat-miR398a-3p | Ca4:724505-727359 | S-(hydroxymethyl)glutathione dehydrogenase | 2 |
| Cat-miR398b-1 | Ca1:46430146-46432252 | PREDICTED: cucumber peeling cupredoxin-like | 3 |
| Cat-miR398b-1 | Ca6:47621472-47626541 | PREDICTED: cytochrome c oxidase subunit 5b-2, mitochondrial-like | 2 |
| Cat-miR398b-1 | Ca7:54935056-54940470 | PREDICTED: FAM10 family protein At4g22670 | 2 |
| Cat-miR398b-1 | Ca6:46206754-46208202 | PREDICTED: uncharacterized protein LOC101502166 | 2 |
| Cat-miR408 | Ca6:62198401-62199252 | Basic blue copper protein | 4 |
| Cat-miR408 | Ca8:14689513-14691283 | 0 | 3 |
| Cat-miR408 | Ca7:53661306-53664808 | PREDICTED: transcription factor MYB1R1 | 2 |
| Cat-miR408 | Ca7:36431559-36445178 | PREDICTED: MATE efflux family protein FRD3-like | 2 |
| Cat-miR408 | Ca7:1013377-1016393 | PREDICTED: probable inactive receptor kinase At5g67200 | 2 |
| Cat-miR408 | Ca1:10230563-10235575 | Casein kinase I-like protein | 2 |
| Cat-miR408 | Ca5:21577686-21598302 | PREDICTED: protein MODIFIER OF SNC1 1-like isoform X1 | 2 |
| Cat-miR408 | Ca7:51849609-51852123 | PREDICTED: dnaJ homolog subfamily C member 21 | 2 |
| Cat-miR408 | Ca4:6535079-6538929 | PREDICTED: pectin acetylesterase 6-like | 2 |
| Cat-miR408 | Ca2:15129926-15131760 | PREDICTED: uncharacterized protein LOC101496923 | 2 |
| Cat-miR408 | Ca4:19207783-19209799 | PREDICTED: zinc finger protein JACKDAW-like | 2 |
| Cat-miR482 | Ca3:46000145-46006977 | Putative FAD synthase | 2 |
| Cat-miR482 | Ca5:13031415-13036085 | PREDICTED: translocase of chloroplast 120, chloroplastic | 2 |
| Cat-miR482 | Ca5:8502861-8503460 | Clathrin heavy chain | 2 |
| Cat-miR482 | Ca3:62098023-62101425 | Kinase 1B | 2 |
| Cat-miR482 | Ca5:38041346-38041591 | 0 | 2 |
| Cat-miR482 | Ca6:2331316-2331927 | PREDICTED: polygalacturonase inhibitor 2-like | 2 |
| Cat-miR5213 | Ca4:15336821-15340954 | PREDICTED: uncharacterized protein LOC101515007 | 2 |
| Cat-miR5213 | Ca6:46206754-46208202 | PREDICTED: uncharacterized protein LOC101502166 | 2 |
| Cat-miR5213 | Ca2:30769415-30776053 | Disease resistance protein (TIR-NBS-LRR class) | 4 |

| miR | Target | Target Annotation | Category |
|---|---|---|---|
| Cat-NovmiR1 | Scaffold4730:239147-242260 | PREDICTED: annexin D5-like | 3 |
| Cat-NovmiR1 | Ca3:40611230-40613412 | P-loopucleoside triphosphate hydrolase superfamily protein | 2 |
| Cat-NovmiR1 | Ca6:65693360-65706527 | Uncharacterized protein | 2 |
| Cat-NovmiR1 | Ca5:33770932-33795513 | Uncharacterized protein | 2 |
| Cat-NovmiR1 | Ca1:493586-501429 | Importin beta-4 | 2 |
| Cat-NovmiR1 | Ca1:13531450-13535357 | Uncharacterized protein | 2 |
| Cat-NovmiR1 | Ca3:32369220-32372317 | PREDICTED: CBS domain-containing protein CBSX6 | 2 |
| Cat-NovmiR1 | Ca7:52636758-52641226 | Hipl2 protein | 2 |
| Cat-NovmiR1 | Ca8:8648978-8654820 | PREDICTED: protein SPT2 homolog | 2 |
| Cat-NovmiR1 | Ca4:35821826-35825849 | Nucleotide/sugar transporter family protein | 2 |
| Cat-NovmiR1 | Ca7:32904864-32909252 | NBS-LRR protein | 4 |
| Cat-NovmiR1 | Ca4:18878310-18881682 | Adenylate kinase | 3 |
| Cat-miR398a-5p | Ca4:28039769-28047455 | PREDICTED: uncharacterized protein LOC101509839 | 2 |
| Cat-miR398a-5p | Ca6:181584-183158 | Nucleobase-ascorbate transporter-like protein | 2 |
| Cat-NovmiR2 | Ca7:49490391-49494954 | Amine oxidase | 2 |
| Cat-NovmiR2 | Ca3:61410220-61413086 | PREDICTED: desumoylating isopeptidase 1 | 2 |
| Cat-NovmiR2 | Ca5:3768745-3777872 | Serine/threonine-protein phosphatase 5 | 2 |
| Cat-NovmiR2 | Ca7:52218005-52226931 | Geranylgeranyl hydrogenase | 2 |
| Cat-NovmiR2 | Ca3:36889979-36893511 | PREDICTED: pentatricopeptide repeat-containing protein At1g80270, mitochondrial-like isoform X1 | 2 |
| Cat-NovmiR2 | Ca4:10430998-10446943 | Exocyst complex component sec5 | 2 |
| Cat-NovmiR4 | Ca1:18680661-18684596 | PREDICTED: metal tolerance protein 1-like | 2 |
| Cat-NovmiR6 | Ca7:46620917-46627159 | PREDICTED: protein trichome birefringence-like 5 | 4 |
| Cat-NovmiR6 | Ca3:47272689-47277573 | PREDICTED: protein YLS9 | 3 |
| Cat-NovmiR6 | Ca4:15962868-15967945 | PREDICTED: dihydrolipoyllysine-residue acetyltransferase component 4 of pyruvate dehydrogenase complex, chloroplastic | 2 |
| Cat-NovmiR6 | Ca1:4797479-4808344 | PREDICTED: F-box/kelch-repeat protein At3g23880-like | 2 |
| Cat-NovmiR6 | Ca6:47569512-47574675 | PREDICTED: uncharacterized protein LOC101501411 | 2 |
| Cat-NovmiR6 | Ca8:5836455-5845794 | Vacuolar protein sorting-associated protein 35 | 2 |
| Cat-NovmiR6 | Ca8:12830063-12831541 | PREDICTED: heat stress transcription factor C-1 | 2 |
| Cat-NovmiR7 | Ca3:31814618-31817847 | PREDICTED: O-acyltransferase WSD1-like | 2 |
| Cat-NovmiR7 | Ca6:11818451-11821061 | #N/A | 2 |
| Cat-NovmiR7 | Ca3:46141135-46146044 | #N/A | 2 |
| Cat-NovmiR7 | Ca4:57258538-57264573 | #N/A | 2 |
| Cat-NovmiR7 | Ca7:3009750-3012053 | #N/A | 2 |
| cat-miR394 | Ca1_35237403-35238547 | Histidine phosphotransferase | 3 |
